# Delayed processing of blood samples impairs the accuracy of mRNA-based biomarkers

**DOI:** 10.1101/2022.01.07.475362

**Authors:** Chace Wilson, Nicolas W. Dias, Stefania Pancini, Vitor Mercadante, Fernando H. Biase

**Affiliations:** Department of Animal and Poultry Sciences, Virginia Polytechnic Institute and State University, Blacksburg, VA, USA

**Keywords:** Transcriptome, RNA degradation, Biomarkers, PWBC, blood

## Abstract

**Background:** The transcriptome of peripheral white blood cells (PWBCs) contains valuable physiological information, thus making them a prime biological sample for investigating mRNA-based biomarkers. However, prolonged storage of whole blood samples can alter gene transcript abundance in PWBCs, compromising the results of biomarker discovery. Here, we designed an experiment to interrogate the impacts of delayed processing of whole blood samples on gene transcript abundance in PWBCs. We hypothesized that storing blood samples for 24 hours at 4°C would cause RNA degradation resulting in altered transcriptome profiles.

**Results:** We produced RNA-sequencing data for 30 samples collected from five estrus synchronized heifers (*Bos taurus*). We quantified transcript abundance for 12,414 protein-coding genes in PWBCs. Analysis of parameters of RNA quality revealed no statistically significant differences (P>0.05) between samples collected from the jugular vein and coccygeal vein, as well as among samples processed after one, three, six, or eight hours. However, samples processed after 24 hours of storage had a lower RNA integrity number value (P=0.03) in comparison to those processed after one hour of storage. Next, we analyzed RNA-sequencing data between samples using those processed after one hour of storage as the baseline for comparison. Interestingly, evaluation of 3’/5’ bias revealed no differences between genes with lower transcript abundance in samples stored for 24 hours relative to one hour. In addition, sequencing coverage of transcripts was similar between samples from the 24-hour and one-hour groups. We identified four and 515 genes with differential transcript abundance in samples processed after storage for eight and 24 hours, respectively, relative to samples processed after one hour.

**Conclusions:** The PWBCs respond to prolonged cold storage by increasing genes related to active chromatin compaction which in turn reduces gene transcription. This alteration in transcriptome profiles can impair the accuracy of mRNA-based biomarkers. Therefore, blood samples collected for mRNA-based biomarker discovery should be refrigerated immediately and processed within six hours post sampling.

## BACKGROUND

Blood is a fluid connective tissue that links the entire biological system of an individual, and is composed of plasma and red and white blood cells [1]. Liew and colleagues [1] coined the idea of the “sentinel principle”, whereby blood can harbor molecular indicators of physiological changes in organs, tissues, and cells. Gene transcripts in peripheral white blood cells (PWBCs) are among these molecular indicators. The transcriptome profile of PWBCs is distinct from one individual to another [2, 3], and the profile is as dynamic as the physiological changes that an individual experiences [4-6]. Most importantly, changes in gene expression are detected in the blood relative to several environmental and pathological factors (reviewed by [1, 7]).

Liquid biopsy, including from blood, has emerged as a powerful source of biological material for studying messenger RNA (mRNA) based biomarkers [8]. For instance, mRNAs have been associated with chemo-sensitivity in advanced gastric cancer patients [9], non-small cell lung cancer [10], acute ischemic stroke [11], neuroendocrine tumor [12], prostate cancer [13], hepatocellular carcinoma [14, 15], and Huntington’s disease [16]. In reproductive health, several studies have focused on changes in genes expressed in PWBCs. A cohort of women who were enrolled in the PREVIENI project [17], and identified as infertile, presented altered levels of diverse genes expressed in the PWBCs relative to fertile women [18, 19]. Recently, we have identified several genes that are differentially expressed when contrasting heifers of different pregnancy outcomes (pregnant by AI, pregnant by natural breeding, or not pregnant) [20, 21]. Therefore, the analysis of mRNAs is one possible avenue for the determination of bloodborne molecules that serve as biomarkers of health.

The processing of blood samples for the separation of the buffy coat followed by resuspension in TRIzol Reagent and immediate cryopreservation at −80°C is very efficacious for the extraction of RNAs with high quality and purity [20-23] suitable for producing data by RNA-sequencing [20, 21, 24, 25]. However, when the site of collection cannot be used to process the blood samples, there is a window of time between sampling and collection. Malentacchi et al detected the alteration of transcript abundance of one gene, out of seven tested by polymerase chain reaction, when samples were stored for 24 hours at 4 degrees Celsius (°C) [26]. To date, no study has carried out a systematic interrogation the transcriptome of PWBCs to understand the consequences of storing blood on the alteration of transcript abundance.

Here, we designed an experiment to systematically interrogate the consequences of storing blood samples at different periods up to 24 hours at 4°C on RNA degradation and the transcriptome profile in PWBCs. We hypothesized that the preservation of blood samples for 24 hours at 4°C would lead to RNA degradation, which would result in an alteration in the transcriptome profile of PWBCs.

## RESULTS

### Overview of the experimental design

We collected blood samples from five estrus-synchronized heifers. Ten mL of blood were drawn from the coccygeal vein and five samples of 10 mL were drawn from the jugular vein within seconds among all samplings within each animal. All samplings were performed within 45 minutes. All tubes were preserved on ice and the samples from the jugular vein were randomly assigned to different groups for delayed processing (one, three, six, eight or 24 hours (hr), Fig. 1A). At the assigned time, PWBCs were isolated, pelleted and resuspended in TRIzol™ Reagent for cryopreservation at −80°C. We extracted total RNA from all samples in one batch, assessed quantity and quality and submitted all samples for library preparation prior to freezing (Fig, 1B).

**Fig. 1.**
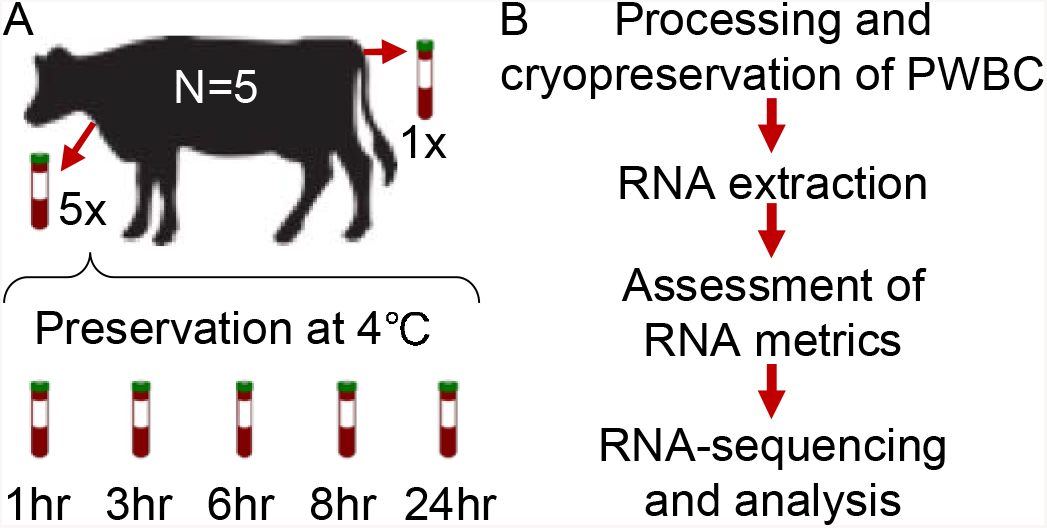
Experimental design and workflow. **A**. Overview of the experimental design with the number of subjects and tubes processed at different times after storage at 4°C. **B**. Workflow of the assays from sample processing to RNA-sequencing.

### Parameters of total RNA based on sampling location and processing delay

We extracted total RNA from 30 samples in one batch, with an average yield of 11.8 μg ±4.5. There was no difference (P>0.05) of the parameters from the samples obtained from the coccygeal versus jugular vein (Table 1, Additional file 1). We then compared the effect of delayed processing on the parameters from samples obtained from the jugular vein. There was no difference (P>0.05) for values of absorbance (A_260_ and A_280_) and the ratio (A_260_/A_280_). However, there was an effect (P=0.03) of the time for delayed processing on the RNA integrity number (RIN). The samples processed 24hr post-collection presented lower RIN relative to the samples processed 1hr post-collection (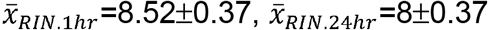, P=0.03, Z-test, Table 1, Additional file 2).

**Table 1.** RNA parameters obtained from peripheral white blood cells.

| Processing Time | RIN | | $A_{260}$ | | $A_{280}$ | | Ratio($A_{260}/A_{280}$) | |
| --- | --- | --- | --- | --- | --- | --- | --- | --- |
| | $\bar{x}$ | $\hat{\sigma}$ | $\bar{x}$ | $\hat{\sigma}$ | $\bar{x}$ | $\hat{\sigma}$ | $\bar{x}$ | $\hat{\sigma}$ |
| **Coccygeal** |  |  |  |  |  |  |  |  |
| 1 hr | 8.48 | 0.16 | 13.27 | 4.02 | 6.85 | 2.03 | 1.93 | 0.02 |
| **Jugular** |  |  |  |  |  |  |  |  |
| 1 hr | 8.52 | 0.37 | 15.99 | 2.75 | 8.28 | 1.37 | 1.93 | 0.02 |
| 3 hr | 8.60 | 0.23 | 17.74 | 6.34 | 9.19 | 3.30 | 1.93 | 0.01 |
| 6 hr | 8.66 | 0.23 | 13.76 | 4.84 | 7.12 | 2.46 | 1.93 | 0.02 |
| 8 hr | 8.34 | 0.33 | 12.40 | 6.85 | 6.40 | 3.49 | 1.91 | 0.06 |
| 24 hr | 8.00 | 0.37 | 14.48 | 2.52 | 7.53 | 1.23 | 1.92 | 0.03 |

### Quality of libraries produced based on sampling location and processing delay

Because the lowest value for RIN was 7.4, which is suitable for transcriptome analysis [27], we proceeded with RNA-sequencing and produced genome-wide transcriptome data for all 30 samples. On average, we produced 29,871,716±3,365,045 pairs of reads per sample (ranging from 21,139,000 to 34,856,707, median 29,941,861, Table 2).

**Table 2.** Metrics for RNA-sequencing data produced from peripheral white blood cells

| Processing | Reads Produced |  | Genes detected before filtering |  | Genes detected after filtering |  | Proportion of reads matching annotation |  | 3'/5' Bias |  |
| --- | --- | --- | --- | --- | --- | --- | --- | --- | --- | --- |
| Time | $\bar{x}$ | $\hat{\sigma}$ | $\bar{x}$ | $\hat{\sigma}$ | $\bar{x}$ | $\hat{\sigma}$ | $\bar{x}$ | $\hat{\sigma}$ | $\bar{x}$ | $\hat{\sigma}$ |
| **Coccygeal** |  |  |  |  |  |  |  |  |  |  |
| 1 hr | 30214519 | 4270887 | 13135 | 167 | 12490 | 51 | 0.56 | 0.04 | 0.51 | 0.02 |
| **Jugular** |  |  |  |  |  |  |  |  |  |  |
| 1 hr | 28452995 | 5098116 | 13055 | 165 | 12462 | 74 | 0.58 | 0.03 | 0.51 | 0.03 |
| 3 hr | 32132121 | 1968340 | 13118 | 128 | 12499 | 49 | 0.56 | 0.03 | 0.50 | 0.01 |
| 6 hr | 28263832 | 2338260 | 13130 | 139 | 12497 | 57 | 0.59 | 0.02 | 0.51 | 0.03 |
| 8 hr | 30483625 | 3040358 | 13102 | 132 | 12478 | 54 | 0.57 | 0.04 | 0.52 | 0.02 |
| 24 hr | 29683203 | 2546563 | 13112 | 105 | 12502 | 40 | 0.53 | 0.08 | 0.52 | 0.01 |

There was no difference (P>0.05) on library parameters of 3’/5’ bias, efficiency of reads assigned to the annotation, number of genes in relationship to the location of blood sampling, nor the delayed processing of the samples (Table 2, Additional files 3 and 4). We noted, that among the samples with delayed processing, the library with the lowest efficiency of reads assigned to the annotation (40%) did not originate from the sample with lowest RIN (7.4), but both samples were processed after 24hr of preservation at 4°C. Further interrogation of the relationship between RIN and library percentage of reads assigned to the annotation showed only a moderate correlation between these two metrics (*Pearson’s r*=0.3799, P=0.0610).

There was no difference (P>0.05) for number of genes detected pre- or post-filtering in relationship to sampling location or delayed processing (Table 2, Additional files 3 and 4). After filtering for lowly expressed genes (RPKM>1 and CPM>1 in five or more samples), we quantified transcript abundance for 12,414 protein-coding genes, followed by 287 long non-coding RNAs and 109 pseudogenes.

### Differential transcript abundance based on sampling location and processing delay

First, we tested whether transcript abundance would be distinguishable based on the location of sampling. The results show no difference (FDR>0.05) between transcripts from samples obtained from coccygeal or jugular veins. Next, we assessed the consistency of transcript abundance within each animal by calculating the correlation of the transcript abundance between the two sources of sampling. The Pearson’s correlation coefficients were greater than 0.99 for all subjects. Both results convergently show no variation in transcript abundance within subject based on sampling source (Fig. 2). Thus, mRNA quantitation data collected from liquid biopsies are consistent regardless of which vein is used for sampling.

**Fig. 2.**
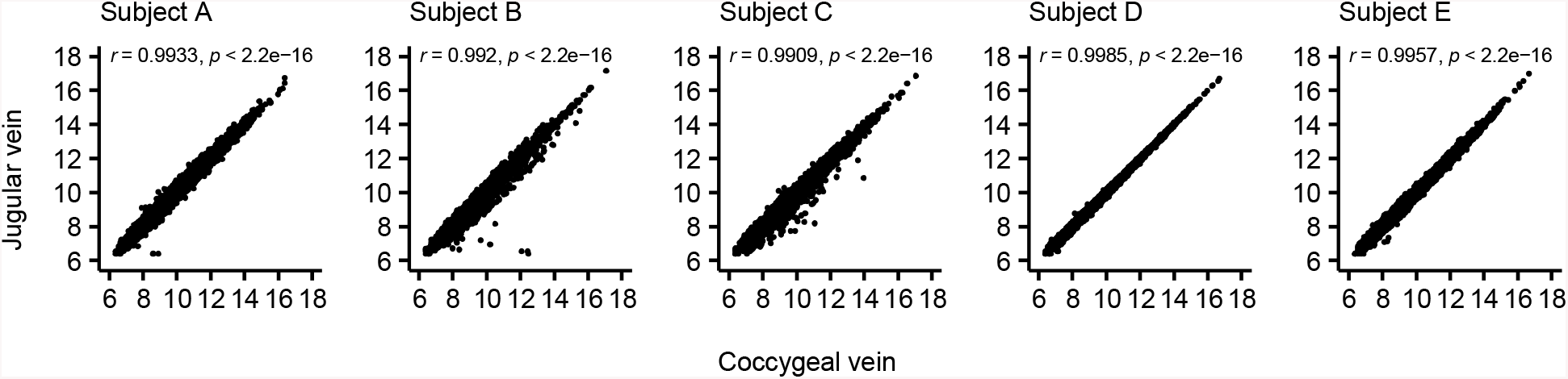
Transcript abundances obtained from the jugular and coccygeal veins from different subjects. The values presented are variance stabilizing read counts obtained from the “DESeq2” package [28].

Second, we asked if the transcript abundance in PWBCs would change if blood samples remained stored at 4°C for different periods of time, relative to the processing of the blood samples within one hour of collection. There was no differential transcript abundance between the samples stored for three or six hours at 4°C relative to the samples processed within one hour of collection (FDR>0.05, Fig. 3A).

**Fig. 3.**
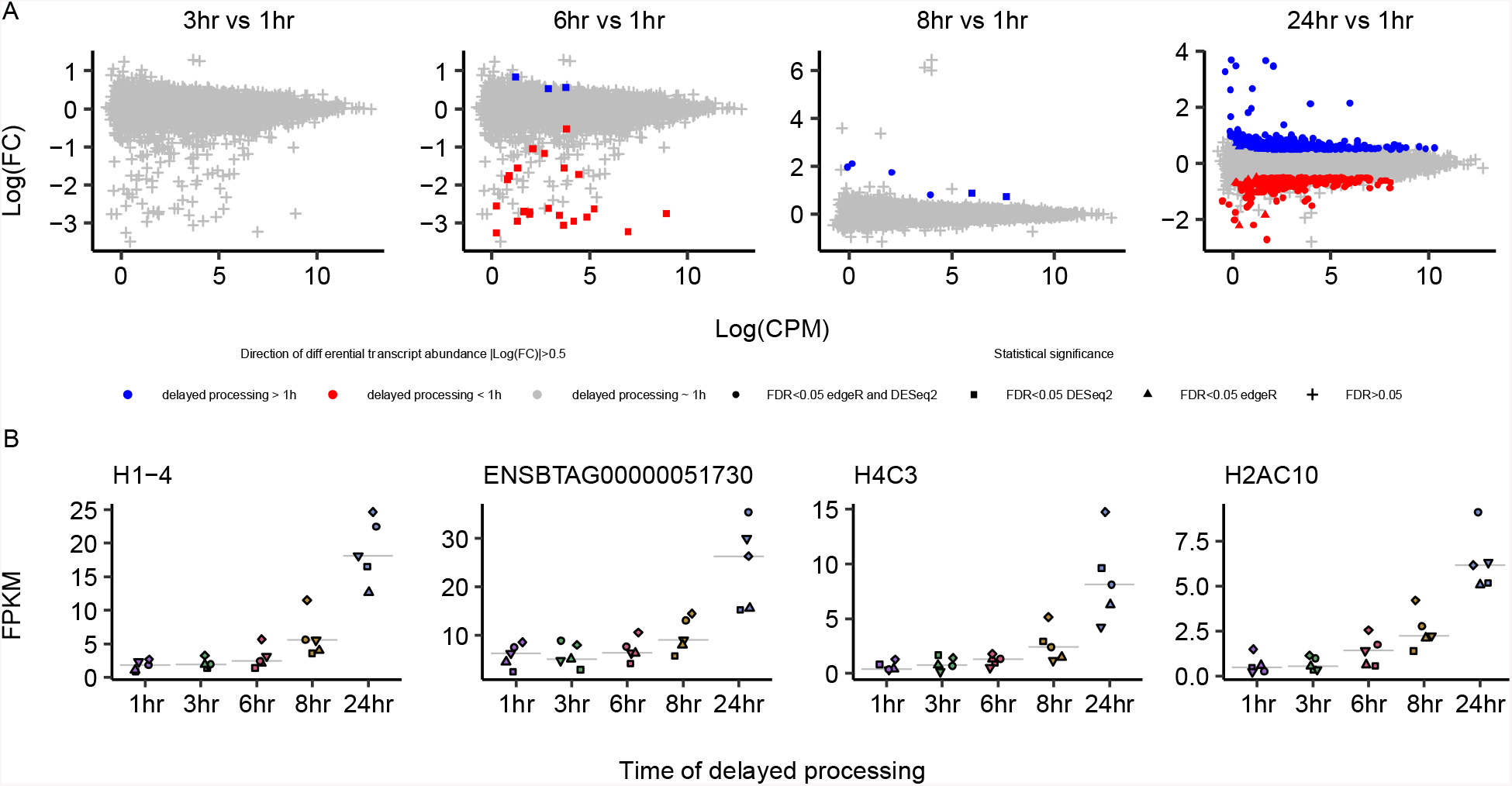
Comparison of transcript abundance in PWBCs from blood samples processed after prolonged storage at 4°C. **A**. M-A plots of the contrasts between each of the prolonged storage times versus samples processed within one hour. **B**. Transcript abundance for genes with differential abundance in both the ‘8hr vs 1hr’ and ‘24hr vs 1hr’ contrasts.

By comparison, we identified four and 515 genes with differential transcript abundance between samples stored for eight and 24 hours, respectively, at 4°C relative to the samples processed within one hour of collection (Fig. 3A). Notably, the four genes detected in the ‘8hr vs 1hr’ contrast were also detected in the ‘24hr vs 1hr’ contrast with higher transcript abundance in the preserved samples relative to those processed within one hour of collection (Fig. 3B). Furthermore, 291 and 224 genes presented greater and lower abundance, respectively, for the ‘24hr vs 1hr’ contrast (please see Additional file 5 for the lists of genes, and Additional file 6 for the individual graphs of transcript abundance for all genes for the contrast ‘24hr vs 1hr’). These results show that storage of blood samples for ≥ 8 hours prior to cryopreservation of PWBCs causes significant changes in the transcriptome profile.

### Analysis of the relationship between a decline in transcript abundance and mRNA degradation

Considering the results of differential transcript abundance, we asked if lower values of FPKM for transcripts in the samples processed after 24hr of storage at 4°C were caused by transcript degradation. First, we inspected whether there was a global trend of transcripts to have prominent degradation in one of the extremities (3’ or 5’). Coverage plots for the 224 genes with lower transcript abundance at 24hr of delayed processing showed a relative nucleotide sequencing depth similar to the 12,264 genes that were not differentially abundant (Fig. 4A).

**Fig. 4.**
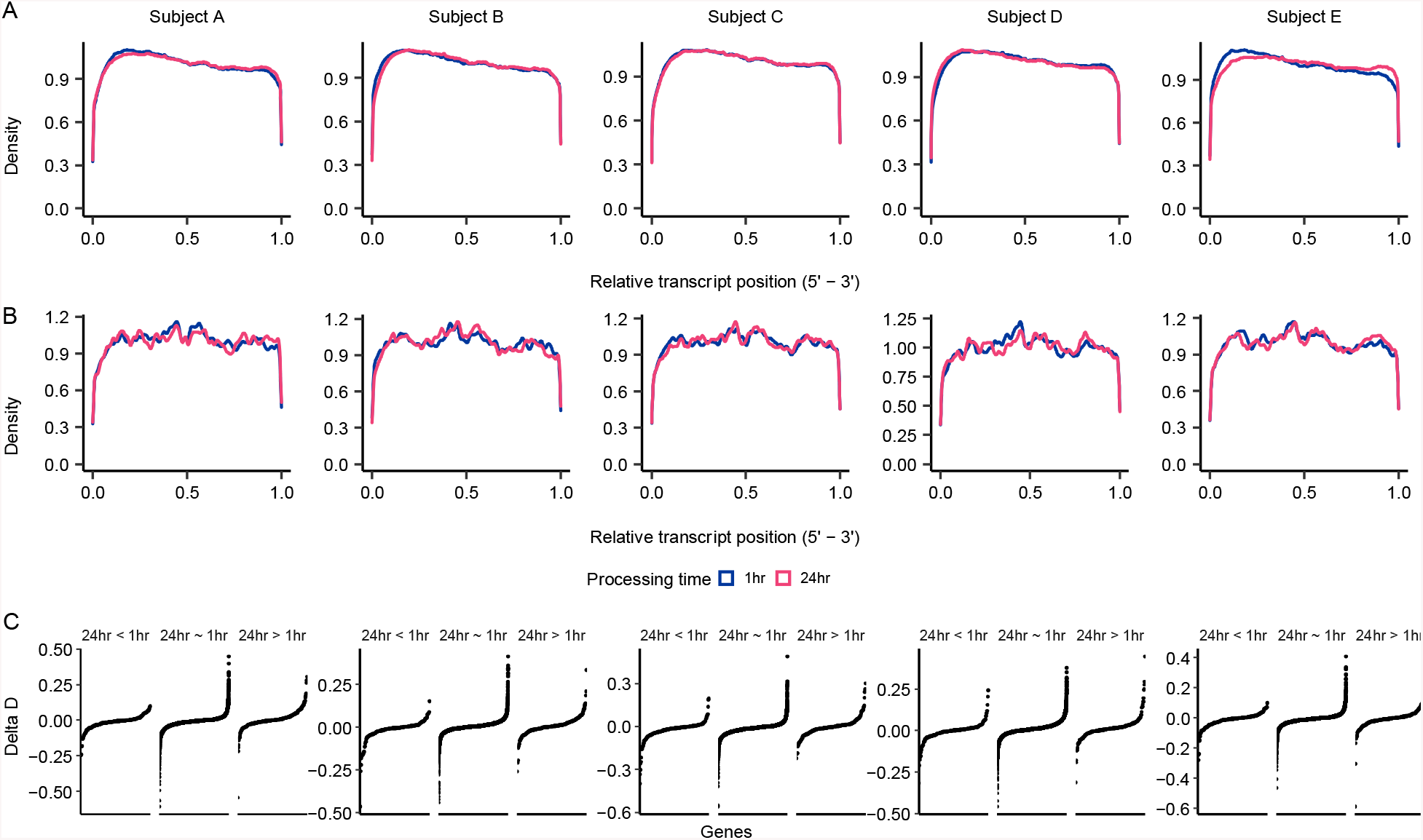
Comparison of overall transcript sequencing coverage from samples that were processed within 1hr and 24hr after blood collection. **A**. Genes with no change in transcript abundance 24hr of storage at 4°C. **B**. Genes with lower transcript abundance after 24hr of storage at 4°C. **C**. Plots of the Delta D-statistic for all genes separated by their results of the contrast ‘24hr vs 1hr’.

We further tested whether the nucleotide coverage of the 224 genes with lower transcript abundance at 24hr of delayed processing was statistically different from distribution of the same genes observed at 1hr of processing. First, we calculated the Kolmogorov-Smirnov D-statistic [29] for the relative nucleotide coverage of genes from samples obtained from the jugular and coccygeal veins (both processed within 1hr of blood collection), which we referred to as (*D*_*j,c*,1*hr*_). We also calculated the D-statistic for the relative nucleotide coverage of genes from samples obtained from the jugular vein processed at 24hr and 1hr post-sampling, which we referred to as (*D*_*j,24h*,1*hr*_). Next, we calculated the difference between the two statistics (*Delta D* = (*D*_(*j*,24*h*,1*hr*)_–*D*_(*j,c*,1*hr*)_). We reasoned that, for a given gene, *Delta D* would approximate to zero if the variation in the sequencing coverage was similar between the samples processed at different times (24hr vs 1hr) and the samples processed at the same time (1hr). Indeed, only seven out of the 224 genes (24hr<1hr) had *Delta D* within the range of −0.25 and 0.25 (Fig. 4B, left plot). Furthermore, the range of *Delta D* calculated for the genes with lower transcript abundance at 24hr of storage (24hr<1hr) was within the range of *Delta D* calculated for the genes with no transcript variation with the passing of 24hr post-collection (Fig. 4B, center plots). Altogether, these results provided strong evidence that the overall sequencing coverage of transcripts was similar between samples processed after 24hr storage at 4°C and within one hour of sampling.

### Gene ontology enrichment analysis of differentially expressed genes

Because we did not observe a systematic reduction in transcript coverage, we reasoned that the differential transcript abundance was a cellular regulatory response to the preservation of blood samples at 4°C. It was noteworthy that three out of four genes with greater transcript abundance at ‘8hr vs 1hr’ (*H1-4, H2AC10* and *H4C3*) were involved in chromatin configuration, specifically annotated with the gene ontology terms ‘nucleosome assembly’ (*H1-4* and *H4C3*), and ‘chromatin silencing’ (*H2AC10*).

Further interrogation of the 291 genes that had greater abundance at ‘24hr vs 1hr’ also revealed an enrichment of the category ‘nucleosome assembly’ with a series of histone related genes (*H1-4, H2BC12, H2BC13, H2BC14, H2BC18, H2BC4, H2BU1, H4C14, H4C3, H4C4* and *H4C8*, fold enrichment=9.78, Fig. 5A, please see Additional file 7 for a complete list of categories and gene annotation). All these genes overlapped with the molecular function ‘DNA binding’ (*BSX, DNMT3B, H1-4, H2AC21, H2AC6, H2AW, H2BC12, H2BC13, H2BC14, H2BC18, H2BC4, H2BU1, H3C6, H4C14, H4C3, H4C4, H4C8, MSH5, PROX2, SNAPC4, TEAD3* and *TERT*, fold enrichment=1, Fig. 5B, Additional file 8). Other biological processes significantly enriched were ‘neutrophil chemotaxis’ (fold enrichment=7.34), ‘cell adhesion’ (fold enrichment=3.19), and ‘transport membrane’ (fold enrichment=2.4) (Fig. 5A, Additional file 7).

**Fig. 5.**
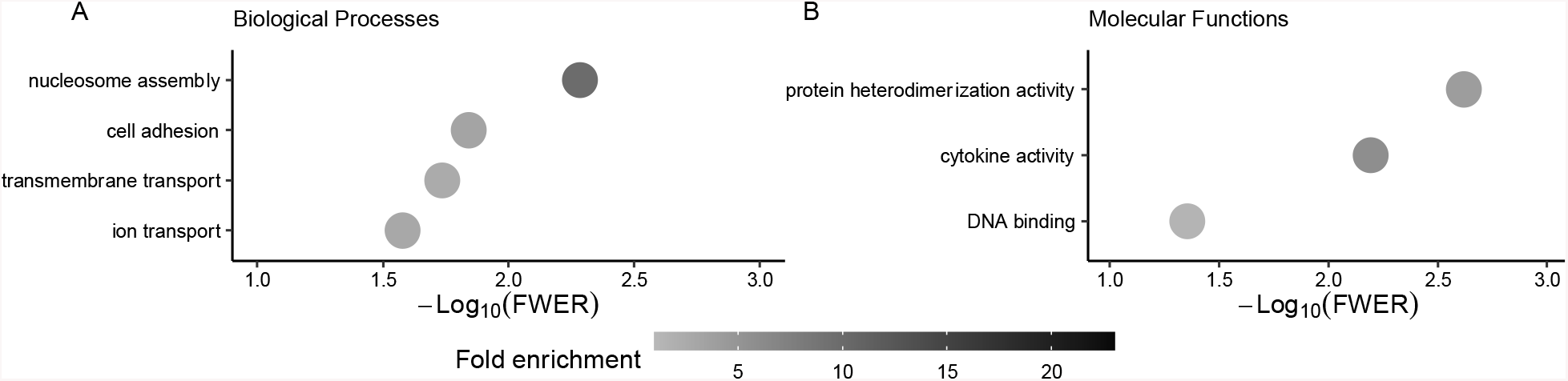
Gene ontology categories enriched in the genes with greater transcript abundance at 24hr versus one hr. **A** Biological processes and **B** Molecular functions. To improve readability, only categories with more than five genes are displayed on the graphs, please, see Additional files 7 and 8 for a full list of categories and the associated genes with their annotation.

We also asked if there was enrichment of gene ontology categories within the 224 genes that had less transcript abundance after 24hr of storage at 4°C, and there were several categories significantly enriched (FWER<0.05, Fig. 6A, please see Additional file 9 for a complete list of categories and gene annotation). Notably, there were a series of signaling related categories such as ‘positive regulation of interferon-gamma production’, ‘positive regulation of interleukin-8 production’, ‘negative regulation of interferon-gamma production’, ‘positive regulation of ERK1 and ERK2 cascade’, ‘positive regulation of MAPK cascade’ ‘positive regulation of interleukin-1 beta production’, ‘positive regulation of interleukin-12 production’, ‘positive regulation of interleukin-6 production’, ‘positive regulation of NF-kappaB transcription factor activity’, ‘positive regulation of NIK/NF-kappaB signaling’, ‘positive regulation of peptidyl-tyrosine phosphorylation’, and ‘positive regulation of phosphatidylinositol 3-kinase signaling’.

**Fig. 6.**
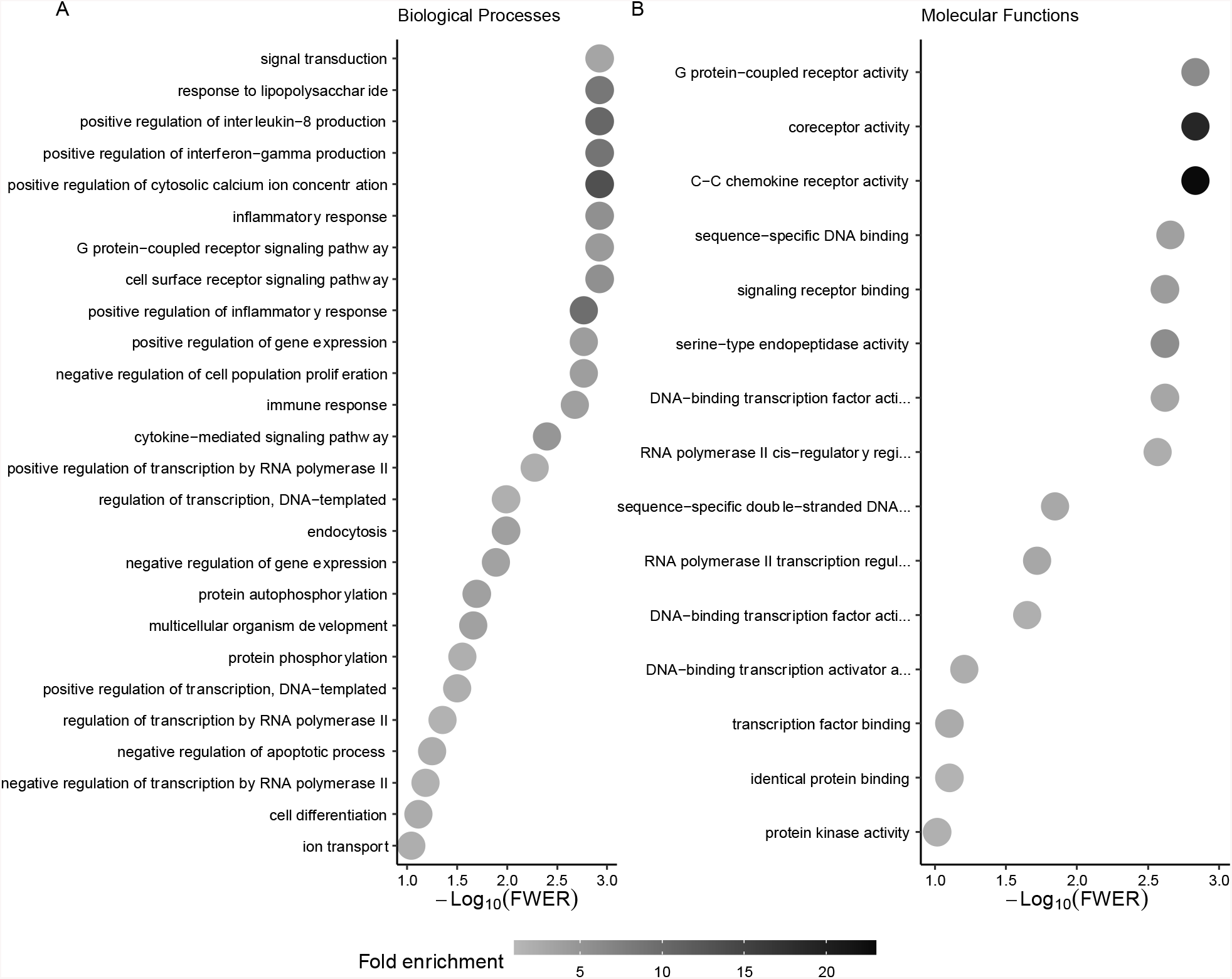
Gene ontology categories enriched in the genes with lower transcript abundance at 24hr versus 1hr. **A** Biological processes and **B** Molecular functions. To improve readability, only categories with more than five genes are displayed on the graphs, please, see Additional files 9 and 10 for a full list of categories and the associated genes with their annotation.

Also within the 224 genes that had less transcript abundance after 24hr of storage at 4°C, there was significant enrichment of a series of categories involved in regulation of transcription and gene expression (FWER<0.05, Fig. 6A, Additional file 9), such as ‘positive regulation of transcription by RNA polymerase II’, ‘regulation of transcription, DNA-templated’, ‘regulation of transcription by RNA polymerase II’, ‘positive regulation of gene expression’, and ‘positive regulation of transcription, DNA-templated’.

In parallel with the identification of the significant enrichment of the categories involved in regulation of signaling and gene expression, the test for enrichment of molecular functions identified that many of those 224 genes were associated with functions that involve interaction with DNA to regulate gene expression, such as ‘RNA polymerase II cis-regulatory region sequence-specific DNA binding’, ‘DNA-binding transcription factor activity, RNA polymerase II-specific’, and ‘DNA-binding transcription factor activity’ (Fig. 6B, Additional file 10).

## DISCUSSION

The main purpose of our study was to understand the dynamics of RNA degradation and the consequences of this RNA degradation on the quantification of transcript abundance in PWBCs from samples stored in the fridge (4°C). We collected multiple samples from the same subject and proceeded with a strategic delay in the processing of samples, followed by immediate cryopreservation of PWBCs. Our methodical interrogation of the RNA quality and systematic analysis of transcriptome data lead us to identify critical factors related to the short-term preservation of blood samples for RNA analysis: (i) the vein used for sampling blood is not a source of significant and systematic changes in the transcriptome profiling of PWBCs; (ii) storing blood samples under refrigeration for 24 hours does reduce their RIN values by approximately one unit, however the drop in RIN values does not interfere with the quantification of transcripts from protein-coding genes or long non-coding RNAs produced in PWBCs; (iii) even if blood samples are refrigerated, the abundance of gene transcripts produced in PWBCs starts to drop irregularly as early as three hours past blood sampling, but changes are consistent across samples after eight hours of refrigeration; and (iv) the transcriptome of PWBCs is severely altered after blood samples are refrigerated for 24 hours post-collection.

### Decline in transcript abundance was not a consequence of mRNA degradation in blood samples preserved at 4°C for 24 hours

According to our hypothesis, we expected that storage of blood tubes at 4°C for a long period of time would reduce the RNA quality through degradation. Indeed, there was a reduction in RIN values from RNA obtained from PWBCs after blood samples were preserved at 4°C for 24 hours (from 8.52±0.37 at 1hr to 8±0.37 at 24hr). The relatively high values of RIN after the preservation of blood samples at 4°C for 24 hours are similar to RIN values reported elsewhere [23]. However, the results observed from the RNA-sequencing did not show indication of reduced RNA quality. The values of the 3’/5’ bias for all libraries ranged from 0.47 to 0.56, with no effect of the processing time on the averages. In samples with degraded RNA, there is a bias towards transcript coverage on the 3’ end, whereas samples with 3’/5’ bias values close to 0.5 have balanced coverage of RNA extremities and are only observed in samples with high RNA quality [30]. Thus, there was no systematic coverage bias towards the 3’ end of polyadenylated transcripts in our samples.

Also based on our hypothesis, we anticipated that transcripts with significantly lower quantification would be a consequence of RNA degradation following a period of storage of blood samples at 4°C. Here we reasoned that coverage plots for the 224 genes with lower abundance at 24 hours in the contrast ‘24hr vs 1hr’ would be distinct between the libraries produced from the 24hr group versus the 1hr group. Contrary to our expectation, the coverage charts (Fig. 4A) showed a virtually identical coverage of transcripts with significantly lower quantification whether on samples processed within one hour or 24 hours of collection. Furthermore, a comparison of the distributions using the Kolmogorov-Smirnov test confirmed no significant changes in transcript coverage based on the amount of time that samples were preserved.

A possible explanation for the discrepancy between the significantly lower values of RIN for samples preserved for 24 hours and the consistent sequencing coverage across transcripts is on the source of data. The RIN values are computed based on data collected from a series of features of an electropherogram, most of which involve information from ribosomal RNAs (5S, 5.8S, 18S and 28S) [31]. On the other hand, RNA-sequencing libraries were produced with enrichment of polyadenylated transcripts, and thus, the results from 3’/5’ bias nor coverage plots do not account for ribosomal RNA. Our results indicate that, although a correlation between transcript coverage and RIN values have been identified [31, 32], this relationship may be prominent in samples with RIN values less than 8.

### Alteration in transcript abundance as a biological response to preservation at 4°C

Our results show that there is a prominent systematic alteration of transcript abundance in PWBCs when blood samples are preserved for 24 hours at 4°C, which is aligned with previous reports [23, 26]. Interestingly, we determined that this alteration of transcript abundance across individuals starts as early as eight hours post-collection.

Because we could not find indication that RNA degradation was a cause of these alterations, we reasoned that the alteration in transcript abundance was a consequence of the PWBCs responding to the cold temperature (4°C) and lack of oxygen. The consequences of long-term exposure of mammalian cells at 4°C have not been well studied, but Al-Fageeh and Smales [33] proposed that the active transcription of a selected group of genes would cause a wide-spread reduction in transcription activity. Well-aligned with this possible mechanism, three out of four genes up regulated in PWBCs after the storage of blood samples for eight hours at 4°C have a role in chromatin organization, including nucleosome assembly, which can be related to a compaction of the chromatin and reduction in transcriptional activity.

The alteration of transcript abundance in PWBCs after storing blood samples at 4°C for 24 hours has been observed before [26]. However, our genome-wide transcriptome analysis shows that the changes are more prominent after 24 hours of storage of blood samples at 4°C. The genes with greater transcript abundance after 24 hours of storage of blood samples at 4°C relative to those processed within one hour of collection seem to be enriched for few biological processes and again with a high enrichment for genes involved in nucleosome assembly. It is possible that the cells increase the transcription of histone related genes to increase the genome-wide compaction of chromatin. The greater number of biological processes enriched for genes with lower transcript abundance after 24 hours of storage of blood samples at 4°C relative to those processed within one hour of collection corroborate the notion of a global silencing in transcription.

### The consequence of preservation of blood samples for the discovery of biomarkers

Considering the results of significant differential transcript abundance observed in the present study, we reasoned that the prolonged storage of blood samples at 4°C would be relevant for investigations searching for mRNA markers in PWBCs. The overlap of our results with transcript abundance of genes also expressed in PWBCs and previously associated with fertility in heifers [20, 21] identified two genes (*NKG2A, PPP1R3B*, Additional file 11) whose analysis of differential transcript abundance would have been compromised by storage of blood samples at 4°C for eight hours or longer. These results strongly indicate that blood samples collected for studies of mRNA biomarkers should: (i) be preserved on ice as soon as they are collected and processed as early as possible, and certainly within 6 hours of collection, for the proper cryopreservation of PWBCs, or (ii) if possible, collected in tubes that allow for the immediate preservation of RNA transcript abundance in the whole blood. However, we note that the chemical or cryopreservation of whole blood for RNA extraction requires further depletion of hemoglobin transcripts if the samples will be used for RNA-sequencing [34, 35].

## CONCLUSIONS AND IMPLICATIONS

The transcriptome of PWBCs changes after blood sampling, even if the samples are refrigerated. A systemic alteration is detected at eight hours post blood collection and follows a pattern where PWBCs increase the transcription of genes related to chromatin compaction. This compaction is likely to reduce the transcription of several genes that function across multiple cellular processes in PWBCs. It is evident that this alteration in transcriptome profiles after prolonged storage can mask the transcriptome signature of a specific physiological phenomenon.

Our findings can be used as a guide for the establishment of protocols for blood processing when samples are supposed to be used for genome-wide quantification of transcripts in PWBCs. Blood samples collected for mRNA-based biomarker discovery should be refrigerated immediately and processed within six hours post-sampling. This recommendation can be considered by investigators working in diverse several areas of life sciences.

## METHODS

### Animal handling and sample collection

Eleven crossbred beef heifers (Angus x Simmental cross), averaging 14 months of age, located at Kentland Farm (Virginia Tech, Blacksburg, VA) were subjected to estrus synchronization. On day zero we administered an intramuscular injection of gonadotrophin-releasing hormone (GnRH, 100 μg; Factrel®; Zoetis Incorporated, Parsippany, NJ) and inserted a controlled internal drug release (CIDR, 1.38 gram Progesterone; Eazi-Breed™ CIDR®; Zoetis Inc.) device in each heifer. On day seven we removed the CIDR insert and administered an intramuscular injection of prostaglandin F2alpha (PGF2α, 25 μg; Lutalyse®; Zoetis Inc.), which was followed by a second injection of GnRH on day ten of the protocol. We used estrus synchronization to mitigate possible effects that the stages of the estrus cycle may have on gene expression [36].

We collected blood samples from heifers that expressed estrus (*n*=5) at the time artificial insemination would normally be performed. Fifty mL of blood were sampled from the jugular vein and 10 mL from the coccygeal vein of each heifer using vacutainers containing 18 mg K2 EDTA (Becton, Dickinson, and Company, Franklin Lakes, NJ). Each tube was inverted several times to prevent blood coagulation and placed on ice immediately until processing.

### Experimental design and blood processing

Blood tubes were sprayed thoroughly with a disinfectant (Lysol®) prior to storage. While on ice, tubes containing samples from the jugular vein were randomly assigned into five groups: 1hr, 3hr, 6hr, 8hr, and 24hr, which correspond to the time the samples remained at 4°C prior to processing. We processed blood samples from the coccygeal vein in group 1hr for comparison of gene expression with samples from the jugular vein.

The buffy coat was separated from whole blood by centrifugation for 20 min at 2000xg at 4°C. The buffy coat of each sample was aspirated and dispensed into 14 mL of a red blood cell lysis buffer solution (1.55 M ammonium chloride, 0.12 M sodium bicarbonate, 1mM EDTA, Cold Spring Harbor Protocols). The mixture was gently mixed on a rocker for 10 min at room temperature, and then centrifuged for 10 min at 800x*g* at 4°C. The supernatant was removed, and each sample was mixed with 200 ml of TRIzol™ Reagent (Invitrogen™, Thermo Fisher Scientific, Waltham, MA). The mixture of TRIzol™ and PWBCs was transferred into cryotubes (Corning Incorporated, Corning, New York) and then snap frozen in liquid nitrogen prior to storage at −80°C [20, 21].

### Total RNA extraction

Total RNA was extracted from the PWBCs using the acid guanidinium thiocyanate-phenol-chloroform procedure [37, 38], with the aid of Phasemaker™ tubes (Invitrogen™, Thermo Fisher Scientific, Waltham, MA), following the manufacturer’s instructions. Briefly, the samples were thawed on ice and 800 μL of TRIzol™ was added to each. Once homogenized, the mixture was transferred into Phasemaker™ tubes, where it was mixed with 200 μL of chloroform and centrifuged for 5 min at 12,000x*g* at 4°C to complete phase separation. Next, the aqueous phase was collected into 1.7 mL microtubes and mixed with 0.5 μL of glycoblue. Then, 500 μL of 100% isopropanol was added to each tube and they were centrifuged for 10 min at 12,000x*g* at 4°C to precipitate the RNA. The RNA pellet was collected and washed twice with 1 mL of 75% ethanol and centrifuged for 2 min at 7,500x*g* at 4°C. Then, the RNA pellet was air-dried briefly and eluted in nuclease free water and maintained on ice for quantification and assessment of quality.

We quantified the total RNA concentration (A_260_) and purity (A_260_/A_280_ ratios) using a NanoDrop™ 2000 Spectrophotometer (Thermo Fisher Scientific, Waltham, MA). We also quantified the RNA using a Qubit RNA High Sensitivity Assay Kit (Invitrogen™, Thermo Fisher Scientific, Waltham, MA) assayed on a Qubit 4 Fluorometer (Invitrogen™, Thermo Fisher Scientific, Waltham, MA). Next, we evaluated the RNA integrity by assaying a sample on an Agilent 2100 Bioanalyzer (Agilent, Santa Clara, CA) using the Agilent RNA 6000 Pico Kit (Agilent, Santa Clara, CA).

### Library preparation and high-throughput sequencing

We diluted the RNA samples to 1 ng/μL for library preparation and confirmed the concentration using a Qubit RNA High Sensitivity Assay Kit (Invitrogen™, Thermo Fisher Scientific, Waltham, MA) and Qubit 4 Fluorometer (Invitrogen™, Thermo Fisher Scientific, Waltham, MA). Five hundred ng were used as starting material for library preparation using the TruSeq® Stranded mRNA Library Prep (Illumina, Inc, San Diego, CA) and the IDT-ILMN TruSeq UD indexes. Sequencing was assayed in a NovaSeq 6000 sequencing platform (Illumina, Inc, San Diego, CA) using a NovaSeq 6000 SP Reagent Kit v1.5, to produce paired-end reads 150 nucleotides long. Preparation of libraries and sequencing assays was performed by staff at the Virginia Tech Genomics Sequencing Center.

### Alignment of sequences and filtering

We removed the sequencing adapters using cutadapt (v. 2.8) and the sequences indicated by the manufacturer (Illumina, Inc, San Diego, CA). Next, we aligned the sequences to the cattle genome [39, 40] (Bos_taurus.ARS-UCD1.2.99) obtained from the Ensembl database [41] using hisat2 (v. 2.2.0 [42]) with the --very-sensitive parameter. Using samtools (v. 1.10 [43]), we filtered the alignment to remove unmapped reads, secondary alignments, alignments whose reads failed quality control, and duplicates. We then utilized biobambam2 (v. 2.0.95 [44]) to mark and eliminate duplicates.

For the estimation of transcript coverage, we aligned the sequences trimmed from adapters to transcript sequences obtained from the Ensembl database [41] with bowtie2 (v.2.4.2 [45]) using the --very-sensitive-local parameter.

### Quantification of transcript abundance and gene filtering

We used featureCounts (subread v. 2.0.1 [46]) to count the fragments that matched to the Ensembl cattle annotation gene (Bos_taurus.ARS-UCD1.2.103). Genes annotated as protein coding, long non-coding RNA and pseudogene were retained. Following the calculation of counts per million (CPM) and reads per million per kilobase (FPKM) we retained genes that presented FPKM and CPM greater than one in five or more samples.

### Quantification of library properties

We calculated the 3’/5’ bias in our libraries using RNA-SeQC (v. 2.4.2 [30]), and the proportion of reads assigned to annotation by dividing the number of reads mapped to the Ensembl annotation divided by the total number of reads sequenced.

### Statistical analyses

#### RNA metrics (RIN, A_280_, A_260_ and A_280_/A_260_) and number of genes detected per library

We used paired Student’s t [47, 48] and Wilcoxon [49] tests to access the null hypothesis of no difference between two sampling locations (H_0_:μ_jugular_=μ_coccygeal_). Within the samples obtained from the jugular vein, we used a generalized linear mixed model to access the null hypothesis of no difference between groups of delayed processing (H_0_:μ_(T1h)_=μ_(T2h)_=…=μ_(T24h)_). The model included time of processing (*T*_*(1hr, 3hr, 6hr, 8hr or 24hr)*_) as fixed effect and animal as random variable (*A*_*(1,2,3,4 or 5)*_). When the model indicated significance of the fixed effect (P<0.05), we used the Z-test [50] and the Dunnett’s approach [51] for simultaneous tests for general linear hypothesis [52, 53] to compare the average of the groups *T*_*(3hr, 6hr, 8hr or 24hr)*_ with the baseline *T*_*(1hr)*_. Averages were inferred as statistically different when Bonferroni adjusted P<0.05.

#### Library 3’/5’ bias, proportion of reads assigned to annotation and genes detected

We used a generalized linear mixed model, with a binomial family and a logistic regression function to access the null hypothesis of no difference between groups of delayed processing (H_0_:μ_(T1h)_=μ_(T2h)_=…=μ_(T24h)_). The time of processing was included in the model as fixed effect and animal was set as random effect. Averages were inferred as statistically different when P<0.05.

#### Differential transcript abundance

We compared the transcript abundance from samples obtained from the jugular and coccygeal veins by using a paired-sample structure (H_0_:μ_jugular_=μ_coccygeal_). Next, we compared the transcript abundance from samples obtained from the jugular vein that were processed at different times. The analyses were performed with the R packages ‘edgeR’ [54] using the quasi-likelihood F-test and ‘DESeq2’ [28] using the Wald’s test. In the case of the delayed processing, we set up contrasts to compare the different processing times versus T_1h_ (H_0_:μ_(T1h)_=μ_(T2h)_; ….; H_0_:μ_(T1h)_=μ_(T24h)_). We adjusted nominal P values for multiple hypothesis testing using the Benjamini-Hochberg false discovery rate [55]. We assumed a difference in transcript abundance to be significant when FDR<0.05 in the results obtained by both ‘edgeR’ and ‘DESeq2’ packages and absolute Log_(fold-change)_>0.5. We utilized this approach to report robust results of differential transcript abundance independent of algorithm biases or limitations [20, 21, 56, 57].

#### Gene ontology enrichment analysis

We tested lists of genes for enrichment of gene ontology terms using the R package ‘GOseq’ [58] and the genes retained after filtering as a background list [59, 60]. Nominal P values were adjusted for multiple hypothesis testing by family wise error rate [61, 62].

#### Contrasts of transcript coverage

We quantified the relative position of each nucleotide in relation to the total number of nucleotides in the transcript, given in percentage. In addition, we calculated the relative proportion of occurrence of each nucleotide in relation to the total coverage of the gene. Then, for each gene in different groups, in a pair-wise manner, we compared the relative position of each nucleotide weighed by the relative coverage using the weighted Kolmogorov-Smirnov test, as described elsewhere [29].

## LIST OF ABBREVIATIONS

CIDR: controlled internal drug release
CPM: counts per million mapped reads
FDR: false discovery rate
FPKM: fragments per kilobase of transcript per million mapped reads
FWER: family-wise error rate
GnRH: gonadotrophin-releasing hormone
hr: hour
IACUC: Institutional Animal Care and Use Committee
K2 EDTA: dipotassium ethylenediaminetetraacetic acid
M: molar
mg: milligram
min: minute
mL: millilitre
mM: millimolar
mRNA: messenger RNA
ng: nanogram
P: p-value
PGF2α: prostaglandin F2alpha
PWBC: peripheral white blood cell
RIN: RNA integrity number
RPKM: reads per kilobase of transcript per million mapped reads
x*g*: relative centrifugal force
μg: microgram
μL: microliter

## DECLARATIONS

### Ethics approval and consent to participate

All animal handling and use was approved by the Institutional Animal Care and Use Committee (IACUC) at Virginia Tech.

### Consent for publication

Not applicable

### Availability of data and material

The raw data generated and analyzed during the current study are available in the GEO NCBI repository, under accession GSE192530 (https://www.ncbi.nlm.nih.gov/geo/query/acc.cgi?acc=GSE192530). To make our work fully reproducible the code utilized for the bioinformatics pipeline and analytical procedures is deposited as Additional file 12, in the figshare repository (doi: 10.6084/m9.figshare.17886068) [63] and can also be accessible at https://biase-lab.github.io/rna_temporal_expression_PWBC/index.html [64].

## Authors’ contributions

FB conceived, supervised, and obtained funding for the study. CW and FB processed the samples, analyzed the data, and wrote the paper. VM supervised the reproductive management and estrus synchronization of the heifers and sample collection. ND and SP contributed to the management and estrus synchronization of the heifers and sample collection. All authors read and approved the final manuscript.

## Acknowledgements

We thank Chad Joines (Director of Beef Operations at Virginia Tech) and the staff from the Beef Cattle Center for the support with animal handling.

## Legend for Additional files

Additional file 1. Ribonucleic acid quality and abundance parameters from samples collected from the jugular and coccygeal veins and processed within one hour of sampling. Animals are indicated by shapes across charts.

Additional file 2. Ribonucleic acid quality and abundance parameters from samples collected from the jugular vein and processed after refrigeration (4°C) for multiple windows of time (x-axis). Animals are indicated by shapes across charts.

Additional file 3. Parameters of library quality from samples collected from the jugular and coccygeal veins and processed within one hour of sampling. Animals are indicated by shapes across charts.

Additional file 4. Parameters of library quality from samples collected from the jugular and processed after refrigeration (4°C) for multiple windows of time (x-axis). Animals are indicated by shapes across charts.

Additional file 5. Results of differential transcript abundance for the contrast ‘24h vs 1h’.

Additional file 6. Charts of transcript abundance (fragment per kilobase per million reads, FPKM) for different times of refrigeration (4°C) prior to processing and cryopreservation of PWBCs. Animals are indicated by shapes across charts.

Additional file 7. Gene ontology enrichment analysis of biological processes for the genes with greater transcript abundance in PWBCs cryopreserved after 24 hours of refrigeration post collection versus one hour within collection.

Additional file 8. Gene ontology enrichment analysis of molecular functions for the genes with greater transcript abundance in PWBCs cryopreserved after 24 hours of refrigeration post collection versus one hour within collection.

Additional file 9. Gene ontology enrichment analysis of biological processes for the genes with lower transcript abundance in PWBCs cryopreserved after 24 hours of refrigeration post collection versus one hour within collection.

Additional file 10. Gene ontology enrichment analysis of molecular functions for the genes with lower transcript abundance in PWBCs cryopreserved after 24 hours of refrigeration post collection versus one hour within collection.

Additional file 11. Differential transcript abundance for genes previously detected as potential biomarkers of heifer fertility. **A**. Genes with no significant variation of transcript abundance following three, six, eight or 24 hours of refrigeration post collection versus one hour within collection. **B**. Genes with significantly differential transcript abundance following 24 hours of refrigeration post collection versus one hour within collection.

Additional file 12. Code utilized for the processing and analysis of the raw data and analytical procedures employed in this study.

