## Supplementary figures and images for "Delayed processing of blood samples impairs the accuracy of mRNA-based biomarkers"

### Additional file 1

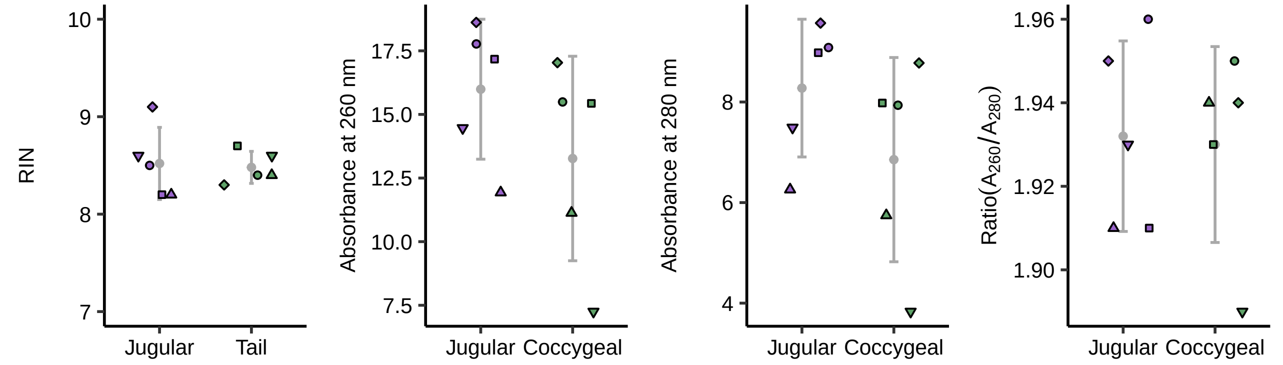

### Additional file 2

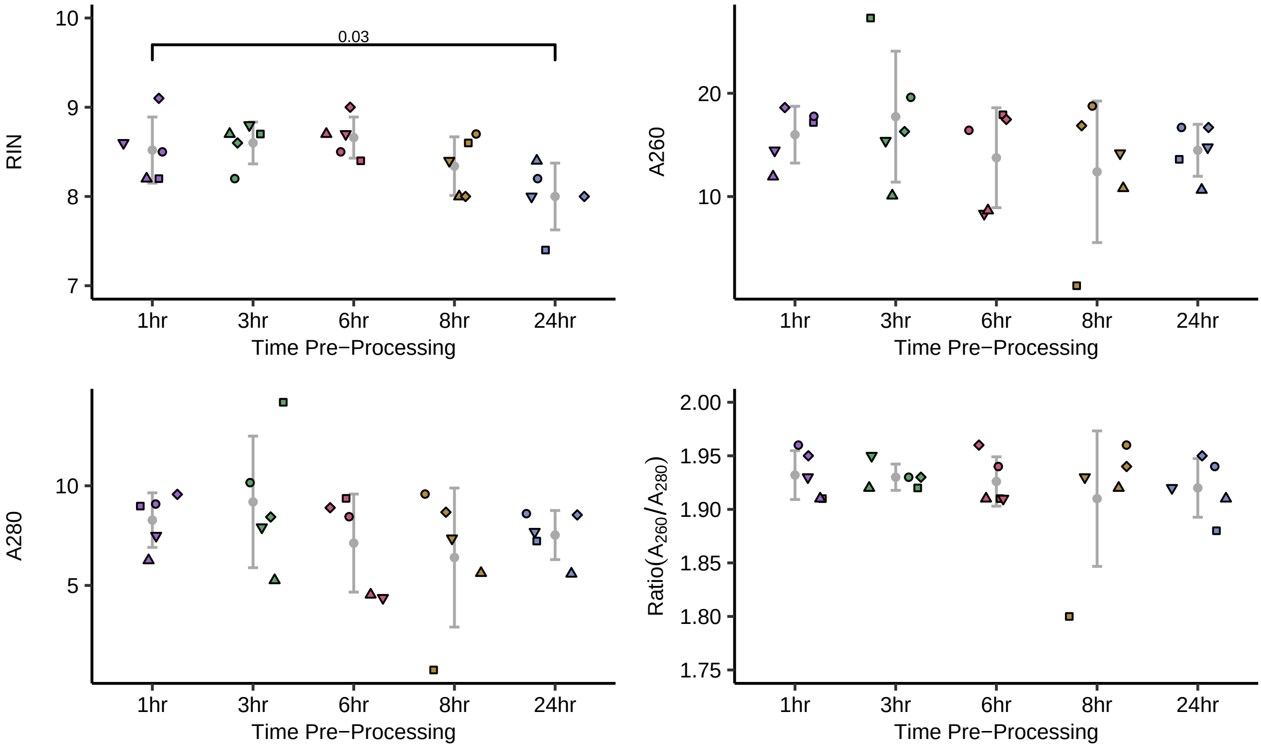

### Additional file 3

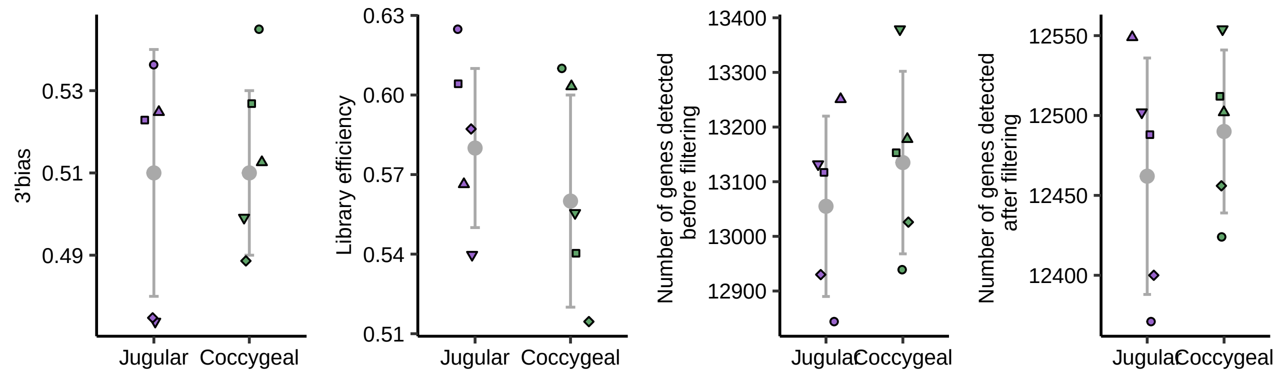

### Additional file 4

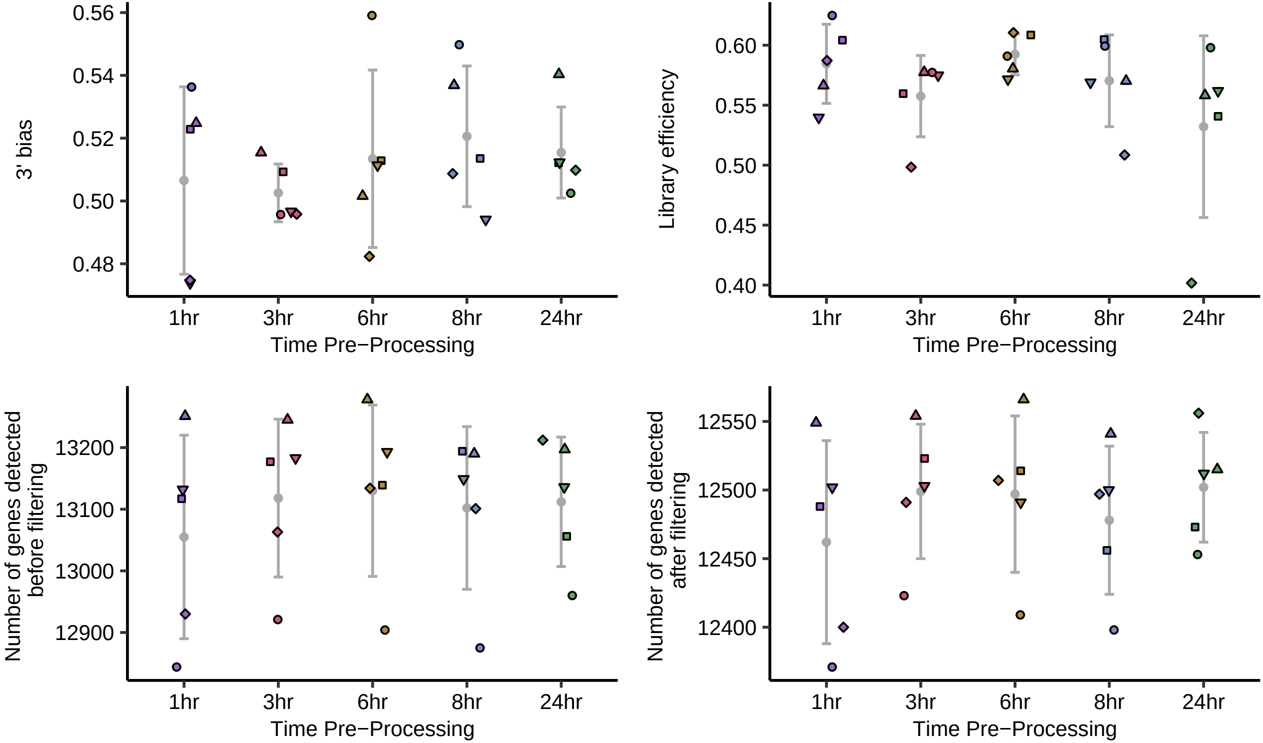

### Additional file 11

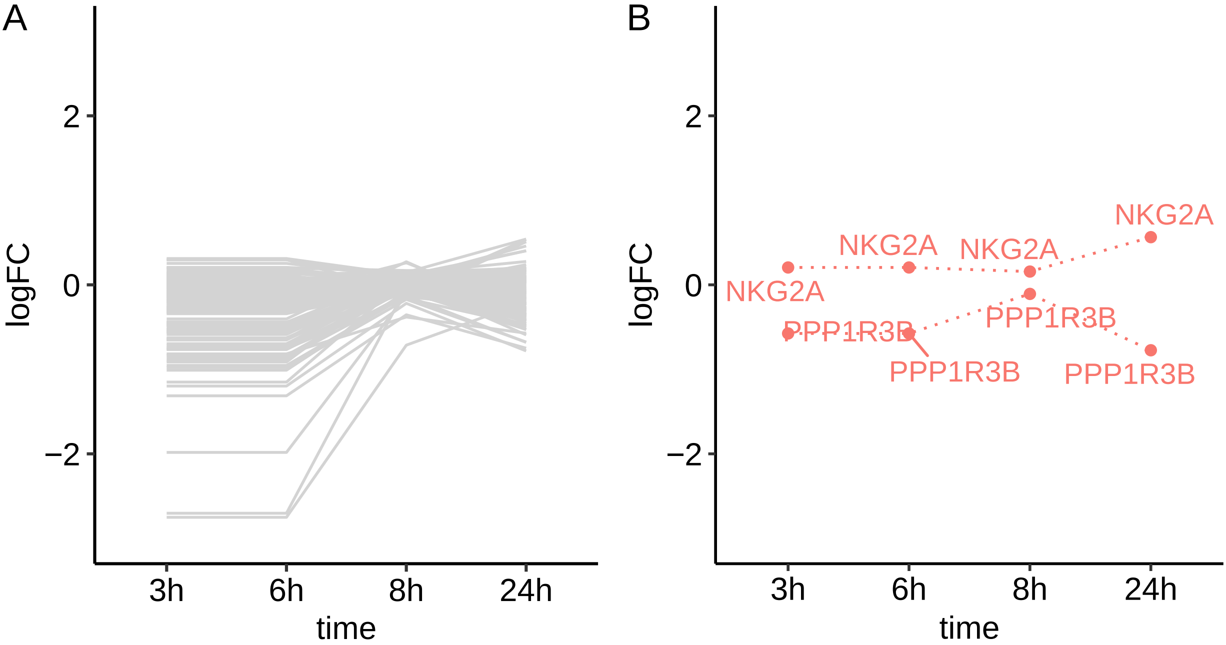
