## Additional file 6 for "Delayed processing of blood samples impairs the accuracy of mRNA-based biomarkers"

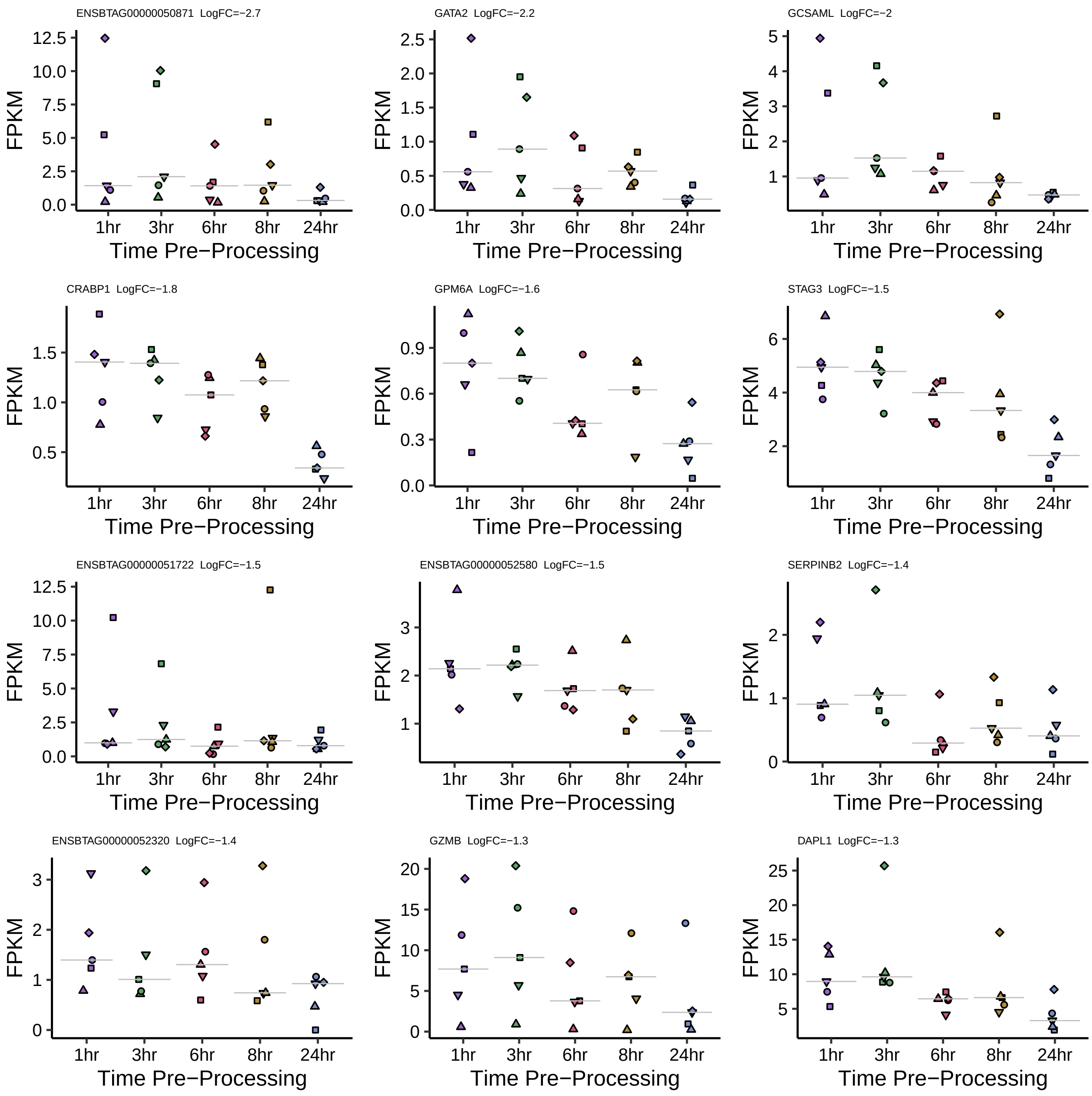

ENSBTAG00000050871 LogFC=−2.7
GATA2 LogFC=−2.2
GCSAML LogFC=−2
CRABP1 LogFC=−1.8
GPM6A LogFC=−1.6
STAG3 LogFC=−1.5
ENSBTAG00000051722 LogFC=−1.5
ENSBTAG00000052580 LogFC=−1.5
SERPINB2 LogFC=−1.4
ENSBTAG00000052320 LogFC=−1.4
GZMB LogFC=−1.3
DAPL1 LogFC=−1.3
FPKM
1hr
3hr
6hr
8hr
24hr
Time Pre−Processing

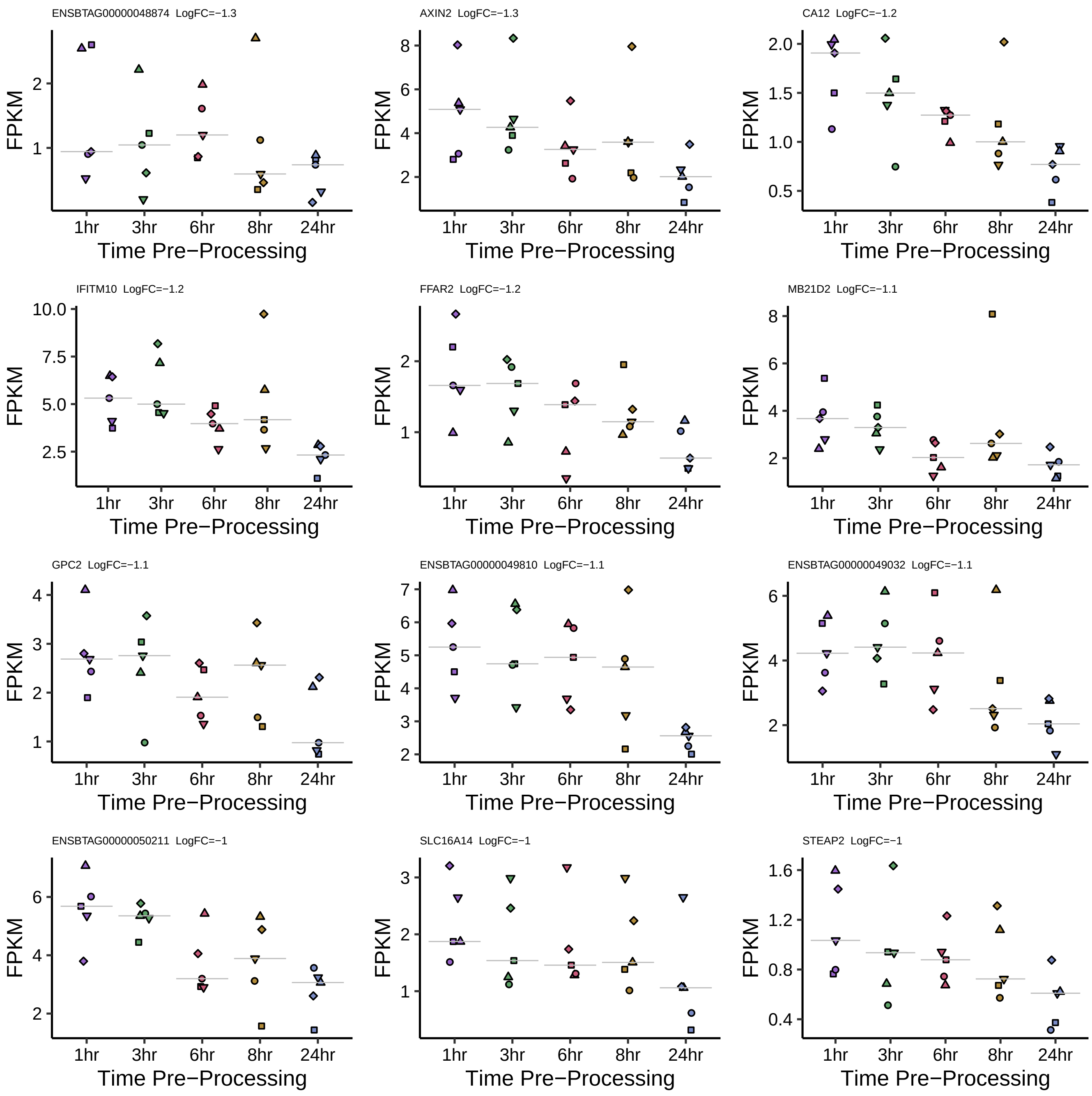
ENSBTAG00000048874 LogFC=−1.3
AXIN2 LogFC=−1.3
CA12 LogFC=−1.2
IFITM10 LogFC=−1.2
FFAR2 LogFC=−1.2
MB21D2 LogFC=−1.1
GPC2 LogFC=−1.1
ENSBTAG00000049810 LogFC=−1.1
ENSBTAG00000049032 LogFC=−1.1
ENSBTAG00000050211 LogFC=−1
SLC16A14 LogFC=−1
STEAP2 LogFC=−1
FPKM
1hr
3hr
6hr
8hr
24hr
Time Pre−Processing

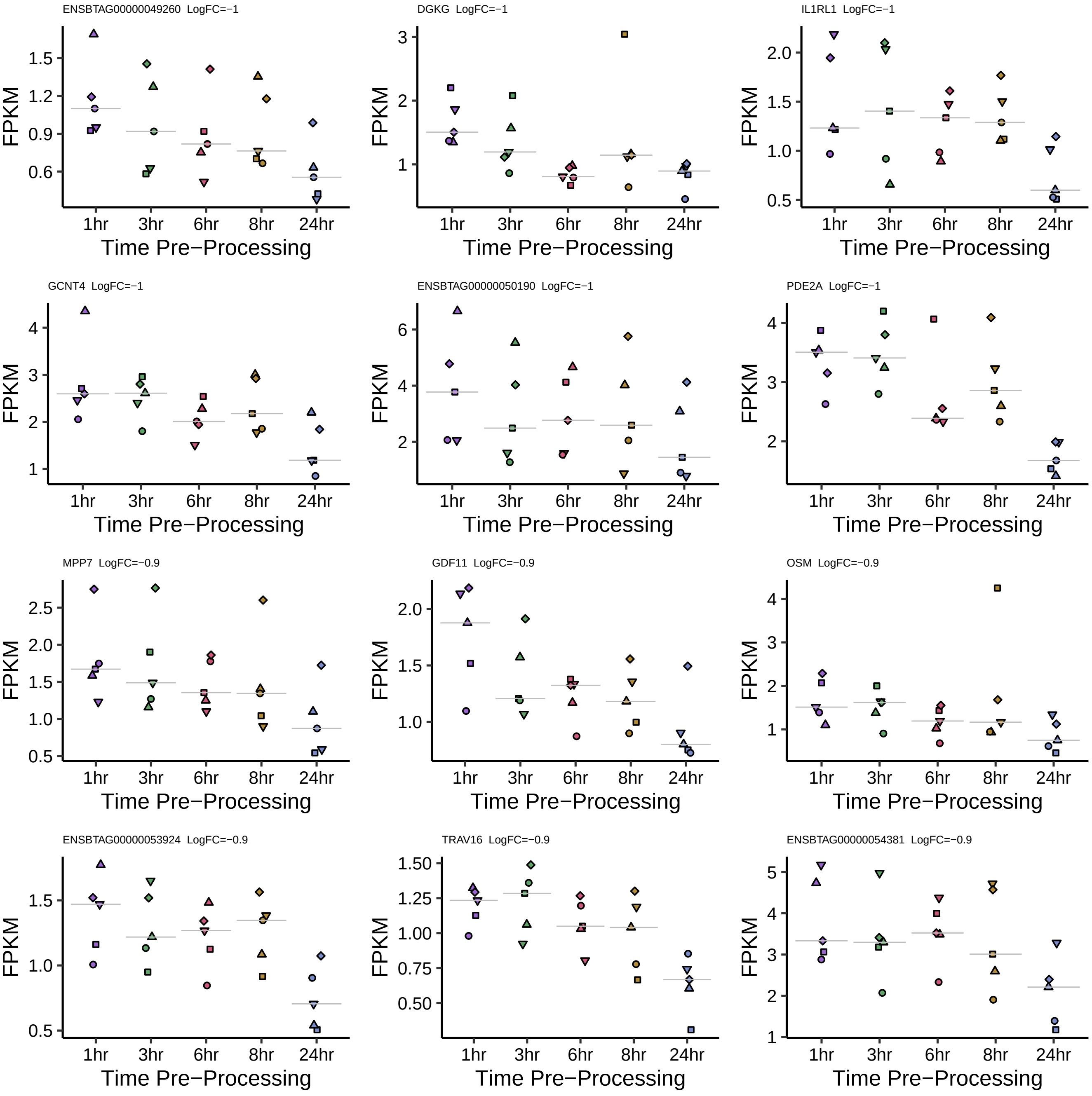
ENSBTAG00000049260 LogFC=−1
FPKM
0.6
0.9
1.2
1.5
1hr
3hr
6hr
8hr
24hr
Time Pre−Processing
DGKG LogFC=−1
FPKM
1
2
3
1hr
3hr
6hr
8hr
24hr
Time Pre−Processing
IL1RL1 LogFC=−1
FPKM
0.5
1.0
1.5
2.0
1hr
3hr
6hr
8hr
24hr
Time Pre−Processing
GCNT4 LogFC=−1
FPKM
1
2
3
4
1hr
3hr
6hr
8hr
24hr
Time Pre−Processing
ENSBTAG00000050190 LogFC=−1
FPKM
2
4
6
1hr
3hr
6hr
8hr
24hr
Time Pre−Processing
PDE2A LogFC=−1
FPKM
2
3
4
1hr
3hr
6hr
8hr
24hr
Time Pre−Processing
MPP7 LogFC=−0.9
FPKM
0.5
1.0
1.5
2.0
2.5
1hr
3hr
6hr
8hr
24hr
Time Pre−Processing
GDF11 LogFC=−0.9
FPKM
1.0
1.5
2.0
1hr
3hr
6hr
8hr
24hr
Time Pre−Processing
OSM LogFC=−0.9
FPKM
1
2
3
4
1hr
3hr
6hr
8hr
24hr
Time Pre−Processing
ENSBTAG00000053924 LogFC=−0.9
FPKM
0.5
1.0
1.5
1hr
3hr
6hr
8hr
24hr
Time Pre−Processing
TRAV16 LogFC=−0.9
FPKM
0.50
0.75
1.00
1.25
1.50
1hr
3hr
6hr
8hr
24hr
Time Pre−Processing
ENSBTAG00000054381 LogFC=−0.9
FPKM
1
2
3
4
5
1hr
3hr
6hr
8hr
24hr
Time Pre−Processing

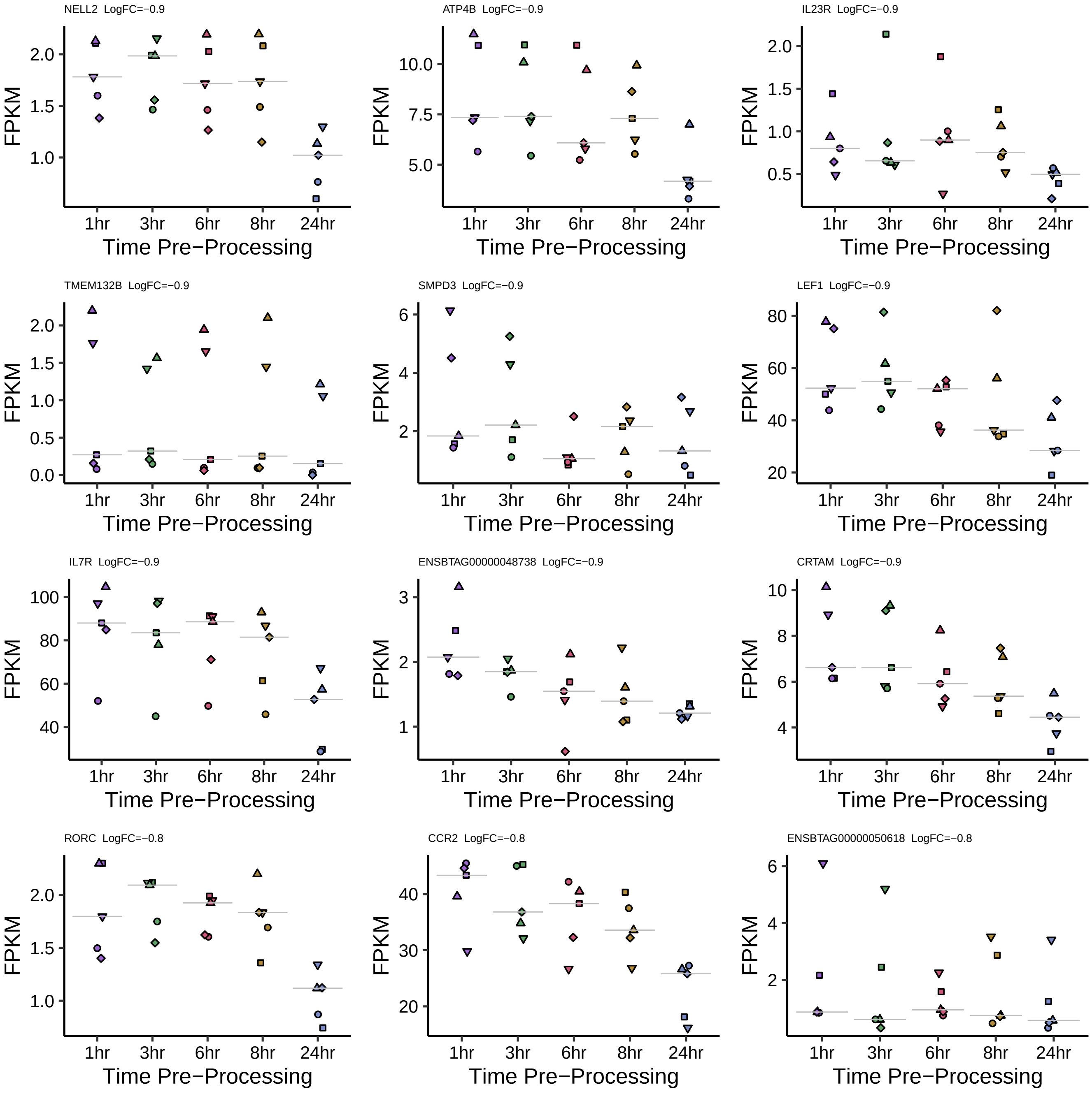
NELL2 LogFC=−0.9
ATP4B LogFC=−0.9
IL23R LogFC=−0.9
TMEM132B LogFC=−0.9
SMPD3 LogFC=−0.9
LEF1 LogFC=−0.9
IL7R LogFC=−0.9
ENSBTAG00000048738 LogFC=−0.9
CRTAM LogFC=−0.9
RORC LogFC=−0.8
CCR2 LogFC=−0.8
ENSBTAG00000050618 LogFC=−0.8
FPKM
1hr
3hr
6hr
8hr
24hr
Time Pre−Processing

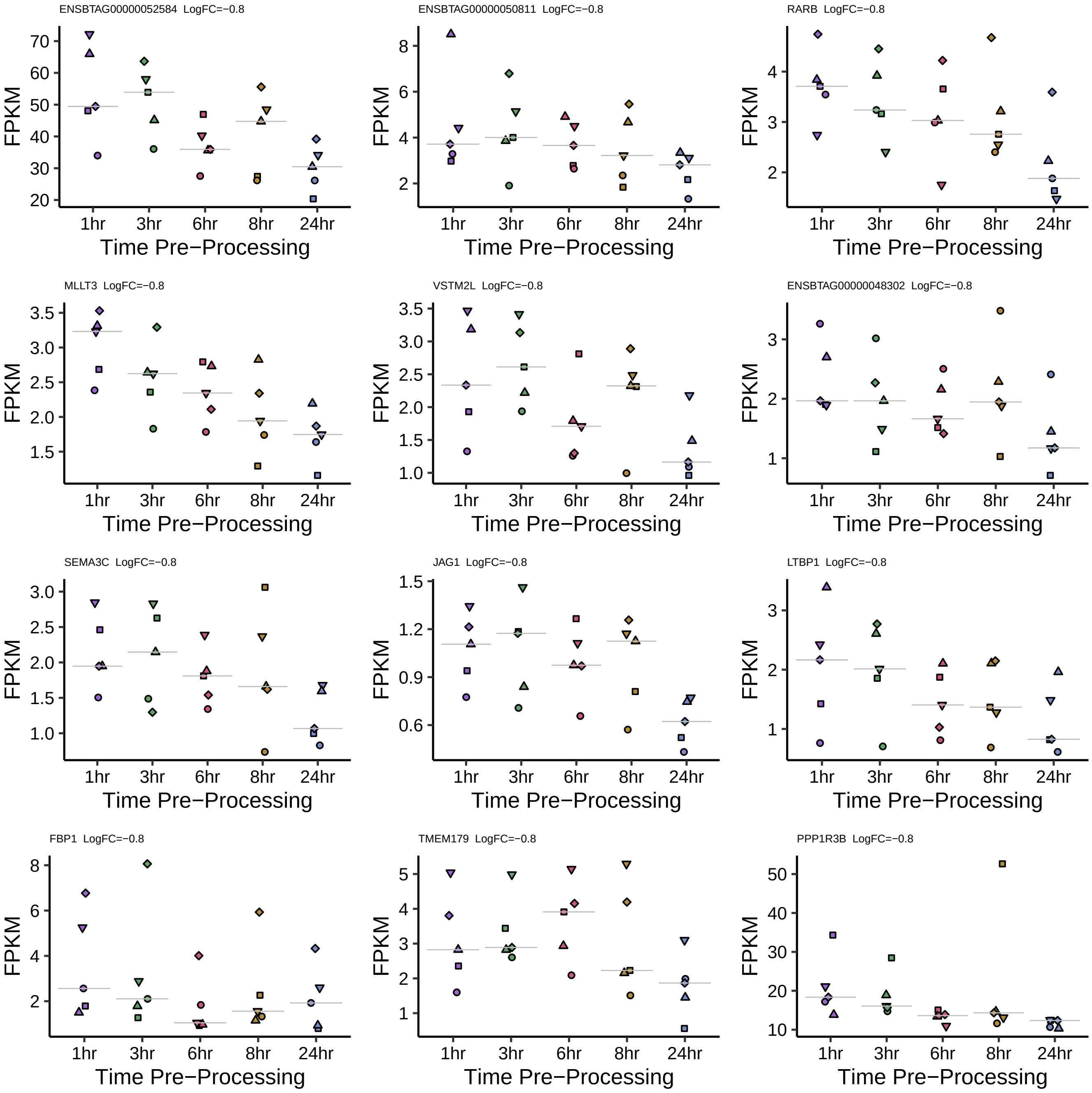

ENSBTAG00000052584 LogFC=−0.8
ENSBTAG00000050811 LogFC=−0.8
RARB LogFC=−0.8
MLLT3 LogFC=−0.8
VSTM2L LogFC=−0.8
ENSBTAG00000048302 LogFC=−0.8
SEMA3C LogFC=−0.8
JAG1 LogFC=−0.8
LTBP1 LogFC=−0.8
FBP1 LogFC=−0.8
TMEM179 LogFC=−0.8
PPP1R3B LogFC=−0.8
FPKM
1hr
3hr
6hr
8hr
24hr
Time Pre−Processing

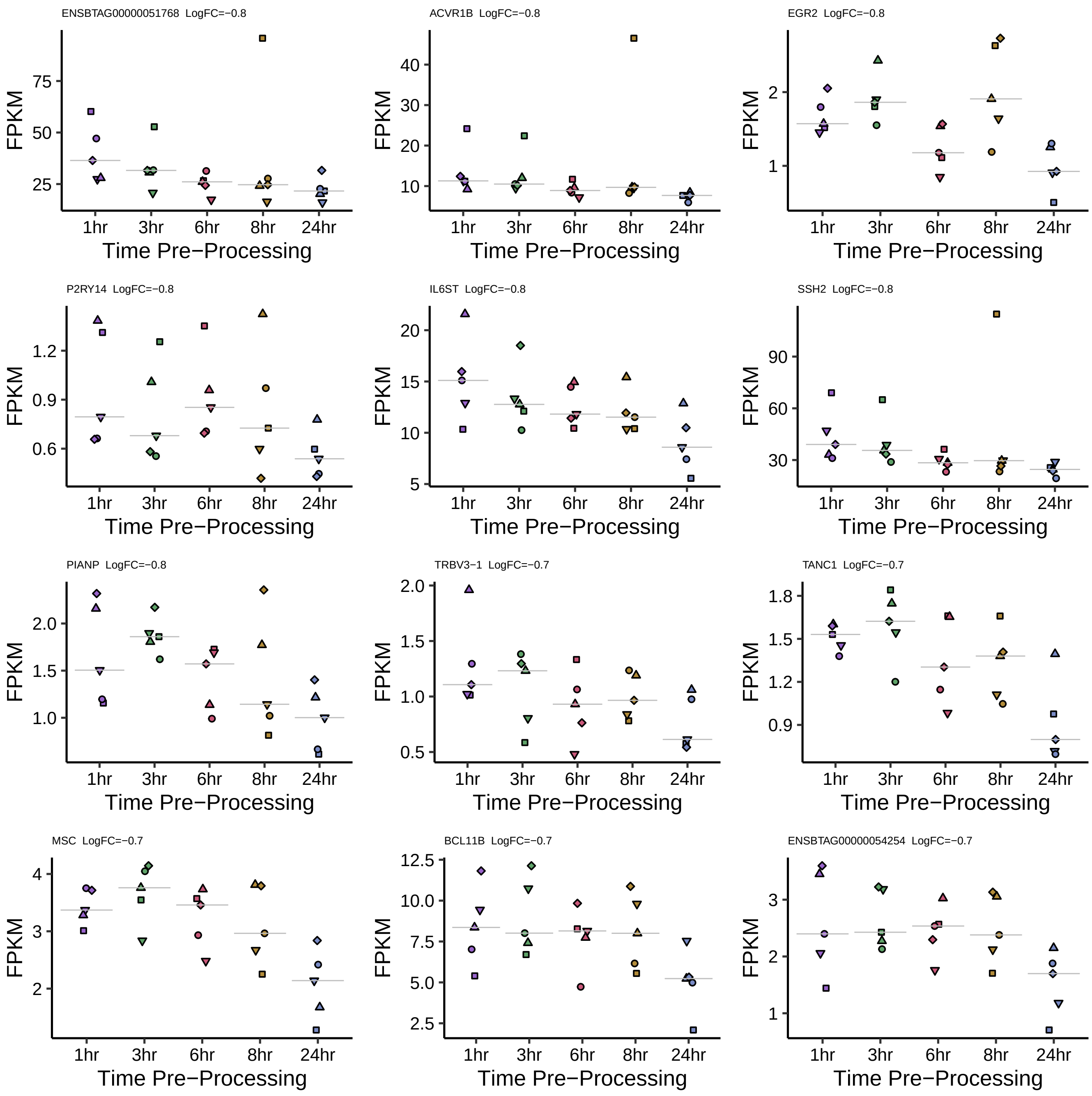

ENSBTAG00000051768 LogFC=−0.8
ACVR1B LogFC=−0.8
EGR2 LogFC=−0.8
P2RY14 LogFC=−0.8
IL6ST LogFC=−0.8
SSH2 LogFC=−0.8
PIANP LogFC=−0.8
TRBV3−1 LogFC=−0.7
TANC1 LogFC=−0.7
MSC LogFC=−0.7
BCL11B LogFC=−0.7
ENSBTAG00000054254 LogFC=−0.7
FPKM
1hr 3hr 6hr 8hr 24hr
Time Pre−Processing

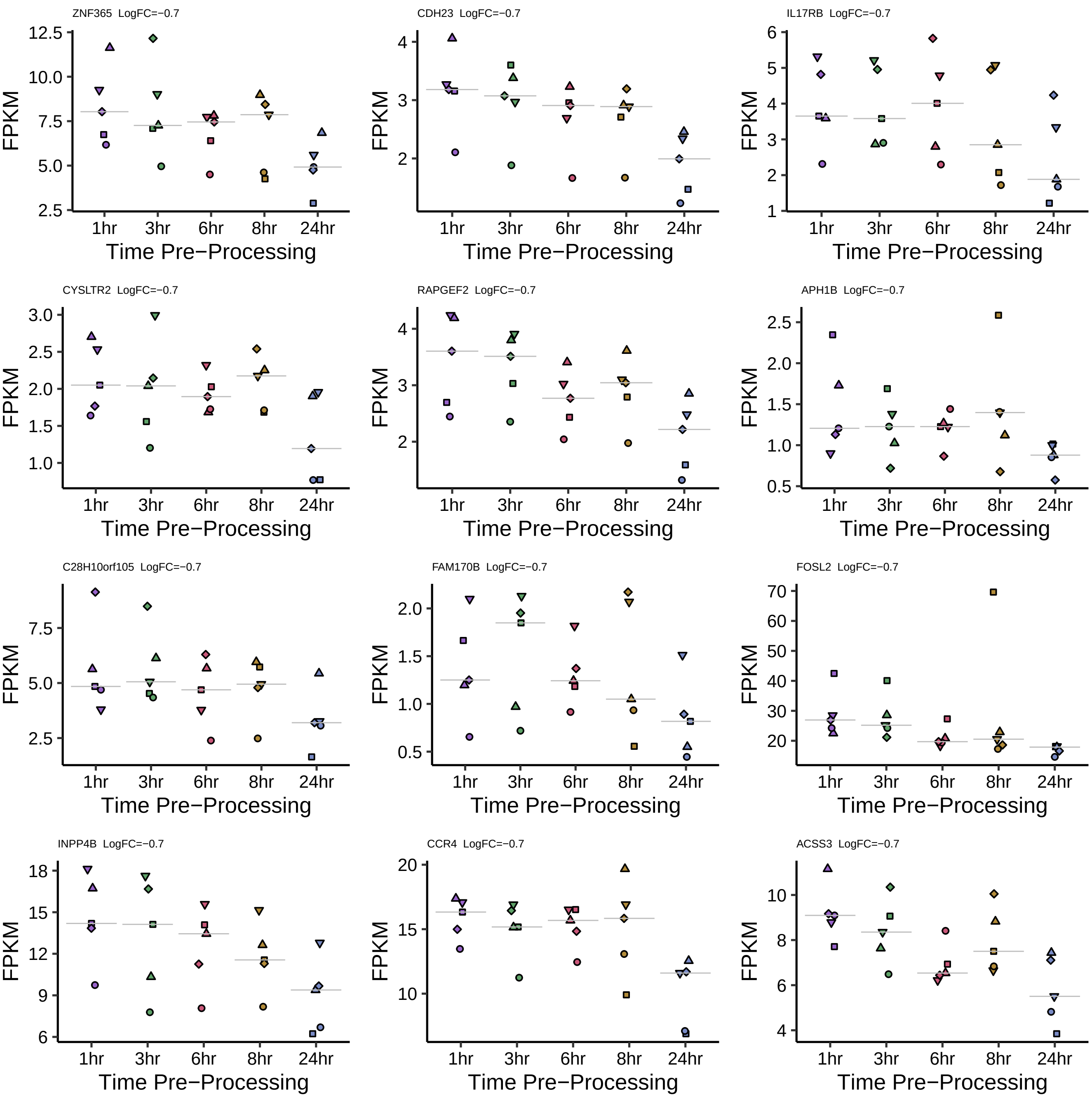

ZNF365 LogFC=−0.7
12.5
10.0
7.5
5.0
2.5
FPKM
1hr
3hr
6hr
8hr
24hr
Time Pre−Processing
CDH23 LogFC=−0.7
4
3
2
FPKM
1hr
3hr
6hr
8hr
24hr
Time Pre−Processing
IL17RB LogFC=−0.7
6
5
4
3
2
1
FPKM
1hr
3hr
6hr
8hr
24hr
Time Pre−Processing
CYSLTR2 LogFC=−0.7
3.0
2.5
2.0
1.5
1.0
FPKM
1hr
3hr
6hr
8hr
24hr
Time Pre−Processing
RAPGEF2 LogFC=−0.7
4
3
2
FPKM
1hr
3hr
6hr
8hr
24hr
Time Pre−Processing
APH1B LogFC=−0.7
2.5
2.0
1.5
1.0
0.5
FPKM
1hr
3hr
6hr
8hr
24hr
Time Pre−Processing
C28H10orf105 LogFC=−0.7
7.5
5.0
2.5
FPKM
1hr
3hr
6hr
8hr
24hr
Time Pre−Processing
FAM170B LogFC=−0.7
2.0
1.5
1.0
0.5
FPKM
1hr
3hr
6hr
8hr
24hr
Time Pre−Processing
FOSL2 LogFC=−0.7
70
60
50
40
30
20
FPKM
1hr
3hr
6hr
8hr
24hr
Time Pre−Processing
INPP4B LogFC=−0.7
18
15
12
9
6
FPKM
1hr
3hr
6hr
8hr
24hr
Time Pre−Processing
CCR4 LogFC=−0.7
20
15
10
FPKM
1hr
3hr
6hr
8hr
24hr
Time Pre−Processing
ACSS3 LogFC=−0.7
10
8
6
4
FPKM
1hr
3hr
6hr
8hr
24hr
Time Pre−Processing

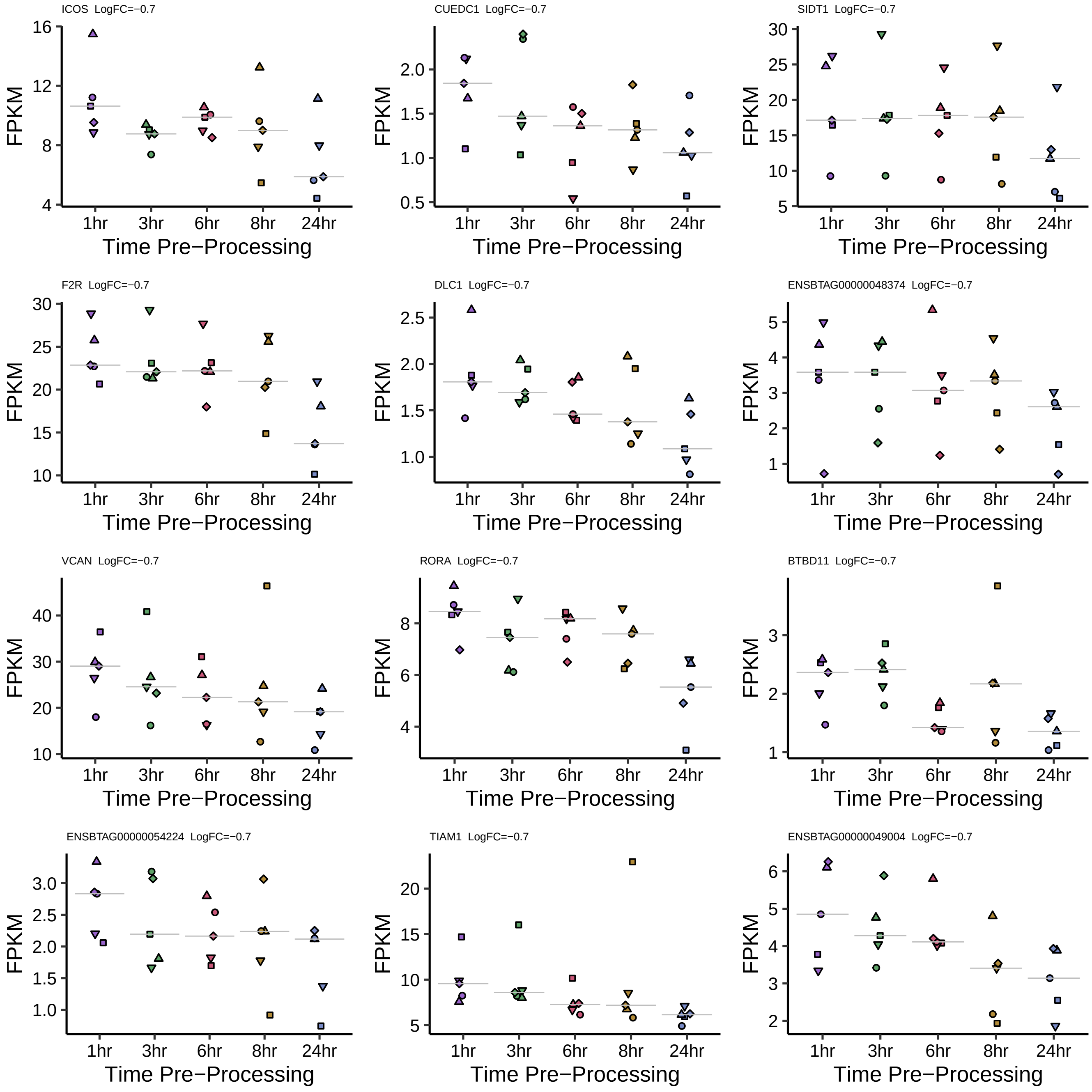

ICOS LogFC=−0.7
CUEDC1 LogFC=−0.7
SIDT1 LogFC=−0.7
F2R LogFC=−0.7
DLC1 LogFC=−0.7
ENSBTAG00000048374 LogFC=−0.7
VCAN LogFC=−0.7
RORA LogFC=−0.7
BTBD11 LogFC=−0.7
ENSBTAG00000054224 LogFC=−0.7
TIAM1 LogFC=−0.7
ENSBTAG00000049004 LogFC=−0.7
FPKM
1hr
3hr
6hr
8hr
24hr
Time Pre−Processing

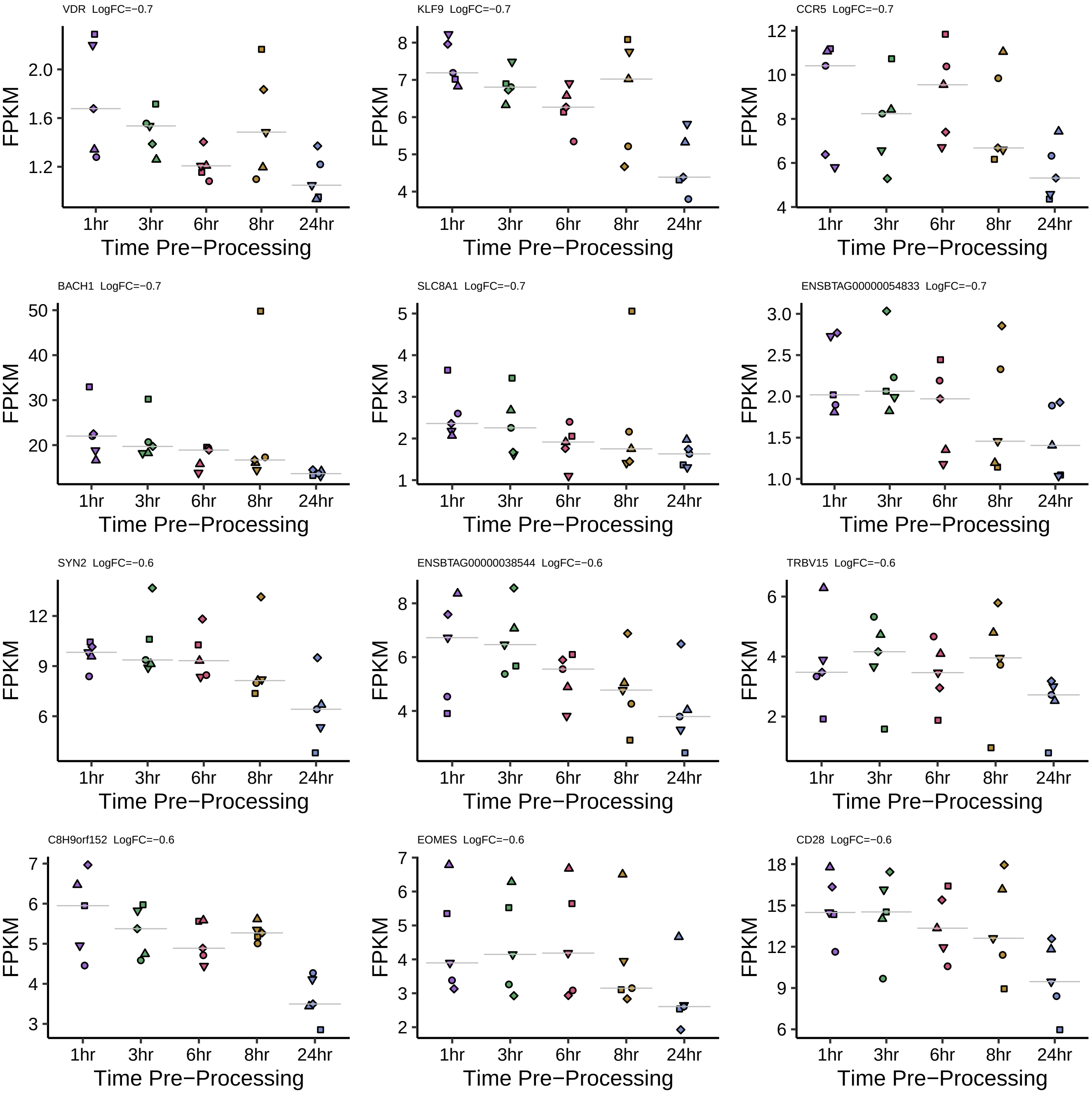
VDR LogFC=−0.7
KLF9 LogFC=−0.7
CCR5 LogFC=−0.7
BACH1 LogFC=−0.7
SLC8A1 LogFC=−0.7
ENSBTAG00000054833 LogFC=−0.7
SYN2 LogFC=−0.6
ENSBTAG00000038544 LogFC=−0.6
TRBV15 LogFC=−0.6
C8H9orf152 LogFC=−0.6
EOMES LogFC=−0.6
CD28 LogFC=−0.6
FPKM
1hr
3hr
6hr
8hr
24hr
Time Pre−Processing

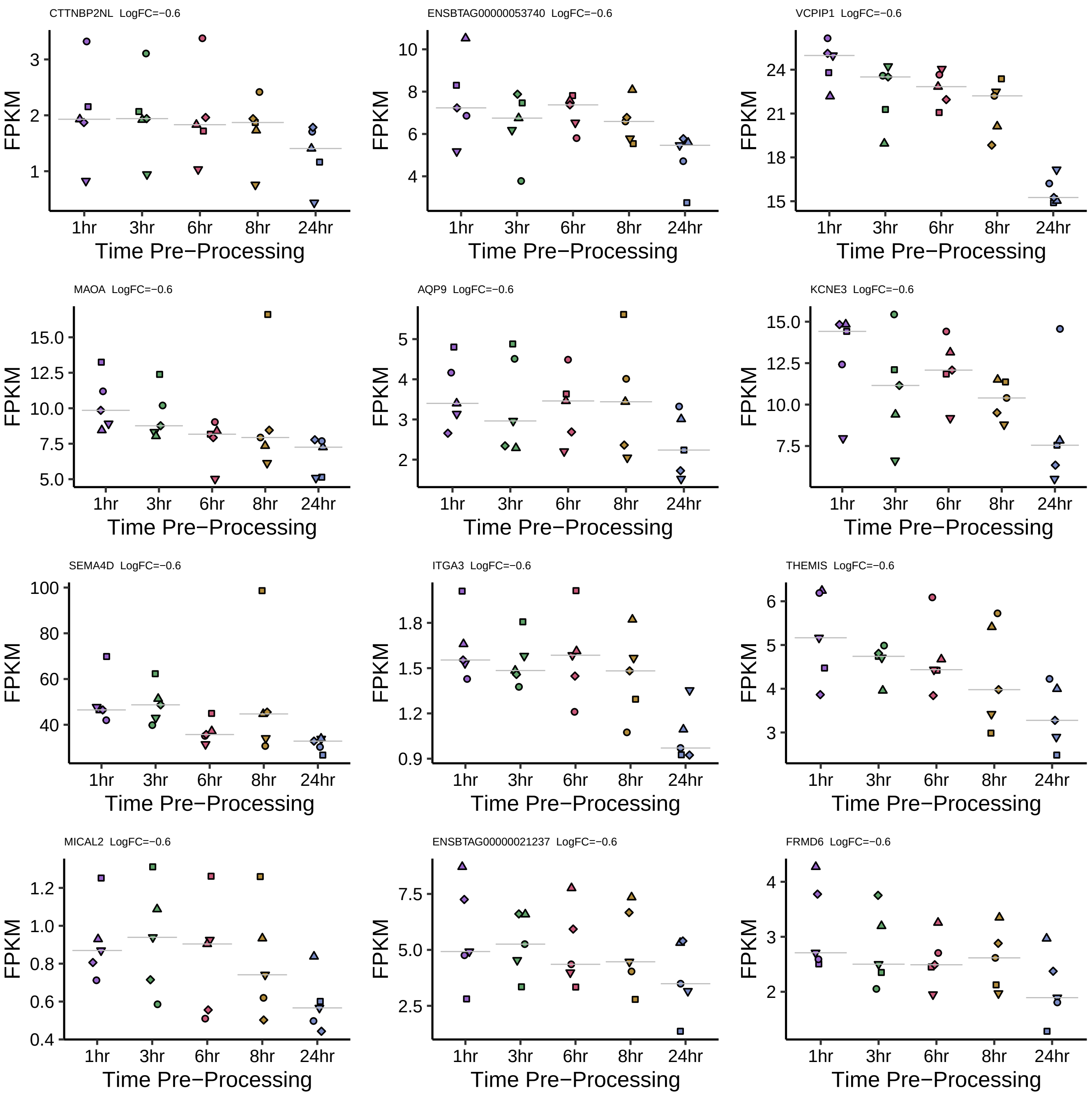
CTTNBP2NL LogFC=−0.6
ENSBTAG00000053740 LogFC=−0.6
VCPIP1 LogFC=−0.6
MAOA LogFC=−0.6
AQP9 LogFC=−0.6
KCNE3 LogFC=−0.6
SEMA4D LogFC=−0.6
ITGA3 LogFC=−0.6
THEMIS LogFC=−0.6
MICAL2 LogFC=−0.6
ENSBTAG00000021237 LogFC=−0.6
FRMD6 LogFC=−0.6
FPKM
1hr
3hr
6hr
8hr
24hr
Time Pre−Processing

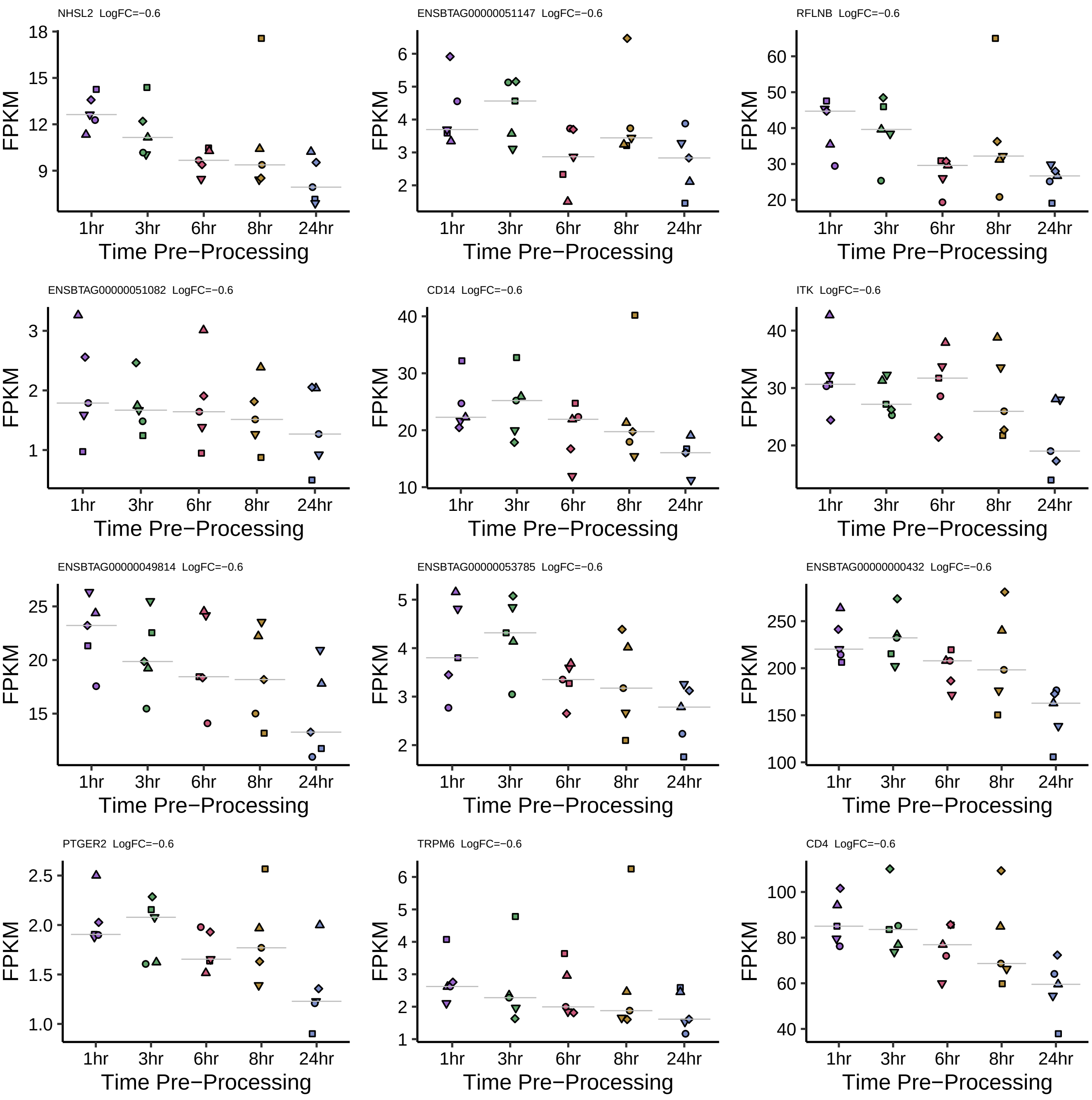

NHSL2 LogFC=−0.6
FPKM
18
15
12
9
1hr
3hr
6hr
8hr
24hr
Time Pre−Processing
ENSBTAG00000051147 LogFC=−0.6
FPKM
6
5
4
3
2
1hr
3hr
6hr
8hr
24hr
Time Pre−Processing
RFLNB LogFC=−0.6
FPKM
60
50
40
30
20
1hr
3hr
6hr
8hr
24hr
Time Pre−Processing
ENSBTAG00000051082 LogFC=−0.6
FPKM
3
2
1
1hr
3hr
6hr
8hr
24hr
Time Pre−Processing
CD14 LogFC=−0.6
FPKM
40
30
20
10
1hr
3hr
6hr
8hr
24hr
Time Pre−Processing
ITK LogFC=−0.6
FPKM
40
30
20
1hr
3hr
6hr
8hr
24hr
Time Pre−Processing
ENSBTAG00000049814 LogFC=−0.6
FPKM
25
20
15
1hr
3hr
6hr
8hr
24hr
Time Pre−Processing
ENSBTAG00000053785 LogFC=−0.6
FPKM
5
4
3
2
1hr
3hr
6hr
8hr
24hr
Time Pre−Processing
ENSBTAG00000000432 LogFC=−0.6
FPKM
250
200
150
100
1hr
3hr
6hr
8hr
24hr
Time Pre−Processing
PTGER2 LogFC=−0.6
FPKM
2.5
2.0
1.5
1.0
1hr
3hr
6hr
8hr
24hr
Time Pre−Processing
TRPM6 LogFC=−0.6
FPKM
6
5
4
3
2
1
1hr
3hr
6hr
8hr
24hr
Time Pre−Processing
CD4 LogFC=−0.6
FPKM
100
80
60
40
1hr
3hr
6hr
8hr
24hr
Time Pre−Processing

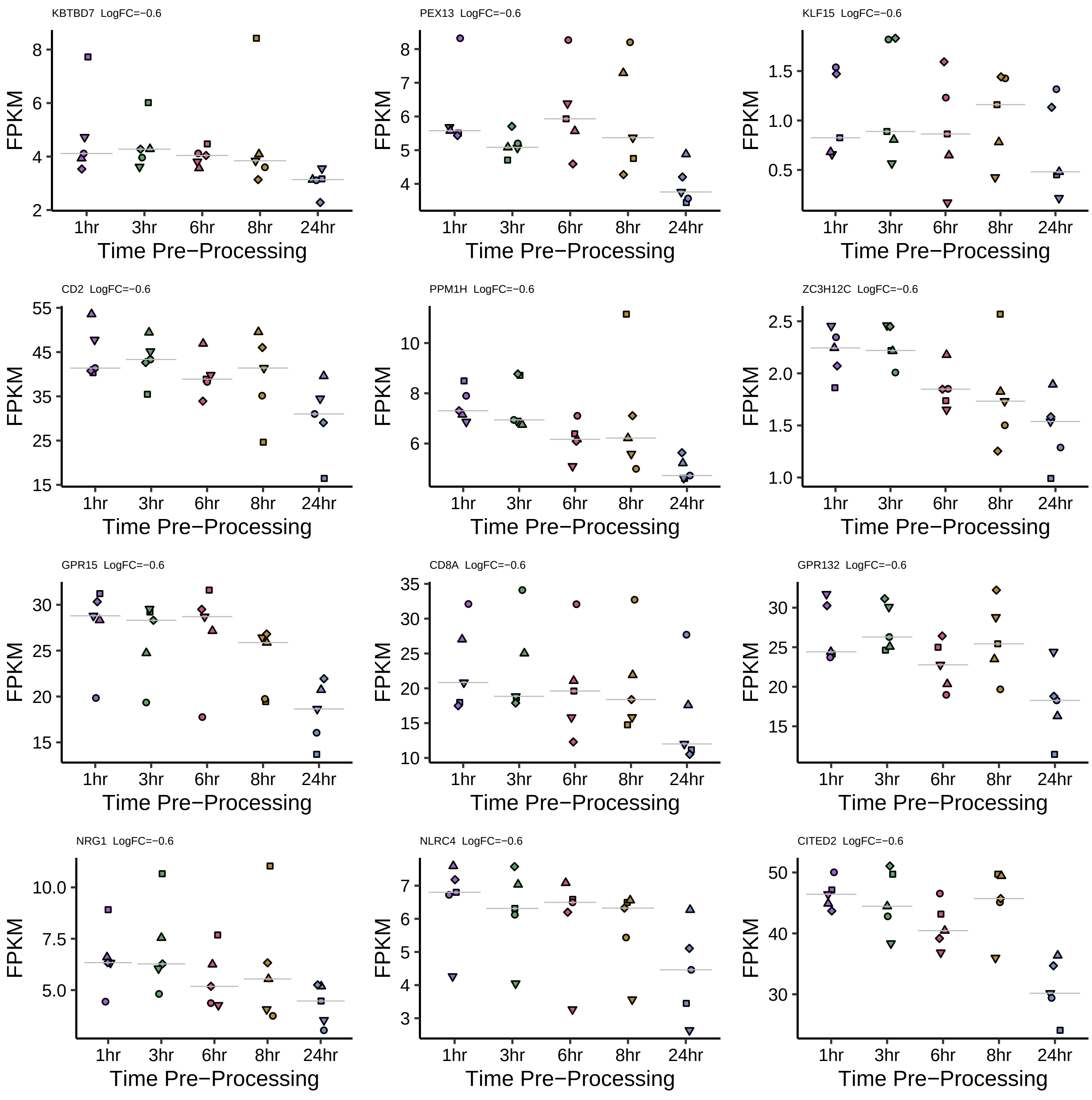

KBTBD7 LogFC=−0.6
PEX13 LogFC=−0.6
KLF15 LogFC=−0.6
CD2 LogFC=−0.6
PPM1H LogFC=−0.6
ZC3H12C LogFC=−0.6
GPR15 LogFC=−0.6
CD8A LogFC=−0.6
GPR132 LogFC=−0.6
NRG1 LogFC=−0.6
NLRC4 LogFC=−0.6
CITED2 LogFC=−0.6
FPKM
1hr
3hr
6hr
8hr
24hr
Time Pre−Processing

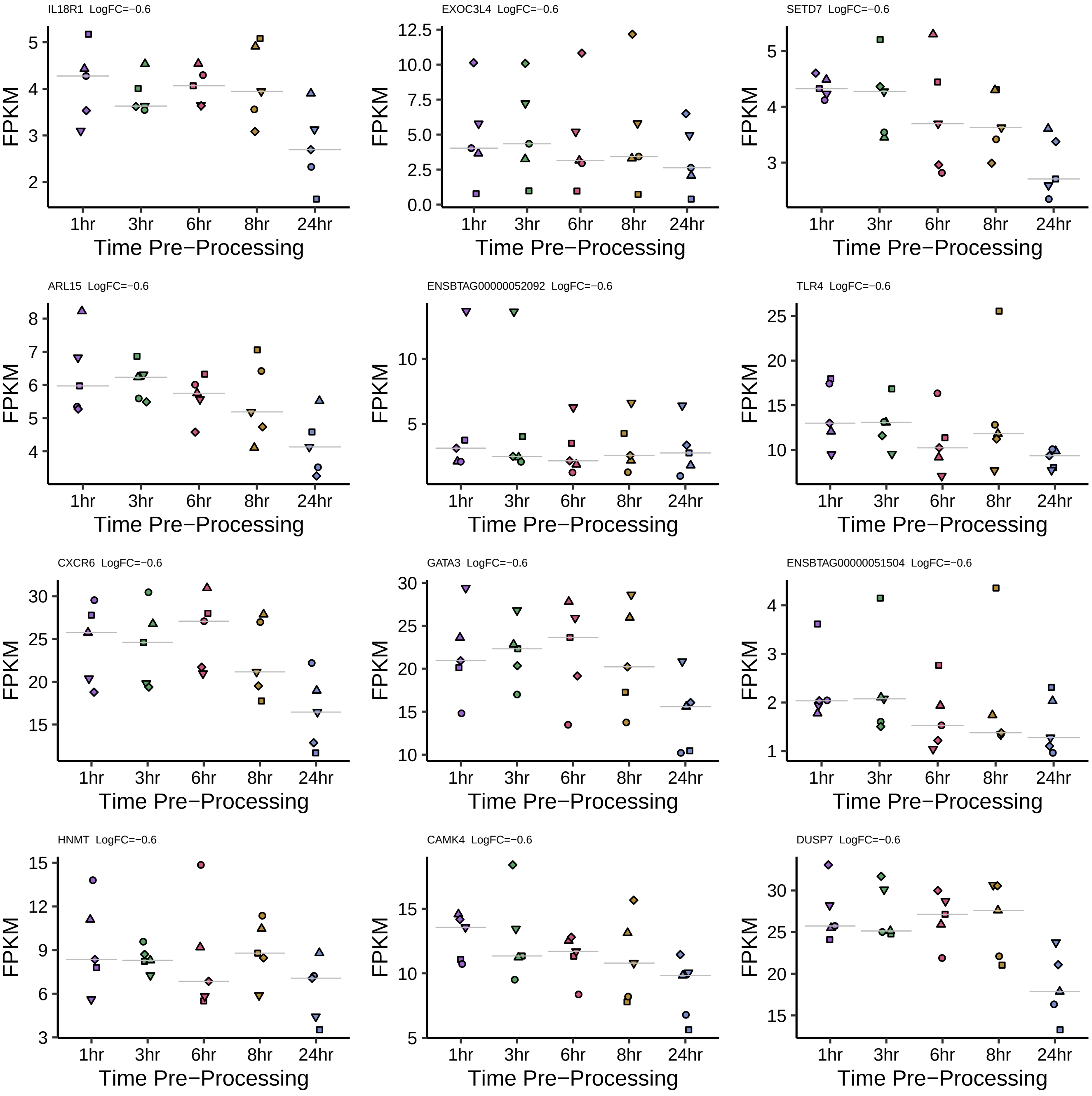
IL18R1 LogFC=−0.6
EXOC3L4 LogFC=−0.6
SETD7 LogFC=−0.6
ARL15 LogFC=−0.6
ENSBTAG00000052092 LogFC=−0.6
TLR4 LogFC=−0.6
CXCR6 LogFC=−0.6
GATA3 LogFC=−0.6
ENSBTAG00000051504 LogFC=−0.6
HNMT LogFC=−0.6
CAMK4 LogFC=−0.6
DUSP7 LogFC=−0.6
FPKM
1hr
3hr
6hr
8hr
24hr
Time Pre−Processing

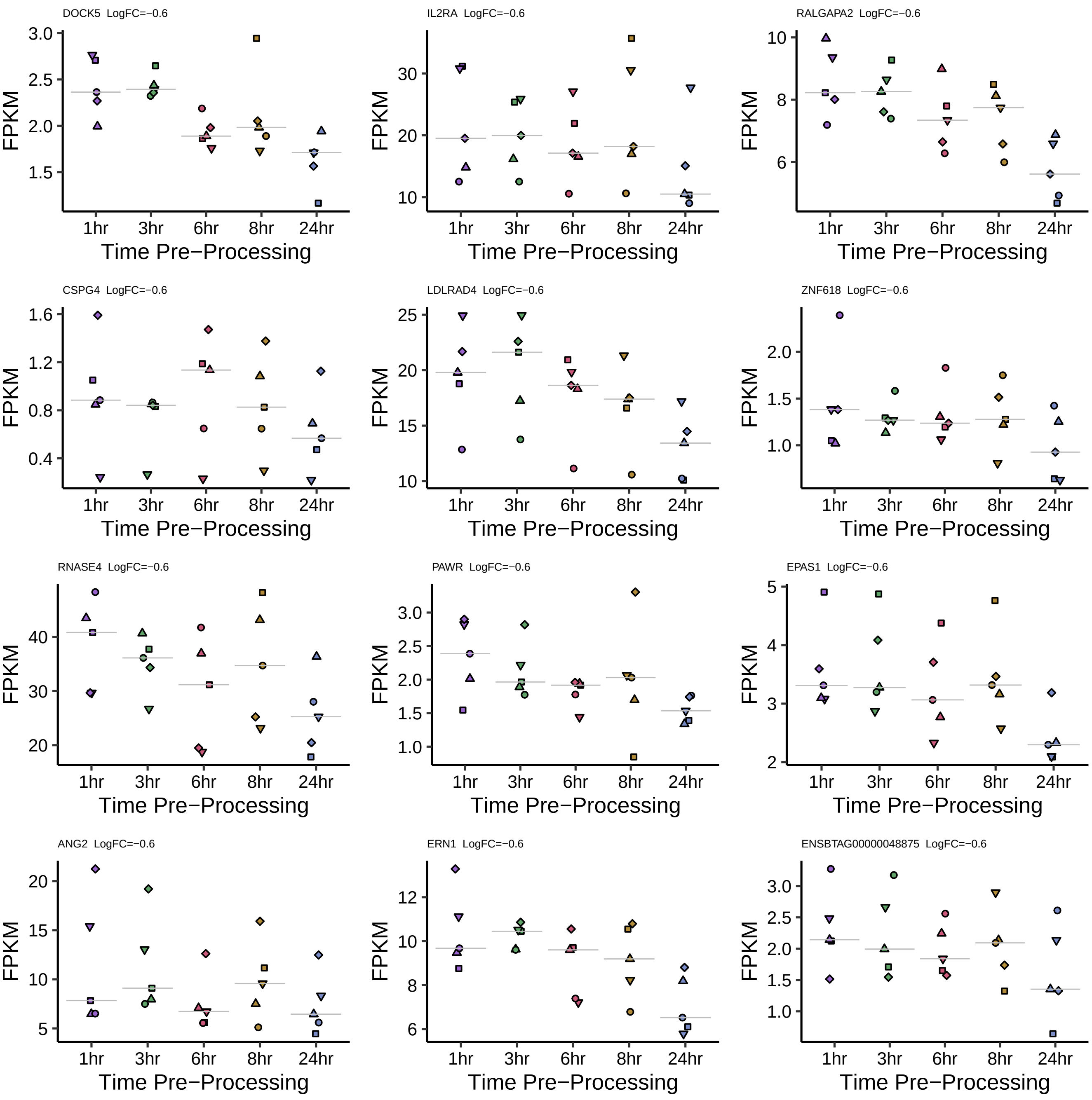
DOCK5 LogFC=−0.6
IL2RA LogFC=−0.6
RALGAPA2 LogFC=−0.6
CSPG4 LogFC=−0.6
LDLRAD4 LogFC=−0.6
ZNF618 LogFC=−0.6
RNASE4 LogFC=−0.6
PAWR LogFC=−0.6
EPAS1 LogFC=−0.6
ANG2 LogFC=−0.6
ERN1 LogFC=−0.6
ENSBTAG00000048875 LogFC=−0.6
FPKM
1hr
3hr
6hr
8hr
24hr
Time Pre−Processing

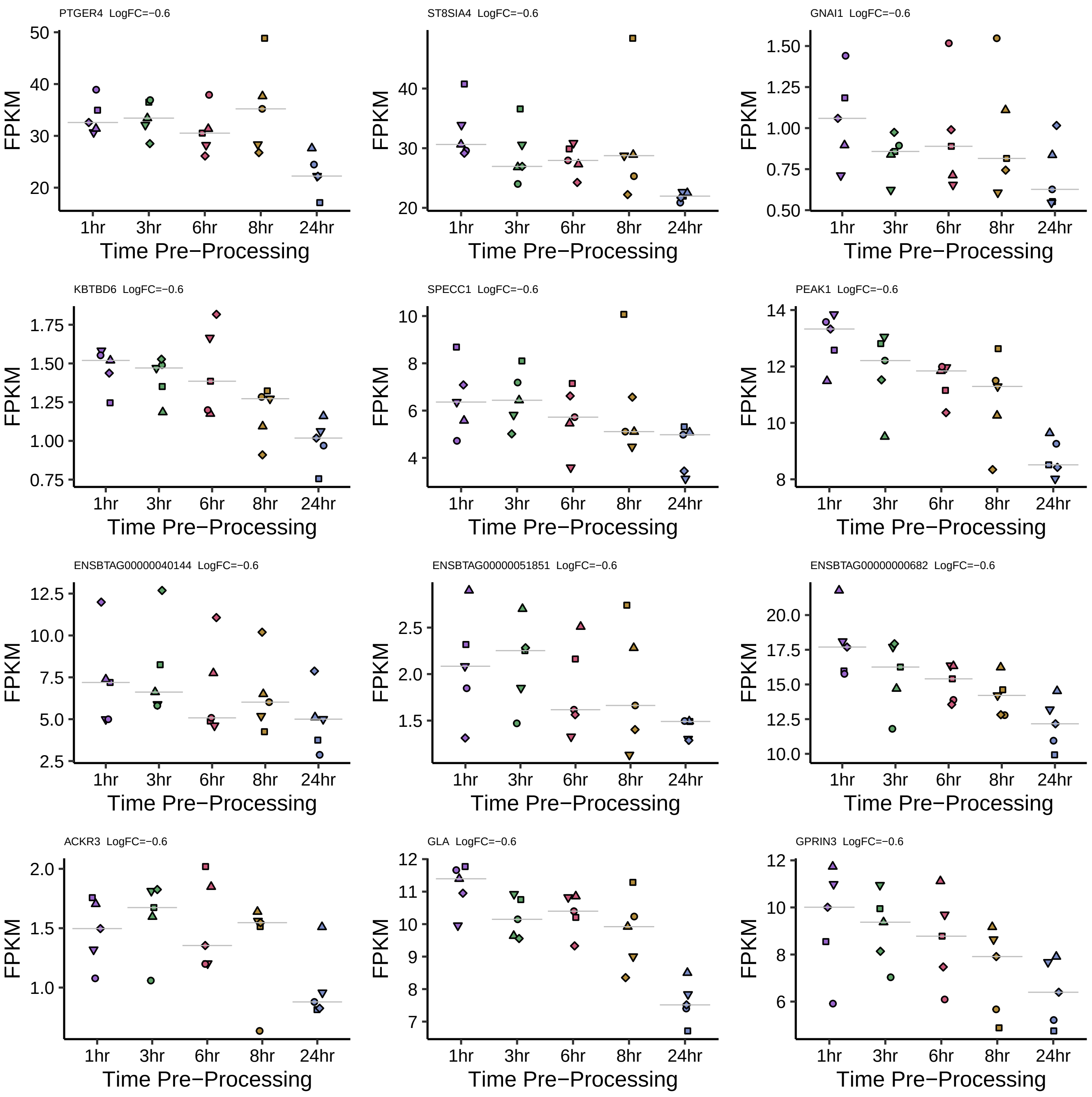

PTGER4 LogFC=−0.6
ST8SIA4 LogFC=−0.6
GNAI1 LogFC=−0.6
KBTBD6 LogFC=−0.6
SPECC1 LogFC=−0.6
PEAK1 LogFC=−0.6
ENSBTAG00000040144 LogFC=−0.6
ENSBTAG00000051851 LogFC=−0.6
ENSBTAG00000000682 LogFC=−0.6
ACKR3 LogFC=−0.6
GLA LogFC=−0.6
GPRIN3 LogFC=−0.6
FPKM
1hr
3hr
6hr
8hr
24hr
Time Pre−Processing

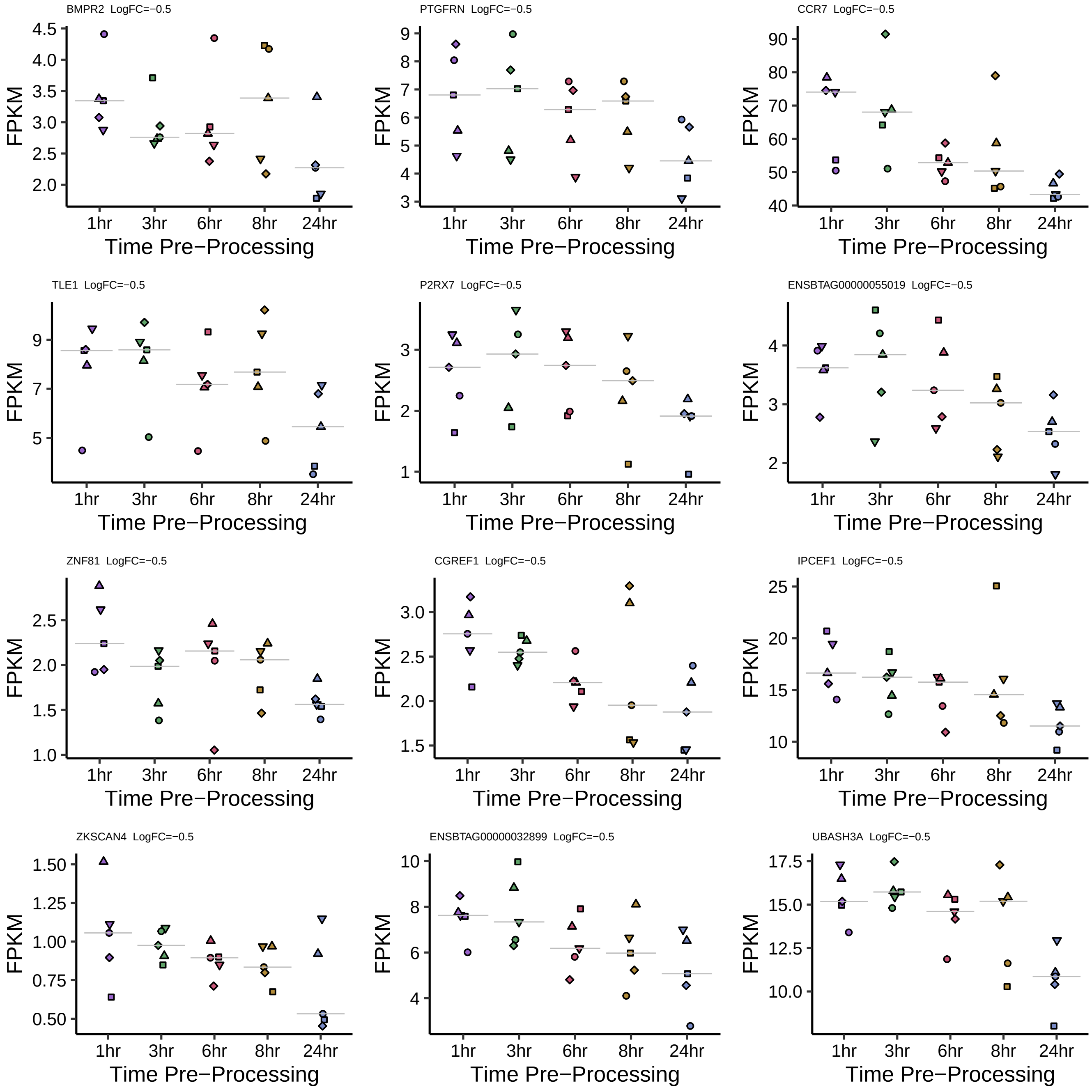
BMPR2 LogFC=−0.5
PTGFRN LogFC=−0.5
CCR7 LogFC=−0.5
TLE1 LogFC=−0.5
P2RX7 LogFC=−0.5
ENSBTAG00000055019 LogFC=−0.5
ZNF81 LogFC=−0.5
CGREF1 LogFC=−0.5
IPCEF1 LogFC=−0.5
ZKSCAN4 LogFC=−0.5
ENSBTAG00000032899 LogFC=−0.5
UBASH3A LogFC=−0.5
FPKM
1hr
3hr
6hr
8hr
24hr
Time Pre−Processing

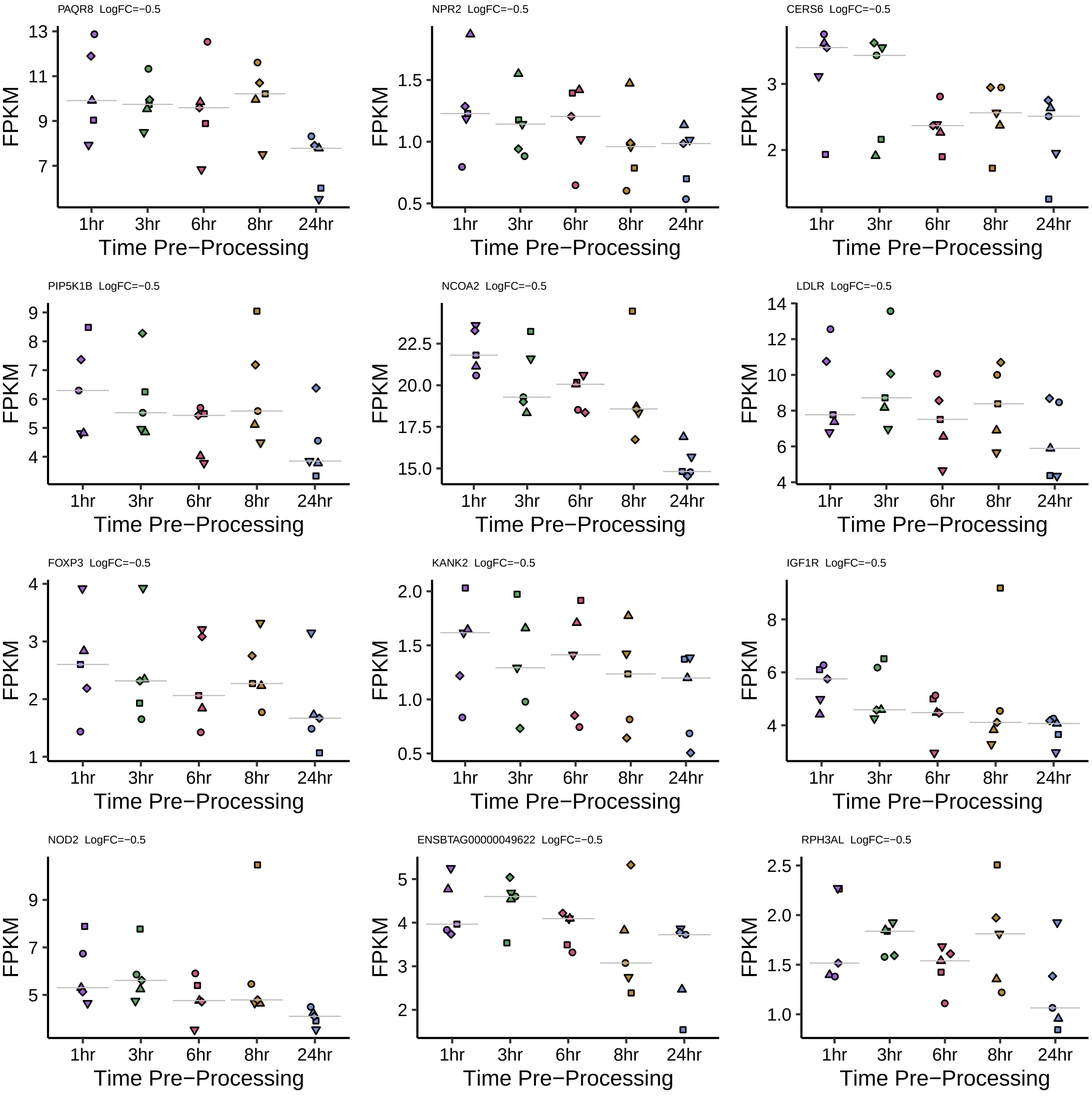
PAQR8 LogFC=−0.5
13
11
9
7
FPKM
1hr 3hr 6hr 8hr 24hr
Time Pre−Processing
NPR2 LogFC=−0.5
1.5
1.0
0.5
FPKM
1hr 3hr 6hr 8hr 24hr
Time Pre−Processing
CERS6 LogFC=−0.5
3
2
FPKM
1hr 3hr 6hr 8hr 24hr
Time Pre−Processing
PIP5K1B LogFC=−0.5
9
8
7
6
5
4
FPKM
1hr 3hr 6hr 8hr 24hr
Time Pre−Processing
NCOA2 LogFC=−0.5
22.5
20.0
17.5
15.0
FPKM
1hr 3hr 6hr 8hr 24hr
Time Pre−Processing
LDLR LogFC=−0.5
14
12
10
8
6
4
FPKM
1hr 3hr 6hr 8hr 24hr
Time Pre−Processing
FOXP3 LogFC=−0.5
4
3
2
1
FPKM
1hr 3hr 6hr 8hr 24hr
Time Pre−Processing
KANK2 LogFC=−0.5
2.0
1.5
1.0
0.5
FPKM
1hr 3hr 6hr 8hr 24hr
Time Pre−Processing
IGF1R LogFC=−0.5
8
6
4
FPKM
1hr 3hr 6hr 8hr 24hr
Time Pre−Processing
NOD2 LogFC=−0.5
9
7
5
FPKM
1hr 3hr 6hr 8hr 24hr
Time Pre−Processing
ENSBTAG00000049622 LogFC=−0.5
5
4
3
2
FPKM
1hr 3hr 6hr 8hr 24hr
Time Pre−Processing
RPH3AL LogFC=−0.5
2.5
2.0
1.5
1.0
FPKM
1hr 3hr 6hr 8hr 24hr
Time Pre−Processing

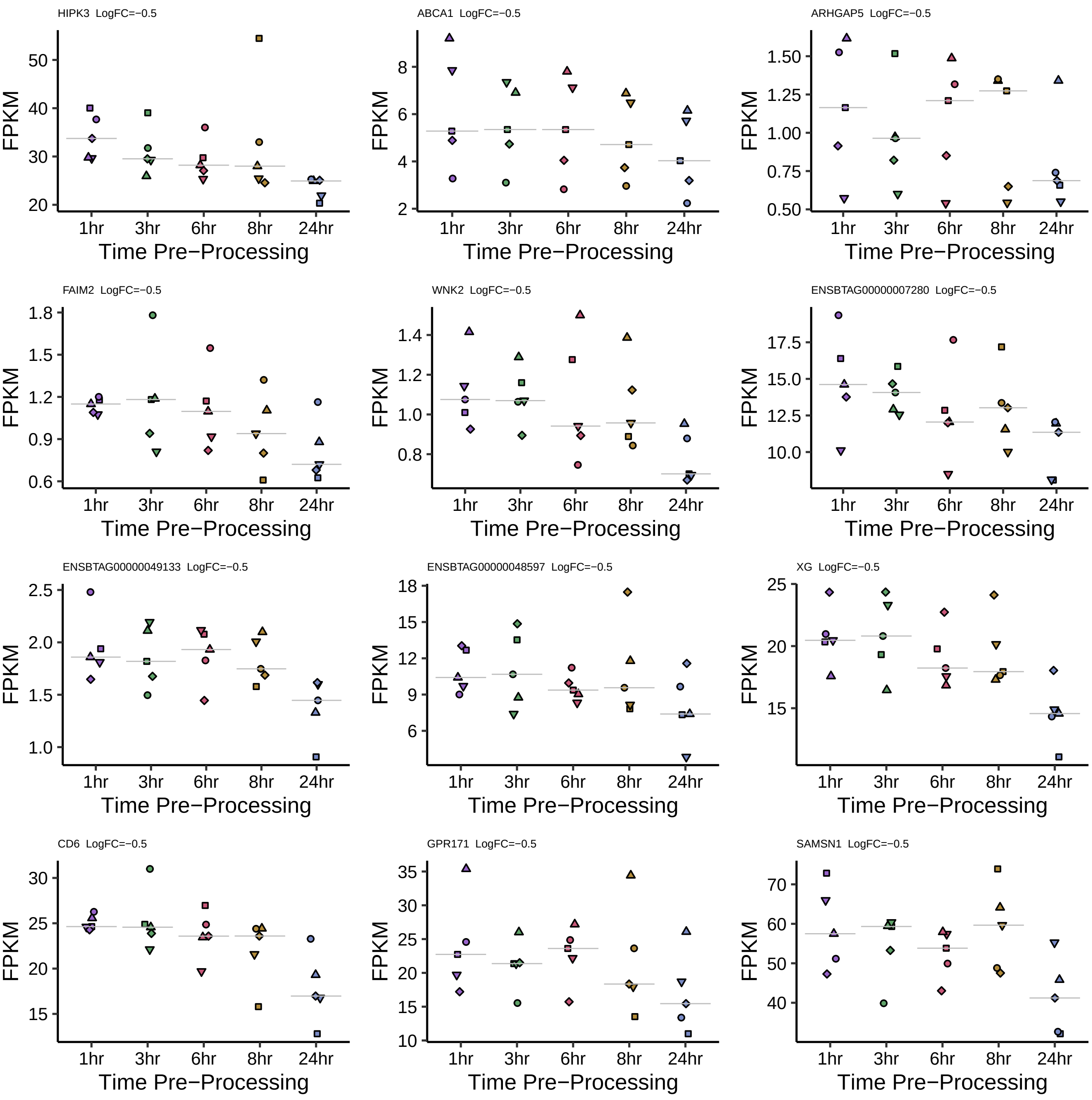
HIPK3 LogFC=−0.5
ABCA1 LogFC=−0.5
ARHGAP5 LogFC=−0.5
FAIM2 LogFC=−0.5
WNK2 LogFC=−0.5
ENSBTAG00000007280 LogFC=−0.5
ENSBTAG00000049133 LogFC=−0.5
ENSBTAG00000048597 LogFC=−0.5
XG LogFC=−0.5
CD6 LogFC=−0.5
GPR171 LogFC=−0.5
SAMSN1 LogFC=−0.5
FPKM
1hr
3hr
6hr
8hr
24hr
Time Pre−Processing

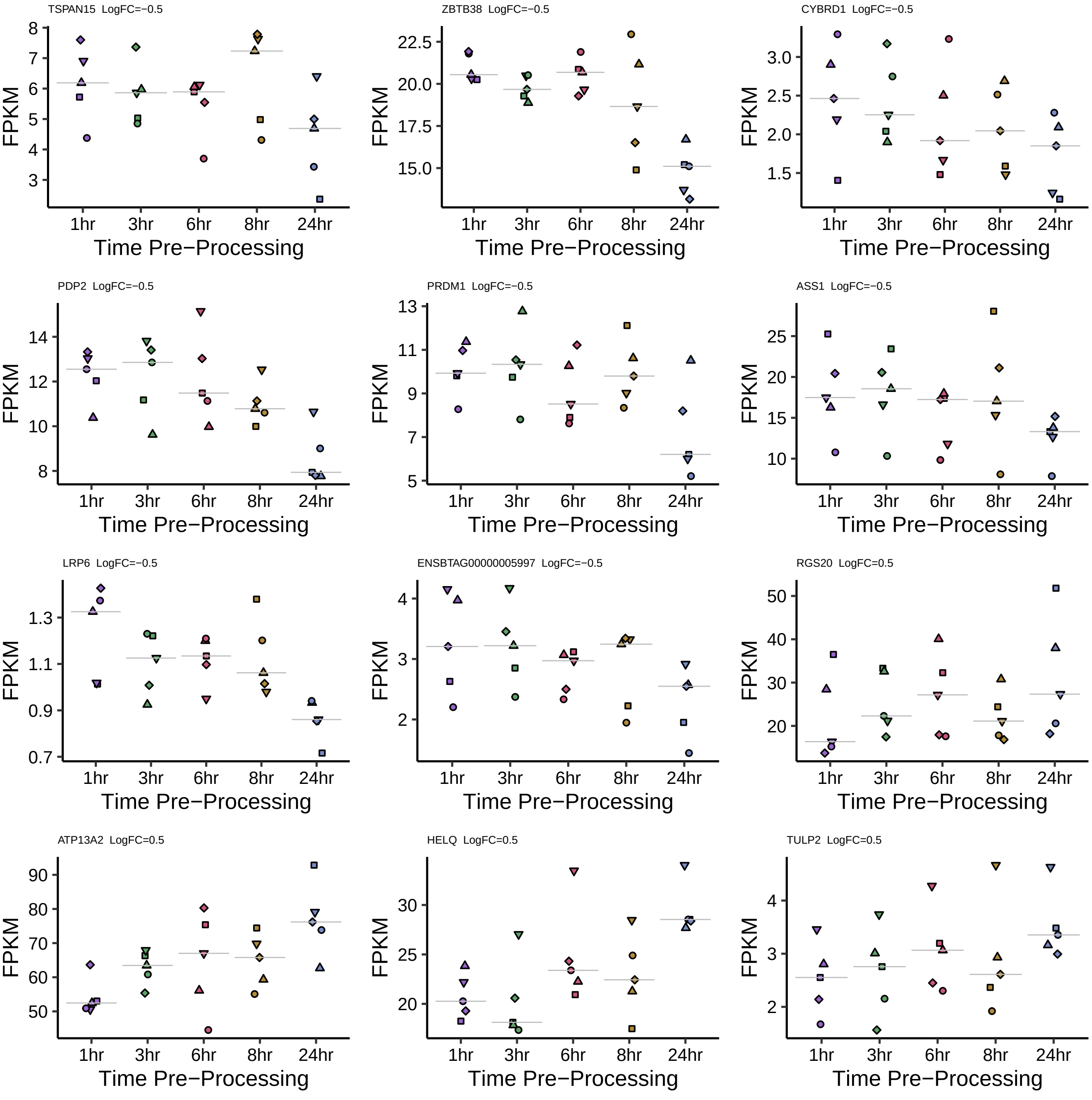
TSPAN15 LogFC=−0.5
ZBTB38 LogFC=−0.5
CYBRD1 LogFC=−0.5
PDP2 LogFC=−0.5
PRDM1 LogFC=−0.5
ASS1 LogFC=−0.5
LRP6 LogFC=−0.5
ENSBTAG00000005997 LogFC=−0.5
RGS20 LogFC=0.5
ATP13A2 LogFC=0.5
HELQ LogFC=0.5
TULP2 LogFC=0.5
FPKM
1hr
3hr
6hr
8hr
24hr
Time Pre−Processing

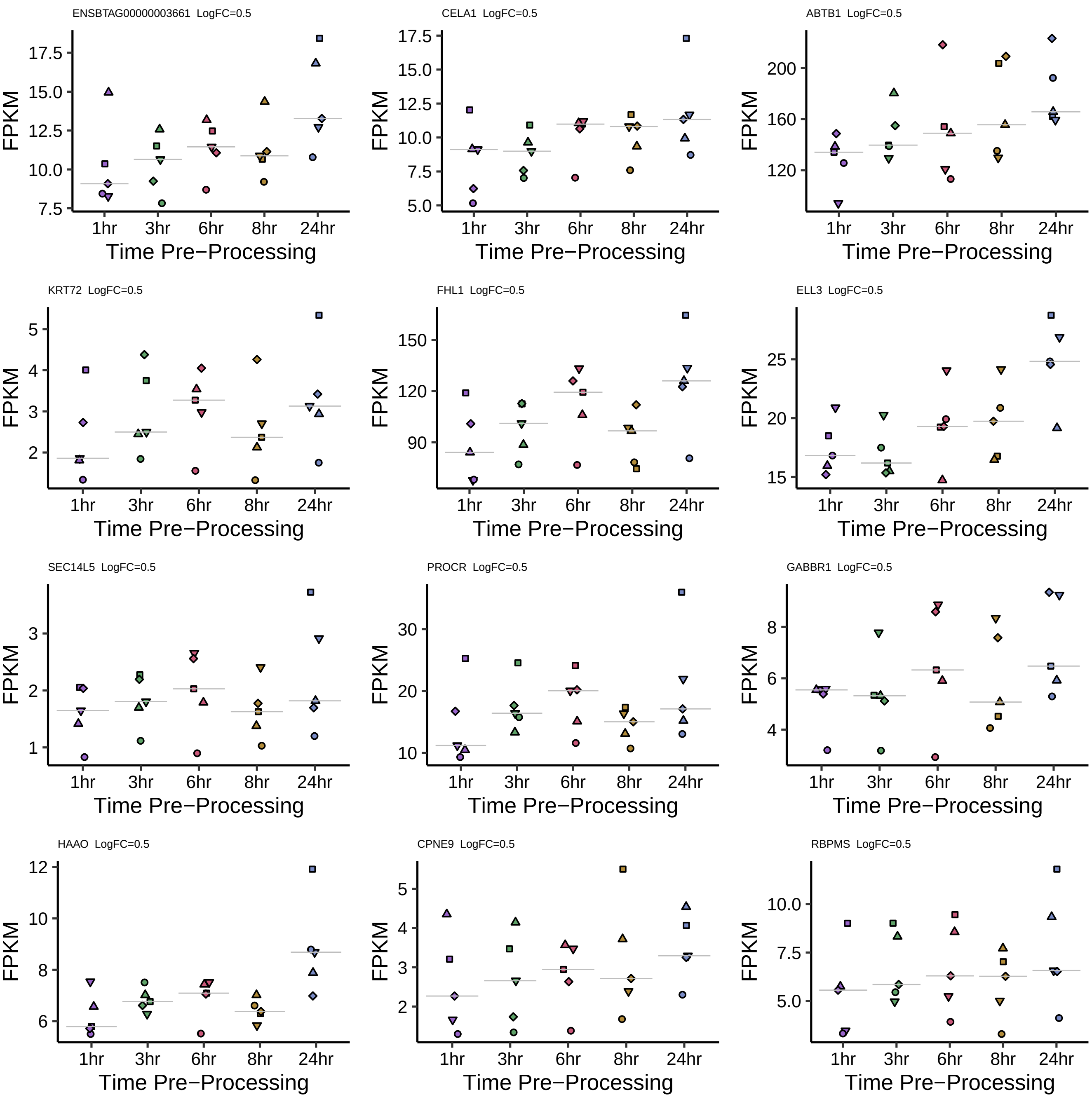
ENSBTAG00000003661 LogFC=0.5
CELA1 LogFC=0.5
ABTB1 LogFC=0.5
KRT72 LogFC=0.5
FHL1 LogFC=0.5
ELL3 LogFC=0.5
SEC14L5 LogFC=0.5
PROCR LogFC=0.5
GABBR1 LogFC=0.5
HAAO LogFC=0.5
CPNE9 LogFC=0.5
RBPMS LogFC=0.5
FPKM
1hr
3hr
6hr
8hr
24hr
Time Pre−Processing

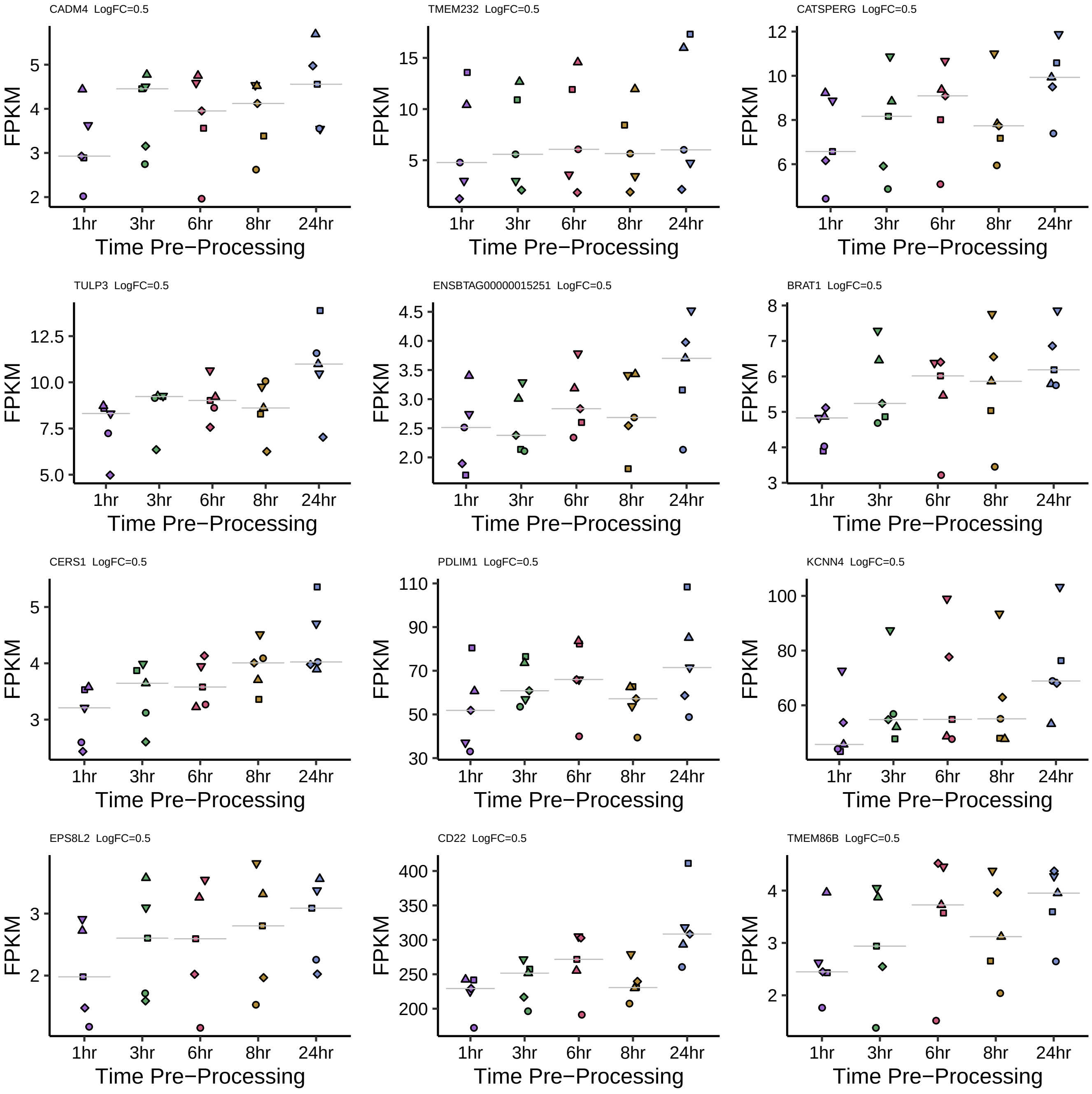
CADM4 LogFC=0.5
TMEM232 LogFC=0.5
CATSPERG LogFC=0.5
TULP3 LogFC=0.5
ENSBTAG00000015251 LogFC=0.5
BRAT1 LogFC=0.5
CERS1 LogFC=0.5
PDLIM1 LogFC=0.5
KCNN4 LogFC=0.5
EPS8L2 LogFC=0.5
CD22 LogFC=0.5
TMEM86B LogFC=0.5
FPKM
1hr
3hr
6hr
8hr
24hr
Time Pre−Processing

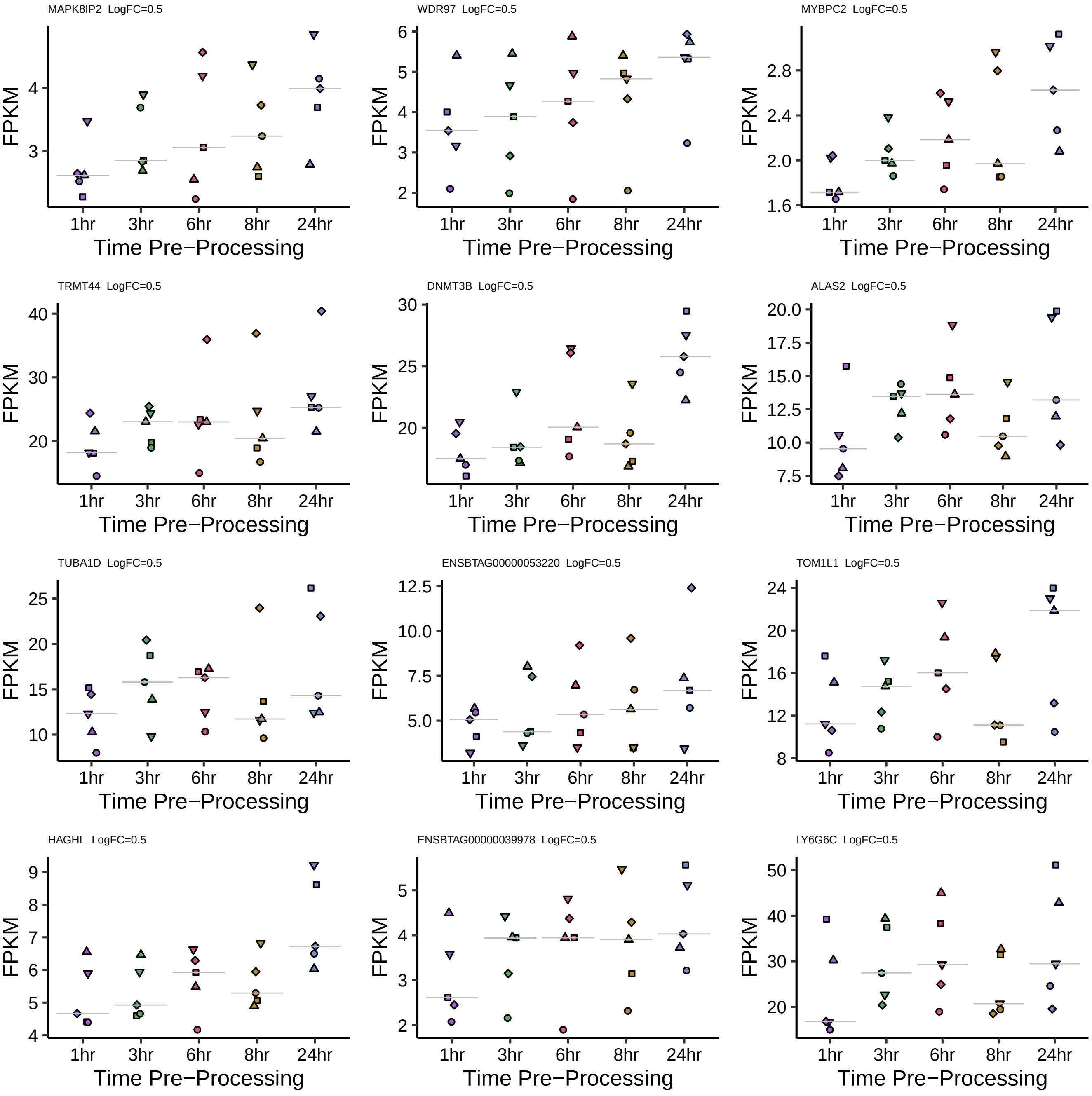
MAPK8IP2 LogFC=0.5
WDR97 LogFC=0.5
MYBPC2 LogFC=0.5
TRMT44 LogFC=0.5
DNMT3B LogFC=0.5
ALAS2 LogFC=0.5
TUBA1D LogFC=0.5
ENSBTAG00000053220 LogFC=0.5
TOM1L1 LogFC=0.5
HAGHL LogFC=0.5
ENSBTAG00000039978 LogFC=0.5
LY6G6C LogFC=0.5
FPKM
1hr
3hr
6hr
8hr
24hr
Time Pre−Processing

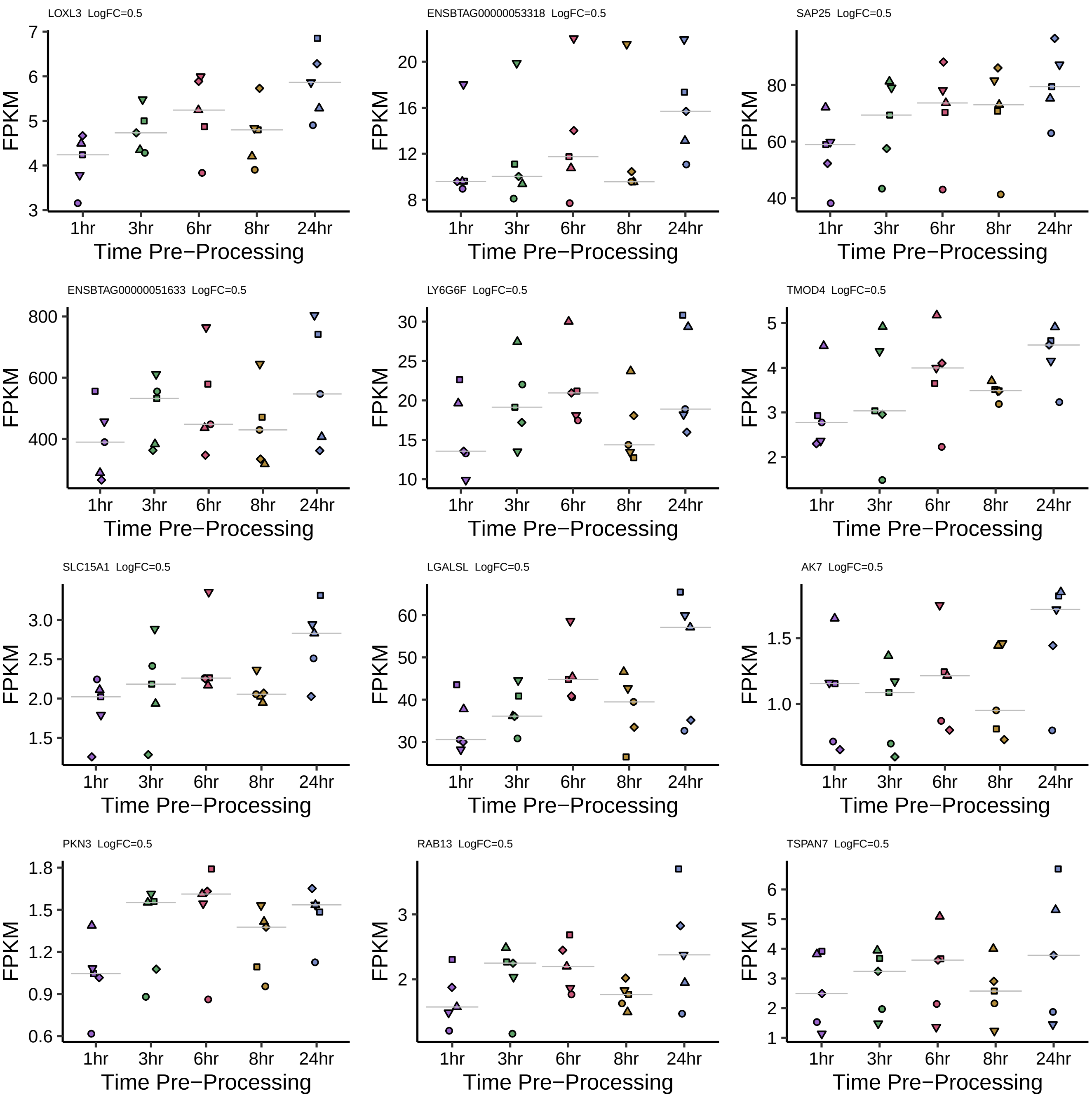
LOXL3 LogFC=0.5
ENSBTAG00000053318 LogFC=0.5
SAP25 LogFC=0.5
ENSBTAG00000051633 LogFC=0.5
LY6G6F LogFC=0.5
TMOD4 LogFC=0.5
SLC15A1 LogFC=0.5
LGALSL LogFC=0.5
AK7 LogFC=0.5
PKN3 LogFC=0.5
RAB13 LogFC=0.5
TSPAN7 LogFC=0.5
FPKM
1hr
3hr
6hr
8hr
24hr
Time Pre−Processing

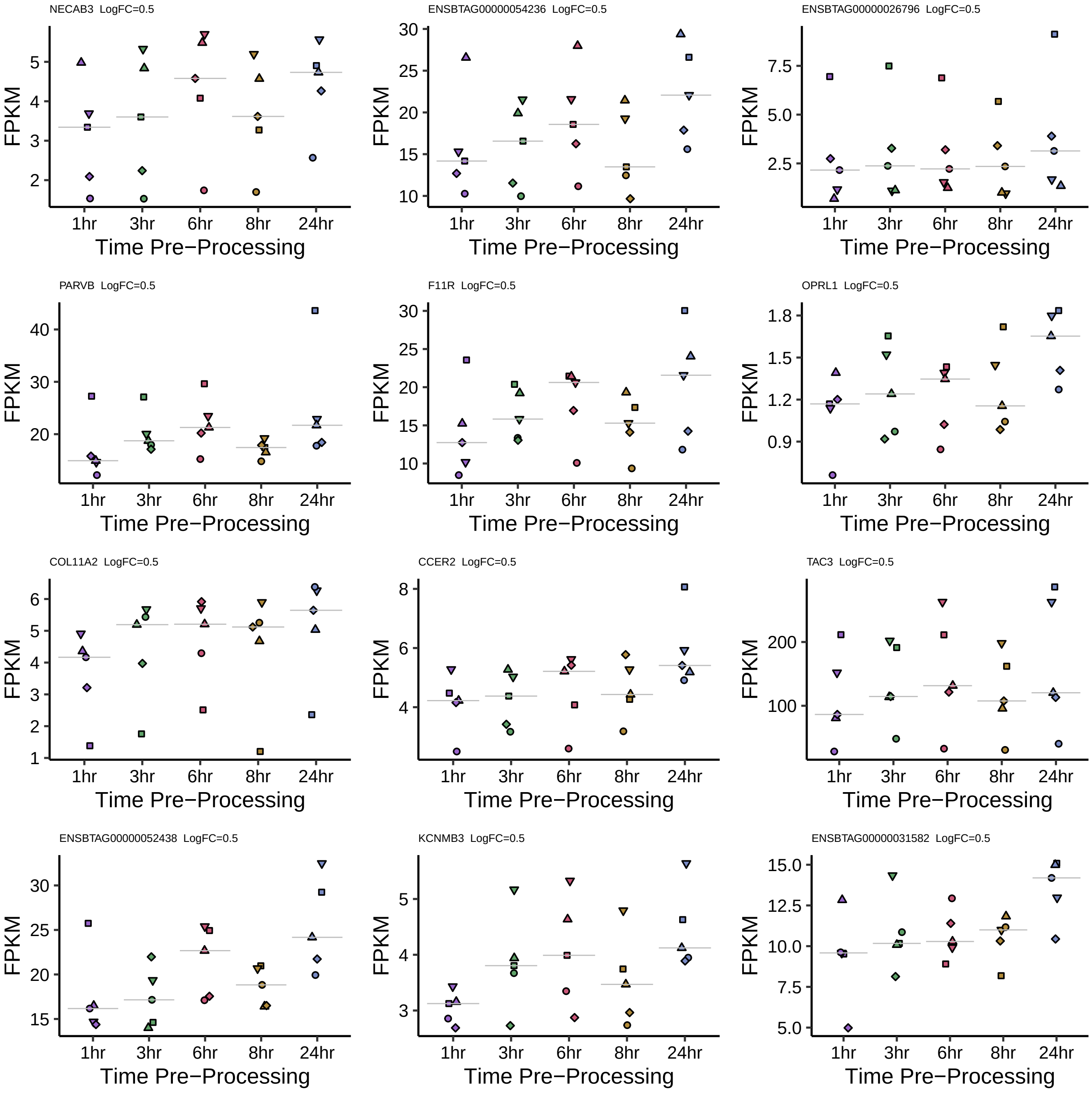

NECAB3 LogFC=0.5
ENSBTAG00000054236 LogFC=0.5
ENSBTAG00000026796 LogFC=0.5
PARVB LogFC=0.5
F11R LogFC=0.5
OPRL1 LogFC=0.5
COL11A2 LogFC=0.5
CCER2 LogFC=0.5
TAC3 LogFC=0.5
ENSBTAG00000052438 LogFC=0.5
KCNMB3 LogFC=0.5
ENSBTAG00000031582 LogFC=0.5
FPKM
Time Pre−Processing
1hr
3hr
6hr
8hr
24hr

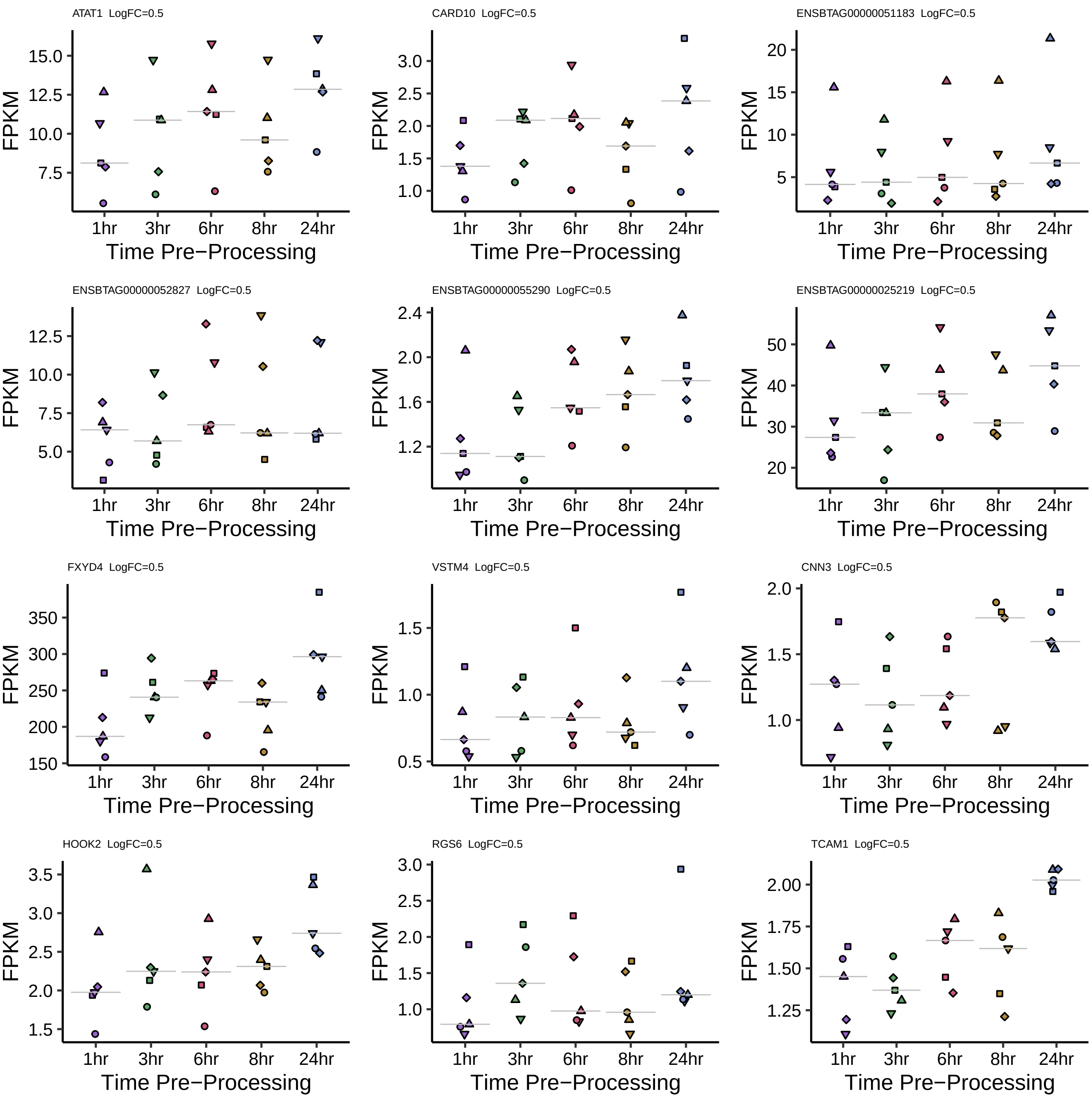
ATAT1 LogFC=0.5
CARD10 LogFC=0.5
ENSBTAG00000051183 LogFC=0.5
ENSBTAG00000052827 LogFC=0.5
ENSBTAG00000055290 LogFC=0.5
ENSBTAG00000025219 LogFC=0.5
FXYD4 LogFC=0.5
VSTM4 LogFC=0.5
CNN3 LogFC=0.5
HOOK2 LogFC=0.5
RGS6 LogFC=0.5
TCAM1 LogFC=0.5
FPKM
1hr
3hr
6hr
8hr
24hr
Time Pre−Processing

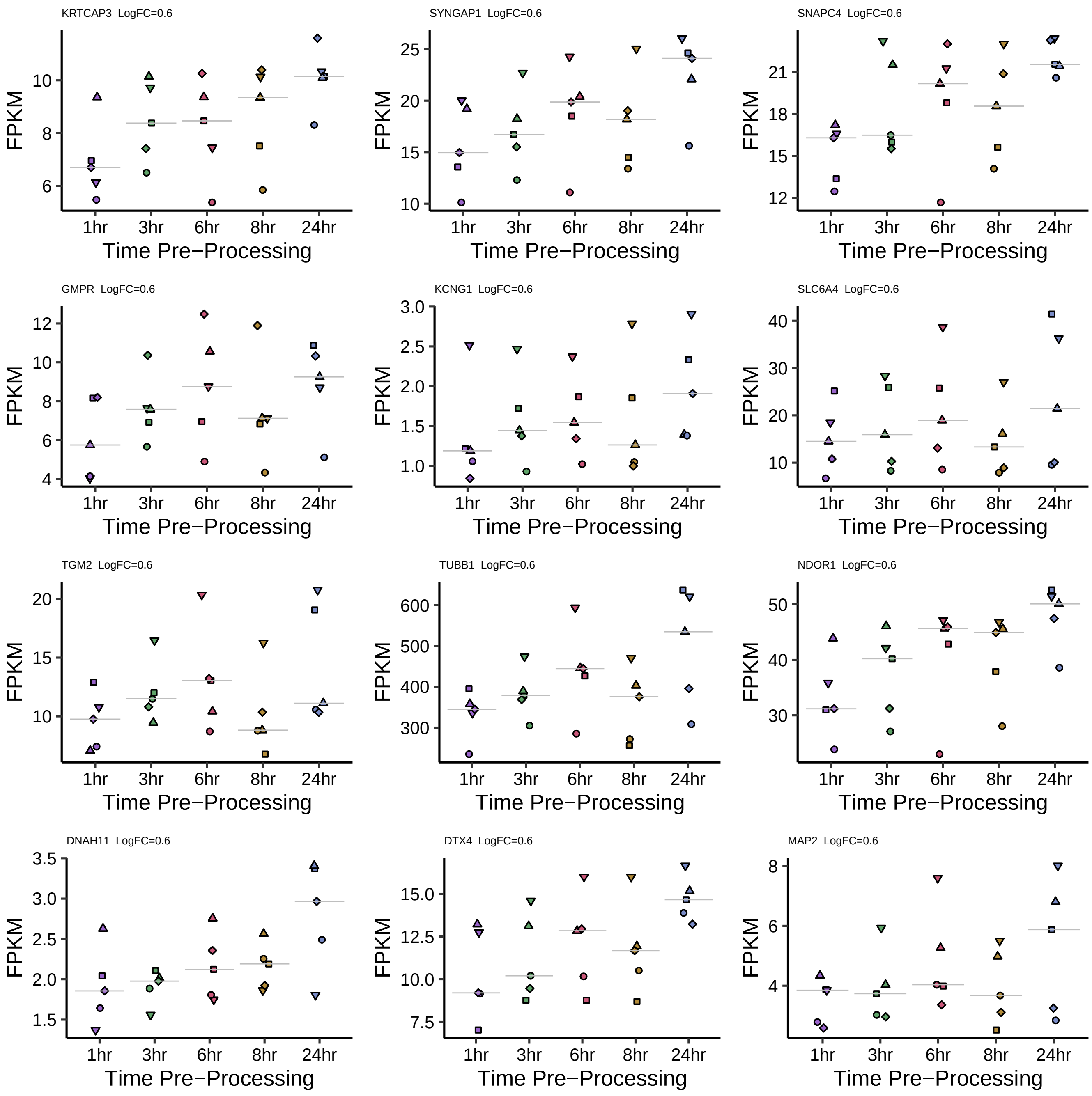

KRTCAP3 LogFC=0.6
SYNGAP1 LogFC=0.6
SNAPC4 LogFC=0.6
GMPR LogFC=0.6
KCNG1 LogFC=0.6
SLC6A4 LogFC=0.6
TGM2 LogFC=0.6
TUBB1 LogFC=0.6
NDOR1 LogFC=0.6
DNAH11 LogFC=0.6
DTX4 LogFC=0.6
MAP2 LogFC=0.6
FPKM
1hr
3hr
6hr
8hr
24hr
Time Pre−Processing

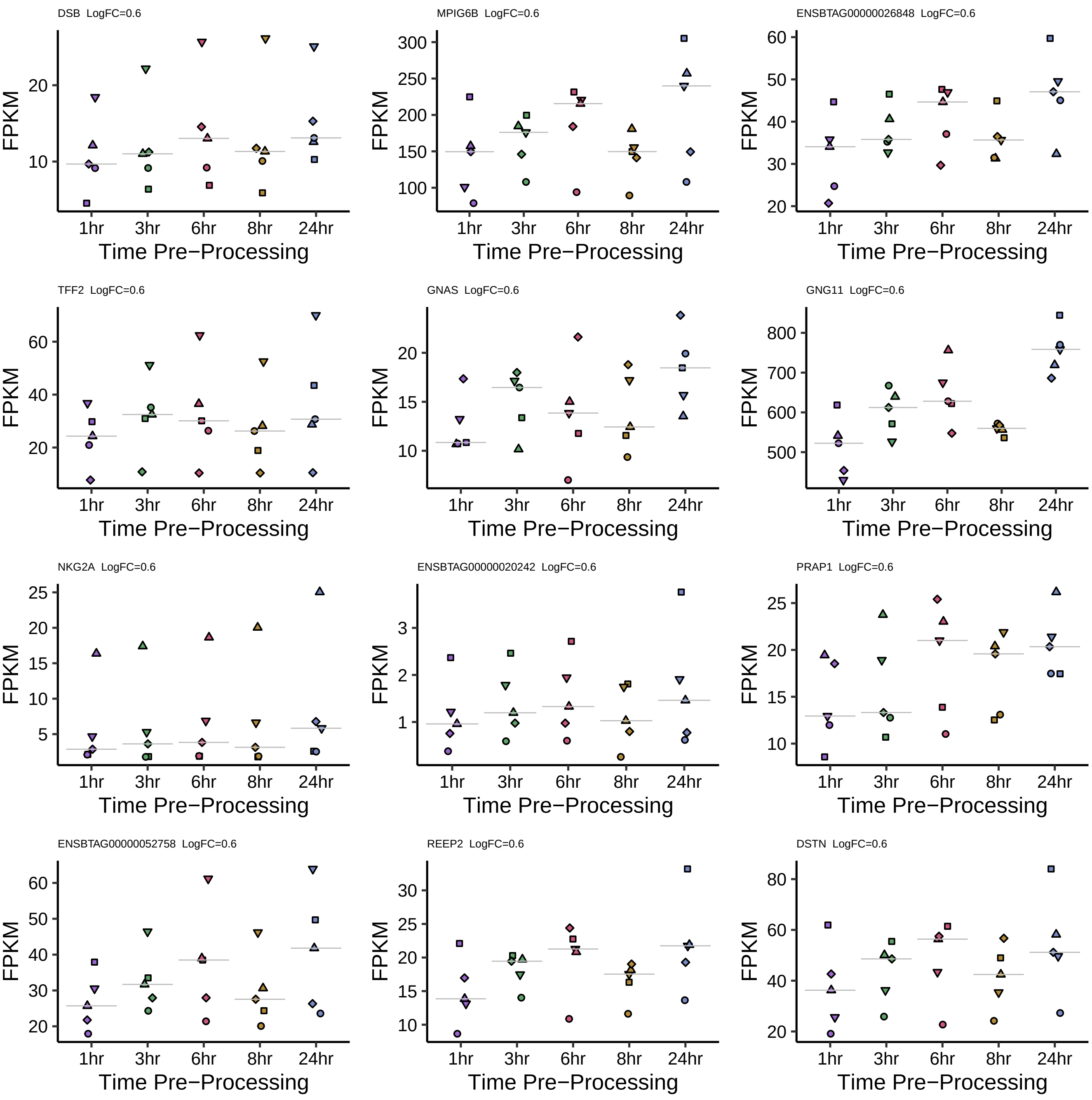

DSB LogFC=0.6
MPIG6B LogFC=0.6
ENSBTAG00000026848 LogFC=0.6
TFF2 LogFC=0.6
GNAS LogFC=0.6
GNG11 LogFC=0.6
NKG2A LogFC=0.6
ENSBTAG00000020242 LogFC=0.6
PRAP1 LogFC=0.6
ENSBTAG00000052758 LogFC=0.6
REEP2 LogFC=0.6
DSTN LogFC=0.6
FPKM
1hr
3hr
6hr
8hr
24hr
Time Pre−Processing

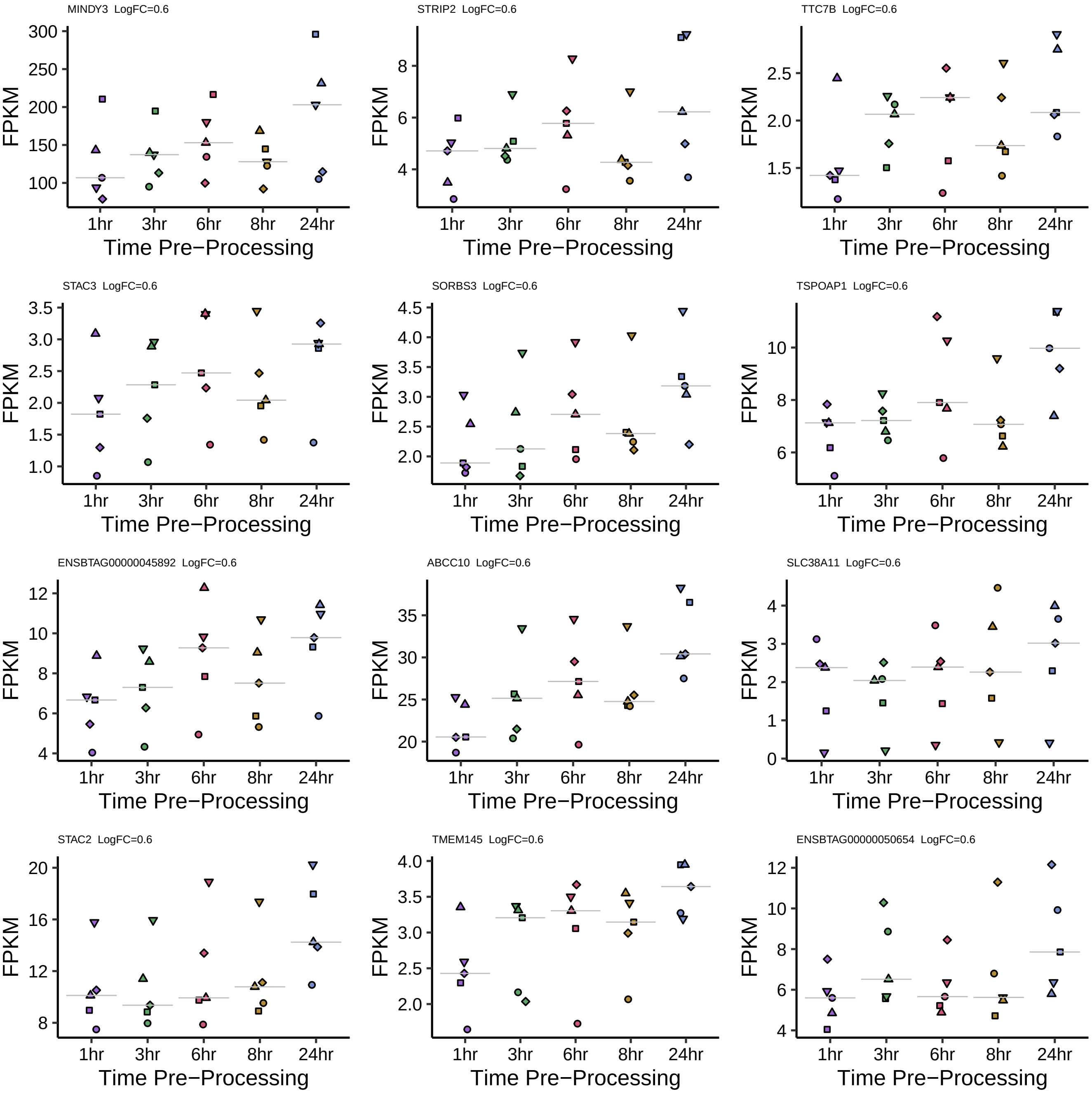

MINDY3 LogFC=0.6
300
250
200
150
100
FPKM
1hr
3hr
6hr
8hr
24hr
Time Pre−Processing
STRIP2 LogFC=0.6
8
6
4
FPKM
1hr
3hr
6hr
8hr
24hr
Time Pre−Processing
TTC7B LogFC=0.6
2.5
2.0
1.5
FPKM
1hr
3hr
6hr
8hr
24hr
Time Pre−Processing
STAC3 LogFC=0.6
3.5
3.0
2.5
2.0
1.5
1.0
FPKM
1hr
3hr
6hr
8hr
24hr
Time Pre−Processing
SORBS3 LogFC=0.6
4.5
4.0
3.5
3.0
2.5
2.0
FPKM
1hr
3hr
6hr
8hr
24hr
Time Pre−Processing
TSPOAP1 LogFC=0.6
10
8
6
FPKM
1hr
3hr
6hr
8hr
24hr
Time Pre−Processing
ENSBTAG00000045892 LogFC=0.6
12
10
8
6
4
FPKM
1hr
3hr
6hr
8hr
24hr
Time Pre−Processing
ABCC10 LogFC=0.6
35
30
25
20
FPKM
1hr
3hr
6hr
8hr
24hr
Time Pre−Processing
SLC38A11 LogFC=0.6
4
3
2
1
0
FPKM
1hr
3hr
6hr
8hr
24hr
Time Pre−Processing
STAC2 LogFC=0.6
20
16
12
8
FPKM
1hr
3hr
6hr
8hr
24hr
Time Pre−Processing
TMEM145 LogFC=0.6
4.0
3.5
3.0
2.5
2.0
FPKM
1hr
3hr
6hr
8hr
24hr
Time Pre−Processing
ENSBTAG00000050654 LogFC=0.6
12
10
8
6
4
FPKM
1hr
3hr
6hr
8hr
24hr
Time Pre−Processing

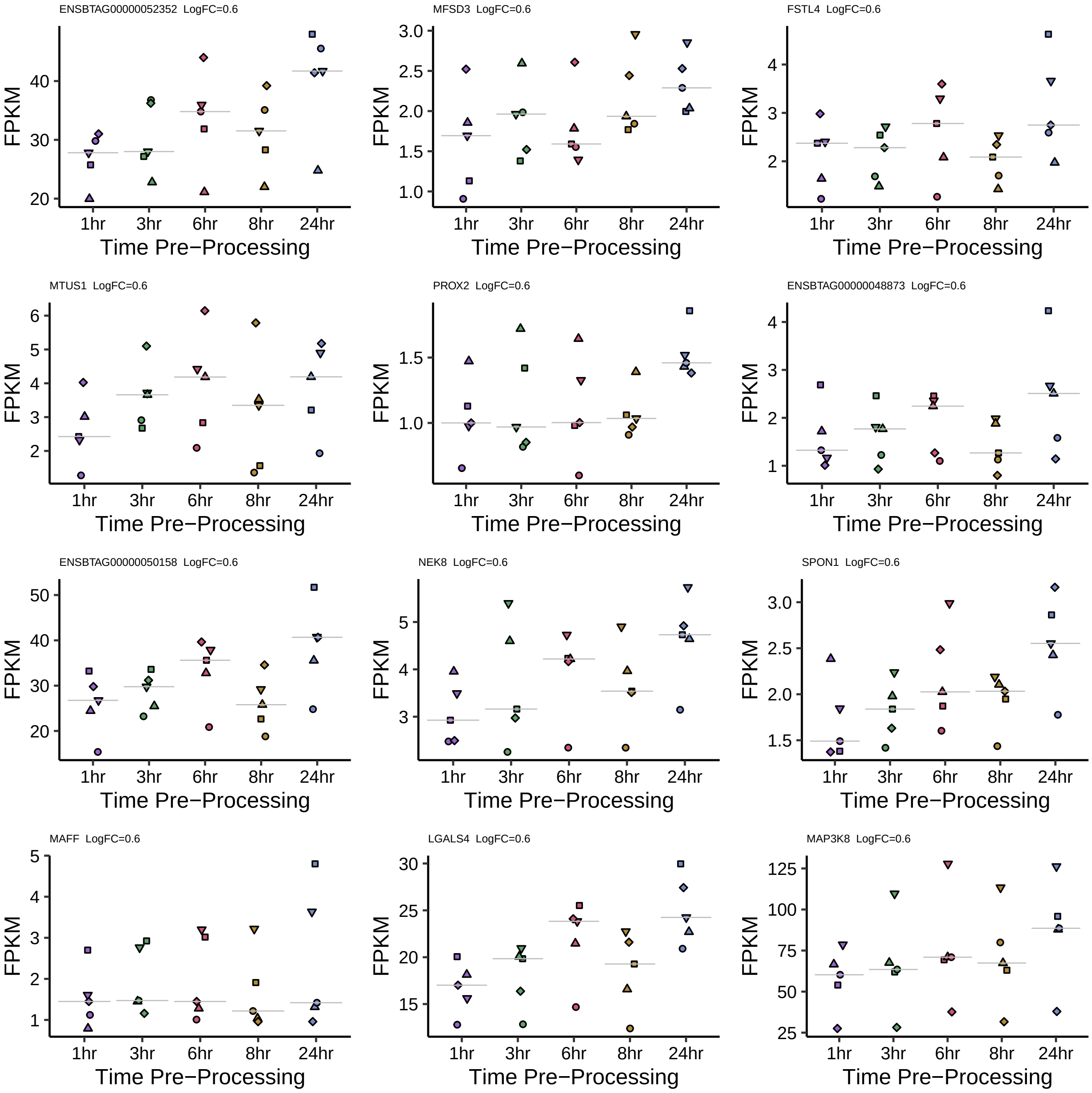
ENSBTAG00000052352 LogFC=0.6
MFSD3 LogFC=0.6
FSTL4 LogFC=0.6
MTUS1 LogFC=0.6
PROX2 LogFC=0.6
ENSBTAG00000048873 LogFC=0.6
ENSBTAG00000050158 LogFC=0.6
NEK8 LogFC=0.6
SPON1 LogFC=0.6
MAFF LogFC=0.6
LGALS4 LogFC=0.6
MAP3K8 LogFC=0.6
FPKM
1hr
3hr
6hr
8hr
24hr
Time Pre−Processing

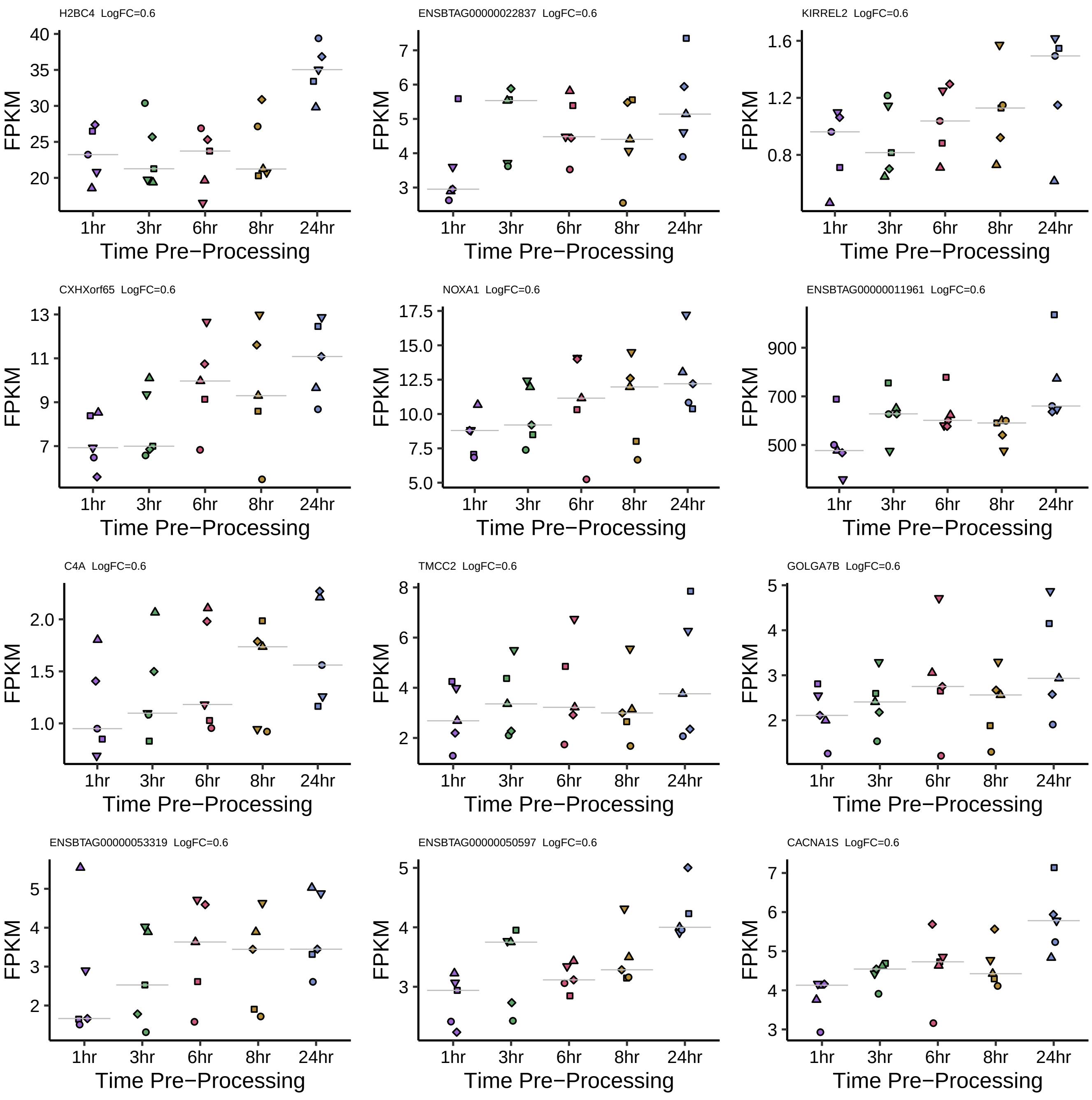
H2BC4 LogFC=0.6
ENSBTAG00000022837 LogFC=0.6
KIRREL2 LogFC=0.6
CXHXorf65 LogFC=0.6
NOXA1 LogFC=0.6
ENSBTAG00000011961 LogFC=0.6
C4A LogFC=0.6
TMCC2 LogFC=0.6
GOLGA7B LogFC=0.6
ENSBTAG00000053319 LogFC=0.6
ENSBTAG00000050597 LogFC=0.6
CACNA1S LogFC=0.6
FPKM
1hr
3hr
6hr
8hr
24hr
Time Pre−Processing

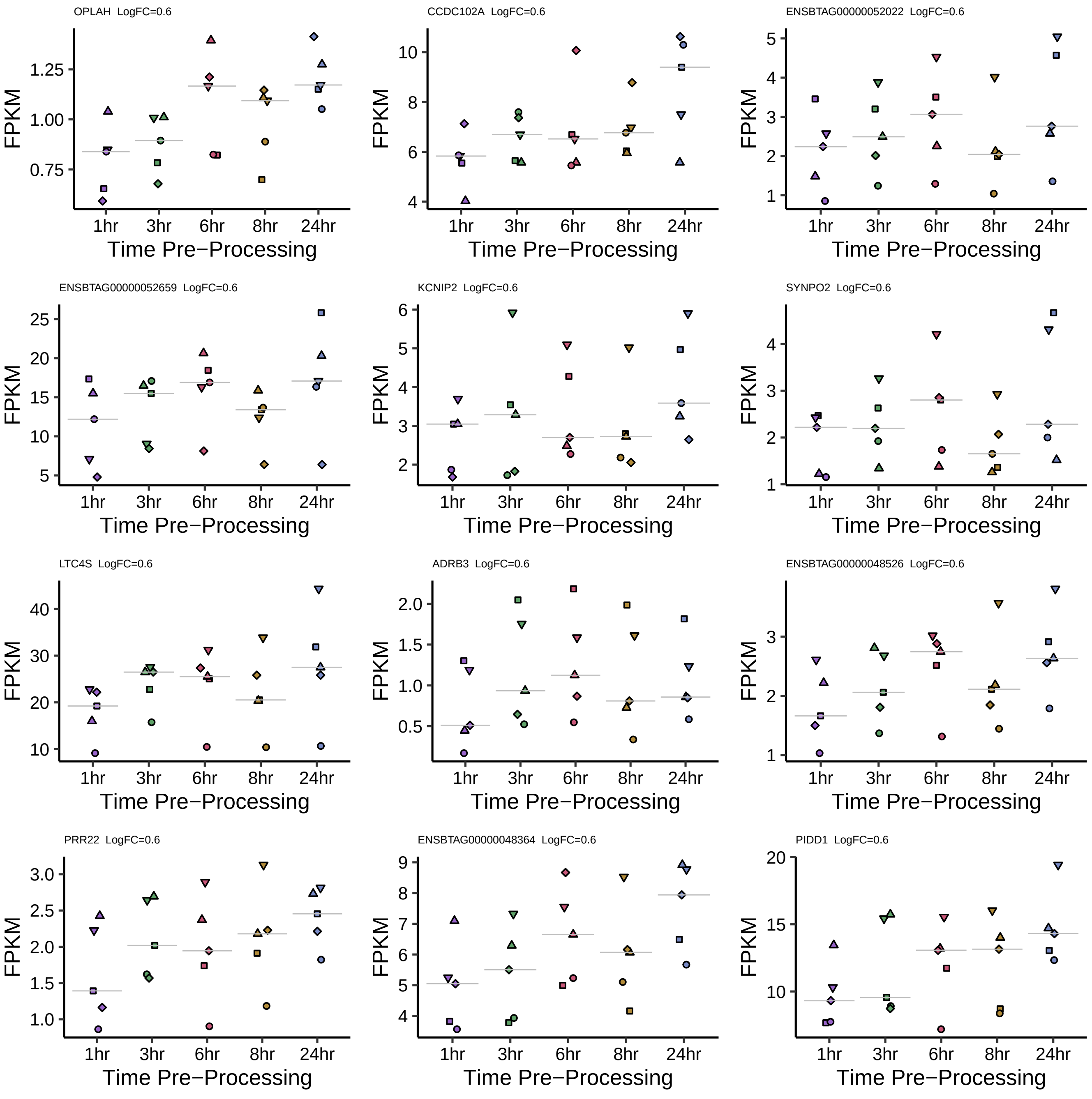
OPLAH LogFC=0.6
CCDC102A LogFC=0.6
ENSBTAG00000052022 LogFC=0.6
ENSBTAG00000052659 LogFC=0.6
KCNIP2 LogFC=0.6
SYNPO2 LogFC=0.6
LTC4S LogFC=0.6
ADRB3 LogFC=0.6
ENSBTAG00000048526 LogFC=0.6
PRR22 LogFC=0.6
ENSBTAG00000048364 LogFC=0.6
PIDD1 LogFC=0.6
FPKM
1hr
3hr
6hr
8hr
24hr
Time Pre−Processing

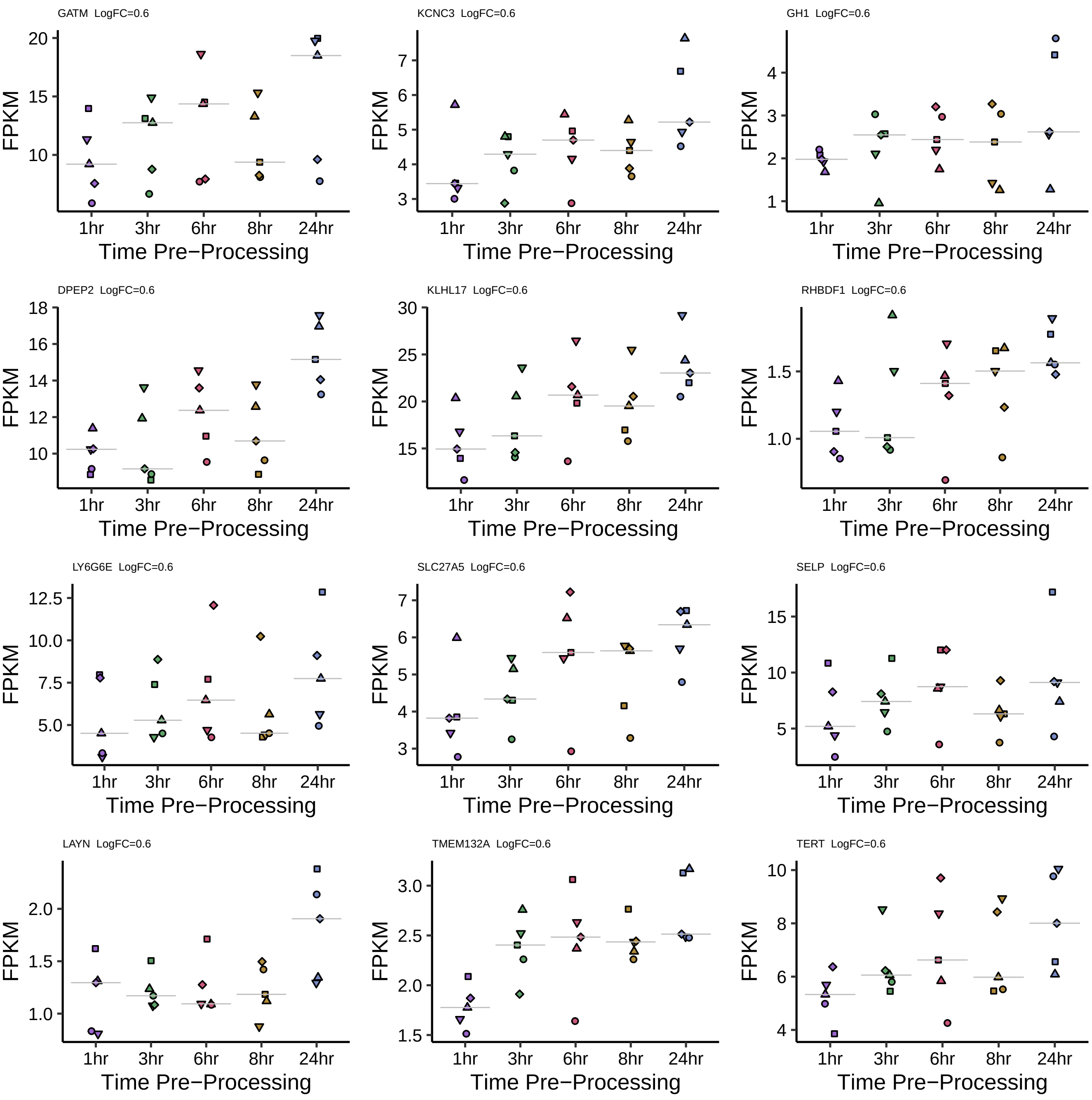
GATM LogFC=0.6
KCNC3 LogFC=0.6
GH1 LogFC=0.6
DPEP2 LogFC=0.6
KLHL17 LogFC=0.6
RHBDF1 LogFC=0.6
LY6G6E LogFC=0.6
SLC27A5 LogFC=0.6
SELP LogFC=0.6
LAYN LogFC=0.6
TMEM132A LogFC=0.6
TERT LogFC=0.6
FPKM
1hr
3hr
6hr
8hr
24hr
Time Pre−Processing

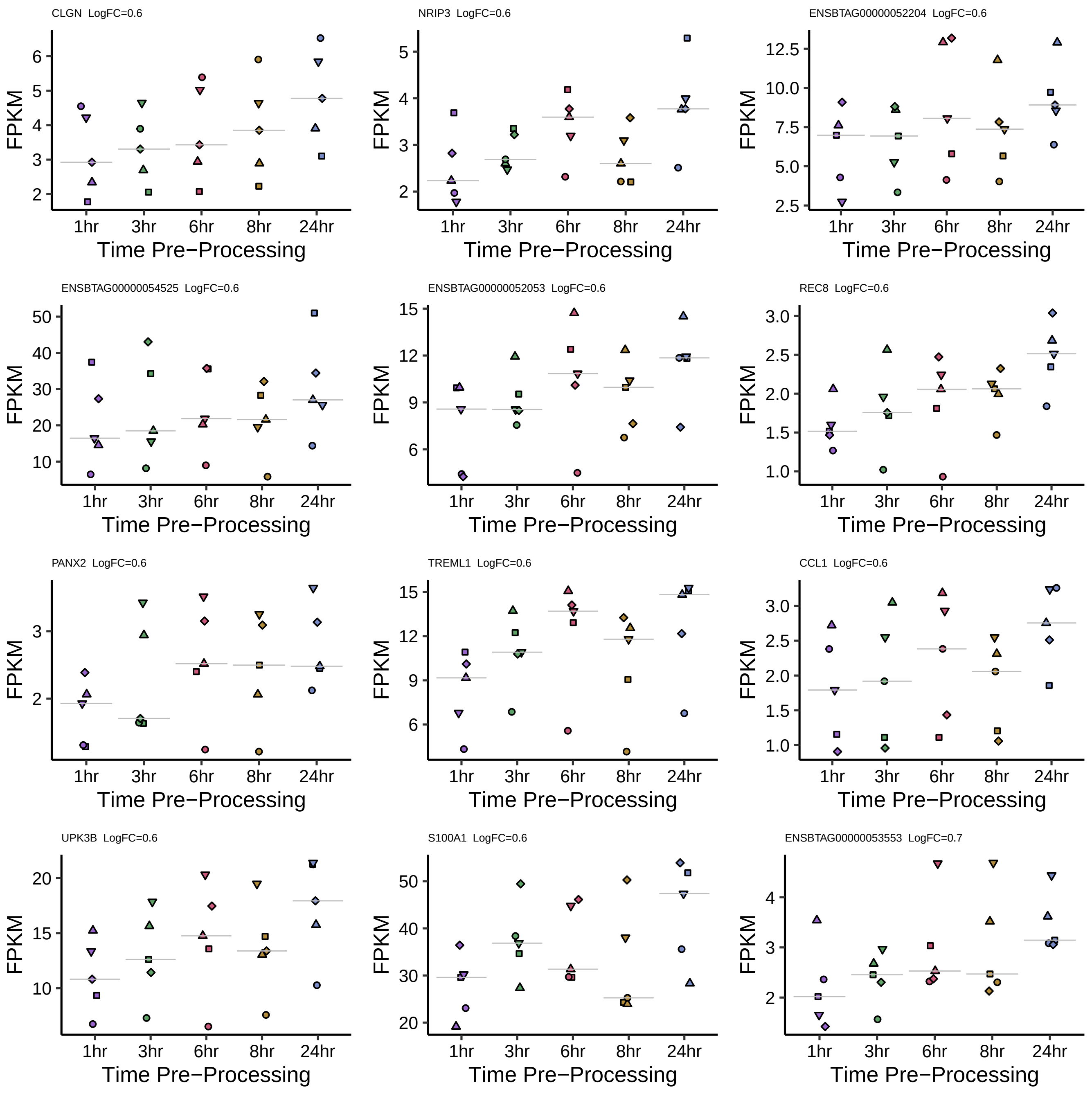

CLGN LogFC=0.6
NRIP3 LogFC=0.6
ENSBTAG00000052204 LogFC=0.6
ENSBTAG00000054525 LogFC=0.6
ENSBTAG00000052053 LogFC=0.6
REC8 LogFC=0.6
PANX2 LogFC=0.6
TREML1 LogFC=0.6
CCL1 LogFC=0.6
UPK3B LogFC=0.6
S100A1 LogFC=0.6
ENSBTAG00000053553 LogFC=0.7
FPKM
1hr
3hr
6hr
8hr
24hr
Time Pre−Processing

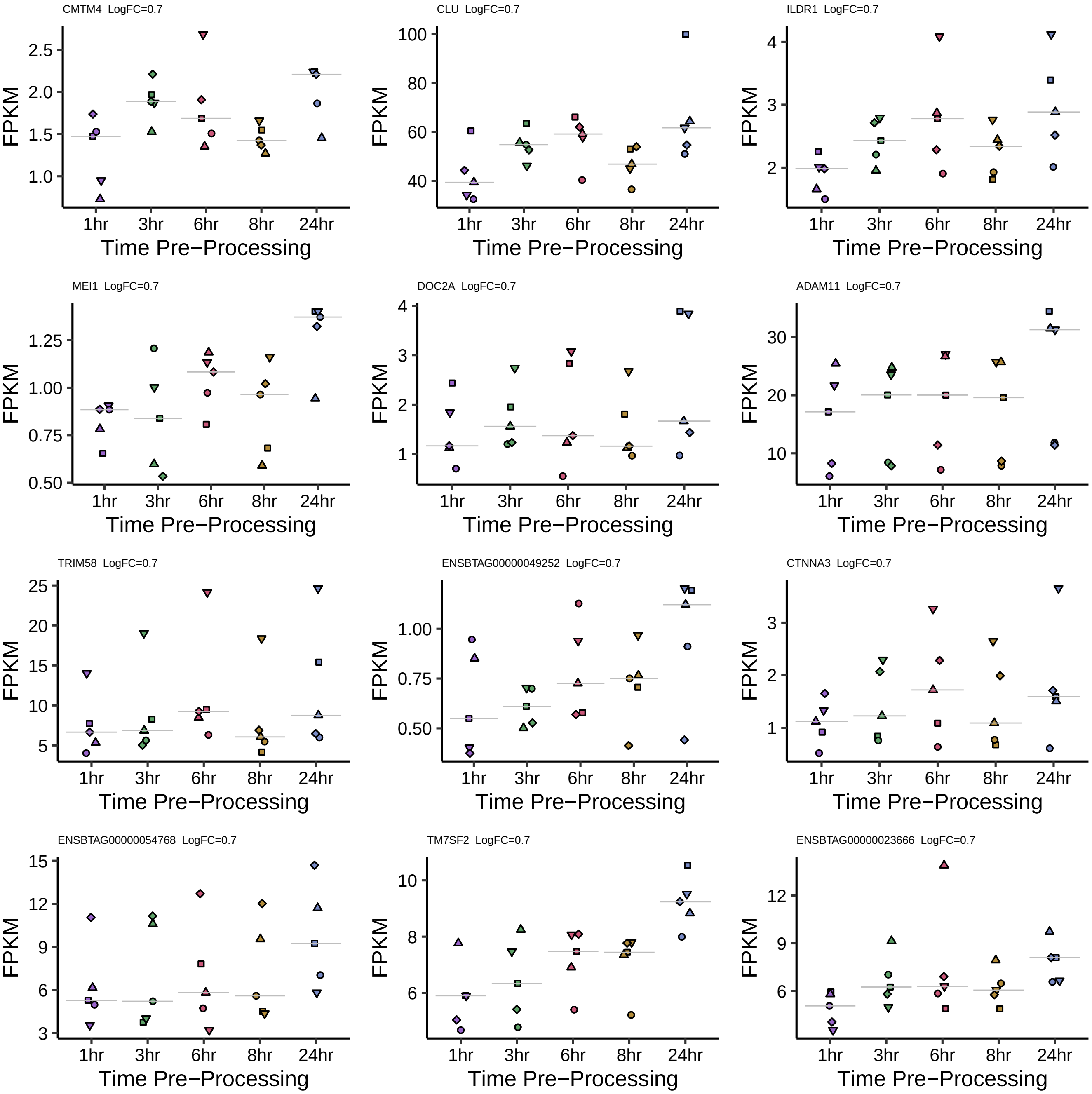

CMTM4 LogFC=0.7
CLU LogFC=0.7
ILDR1 LogFC=0.7
MEI1 LogFC=0.7
DOC2A LogFC=0.7
ADAM11 LogFC=0.7
TRIM58 LogFC=0.7
ENSBTAG00000049252 LogFC=0.7
CTNNA3 LogFC=0.7
ENSBTAG00000054768 LogFC=0.7
TM7SF2 LogFC=0.7
ENSBTAG00000023666 LogFC=0.7
FPKM
1hr
3hr
6hr
8hr
24hr
Time Pre−Processing

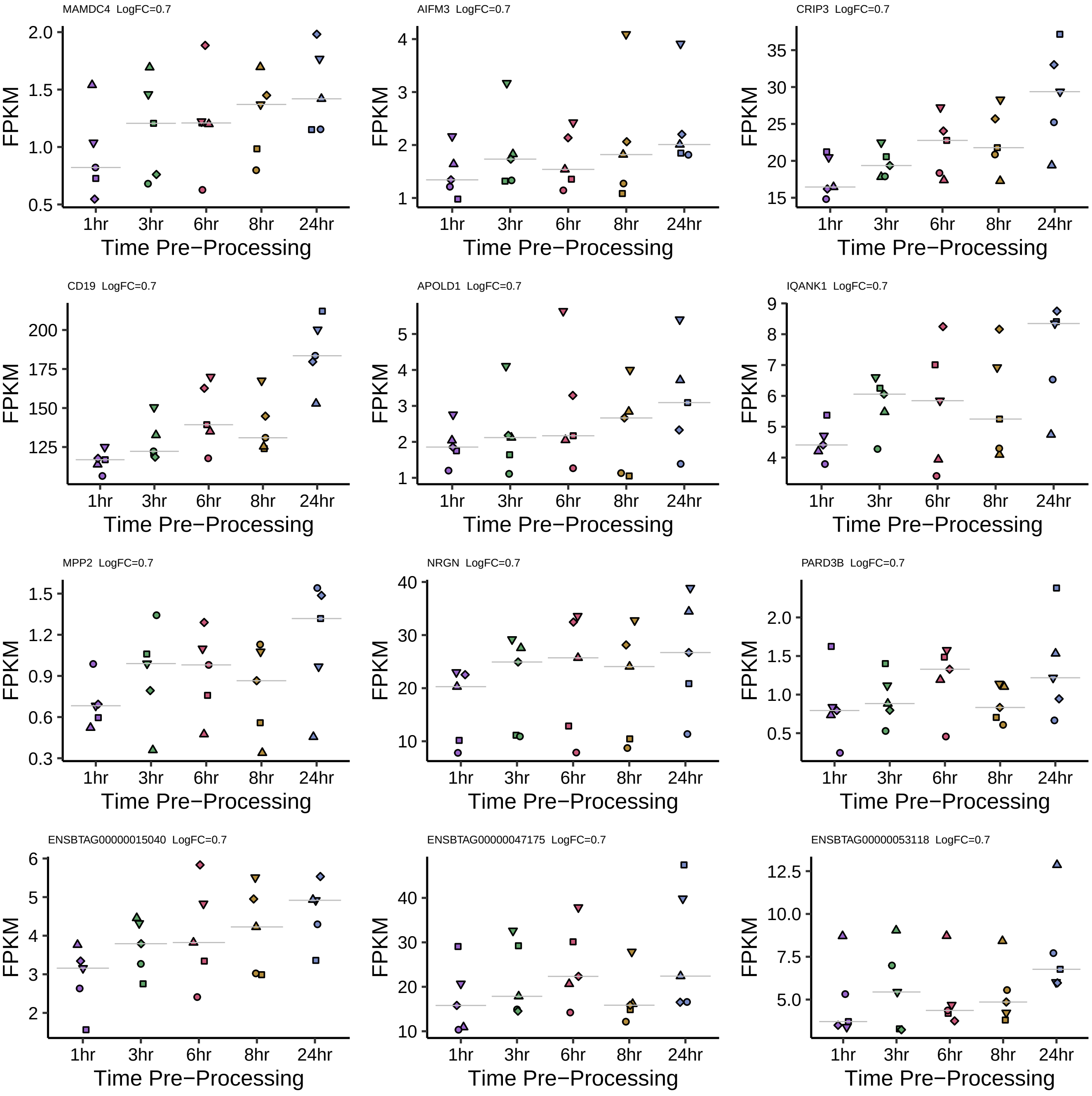

MAMDC4 LogFC=0.7
2.0
1.5
1.0
0.5
FPKM
1hr 3hr 6hr 8hr 24hr
Time Pre−Processing
AIFM3 LogFC=0.7
4
3
2
1
FPKM
1hr 3hr 6hr 8hr 24hr
Time Pre−Processing
CRIP3 LogFC=0.7
35
30
25
20
15
FPKM
1hr 3hr 6hr 8hr 24hr
Time Pre−Processing
CD19 LogFC=0.7
200
175
150
125
FPKM
1hr 3hr 6hr 8hr 24hr
Time Pre−Processing
APOLD1 LogFC=0.7
5
4
3
2
1
FPKM
1hr 3hr 6hr 8hr 24hr
Time Pre−Processing
IQANK1 LogFC=0.7
9
8
7
6
5
4
FPKM
1hr 3hr 6hr 8hr 24hr
Time Pre−Processing
MPP2 LogFC=0.7
1.5
1.2
0.9
0.6
0.3
FPKM
1hr 3hr 6hr 8hr 24hr
Time Pre−Processing
NRGN LogFC=0.7
40
30
20
10
FPKM
1hr 3hr 6hr 8hr 24hr
Time Pre−Processing
PARD3B LogFC=0.7
2.0
1.5
1.0
0.5
FPKM
1hr 3hr 6hr 8hr 24hr
Time Pre−Processing
ENSBTAG00000015040 LogFC=0.7
6
5
4
3
2
FPKM
1hr 3hr 6hr 8hr 24hr
Time Pre−Processing
ENSBTAG00000047175 LogFC=0.7
40
30
20
10
FPKM
1hr 3hr 6hr 8hr 24hr
Time Pre−Processing
ENSBTAG00000053118 LogFC=0.7
12.5
10.0
7.5
5.0
FPKM
1hr 3hr 6hr 8hr 24hr
Time Pre−Processing

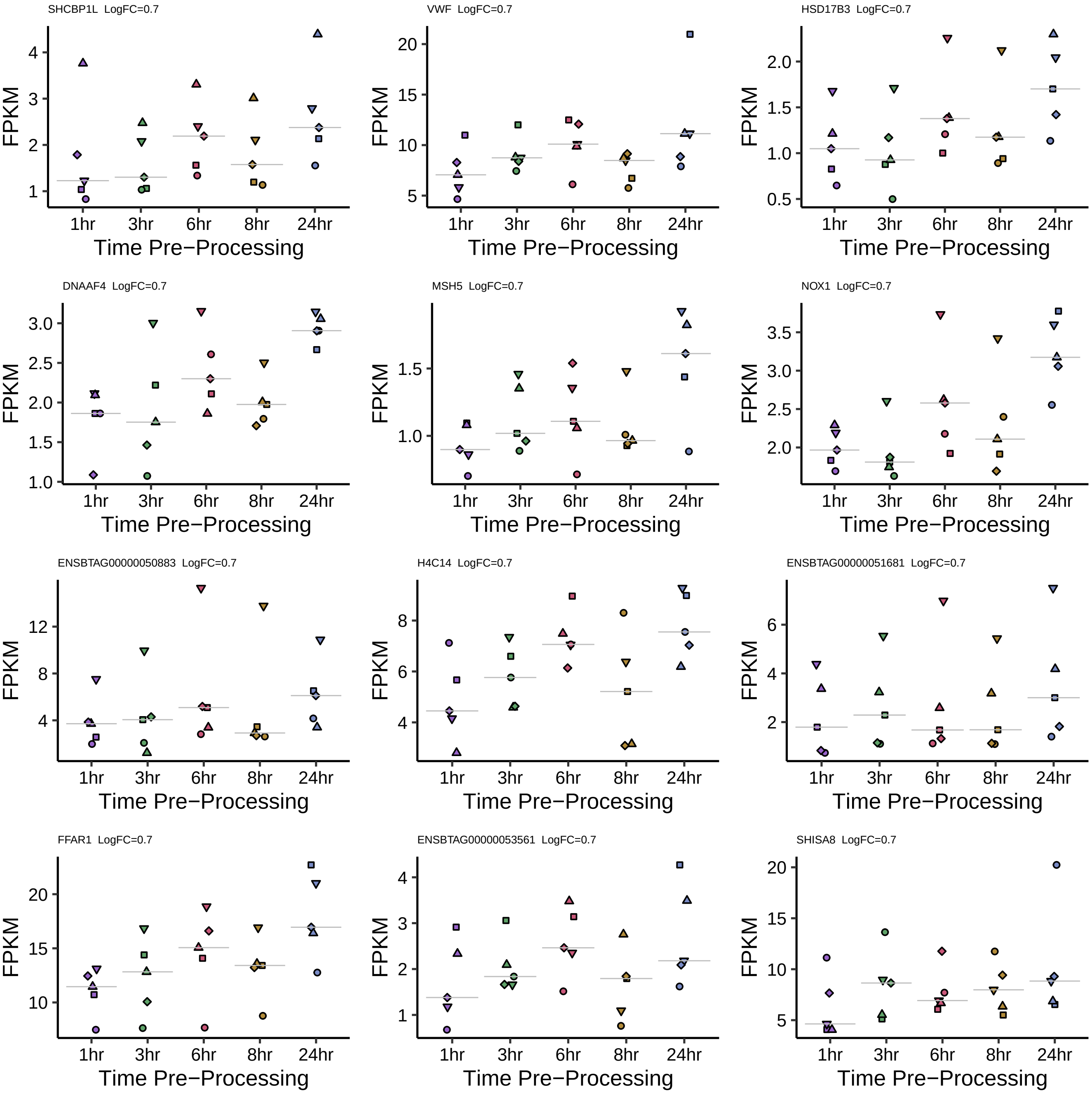
SHCBP1L LogFC=0.7
VWF LogFC=0.7
HSD17B3 LogFC=0.7
DNAAF4 LogFC=0.7
MSH5 LogFC=0.7
NOX1 LogFC=0.7
ENSBTAG00000050883 LogFC=0.7
H4C14 LogFC=0.7
ENSBTAG00000051681 LogFC=0.7
FFAR1 LogFC=0.7
ENSBTAG00000053561 LogFC=0.7
SHISA8 LogFC=0.7
FPKM
1hr
3hr
6hr
8hr
24hr
Time Pre−Processing

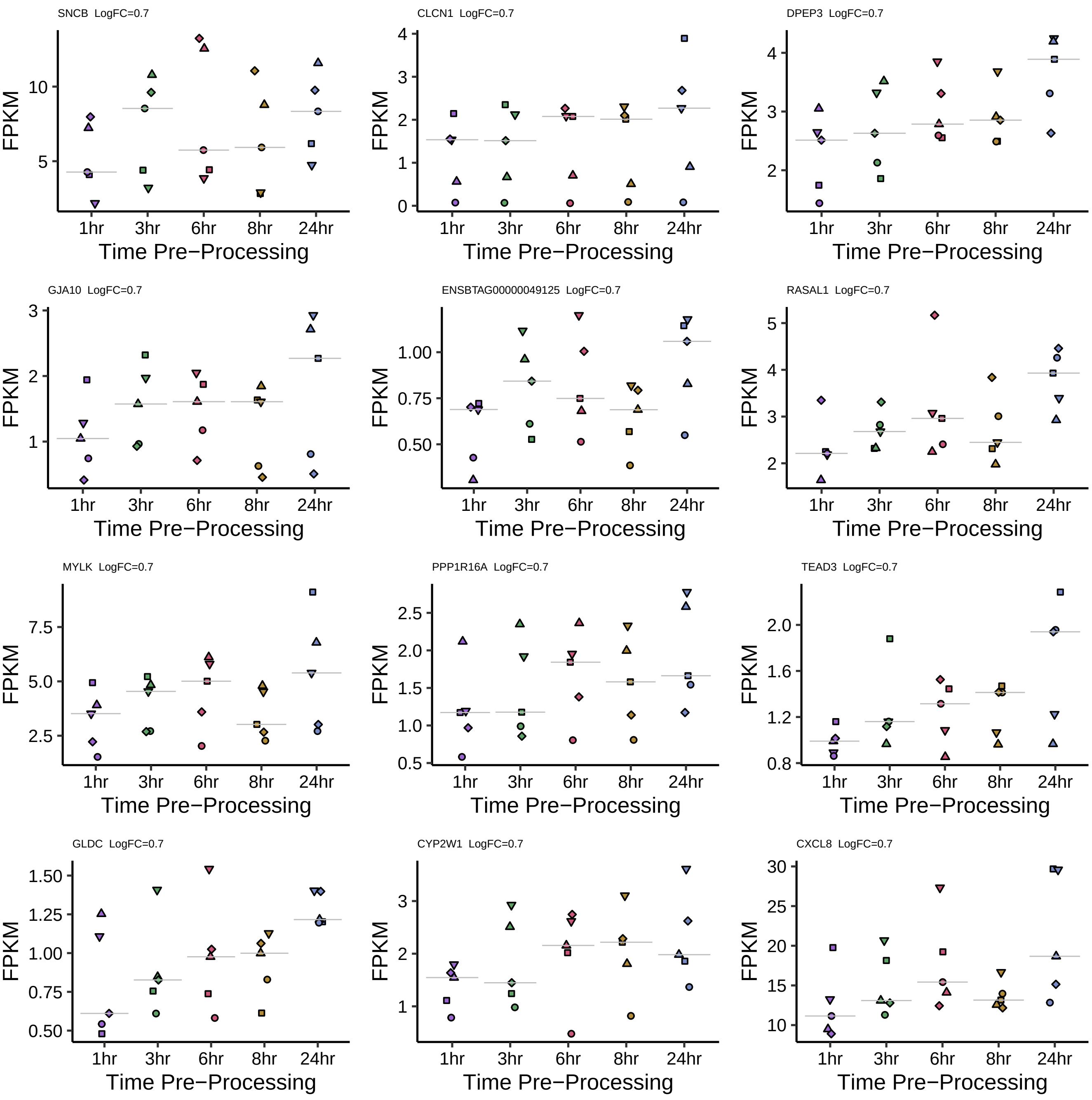
SNCB LogFC=0.7
CLCN1 LogFC=0.7
DPEP3 LogFC=0.7
GJA10 LogFC=0.7
ENSBTAG00000049125 LogFC=0.7
RASAL1 LogFC=0.7
MYLK LogFC=0.7
PPP1R16A LogFC=0.7
TEAD3 LogFC=0.7
GLDC LogFC=0.7
CYP2W1 LogFC=0.7
CXCL8 LogFC=0.7
FPKM
1hr
3hr
6hr
8hr
24hr
Time Pre−Processing

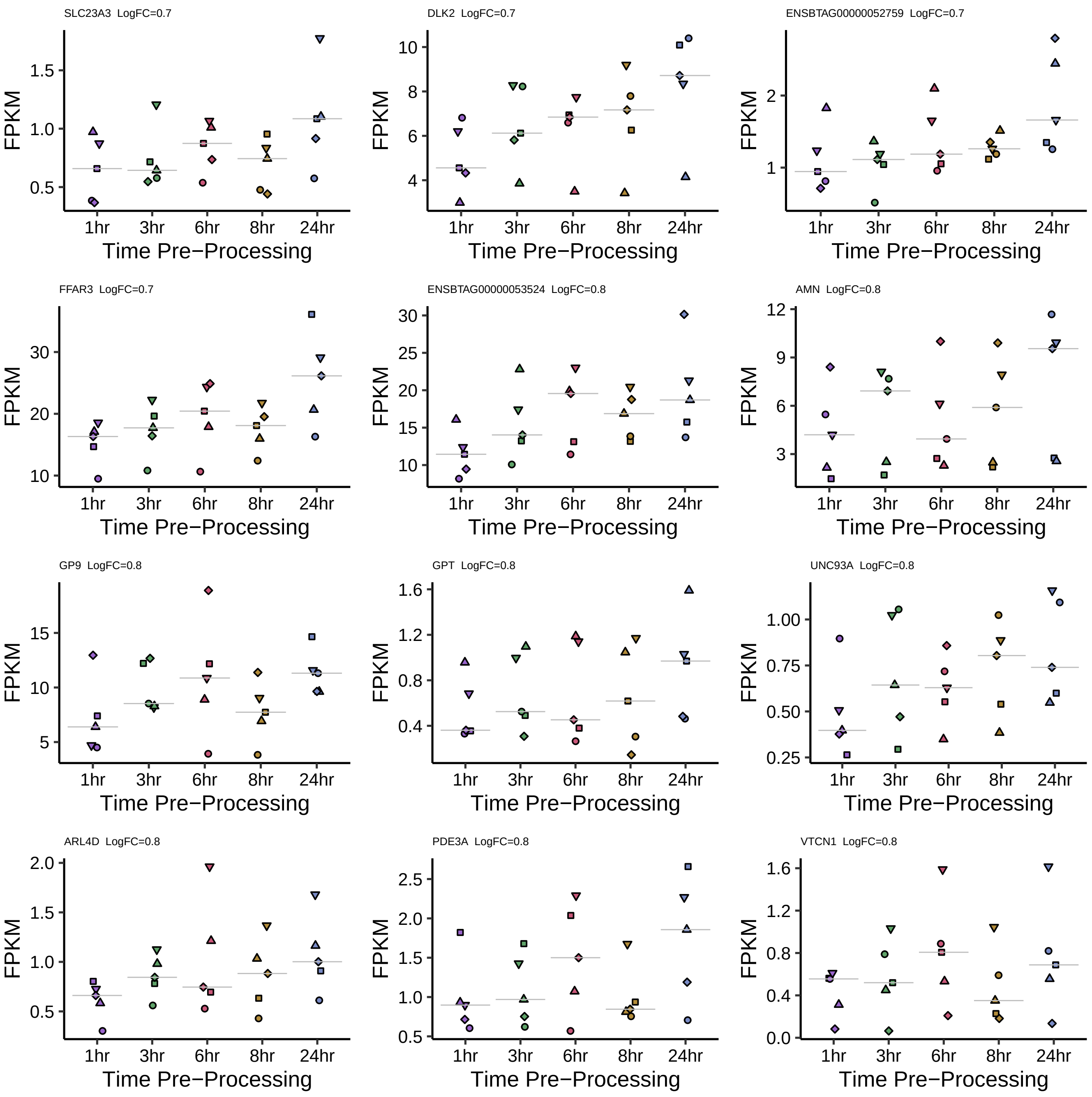
SLC23A3 LogFC=0.7
DLK2 LogFC=0.7
ENSBTAG00000052759 LogFC=0.7
FFAR3 LogFC=0.7
ENSBTAG00000053524 LogFC=0.8
AMN LogFC=0.8
GP9 LogFC=0.8
GPT LogFC=0.8
UNC93A LogFC=0.8
ARL4D LogFC=0.8
PDE3A LogFC=0.8
VTCN1 LogFC=0.8
FPKM
1hr
3hr
6hr
8hr
24hr
Time Pre−Processing

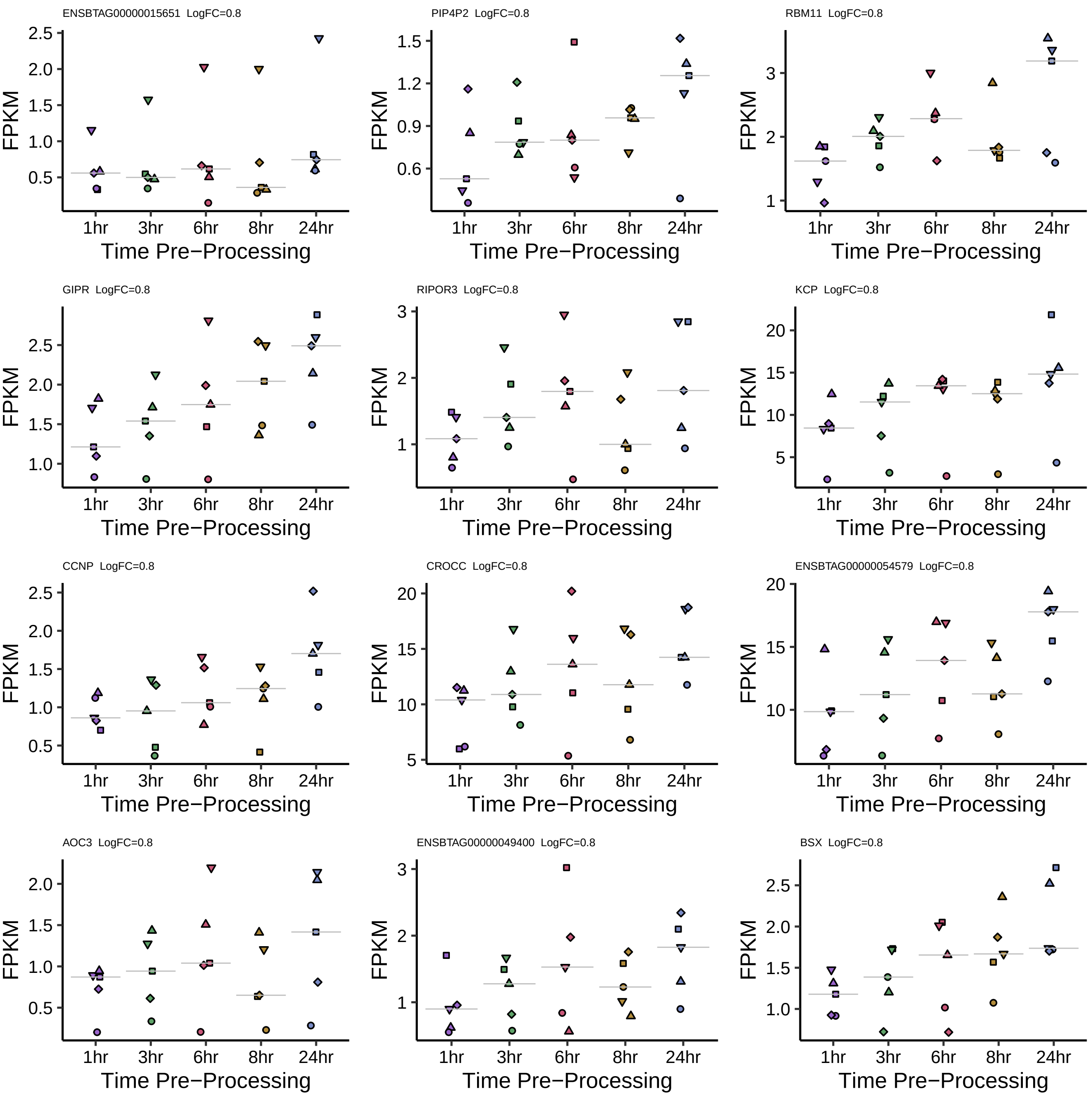
ENSBTAG00000015651 LogFC=0.8
PIP4P2 LogFC=0.8
RBM11 LogFC=0.8
GIPR LogFC=0.8
RIPOR3 LogFC=0.8
KCP LogFC=0.8
CCNP LogFC=0.8
CROCC LogFC=0.8
ENSBTAG00000054579 LogFC=0.8
AOC3 LogFC=0.8
ENSBTAG00000049400 LogFC=0.8
BSX LogFC=0.8
FPKM
1hr
3hr
6hr
8hr
24hr
Time Pre−Processing

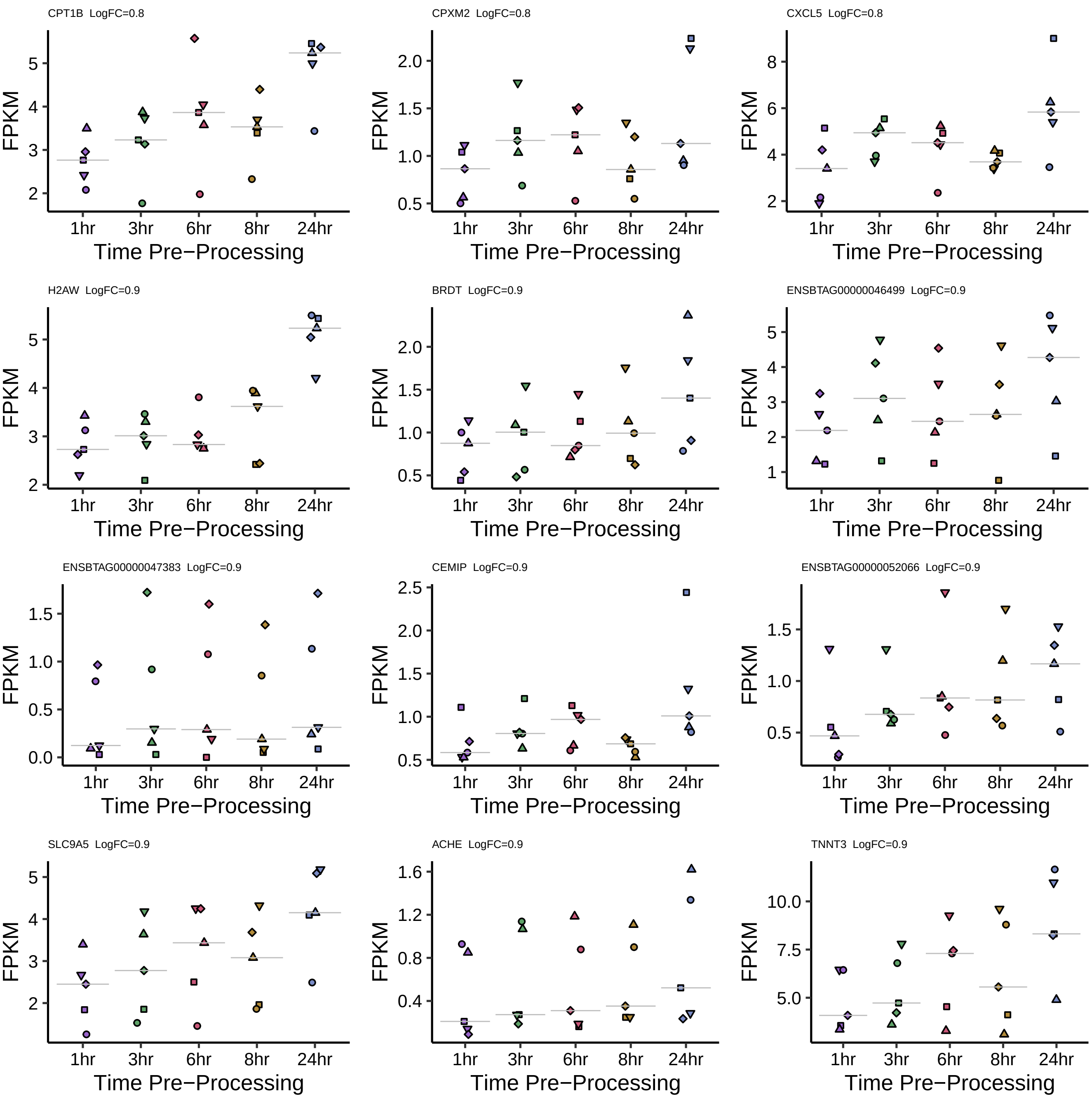
CPT1B LogFC=0.8
CPXM2 LogFC=0.8
CXCL5 LogFC=0.8
H2AW LogFC=0.9
BRDT LogFC=0.9
ENSBTAG00000046499 LogFC=0.9
ENSBTAG00000047383 LogFC=0.9
CEMIP LogFC=0.9
ENSBTAG00000052066 LogFC=0.9
SLC9A5 LogFC=0.9
ACHE LogFC=0.9
TNNT3 LogFC=0.9
FPKM
Time Pre−Processing
1hr
3hr
6hr
8hr
24hr

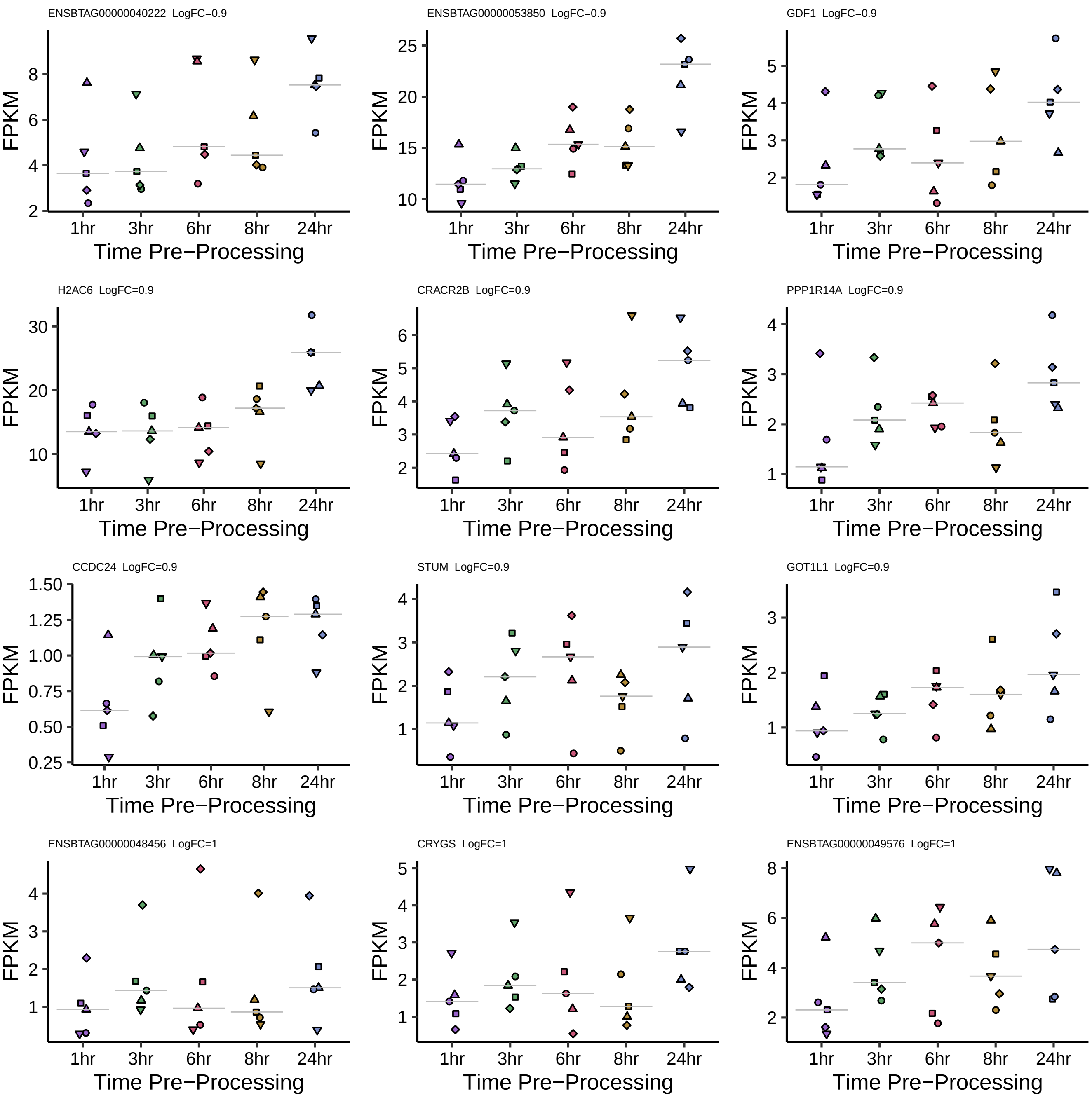
ENSBTAG00000040222 LogFC=0.9
ENSBTAG00000053850 LogFC=0.9
GDF1 LogFC=0.9
H2AC6 LogFC=0.9
CRACR2B LogFC=0.9
PPP1R14A LogFC=0.9
CCDC24 LogFC=0.9
STUM LogFC=0.9
GOT1L1 LogFC=0.9
ENSBTAG00000048456 LogFC=1
CRYGS LogFC=1
ENSBTAG00000049576 LogFC=1
FPKM
1hr
3hr
6hr
8hr
24hr
Time Pre−Processing

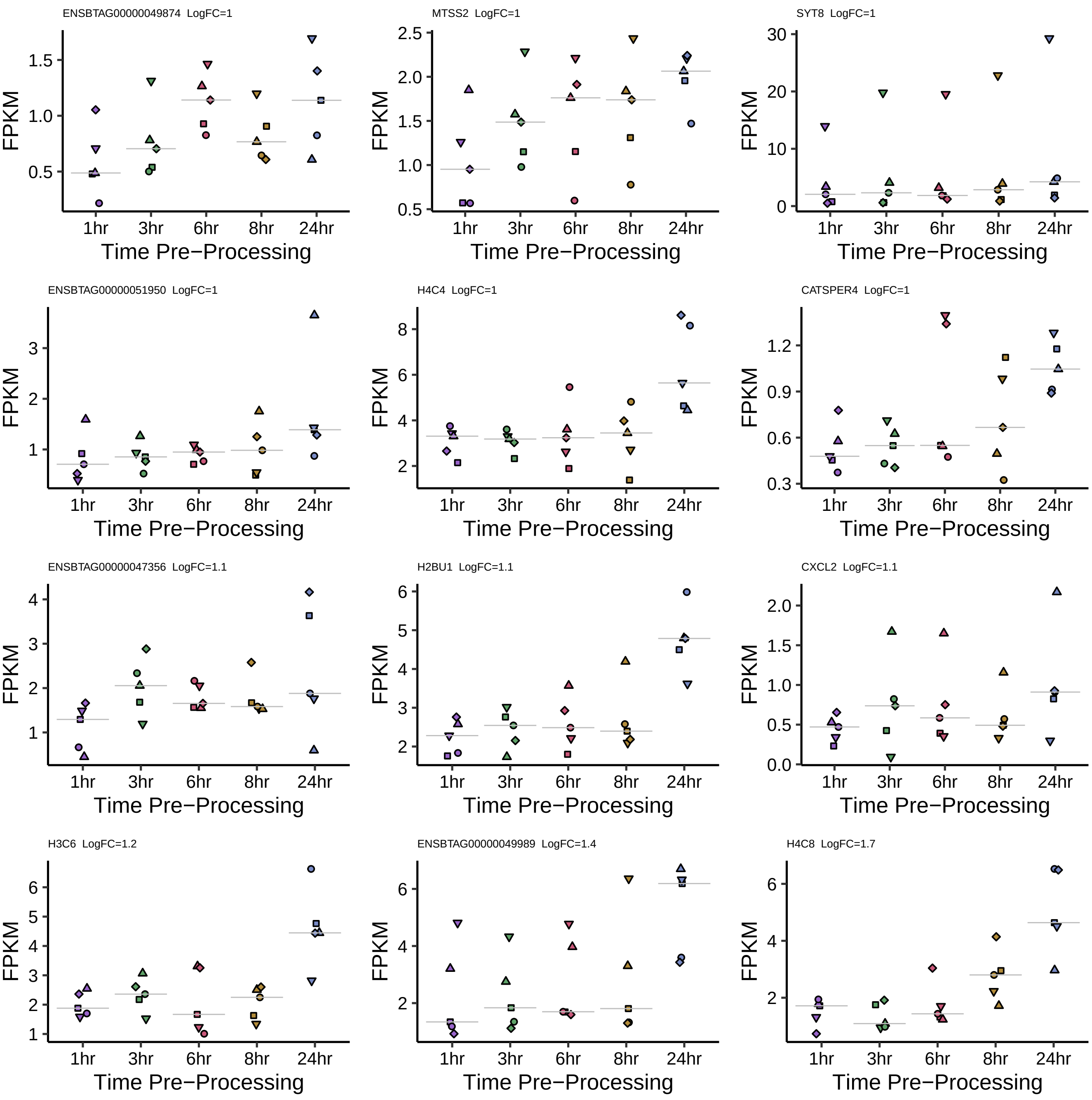
ENSBTAG00000049874 LogFC=1
MTSS2 LogFC=1
SYT8 LogFC=1
ENSBTAG00000051950 LogFC=1
H4C4 LogFC=1
CATSPER4 LogFC=1
ENSBTAG00000047356 LogFC=1.1
H2BU1 LogFC=1.1
CXCL2 LogFC=1.1
H3C6 LogFC=1.2
ENSBTAG00000049989 LogFC=1.4
H4C8 LogFC=1.7
FPKM
1hr
3hr
6hr
8hr
24hr
Time Pre−Processing

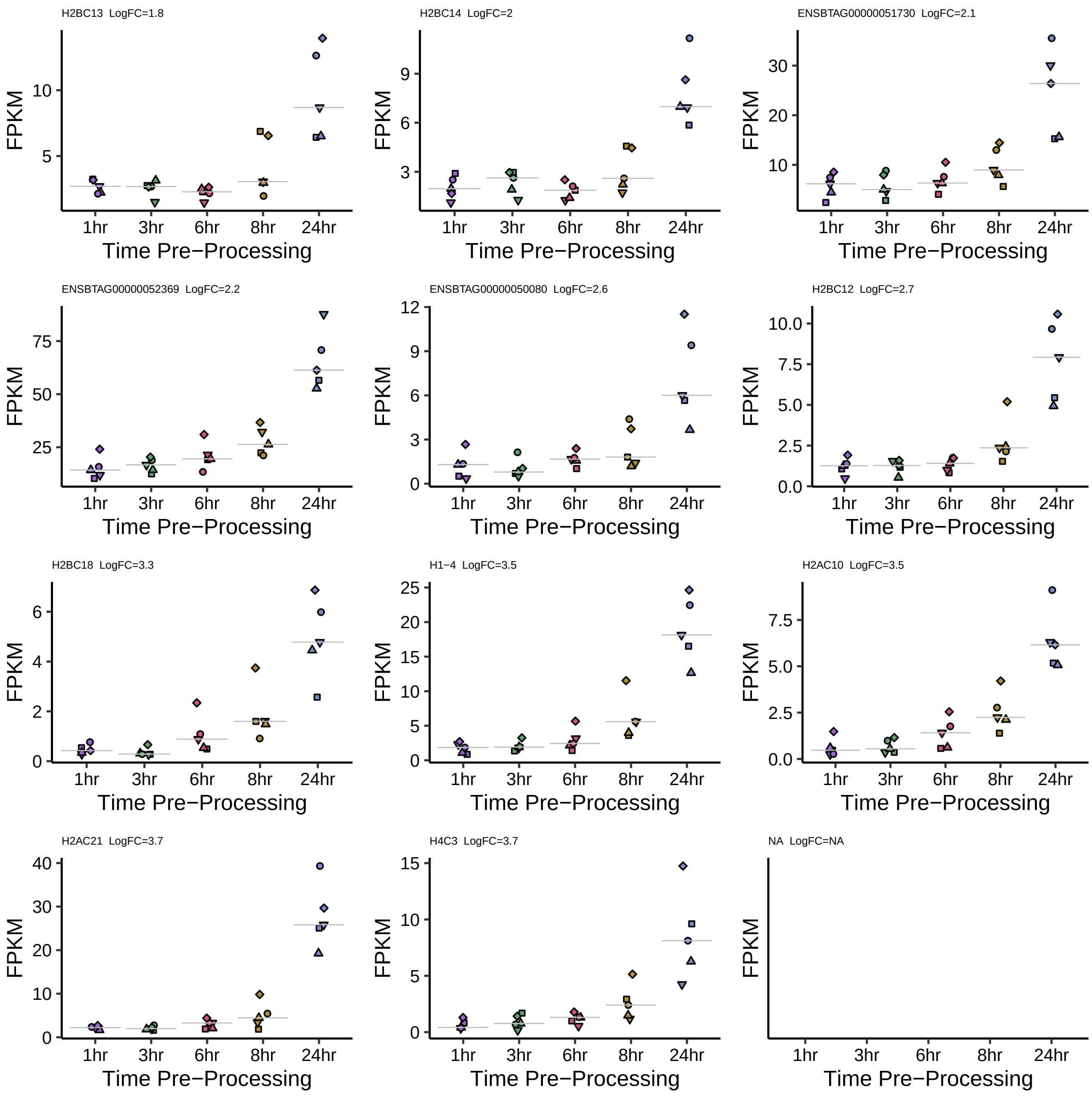
H2BC13 LogFC=1.8
H2BC14 LogFC=2
ENSBTAG00000051730 LogFC=2.1
ENSBTAG00000052369 LogFC=2.2
ENSBTAG00000050080 LogFC=2.6
H2BC12 LogFC=2.7
H2BC18 LogFC=3.3
H1−4 LogFC=3.5
H2AC10 LogFC=3.5
H2AC21 LogFC=3.7
H4C3 LogFC=3.7
NA LogFC=NA
FPKM
1hr
3hr
6hr
8hr
24hr
Time Pre−Processing
