## Additional file 12 for "Delayed processing of blood samples impairs the accuracy of mRNA-based biomarkers": Additional_file_12.html

#### 2022-01-06

*Background*: The transcriptome of peripheral white blood cells (PWBCs) contains valuable physiological information, thus making them a prime biological sample for investigating mRNA-based biomarkers. However, prolonged storage of whole blood samples can alter gene transcript abundance in PWBCs, compromising the results of biomarker discovery. Here, we designed an experiment to interrogate the impacts of delayed processing of whole blood samples on gene transcript abundance in PWBCs. We hypothesized that storing blood samples for 24 hours at 4°C would cause RNA degradation resulting in altered transcriptome profiles.   
 *Results*: We produced RNA-sequencing data for 30 samples collected from five estrus synchronized heifers (Bos taurus). We quantified transcript abundance for 12,414 protein-coding genes in PWBCs. Analysis of parameters of RNA quality revealed no statistically significant differences (P>0.05) between samples collected from the jugular vein and coccygeal vein, as well as among samples processed after one, three, six, or eight hours. However, samples processed after 24 hours of storage had a lower RNA integrity number value (P=0.03) in comparison to those processed after one hour of storage. Next, we analyzed RNA-sequencing data between samples using those processed after one hour of storage as the baseline for comparison. Interestingly, evaluation of 3’/5’ bias revealed no differences between genes with lower transcript abundance in samples stored for 24 hours relative to one hour. In addition, sequencing coverage of transcripts was similar between samples from the 24-hour and one-hour groups. We identified four and 515 genes with differential transcript abundance in samples processed after storage for eight and 24 hours, respectively, relative to samples processed after one hour.   
 *Conclusions*: The PWBCs respond to prolonged cold storage by increasing genes related to active chromatin compaction which in turn reduces gene transcription. This alteration in transcriptome profiles can impair the accuracy of mRNA-based biomarkers. Therefore, blood samples collected for mRNA-based biomarker discovery should be refrigerated immediately and processed within six hours post sampling.

### Overview

Code produced by Fernando Biase and Chace Wilson. We created this file to permit reproducibility of the findings described in the paper. Please direct questions to Fernando Biase: ***fbiase*** at ***vt.edu***   
 The raw data is deposited on GEO repository under the following access GSE192530. For the reproduction of this code please download the .RData file containing all the objects corresponding to the data used in this work: resource\_data\_2022\_01\_05.RData

### Code

Use the tab below to navigate the different segments of our code.

#### Scripts to process raw files

##### Align raw reads to the Cattle genome

```
#!/bin/bash
##Paths: 
path_to_index=/Bos_taurus.ARS-UCD1.2.99
splice_sites=/hisat_splice.txt
path_to_picard=/
fastq=/fastq

for sample in $(ls $fastq/*.gz | cut -c 64-71 | sort | uniq); do
fastq1=$(ls $fastq/* | grep $sample | grep "trimmed" |  grep "R1")
fastq2=$(ls $fastq/* | grep $sample | grep "trimmed" |  grep "R2")
echo "aligning"
echo $fastq1
echo $fastq2
mkdir -p /alignment_a/$sample
cd /alignment_a/$sample
output_alignment=/alignment_a/$sample
hisat2  -p 30 -k 1 -x $path_to_index -1 $fastq1  -2 $fastq2  --bowtie2-dp 2 --score-min L,0,-1 --dta --known-splicesite-infile $splice_sites  -S $output_alignment/alignment.sam  --summary-file $output_alignment/alignment_summary.txt
done;
```

##### Convert sam files into bam files

```
#!/bin/bash
for folder in /alignment_a/GS*; do
samtools sort -m 4G -@ 20 -O BAM -l 9 -o $folder/alignment.bam  $folder/alignment.sam
rm $folder/alignment.sam
done;

for folder in /alignment_a/GS*; do
samtools index -@ 20  $folder/alignment.bam 
done;
```

##### Filtering

```
#!/bin/bash
nproc=0     
maxProc=5 # Maximum number of processor to use

for folder in /alignment_a/GS*; do
samtools view -b -h -F 1796 -o $folder/alignment.filtered.bam $folder/alignment.bam  &

 (( nproc++ ))
    if [ "$nproc" -ge "$maxProc" ]; then
        wait
        nproc=0
        echo RESET-------
    fi
    
done;

for folder in /alignment_a/GS*; do
samtools sort  -m 4G -@ 20 -O BAM -l 9 -o $folder/alignment.filtered.sorted.bam  $folder/alignment.filtered.bam
done;
```

##### Duplicate Removal

```
#!/bin/bash
for folder in /alignment_a/GS*; do
#input files

alignment=$folder/alignment.filtered.sorted.bam
echo 'working on'
echo $alignment

#output files
undup_output=$folder/alignment.filtered.sorted.undup.bam
bammarkduplicates I=$alignment O=$undup_output  level=9 markthreads=30 rmdup=1 index=1 dupindex=0  verbose=0 
done;
```

##### Counting

```
#!/bin/bash
for folder in /alignment_a/GS*; do
gtf_file=/Bos_taurus.ARS-UCD1.2.98.gtf
undup_output=$folder/alignment.filtered.sorted.undup.bam
#output files
sample=$(echo $folder | awk '{n=split($0,a,"/");print a[8]}')
output=/mRNA_count_a/$sample.count
featureCounts -s 0 -a $gtf_file -o $output -F 'GTF' -t 'exon' -g 'gene_id' --ignoreDup -T 30 -p  $undup_output
done;
```

##### Count absolute nucleotide coverage

```
bedtools coverage -d -s -b /alignment_a/GSC22336/alignment.filtered.sorted.undup.bam -a  /Bos_taurus.ARS-UCD1.2.98_exon.gtf  > /coverage_files/coverage_GSC22336.txt
bedtools coverage -d -s -b /alignment_a/GSC22342/alignment.filtered.sorted.undup.bam -a  /Bos_taurus.ARS-UCD1.2.98_exon.gtf  > /coverage_files/coverage_GSC22342.txt
bedtools coverage -d -s -b /alignment_a/GSC22348/alignment.filtered.sorted.undup.bam -a  /Bos_taurus.ARS-UCD1.2.98_exon.gtf  > /coverage_files/coverage_GSC22348.txt
bedtools coverage -d -s -b /alignment_a/GSC22354/alignment.filtered.sorted.undup.bam -a  /Bos_taurus.ARS-UCD1.2.98_exon.gtf  > /coverage_files/coverage_GSC22354.txt
bedtools coverage -d -s -b /alignment_a/GSC22360/alignment.filtered.sorted.undup.bam -a  /Bos_taurus.ARS-UCD1.2.98_exon.gtf  > /coverage_files/coverage_GSC22360.txt
bedtools coverage -d -s -b /alignment_a/GSC22340/alignment.filtered.sorted.undup.bam -a  /Bos_taurus.ARS-UCD1.2.98_exon.gtf  > /coverage_files/coverage_GSC22340.txt
bedtools coverage -d -s -b /alignment_a/GSC22346/alignment.filtered.sorted.undup.bam -a  /Bos_taurus.ARS-UCD1.2.98_exon.gtf  > /coverage_files/coverage_GSC22346.txt
bedtools coverage -d -s -b /alignment_a/GSC22352/alignment.filtered.sorted.undup.bam -a  /Bos_taurus.ARS-UCD1.2.98_exon.gtf  > /coverage_files/coverage_GSC22352.txt
bedtools coverage -d -s -b /alignment_a/GSC22358/alignment.filtered.sorted.undup.bam -a  /Bos_taurus.ARS-UCD1.2.98_exon.gtf  > /coverage_files/coverage_GSC22358.txt
bedtools coverage -d -s -b /alignment_a/GSC22364/alignment.filtered.sorted.undup.bam -a  /Bos_taurus.ARS-UCD1.2.98_exon.gtf  > /coverage_files/coverage_GSC22364.txt
bedtools coverage -d -s -b /alignment_a/GSC22335/alignment.filtered.sorted.undup.bam -a  /Bos_taurus.ARS-UCD1.2.98_exon.gtf  > /coverage_files/coverage_GSC22335.txt
bedtools coverage -d -s -b /alignment_a/GSC22341/alignment.filtered.sorted.undup.bam -a  /Bos_taurus.ARS-UCD1.2.98_exon.gtf  > /coverage_files/coverage_GSC22341.txt
bedtools coverage -d -s -b /alignment_a/GSC22347/alignment.filtered.sorted.undup.bam -a  /Bos_taurus.ARS-UCD1.2.98_exon.gtf  > /coverage_files/coverage_GSC22347.txt
bedtools coverage -d -s -b /alignment_a/GSC22353/alignment.filtered.sorted.undup.bam -a  /Bos_taurus.ARS-UCD1.2.98_exon.gtf  > /coverage_files/coverage_GSC22353.txt
bedtools coverage -d -s -b /alignment_a/GSC22359/alignment.filtered.sorted.undup.bam -a  /Bos_taurus.ARS-UCD1.2.98_exon.gtf  > /coverage_files/coverage_GSC22359.txt
```

##### RNASeQC

```
gtf=/Bos_taurus.ARS-UCD1.2.103_collapsed.gtf
fasta=/Bos_taurus.ARS-UCD1.2.dna.toplevel.fa
output=/RNA_SeQC

bam=/alignment_a/GSC22335/alignment.bam
/rnaseqc.v2.4.2.linux  $gtf $bam  $output --unpaired --fasta=$fasta  --gene-length=500 --verbose --legacy --sample=GSC22335
bam=/alignment_a/GSC22336/alignment.bam
/rnaseqc.v2.4.2.linux  $gtf $bam  $output --unpaired --fasta=$fasta  --gene-length=500 --verbose --legacy --sample=GSC22336
bam=/alignment_a/GSC22337/alignment.bam
/rnaseqc.v2.4.2.linux  $gtf $bam  $output --unpaired --fasta=$fasta  --gene-length=500 --verbose --legacy --sample=GSC22337
bam=/alignment_a/GSC22338/alignment.bam
/rnaseqc.v2.4.2.linux  $gtf $bam  $output --unpaired --fasta=$fasta  --gene-length=500 --verbose --legacy --sample=GSC22338
bam=/alignment_a/GSC22339/alignment.bam
/rnaseqc.v2.4.2.linux  $gtf $bam  $output --unpaired --fasta=$fasta  --gene-length=500 --verbose --legacy --sample=GSC22339
bam=/alignment_a/GSC22340/alignment.bam
/rnaseqc.v2.4.2.linux  $gtf $bam  $output --unpaired --fasta=$fasta  --gene-length=500 --verbose --legacy --sample=GSC22340
bam=/alignment_a/GSC22341/alignment.bam
/rnaseqc.v2.4.2.linux  $gtf $bam  $output --unpaired --fasta=$fasta  --gene-length=500 --verbose --legacy --sample=GSC22341
bam=/alignment_a/GSC22342/alignment.bam
/rnaseqc.v2.4.2.linux  $gtf $bam  $output --unpaired --fasta=$fasta  --gene-length=500 --verbose --legacy --sample=GSC22342
bam=/alignment_a/GSC22343/alignment.bam
/rnaseqc.v2.4.2.linux  $gtf $bam  $output --unpaired --fasta=$fasta  --gene-length=500 --verbose --legacy --sample=GSC22343
bam=/alignment_a/GSC22344/alignment.bam
/rnaseqc.v2.4.2.linux  $gtf $bam  $output --unpaired --fasta=$fasta  --gene-length=500 --verbose --legacy --sample=GSC22344
bam=/alignment_a/GSC22344/alignment.bam
/rnaseqc.v2.4.2.linux  $gtf $bam  $output --unpaired --fasta=$fasta  --gene-length=500 --verbose --legacy --sample=GSC22344
bam=/alignment_a/GSC22345/alignment.bam
/rnaseqc.v2.4.2.linux  $gtf $bam  $output --unpaired --fasta=$fasta  --gene-length=500 --verbose --legacy --sample=GSC22345
bam=/alignment_a/GSC22346/alignment.bam
/rnaseqc.v2.4.2.linux  $gtf $bam  $output --unpaired --fasta=$fasta  --gene-length=500 --verbose --legacy --sample=GSC22346
bam=/alignment_a/GSC22347/alignment.bam
/rnaseqc.v2.4.2.linux  $gtf $bam  $output --unpaired --fasta=$fasta  --gene-length=500 --verbose --legacy --sample=GSC22347
bam=/alignment_a/GSC22348/alignment.bam
/rnaseqc.v2.4.2.linux  $gtf $bam  $output --unpaired --fasta=$fasta  --gene-length=500 --verbose --legacy --sample=GSC22348
bam=/alignment_a/GSC22349/alignment.bam
/rnaseqc.v2.4.2.linux  $gtf $bam  $output --unpaired --fasta=$fasta  --gene-length=500 --verbose --legacy --sample=GSC22349
bam=/alignment_a/GSC22350/alignment.bam
/rnaseqc.v2.4.2.linux  $gtf $bam  $output --unpaired --fasta=$fasta  --gene-length=500 --verbose --legacy --sample=GSC22350
bam=/alignment_a/GSC22351/alignment.bam
/rnaseqc.v2.4.2.linux  $gtf $bam  $output --unpaired --fasta=$fasta  --gene-length=500 --verbose --legacy --sample=GSC22351
bam=/alignment_a/GSC22352/alignment.bam
/rnaseqc.v2.4.2.linux  $gtf $bam  $output --unpaired --fasta=$fasta  --gene-length=500 --verbose --legacy --sample=GSC22352
bam=/alignment_a/GSC22353/alignment.bam
/rnaseqc.v2.4.2.linux  $gtf $bam  $output --unpaired --fasta=$fasta  --gene-length=500 --verbose --legacy --sample=GSC22353
bam=/alignment_a/GSC22354/alignment.bam
/rnaseqc.v2.4.2.linux  $gtf $bam  $output --unpaired --fasta=$fasta  --gene-length=500 --verbose --legacy --sample=GSC22354
bam=/alignment_a/GSC22355/alignment.bam
/rnaseqc.v2.4.2.linux  $gtf $bam  $output --unpaired --fasta=$fasta  --gene-length=500 --verbose --legacy --sample=GSC22355
bam=/alignment_a/GSC22356/alignment.bam
/rnaseqc.v2.4.2.linux  $gtf $bam  $output --unpaired --fasta=$fasta  --gene-length=500 --verbose --legacy --sample=GSC22356
bam=/alignment_a/GSC22357/alignment.bam
/rnaseqc.v2.4.2.linux  $gtf $bam  $output --unpaired --fasta=$fasta  --gene-length=500 --verbose --legacy --sample=GSC22357
bam=/alignment_a/GSC22358/alignment.bam
/rnaseqc.v2.4.2.linux  $gtf $bam  $output --unpaired --fasta=$fasta  --gene-length=500 --verbose --legacy --sample=GSC22358
bam=/alignment_a/GSC22359/alignment.bam
/rnaseqc.v2.4.2.linux  $gtf $bam  $output --unpaired --fasta=$fasta  --gene-length=500 --verbose --legacy --sample=GSC22359
bam=/alignment_a/GSC22360/alignment.bam
/rnaseqc.v2.4.2.linux  $gtf $bam  $output --unpaired --fasta=$fasta  --gene-length=500 --verbose --legacy --sample=GSC22360
bam=/alignment_a/GSC22361/alignment.bam
/rnaseqc.v2.4.2.linux  $gtf $bam  $output --unpaired --fasta=$fasta  --gene-length=500 --verbose --legacy --sample=GSC22361
bam=/alignment_a/GSC22362/alignment.bam
/rnaseqc.v2.4.2.linux  $gtf $bam  $output --unpaired --fasta=$fasta  --gene-length=500 --verbose --legacy --sample=GSC22362
bam=/alignment_a/GSC22363/alignment.bam
/rnaseqc.v2.4.2.linux  $gtf $bam  $output --unpaired --fasta=$fasta  --gene-length=500 --verbose --legacy --sample=GSC22363
bam=/alignment_a/GSC22364/alignment.bam
/rnaseqc.v2.4.2.linux  $gtf $bam  $output --unpaired --fasta=$fasta  --gene-length=500 --verbose --legacy --sample=GSC22364
```

#### Import files needed for reproducibility

This was run once to create the resource\_data\_2022\_01\_05.RData file which will be loaded next

##### RNA quantification and quality data

```
RNA_data_frame<- as.data.frame(read_excel("/mnt/storage/lab_folder/shared_R_codes/fernando/RNA_degradation/resources/RNA_degradation_study_sample_information.xlsx"))
```

##### Obtain ensembl annotation for reproducibility

```
library('biomaRt', lib.loc="/usr/lib/R/site-library")
cow<-useMart("ensembl", dataset = "btaurus_gene_ensembl", host="www.ensembl.org") 
annotation.ensembl.symbol<-getBM(attributes = c('ensembl_gene_id','external_gene_name','description','hgnc_symbol','gene_biotype','transcript_length','chromosome_name','start_position','end_position','strand'),  values = "*", mart = cow)

cow<-useMart("ensembl", dataset = "btaurus_gene_ensembl", host="www.ensembl.org") 
annotation.ensembl.transcript<-getBM(attributes = c('ensembl_gene_id','gene_biotype','ensembl_transcript_id','transcript_length'),  values = "*", mart = cow)
annotation.ensembl.transcript<-annotation.ensembl.transcript[annotation.ensembl.transcript$gene_biotype =="protein_coding",]
annotation.ensembl.transcript<-annotation.ensembl.transcript[order(annotation.ensembl.transcript$ensembl_gene_id, -annotation.ensembl.transcript$transcript_length),]
annotation.ensembl.transcript<-annotation.ensembl.transcript[!duplicated(annotation.ensembl.transcript$ensembl_gene_id),]
annotation.ensembl.transcript<-annotation.ensembl.transcript[annotation.ensembl.transcript$transcript_length > 400,]

annotation.ensembl.symbol<-annotation.ensembl.symbol[order(annotation.ensembl.symbol$ensembl_gene_id, -annotation.ensembl.symbol$transcript_length),]
annotation.ensembl.symbol<-annotation.ensembl.symbol[!duplicated(annotation.ensembl.symbol$ensembl_gene_id),]
gene.length<-annotation.ensembl.symbol[,c( "ensembl_gene_id", "transcript_length" )]
annotation.GO.biomart<-getBM(attributes = c('ensembl_gene_id', 'external_gene_name','go_id','name_1006','namespace_1003'),  values = "*", mart = cow)

#write.table(annotation.ensembl.symbol,file="/mnt/storage/lab_folder/shared_R_codes/fernando/RNA_degradation/resources/2021_02_19_annotation.ensembl.symbol.txt", sep = "\t",append = FALSE, quote = FALSE)
#system('bzip2 --best /mnt/storage/lab_folder/shared_R_codes/fernando/RNA_degradation/resources/2021_02_19_annotation.ensembl.symbol.txt')

#write.table(gene.length,file="/mnt/storage/lab_folder/shared_R_codes/fernando/RNA_degradation/resources/2021_02_19_gene.length.txt", sep = "\t",append = FALSE, quote = FALSE)
#system('bzip2 --best /mnt/storage/lab_folder/shared_R_codes/fernando/RNA_degradation/resources/2021_02_19_gene.length.txt')

#write.table(annotation.GO.biomart,file="/mnt/storage/lab_folder/shared_R_codes/fernando/RNA_degradation/resources/2021_02_19_annotation.GO.biomart.txt", sep = "\t",append = FALSE, quote = FALSE)
#system('bzip2 --best /mnt/storage/lab_folder/shared_R_codes/fernando/RNA_degradation/resources/2021_02_19_annotation.GO.biomart.txt')

#write.table(annotation.ensembl.transcript,file="/mnt/storage/lab_folder/shared_R_codes/fernando/RNA_degradation/resources/2021_02_19_annotation.ensembl.transcript.txt", sep = "\t",append = FALSE, quote = FALSE)
#system('bzip2 --best /mnt/storage/lab_folder/shared_R_codes/fernando/RNA_degradation/resources/2021_02_19_annotation.ensembl.transcript.txt')
```

##### Import annotation

```
annotation.ensembl.symbol<-read.delim("/mnt/storage/lab_folder/shared_R_codes/fernando/RNA_degradation/resources/2021_02_19_annotation.ensembl.symbol.txt.bz2", header=TRUE, sep= "\t",row.names=1, stringsAsFactors = FALSE)
gene.length<-read.delim("/mnt/storage/lab_folder/shared_R_codes/fernando/RNA_degradation/resources/2021_02_19_gene.length.txt.bz2", header=TRUE, sep= "\t",row.names=1, stringsAsFactors = FALSE)
annotation.GO.biomart<-read.delim("/mnt/storage/lab_folder/shared_R_codes/fernando/RNA_degradation/resources/2021_02_19_annotation.GO.biomart.txt.bz2", header=TRUE, sep= "\t",row.names=1, stringsAsFactors = FALSE)
annotation.ensembl.transcript<-read.delim("/mnt/storage/lab_folder/shared_R_codes/fernando/RNA_degradation/resources/2021_02_19_annotation.ensembl.transcript.txt.bz2", header=TRUE, sep= "\t",row.names=1, stringsAsFactors = FALSE)
```

##### Import read counts

```
files<-list.files("/mnt/storage/lab_folder/heifer_infertility/RNA_integrity/alignment_a", recursive=T, pattern="summary.txt", full.names = TRUE)
#length(files)

reads_produced_a<-data.frame()
reads_produced_b<-data.frame()
#length(files)

for (n in 1:30) {
reads_produced<-read.delim(files[n], sep= " ",header=FALSE, stringsAsFactors = FALSE, comment.char= "#")
sample<-substring(files[n], 70,77)
number_of_reads_produced<-as.integer(reads_produced[1,1])

reads_produced_a<-data.frame(sample,number_of_reads_produced)
reads_produced_b<-rbind(reads_produced_b,reads_produced_a)
}

files<-list.files("/mnt/storage/lab_folder/heifer_infertility/RNA_integrity/mRNA_count_a", recursive=T, pattern="count.summary", full.names = TRUE)
#length(files)
counting_summary_a<-data.frame()
counting_summary_b<-data.frame()
#length(files)

for (n in 1:30) {
counting_summary<-read.delim(files[n], sep= "\t",header=TRUE, stringsAsFactors = FALSE, comment.char= "#")
sample<-substring(files[n], 71,78)
number_of_reads_assigned<-counting_summary[1,2]
unassigned_NoFeatures<-counting_summary[12,2]
unassigned_Ambiguity<-counting_summary[14,2]
counting_summary_a<-data.frame(sample,number_of_reads_assigned, unassigned_NoFeatures, unassigned_Ambiguity)
counting_summary_b<-rbind(counting_summary_b,counting_summary_a)
}

counting_summary_b$reads_sequenced<-reads_produced_b$number_of_reads_produced
counting_summary_b$reads_retained<-counting_summary_b$number_of_reads_assigned + counting_summary_b$unassigned_NoFeatures + counting_summary_b$unassigned_Ambiguity
counting_summary_b$reads_discarted<-counting_summary_b$reads_sequenced - counting_summary_b$reads_retained
counting_summary_b$perc_number_of_reads_discarted<-counting_summary_b$reads_discarted/counting_summary_b$reads_sequenced
counting_summary_b$perc_number_of_reads_retained<-counting_summary_b$reads_retained/counting_summary_b$reads_sequenced
counting_summary_b$perc_number_of_reads_assigned<-counting_summary_b$number_of_reads_assigned/counting_summary_b$reads_sequenced
counting_summary_b$perc_unassigned_NoFeatures<-counting_summary_b$unassigned_NoFeatures/counting_summary_b$reads_sequenced
counting_summary_b$perc_unassigned_Ambiguity<-counting_summary_b$unassigned_Ambiguity/counting_summary_b$reads_sequenced

#write.table(counting_summary_b, file="/counting_summary_b_second_alignment.txt",quote = FALSE, sep = "\t",row.names = FALSE)
```

##### Obtain counts per sample

```
files<-list.files("/mnt/storage/lab_folder/heifer_infertility/RNA_integrity/mRNA_count_a", recursive=T, pattern="count", full.names = TRUE)
files<-files[grep("summary", files, invert = TRUE)]
files<-files[grep(".count1", files, invert = TRUE)]
files<-files[grep(".count2", files, invert = TRUE)]
files<-files[grep(".sh", files, invert = TRUE)]
#length(files)
count_data<-data.frame(matrix(nrow=27607))
for (n in 1:length(files)) {
  count<-read.delim(files[n], header=TRUE, sep= "\t", stringsAsFactors = FALSE, comment.char= "#")
  count<-count[,c(1,7)]
  count_data<-cbind(count_data,count)
}
rownames(count_data)<-count_data[,2]
count_data<-count_data[,seq(from = 3, to = 61, by = 2)]
colnames(count_data)<- substr(colnames(count_data), 71, 78)

#write_delim(count_data, "/2021_12_21_unfiltered_count_data.txt", delim = "\t", quote =  "none")
#system("bzip2 /2021_12_21_unfiltered_count_data.txt")
```

##### Import parameters regarding sequencing efficiency

```
files<-list.files("/mnt/storage/lab_folder/heifer_infertility/RNA_integrity/RNA_SeQC", recursive=T, pattern="metrics.tsv", full.names = TRUE)
#length(files)

bias_metrics_a<-data.frame()
bias_metrics_b<-data.frame()

for (n in 1:30) {
bias_metrics<-read.delim(files[n], sep= "\t",header=FALSE, stringsAsFactors = FALSE, comment.char= "#")
sample<-substring(files[n], 67,74)
three_prime_bias_mean<-as.numeric(bias_metrics[69,2])
three_prime_bias_median<-as.numeric(bias_metrics[70,2])

bias_metrics_a<-data.frame(sample,three_prime_bias_mean,three_prime_bias_median)
bias_metrics_b<-rbind(bias_metrics_a,bias_metrics_b)
}
```

##### Calculate transcript coverage

This chunk was used to produce the objects with the relative coverage of transcripts. The results were saved separately and loaded together with the .RData file.

```
library("foreach", lib.loc="/usr/lib/R/site-library")
library("parallel", lib.loc="/usr/lib/R/site-library")
library("doParallel", lib.loc="/usr/lib/R/site-library")
library("data.table", lib.loc="/usr/lib/R/site-library")
library("spatstat", lib.loc="/usr/lib/R/site-library")

gc()
GSC22335<-data.table::fread(file="/coverage_files/coverage_GSC22335.txt.gz", sep = "\t", header = FALSE ,colClasses= c( rep("NULL", 8), "character", "NULL", "integer"), quote=FALSE, stringsAsFactors=FALSE, nThread=2, showProgress=FALSE)
GSC22335<-GSC22335[,"gene_id" := tstrsplit(V9, " ", fixed=TRUE)[2]]
GSC22335<-GSC22335[,c("V11","gene_id")]
GSC22335<-GSC22335[,gene_id := substr(gene_id, 2,19)]
GSC22335<-GSC22335[gene_id %in% genes_expressed,]
GSC22335<-GSC22335[ , ':=' (Counter= 1:.N, Length = .N, Sum = sum(V11)), by = gene_id ]
GSC22335<-GSC22335[V11 > 0,]
GSC22335<-GSC22335[Sum > 100,]
GSC22335<-GSC22335[Length > 500,]
GSC22335<-GSC22335[ , prop := round(V11 / Sum,5)]
GSC22335<-GSC22335[ , rel_position := round(Counter / Length,5)]
GSC22335<-GSC22335[ , c("gene_id", "prop" ,"rel_position")]

GSC22336<-data.table::fread(file="/coverage_files/coverage_GSC22336.txt.gz", sep = "\t", header = FALSE ,colClasses= c( rep("NULL", 8), "character", "NULL", "integer"), quote=FALSE, stringsAsFactors=FALSE, nThread=2, showProgress=FALSE)
GSC22336<-GSC22336[,"gene_id" := tstrsplit(V9, " ", fixed=TRUE)[2]]
GSC22336<-GSC22336[,c("V11","gene_id")]
GSC22336<-GSC22336[,gene_id := substr(gene_id, 2,19)]
GSC22336<-GSC22336[gene_id %in% genes_expressed,]
GSC22336<-GSC22336[ , ':=' (Counter= 1:.N, Length = .N, Sum = sum(V11)), by = gene_id ]
GSC22336<-GSC22336[V11 > 0,]
GSC22336<-GSC22336[Sum > 100,]
GSC22336<-GSC22336[Length > 500,]
GSC22336<-GSC22336[ , prop := round(V11 / Sum,5)]
GSC22336<-GSC22336[ , rel_position := round(Counter / Length,5)]
GSC22336<-GSC22336[ , c("gene_id", "prop" ,"rel_position")]

GSC22340<-data.table::fread(file="/coverage_files/coverage_GSC22340.txt.gz", sep = "\t", header = FALSE ,colClasses= c( rep("NULL", 8), "character", "NULL", "integer"), quote=FALSE, stringsAsFactors=FALSE, nThread=2, showProgress=FALSE)
GSC22340<-GSC22340[,"gene_id" := tstrsplit(V9, " ", fixed=TRUE)[2]]
GSC22340<-GSC22340[,c("V11","gene_id")]
GSC22340<-GSC22340[,gene_id := substr(gene_id, 2,19)]
GSC22340<-GSC22340[gene_id %in% genes_expressed,]
GSC22340<-GSC22340[ , ':=' (Counter= 1:.N, Length = .N, Sum = sum(V11)), by = gene_id ]
GSC22340<-GSC22340[V11 > 0,]
GSC22340<-GSC22340[Sum > 100,]
GSC22340<-GSC22340[Length > 500,]
GSC22340<-GSC22340[ , prop := round(V11 / Sum,5)]
GSC22340<-GSC22340[ , rel_position := round(Counter / Length,5)]
GSC22340<-GSC22340[ , c("gene_id", "prop" ,"rel_position")]

GSC22341<-data.table::fread(file="/coverage_files/coverage_GSC22341.txt.gz", sep = "\t", header = FALSE ,colClasses= c( rep("NULL", 8), "character", "NULL", "integer"), quote=FALSE, stringsAsFactors=FALSE, nThread=2, showProgress=FALSE)
GSC22341<-GSC22341[,"gene_id" := tstrsplit(V9, " ", fixed=TRUE)[2]]
GSC22341<-GSC22341[,c("V11","gene_id")]
GSC22341<-GSC22341[,gene_id := substr(gene_id, 2,19)]
GSC22341<-GSC22341[gene_id %in% genes_expressed,]
GSC22341<-GSC22341[ , ':=' (Counter= 1:.N, Length = .N, Sum = sum(V11)), by = gene_id ]
GSC22341<-GSC22341[V11 > 0,]
GSC22341<-GSC22341[Sum > 100,]
GSC22341<-GSC22341[Length > 500,]
GSC22341<-GSC22341[ , prop := round(V11 / Sum,5)]
GSC22341<-GSC22341[ , rel_position := round(Counter / Length,5)]
GSC22341<-GSC22341[ , c("gene_id", "prop" ,"rel_position")]

GSC22342<-data.table::fread(file="/coverage_files/coverage_GSC22342.txt.gz", sep = "\t", header = FALSE ,colClasses= c( rep("NULL", 8), "character", "NULL", "integer"), quote=FALSE, stringsAsFactors=FALSE, nThread=2, showProgress=FALSE)
GSC22342<-GSC22342[,"gene_id" := tstrsplit(V9, " ", fixed=TRUE)[2]]
GSC22342<-GSC22342[,c("V11","gene_id")]
GSC22342<-GSC22342[,gene_id := substr(gene_id, 2,19)]
GSC22342<-GSC22342[gene_id %in% genes_expressed,]
GSC22342<-GSC22342[ , ':=' (Counter= 1:.N, Length = .N, Sum = sum(V11)), by = gene_id ]
GSC22342<-GSC22342[V11 > 0,]
GSC22342<-GSC22342[Sum > 100,]
GSC22342<-GSC22342[Length > 500,]
GSC22342<-GSC22342[ , prop := round(V11 / Sum,5)]
GSC22342<-GSC22342[ , rel_position := round(Counter / Length,5)]
GSC22342<-GSC22342[ , c("gene_id", "prop" ,"rel_position")]

GSC22346<-data.table::fread(file="/coverage_files/coverage_GSC22346.txt.gz", sep = "\t", header = FALSE ,colClasses= c( rep("NULL", 8), "character", "NULL", "integer"), quote=FALSE, stringsAsFactors=FALSE, nThread=2, showProgress=FALSE)
GSC22346<-GSC22346[,"gene_id" := tstrsplit(V9, " ", fixed=TRUE)[2]]
GSC22346<-GSC22346[,c("V11","gene_id")]
GSC22346<-GSC22346[,gene_id := substr(gene_id, 2,19)]
GSC22346<-GSC22346[gene_id %in% genes_expressed,]
GSC22346<-GSC22346[ , ':=' (Counter= 1:.N, Length = .N, Sum = sum(V11)), by = gene_id ]
GSC22346<-GSC22346[V11 > 0,]
GSC22346<-GSC22346[Sum > 100,]
GSC22346<-GSC22346[Length > 500,]
GSC22346<-GSC22346[ , prop := round(V11 / Sum,5)]
GSC22346<-GSC22346[ , rel_position := round(Counter / Length,5)]
GSC22346<-GSC22346[ , c("gene_id", "prop" ,"rel_position")]

GSC22347<-data.table::fread(file="/coverage_files/coverage_GSC22347.txt.gz", sep = "\t", header = FALSE ,colClasses= c( rep("NULL", 8), "character", "NULL", "integer"), quote=FALSE, stringsAsFactors=FALSE, nThread=2, showProgress=FALSE)
GSC22347<-GSC22347[,"gene_id" := tstrsplit(V9, " ", fixed=TRUE)[2]]
GSC22347<-GSC22347[,c("V11","gene_id")]
GSC22347<-GSC22347[,gene_id := substr(gene_id, 2,19)]
GSC22347<-GSC22347[gene_id %in% genes_expressed,]
GSC22347<-GSC22347[ , ':=' (Counter= 1:.N, Length = .N, Sum = sum(V11)), by = gene_id ]
GSC22347<-GSC22347[V11 > 0,]
GSC22347<-GSC22347[Sum > 100,]
GSC22347<-GSC22347[Length > 500,]
GSC22347<-GSC22347[ , prop := round(V11 / Sum,5)]
GSC22347<-GSC22347[ , rel_position := round(Counter / Length,5)]
GSC22347<-GSC22347[ , c("gene_id", "prop" ,"rel_position")]

GSC22348<-data.table::fread(file="/coverage_files/coverage_GSC22348.txt.gz", sep = "\t", header = FALSE ,colClasses= c( rep("NULL", 8), "character", "NULL", "integer"), quote=FALSE, stringsAsFactors=FALSE, nThread=2, showProgress=FALSE)
GSC22348<-GSC22348[,"gene_id" := tstrsplit(V9, " ", fixed=TRUE)[2]]
GSC22348<-GSC22348[,c("V11","gene_id")]
GSC22348<-GSC22348[,gene_id := substr(gene_id, 2,19)]
GSC22348<-GSC22348[gene_id %in% genes_expressed,]
GSC22348<-GSC22348[ , ':=' (Counter= 1:.N, Length = .N, Sum = sum(V11)), by = gene_id ]
GSC22348<-GSC22348[V11 > 0,]
GSC22348<-GSC22348[Sum > 100,]
GSC22348<-GSC22348[Length > 500,]
GSC22348<-GSC22348[ , prop := round(V11 / Sum,5)]
GSC22348<-GSC22348[ , rel_position := round(Counter / Length,5)]
GSC22348<-GSC22348[ , c("gene_id", "prop" ,"rel_position")]

GSC22352<-data.table::fread(file="/coverage_files/coverage_GSC22352.txt.gz", sep = "\t", header = FALSE ,colClasses= c( rep("NULL", 8), "character", "NULL", "integer"), quote=FALSE, stringsAsFactors=FALSE, nThread=2, showProgress=FALSE)
GSC22352<-GSC22352[,"gene_id" := tstrsplit(V9, " ", fixed=TRUE)[2]]
GSC22352<-GSC22352[,c("V11","gene_id")]
GSC22352<-GSC22352[,gene_id := substr(gene_id, 2,19)]
GSC22352<-GSC22352[gene_id %in% genes_expressed,]
GSC22352<-GSC22352[ , ':=' (Counter= 1:.N, Length = .N, Sum = sum(V11)), by = gene_id ]
GSC22352<-GSC22352[V11 > 0,]
GSC22352<-GSC22352[Sum > 100,]
GSC22352<-GSC22352[Length > 500,]
GSC22352<-GSC22352[ , prop := round(V11 / Sum,5)]
GSC22352<-GSC22352[ , rel_position := round(Counter / Length,5)]
GSC22352<-GSC22352[ , c("gene_id", "prop" ,"rel_position")]

GSC22353<-data.table::fread(file="/coverage_files/coverage_GSC22353.txt.gz", sep = "\t", header = FALSE ,colClasses= c( rep("NULL", 8), "character", "NULL", "integer"), quote=FALSE, stringsAsFactors=FALSE, nThread=2, showProgress=FALSE)
GSC22353<-GSC22353[,"gene_id" := tstrsplit(V9, " ", fixed=TRUE)[2]]
GSC22353<-GSC22353[,c("V11","gene_id")]
GSC22353<-GSC22353[,gene_id := substr(gene_id, 2,19)]
GSC22353<-GSC22353[gene_id %in% genes_expressed,]
GSC22353<-GSC22353[ , ':=' (Counter= 1:.N, Length = .N, Sum = sum(V11)), by = gene_id ]
GSC22353<-GSC22353[V11 > 0,]
GSC22353<-GSC22353[Sum > 100,]
GSC22353<-GSC22353[Length > 500,]
GSC22353<-GSC22353[ , prop := round(V11 / Sum,5)]
GSC22353<-GSC22353[ , rel_position := round(Counter / Length,5)]
GSC22353<-GSC22353[ , c("gene_id", "prop" ,"rel_position")]

GSC22354<-data.table::fread(file="/coverage_files/coverage_GSC22354.txt.gz", sep = "\t", header = FALSE ,colClasses= c( rep("NULL", 8), "character", "NULL", "integer"), quote=FALSE, stringsAsFactors=FALSE, nThread=2, showProgress=FALSE)
GSC22354<-GSC22354[,"gene_id" := tstrsplit(V9, " ", fixed=TRUE)[2]]
GSC22354<-GSC22354[,c("V11","gene_id")]
GSC22354<-GSC22354[,gene_id := substr(gene_id, 2,19)]
GSC22354<-GSC22354[gene_id %in% genes_expressed,]
GSC22354<-GSC22354[ , ':=' (Counter= 1:.N, Length = .N, Sum = sum(V11)), by = gene_id ]
GSC22354<-GSC22354[V11 > 0,]
GSC22354<-GSC22354[Sum > 100,]
GSC22354<-GSC22354[Length > 500,]
GSC22354<-GSC22354[ , prop := round(V11 / Sum,5)]
GSC22354<-GSC22354[ , rel_position := round(Counter / Length,5)]
GSC22354<-GSC22354[ , c("gene_id", "prop" ,"rel_position")]

GSC22358<-data.table::fread(file="/coverage_files/coverage_GSC22358.txt.gz", sep = "\t", header = FALSE ,colClasses= c( rep("NULL", 8), "character", "NULL", "integer"), quote=FALSE, stringsAsFactors=FALSE, nThread=2, showProgress=FALSE)
GSC22358<-GSC22358[,"gene_id" := tstrsplit(V9, " ", fixed=TRUE)[2]]
GSC22358<-GSC22358[,c("V11","gene_id")]
GSC22358<-GSC22358[,gene_id := substr(gene_id, 2,19)]
GSC22358<-GSC22358[gene_id %in% genes_expressed,]
GSC22358<-GSC22358[ , ':=' (Counter= 1:.N, Length = .N, Sum = sum(V11)), by = gene_id ]
GSC22358<-GSC22358[V11 > 0,]
GSC22358<-GSC22358[Sum > 100,]
GSC22358<-GSC22358[Length > 500,]
GSC22358<-GSC22358[ , prop := round(V11 / Sum,5)]
GSC22358<-GSC22358[ , rel_position := round(Counter / Length,5)]
GSC22358<-GSC22358[ , c("gene_id", "prop" ,"rel_position")]

GSC22359<-data.table::fread(file="/coverage_files/coverage_GSC22359.txt.gz", sep = "\t", header = FALSE ,colClasses= c( rep("NULL", 8), "character", "NULL", "integer"), quote=FALSE, stringsAsFactors=FALSE, nThread=2, showProgress=FALSE)
GSC22359<-GSC22359[,"gene_id" := tstrsplit(V9, " ", fixed=TRUE)[2]]
GSC22359<-GSC22359[,c("V11","gene_id")]
GSC22359<-GSC22359[,gene_id := substr(gene_id, 2,19)]
GSC22359<-GSC22359[gene_id %in% genes_expressed,]
GSC22359<-GSC22359[ , ':=' (Counter= 1:.N, Length = .N, Sum = sum(V11)), by = gene_id ]
GSC22359<-GSC22359[V11 > 0,]
GSC22359<-GSC22359[Sum > 100,]
GSC22359<-GSC22359[Length > 500,]
GSC22359<-GSC22359[ , prop := round(V11 / Sum,5)]
GSC22359<-GSC22359[ , rel_position := round(Counter / Length,5)]
GSC22359<-GSC22359[ , c("gene_id", "prop" ,"rel_position")]

GSC22360<-data.table::fread(file="/coverage_files/coverage_GSC22360.txt.gz", sep = "\t", header = FALSE ,colClasses= c( rep("NULL", 8), "character", "NULL", "integer"), quote=FALSE, stringsAsFactors=FALSE, nThread=2, showProgress=FALSE)
GSC22360<-GSC22360[,"gene_id" := tstrsplit(V9, " ", fixed=TRUE)[2]]
GSC22360<-GSC22360[,c("V11","gene_id")]
GSC22360<-GSC22360[,gene_id := substr(gene_id, 2,19)]
GSC22360<-GSC22360[gene_id %in% genes_expressed,]
GSC22360<-GSC22360[ , ':=' (Counter= 1:.N, Length = .N, Sum = sum(V11)), by = gene_id ]
GSC22360<-GSC22360[V11 > 0,]
GSC22360<-GSC22360[Sum > 100,]
GSC22360<-GSC22360[Length > 500,]
GSC22360<-GSC22360[ , prop := round(V11 / Sum,5)]
GSC22360<-GSC22360[ , rel_position := round(Counter / Length,5)]
GSC22360<-GSC22360[ , c("gene_id", "prop" ,"rel_position")]

GSC22364<-data.table::fread(file="/coverage_files/coverage_GSC22364.txt.gz", sep = "\t", header = FALSE ,colClasses= c( rep("NULL", 8), "character", "NULL", "integer"), quote=FALSE, stringsAsFactors=FALSE, nThread=2, showProgress=FALSE)
GSC22364<-GSC22364[,"gene_id" := tstrsplit(V9, " ", fixed=TRUE)[2]]
GSC22364<-GSC22364[,c("V11","gene_id")]
GSC22364<-GSC22364[,gene_id := substr(gene_id, 2,19)]
GSC22364<-GSC22364[gene_id %in% genes_expressed,]
GSC22364<-GSC22364[ , ':=' (Counter= 1:.N, Length = .N, Sum = sum(V11)), by = gene_id ]
GSC22364<-GSC22364[V11 > 0,]
GSC22364<-GSC22364[Sum > 100,]
GSC22364<-GSC22364[Length > 500,]
GSC22364<-GSC22364[ , prop := round(V11 / Sum,5)]
GSC22364<-GSC22364[ , rel_position := round(Counter / Length,5)]
GSC22364<-GSC22364[ , c("gene_id", "prop" ,"rel_position")]


saveRDS(GSC22335,file="/GSC22335.rds")
saveRDS(GSC22336,file="/GSC22336.rds")
saveRDS(GSC22340,file="/GSC22340.rds")
saveRDS(GSC22341,file="/GSC22341.rds")
saveRDS(GSC22342,file="/GSC22342.rds")
saveRDS(GSC22346,file="/GSC22346.rds")
saveRDS(GSC22347,file="/GSC22347.rds")
saveRDS(GSC22348,file="/GSC22348.rds")
saveRDS(GSC22352,file="/GSC22352.rds")
saveRDS(GSC22353,file="/GSC22353.rds")
saveRDS(GSC22354,file="/GSC22354.rds")
saveRDS(GSC22358,file="/GSC22358.rds")
saveRDS(GSC22359,file="/GSC22359.rds")
saveRDS(GSC22360,file="/GSC22360.rds")
saveRDS(GSC22364,file="/GSC22364.rds")
```

##### Import summary of gene transcript coverage

```
GSC22335<-readRDS("/GSC22335.rds")
GSC22336<-readRDS("/GSC22336.rds")
GSC22340<-readRDS("/GSC22340.rds")
GSC22341<-readRDS("/GSC22341.rds")
GSC22342<-readRDS("/GSC22342.rds")
GSC22346<-readRDS("/GSC22346.rds")
GSC22347<-readRDS("/GSC22347.rds")
GSC22348<-readRDS("/GSC22348.rds")
GSC22352<-readRDS("/GSC22352.rds")
GSC22353<-readRDS("/GSC22353.rds")
GSC22354<-readRDS("/GSC22354.rds")
GSC22358<-readRDS("/GSC22358.rds")
GSC22359<-readRDS("/GSC22359.rds")
GSC22360<-readRDS("/GSC22360.rds")
GSC22364<-readRDS("/GSC22364.rds")
```

##### Test for differences in distribution of transcript coverage

```
GSC22335<-na.omit(GSC22335)
GSC22336<-na.omit(GSC22336)

ks_test_GSC22335_GSC22336<-data.frame()

cl <- makeCluster(34)
registerDoParallel(cl)
ks_test_GSC22335_GSC22336<-foreach(i = genes_expressed, .combine='rbind',.packages=c( "data.table", "spatstat"), .inorder=FALSE, .errorhandling='remove',.verbose=TRUE ) %dopar%  {
   test<-ks_weighted(GSC22335[gene_id==i]$rel_position, GSC22336[gene_id==i]$rel_position, GSC22335[gene_id==i]$prop, GSC22336[gene_id==i]$prop)
   data.frame(gene_id=i, statistic=test[[1]], p_value=test[[2]])
  }
stopCluster(cl)

setDT(ks_test_GSC22335_GSC22336)

ks_test_GSC22336_GSC22340<-data.frame()

cl <- makeCluster(34)
registerDoParallel(cl)
ks_test_GSC22336_GSC22340<-foreach(i = genes_expressed, .combine='rbind',.packages=c( "data.table", "spatstat"), .inorder=FALSE, .errorhandling='remove',.verbose=TRUE ) %dopar%  {
   test<-ks_weighted(GSC22336[gene_id==i]$rel_position, GSC22340[gene_id==i]$rel_position, GSC22336[gene_id==i]$prop, GSC22340[gene_id==i]$prop)
   data.frame(gene_id=i, statistic=test[[1]], p_value=test[[2]])
  }
stopCluster(cl)

setDT(ks_test_GSC22336_GSC22340)

##
ks_test_GSC22341_GSC22342<-data.frame()
cl <- makeCluster(34)
registerDoParallel(cl)
ks_test_GSC22341_GSC22342<-foreach(i = genes_expressed, .combine='rbind',.packages=c( "data.table", "spatstat"), .inorder=FALSE, .errorhandling='remove',.verbose=TRUE ) %dopar%  {
   test<-ks_weighted(GSC22341[gene_id==i]$rel_position, GSC22342[gene_id==i]$rel_position, GSC22341[gene_id==i]$prop, GSC22342[gene_id==i]$prop)
   data.frame(gene_id=i, statistic=test[[1]], p_value=test[[2]])
  }
stopCluster(cl)

setDT(ks_test_GSC22341_GSC22342)

ks_test_GSC22342_GSC22346<-data.frame()

cl <- makeCluster(34)
registerDoParallel(cl)
ks_test_GSC22342_GSC22346<-foreach(i = genes_expressed, .combine='rbind',.packages=c( "data.table", "spatstat"), .inorder=FALSE, .errorhandling='remove',.verbose=TRUE ) %dopar%  {
   test<-ks_weighted(GSC22342[gene_id==i]$rel_position, GSC22346[gene_id==i]$rel_position, GSC22342[gene_id==i]$prop, GSC22346[gene_id==i]$prop)
   data.frame(gene_id=i, statistic=test[[1]], p_value=test[[2]])
  }
stopCluster(cl)

setDT(ks_test_GSC22342_GSC22346)

##

ks_test_GSC22347_GSC22348<-data.frame()

cl <- makeCluster(34)
registerDoParallel(cl)
ks_test_GSC22347_GSC22348<-foreach(i = genes_expressed, .combine='rbind',.packages=c( "data.table", "spatstat"), .inorder=FALSE, .errorhandling='remove',.verbose=TRUE ) %dopar%  {
   test<-ks_weighted(GSC22347[gene_id==i]$rel_position, GSC22348[gene_id==i]$rel_position, GSC22347[gene_id==i]$prop, GSC22348[gene_id==i]$prop)
   data.frame(gene_id=i, statistic=test[[1]], p_value=test[[2]])
  }
stopCluster(cl)

setDT(ks_test_GSC22347_GSC22348)


ks_test_GSC22348_GSC22352<-data.frame()

cl <- makeCluster(34)
registerDoParallel(cl)
ks_test_GSC22348_GSC22352<-foreach(i = genes_expressed, .combine='rbind',.packages=c( "data.table", "spatstat"), .inorder=FALSE, .errorhandling='remove',.verbose=TRUE ) %dopar%  {
   test<-ks_weighted(GSC22348[gene_id==i]$rel_position, GSC22352[gene_id==i]$rel_position, GSC22348[gene_id==i]$prop, GSC22352[gene_id==i]$prop)
   data.frame(gene_id=i, statistic=test[[1]], p_value=test[[2]])
  }
stopCluster(cl)

setDT(ks_test_GSC22348_GSC22352)

##

ks_test_GSC22353_GSC22354<-data.frame()

cl <- makeCluster(34)
registerDoParallel(cl)
ks_test_GSC22353_GSC22354<-foreach(i = genes_expressed, .combine='rbind',.packages=c( "data.table", "spatstat"), .inorder=FALSE, .errorhandling='remove',.verbose=TRUE ) %dopar%  {
   test<-ks_weighted(GSC22353[gene_id==i]$rel_position, GSC22354[gene_id==i]$rel_position, GSC22353[gene_id==i]$prop, GSC22354[gene_id==i]$prop)
   data.frame(gene_id=i, statistic=test[[1]], p_value=test[[2]])
  }
stopCluster(cl)

setDT(ks_test_GSC22353_GSC22354)

ks_test_GSC22354_GSC22358<-data.frame()

cl <- makeCluster(34)
registerDoParallel(cl)
ks_test_GSC22354_GSC22358<-foreach(i = genes_expressed, .combine='rbind',.packages=c( "data.table", "spatstat"), .inorder=FALSE, .errorhandling='remove',.verbose=TRUE ) %dopar%  {
   test<-ks_weighted(GSC22354[gene_id==i]$rel_position, GSC22358[gene_id==i]$rel_position, GSC22354[gene_id==i]$prop, GSC22358[gene_id==i]$prop)
   data.frame(gene_id=i, statistic=test[[1]], p_value=test[[2]])
  }
stopCluster(cl)

setDT(ks_test_GSC22354_GSC22358)

##

ks_test_GSC22359_GSC22360<-data.frame()

cl <- makeCluster(34)
registerDoParallel(cl)
ks_test_GSC22359_GSC22360<-foreach(i = genes_expressed, .combine='rbind',.packages=c( "data.table", "spatstat"), .inorder=FALSE, .errorhandling='remove',.verbose=TRUE ) %dopar%  {
   test<-ks_weighted(GSC22359[gene_id==i]$rel_position, GSC22360[gene_id==i]$rel_position, GSC22359[gene_id==i]$prop, GSC22360[gene_id==i]$prop)
   data.frame(gene_id=i, statistic=test[[1]], p_value=test[[2]])
  }
stopCluster(cl)

setDT(ks_test_GSC22359_GSC22360)

ks_test_GSC22360_GSC22364<-data.frame()

cl <- makeCluster(34)
registerDoParallel(cl)
ks_test_GSC22360_GSC22364<-foreach(i = genes_expressed, .combine='rbind',.packages=c( "data.table", "spatstat"), .inorder=FALSE, .errorhandling='remove',.verbose=TRUE ) %dopar%  {
   test<-ks_weighted(GSC22360[gene_id==i]$rel_position, GSC22364[gene_id==i]$rel_position, GSC22360[gene_id==i]$prop, GSC22364[gene_id==i]$prop)
   data.frame(gene_id=i, statistic=test[[1]], p_value=test[[2]])
  }
stopCluster(cl)

setDT(ks_test_GSC22360_GSC22364)

saveRDS( ks_test_GSC22335_GSC22336,file="/ks_test_GSC22335_GSC22336.rds")
saveRDS( ks_test_GSC22336_GSC22340,file="/ks_test_GSC22336_GSC22340.rds")
saveRDS( ks_test_GSC22341_GSC22342,file="/ks_test_GSC22341_GSC22342.rds")
saveRDS( ks_test_GSC22342_GSC22346,file="/ks_test_GSC22342_GSC22346.rds")
saveRDS( ks_test_GSC22347_GSC22348,file="/ks_test_GSC22347_GSC22348.rds")
saveRDS( ks_test_GSC22348_GSC22352,file="/ks_test_GSC22348_GSC22352.rds")
saveRDS( ks_test_GSC22353_GSC22354,file="/ks_test_GSC22353_GSC22354.rds")
saveRDS( ks_test_GSC22354_GSC22358,file="/ks_test_GSC22354_GSC22358.rds")
saveRDS( ks_test_GSC22359_GSC22360,file="/ks_test_GSC22359_GSC22360.rds")
saveRDS( ks_test_GSC22360_GSC22364,file="/ks_test_GSC22360_GSC22364.rds")
```

##### Import resutls from the Kolmogorov-Smirnov test of distribution

```
ks_test_GSC22335_GSC22336<-readRDS( file="/ks_test_GSC22335_GSC22336.rds")
ks_test_GSC22336_GSC22340<-readRDS( file="/ks_test_GSC22336_GSC22340.rds")
ks_test_GSC22341_GSC22342<-readRDS( file="/ks_test_GSC22341_GSC22342.rds")
ks_test_GSC22342_GSC22346<-readRDS( file="/ks_test_GSC22342_GSC22346.rds")
ks_test_GSC22347_GSC22348<-readRDS( file="/ks_test_GSC22347_GSC22348.rds")
ks_test_GSC22348_GSC22352<-readRDS( file="/ks_test_GSC22348_GSC22352.rds")
ks_test_GSC22353_GSC22354<-readRDS( file="/ks_test_GSC22353_GSC22354.rds")
ks_test_GSC22354_GSC22358<-readRDS( file="/ks_test_GSC22354_GSC22358.rds")
ks_test_GSC22359_GSC22360<-readRDS( file="/ks_test_GSC22359_GSC22360.rds")
ks_test_GSC22360_GSC22364<-readRDS( file="/ks_test_GSC22360_GSC22364.rds")
ks_test_GSC22335_GSC22336<-readRDS( file="/ks_test_GSC22335_GSC22336.rds")
```

##### Import resutls of DEGs based on pregnancy outcome from Moorey et al (2020)

```
DEGs_pregnancy_outcome_past_publications<-as.data.frame(read_excel("/DEGs_pregnancy_outcome.xlsx"))
```

##### Save all objects into one .RData file

```
rm("bias_metrics","bias_metrics_a", "counting_summary","counting_summary_a" , "files" ,"count","sample")
save.image( file = "/resource_data_2022_01_05.RData", compress="bzip2", safe=TRUE)
```

#### Analytical procedures

Load all libraries.

```
.libPaths("/usr/lib/R/site-library")

library('dplyr', quietly = TRUE,lib.loc="/usr/lib/R/site-library")
library('ggplot2', quietly = TRUE,lib.loc="/usr/lib/R/site-library")
library('edgeR',quietly = TRUE, lib.loc="/usr/lib/R/site-library")
library("cowplot", quietly = TRUE,lib.loc="/usr/lib/R/site-library")
library("GGally",quietly = TRUE,lib.loc="/usr/lib/R/site-library")
library("DESeq2",quietly = TRUE,lib.loc="/usr/lib/R/site-library")
library('readxl',quietly = TRUE,lib.loc="/usr/lib/R/site-library")
library('DEXSeq',quietly = TRUE,lib.loc="/usr/lib/R/site-library")
library("VennDiagram", quietly = TRUE, lib.loc="/usr/lib/R/site-library")
library("ComplexHeatmap",quietly = TRUE, lib.loc="/usr/lib/R/site-library")
library("flashClust", quietly = TRUE, lib.loc="/usr/lib/R/site-library")
library("circlize", lib.loc="/usr/lib/R/site-library")
library("nlme", quietly = TRUE, lib.loc="/usr/lib/R/site-library")
library("multcomp", quietly = TRUE, lib.loc="/usr/lib/R/site-library")
library("emmeans", quietly = TRUE, lib.loc="/usr/lib/R/site-library")
library('plotly' , quietly = TRUE, lib.loc="/usr/lib/R/site-library")
library('tidyverse' ,quietly = TRUE , lib.loc="/usr/lib/R/site-library")
library('htmlwidgets' , quietly = TRUE, lib.loc="/usr/lib/R/site-library")
library('reshape2' ,quietly = TRUE, lib.loc="/usr/lib/R/site-library")
library("lme4" ,quietly = TRUE, lib.loc="/usr/lib/R/site-library")
library("predictmeans" ,quietly = TRUE, lib.loc="/usr/lib/R/site-library")
library("fitdistrplus" ,quietly = TRUE, lib.loc="/usr/lib/R/site-library")
library("logspline" ,quietly = TRUE, lib.loc="/usr/lib/R/site-library")
library("betareg" ,quietly = TRUE, lib.loc="/usr/lib/R/site-library")
library("ggpubr" ,quietly = TRUE, lib.loc="/usr/lib/R/site-library")
library("glmmTMB" ,quietly = TRUE, lib.loc="/usr/lib/R/site-library") 
library("car" ,quietly = TRUE, lib.loc="/usr/lib/R/site-library") 
library("goseq" ,quietly = TRUE, lib.loc="/usr/lib/R/site-library")
library("Rsamtools", quietly = TRUE, lib.loc="/usr/lib/R/site-library")
library("stringr",quietly = TRUE , lib.loc="/usr/lib/R/site-library")
library("dgof",quietly = TRUE , lib.loc="/usr/lib/R/site-library")
library("data.table",quietly = TRUE , lib.loc="/usr/lib/R/site-library")
library("tidyr",quietly = TRUE , lib.loc="/usr/lib/R/site-library")
library("dplyr",quietly = TRUE , lib.loc="/usr/lib/R/site-library")
library("ggsignif",quietly = TRUE , lib.loc="/usr/lib/R/site-library")
library("kableExtra",quietly = TRUE , lib.loc="/usr/lib/R/site-library")
library("tidyverse",quietly = TRUE , lib.loc="/usr/lib/R/site-library")
library("grid",quietly = TRUE , lib.loc="/usr/lib/R/site-library")
library("gridExtra",quietly = TRUE , lib.loc="/usr/lib/R/site-library")
```

Load all the objects required to reproduce our results.

```
load( file = "/mnt/storage/lab_folder/shared_R_codes/fernando/RNA_degradation/resources/resource_data_2022_01_05.RData")
gc()
```

Test for Normality of RNA parameters.

```
fitdistrplus::descdist(log2(RNA_data_frame$RIN), discrete = FALSE, boot=1000)

fn <- fitdistrplus::fitdist(RNA_data_frame$RIN, "norm", method = "mle")
flg <- fitdistrplus::fitdist(RNA_data_frame$RIN, "logis", method = "mle")
fw <- fitdistrplus::fitdist(RNA_data_frame$RIN, "weibull", method = "mle")
fln <- fitdistrplus::fitdist(RNA_data_frame$RIN, "lnorm", method = "mle")
fg <- fitdistrplus::fitdist(RNA_data_frame$RIN, "gamma", method = "mle")

fitdistrplus::gofstat(list(fn, flg,  fw, fln, fg), discrete=FALSE, fitnames= c("Normal", "Logistic","Weibull", "lognormal", "gamma"))
fitdistrplus::gofstat(list(fn, flg,  fw, fln, fg), discrete=FALSE, fitnames= c("Normal", "Logistic","Weibull", "lognormal", "gamma"))$chisqpvalue
fitdistrplus::gofstat(list(fn, flg,  fw, fln, fg), discrete=FALSE, fitnames= c("Normal", "Logistic","Weibull", "lognormal", "gamma"))$cvmtest
fitdistrplus::gofstat(list(fn, flg,  fw, fln, fg), discrete=FALSE, fitnames= c("Normal", "Logistic","Weibull", "lognormal", "gamma"))$adtest

plot.legend <- c("Normal", "Logistic","Weibull", "lognormal", "gamma")
par(mfrow = c(2, 2))
fitdistrplus::denscomp(list(fn, flg,  fw, fln, fg))
fitdistrplus::qqcomp(list(fn, flg,  fw, fln, fg))
fitdistrplus::ppcomp(list(fn, flg,  fw, fln, fg))
fitdistrplus::cdfcomp(list(fn, flg,  fw, fln, fg), xlogscale=TRUE, ylogscale=TRUE,legendtext = plot.legend)


fitdistrplus::descdist(log2(RNA_data_frame$ratio), discrete = FALSE, boot=1000)
fg <- fitdistrplus::fitdist(RNA_data_frame$ratio, "gamma", method = "mle")
fn <- fitdistrplus::fitdist(RNA_data_frame$ratio, "norm", method = "mle")
fw <- fitdistrplus::fitdist(RNA_data_frame$ratio, "weibull", method = "mle")

fitdistrplus::gofstat(list(fn,  fg, fw), discrete=FALSE, fitnames= c("Normal",  "gamma", "weibull"))
fitdistrplus::gofstat(list(fn,  fg, fw), discrete=FALSE, fitnames= c("Normal",  "gamma", "weibull"))$chisqpvalue
fitdistrplus::gofstat(list(fn,  fg, fw), discrete=FALSE, fitnames= c("Normal",  "gamma", "weibull"))$cvmtest
fitdistrplus::gofstat(list(fn, fg, fw), discrete=FALSE, fitnames= c("Normal",  "gamma", "weibull"))$adtest

fitdistrplus::cdfcomp(list(fn,   fw,  fg), xlogscale=TRUE, ylogscale=TRUE)

plot.legend <- c("Normal","gamma", "Weibull"  )
par(mfrow = c(2, 2))
fitdistrplus::denscomp(list(fn,   fw,  fg))
fitdistrplus::qqcomp(list(fn,   fw,  fg))
fitdistrplus::ppcomp(list(fn,   fw,  fg))
fitdistrplus::cdfcomp(list(fn,   fw,  fg), xlogscale=TRUE, ylogscale=TRUE,legendtext = plot.legend)
```

##### Table 1

```
RNA_data_table<-as.data.table(RNA_data_frame)
RNA_data_frame_summary<-RNA_data_table[, .(RIN_average=round(mean(RIN),2),RIN_sd=round(sd(RIN),2), 
                                                  A260_average=round(mean(A260),2),A260_sd=round(sd(A260),2),
                                                  A280_average=round(mean(A280),2),A280_sd=round(sd(A280),2),
                                                  ratio_average=round(mean(ratio),2),ratio_sd=round(sd(ratio),2)),  by = c('location_blood_drawing','Time_pre_processing') ]

RNA_data_frame_summary[location_blood_drawing == "tail", location_blood_drawing := "coccygeal"  ]

colnames(RNA_data_frame_summary)<-c("Vein","Processing time","RIN_average","RIN_sd","ratio_average","ratio_sd","A260_average","A260_sd","A280_average","A280_sd" )
kbl(RNA_data_frame_summary, col.names=c("Vein","Processing time","$\\overline{x}$","$\\hat{\\sigma}$","$\\bar{x}$","$\\hat{\\sigma}$","$\\bar{x}$","$\\hat{\\sigma}$","$\\bar{x}$","$\\hat{\\sigma}$"),
    caption = "Table 1. RNA parameters obtained from peripheral white blood cells.") %>%
  kable_classic() %>%
#  kable_styling(full_width = F) %>%
  add_header_above(c(" "=2, "RIN"=2, "A260"=2, "A280"=2,"Ratio"=2)) %>%
  #pack_rows(group_label="Coccygeal", 1, 1) %>%
  #pack_rows(group_label="Jugular", 2, 6) %>%
  collapse_rows(columns = 1, valign = "top") %>%
  column_spec(column = 1, width = "1.1in")  %>%
  column_spec(column = 3:9, width = "0.5in")
```

Table 1. RNA parameters obtained from peripheral white blood cells.

|  | | RIN | | A260 | | A280 | | Ratio | |
| --- | --- | --- | --- | --- | --- | --- | --- | --- | --- |
| Vein | Processing time | \(\overline{x}\) | \(\hat{\sigma}\) | \(\bar{x}\) | \(\hat{\sigma}\) | \(\bar{x}\) | \(\hat{\sigma}\) | \(\bar{x}\) | \(\hat{\sigma}\) |
| coccygeal | 1h | 8.48 | 0.16 | 13.27 | 4.02 | 6.85 | 2.03 | 1.93 | 0.02 |
| jugular | 1h | 8.52 | 0.37 | 15.99 | 2.75 | 8.28 | 1.37 | 1.93 | 0.02 |
|  | 3h | 8.60 | 0.23 | 17.74 | 6.34 | 9.19 | 3.30 | 1.93 | 0.01 |
|  | 6h | 8.66 | 0.23 | 13.76 | 4.84 | 7.12 | 2.46 | 1.93 | 0.02 |
|  | 8h | 8.34 | 0.33 | 12.40 | 6.85 | 6.40 | 3.49 | 1.91 | 0.06 |
|  | 24h | 8.00 | 0.37 | 14.48 | 2.52 | 7.53 | 1.23 | 1.92 | 0.03 |

##### Compare RNA parameters between Coccygeal vs Jugular

```
tail_jugular<-RNA_data_frame[RNA_data_frame$Time_pre_processing=="1h",]
tail_jugular<-tail_jugular[order(tail_jugular$location_blood_drawing, tail_jugular$Heifer_ID),]
tail_jugular$location_blood_drawing<-as.factor(tail_jugular$location_blood_drawing)
tail_jugular$Heifer_ID<-as.factor(tail_jugular$Heifer_ID)


RIN_tail<-tail_jugular[tail_jugular$location_blood_drawing=="tail",7]
RIN_jugular<-tail_jugular[tail_jugular$location_blood_drawing=="jugular",7]
wilcox.test(RIN_tail, RIN_jugular, paired = TRUE, alternative = "two.sided", exact = FALSE)
```

```
## 
##  Wilcoxon signed rank test with continuity correction
## 
## data:  RIN_tail and RIN_jugular
## V = 5, p-value = 1
## alternative hypothesis: true location shift is not equal to 0
```

```
coin::wilcoxsign_test(RIN ~ location_blood_drawing | Heifer_ID, data = tail_jugular,alternative = "two.sided", distribution="exact", zero.method =  "Wilcoxon")
```

```
## 
##  Exact Wilcoxon Signed-Rank Test
## 
## data:  y by x (pos, neg) 
##   stratified by block
## Z = 0, p-value = 1
## alternative hypothesis: true mu is not equal to 0
```

```
t.test(RIN_tail, RIN_jugular, paired=TRUE, alternate="two.sided")
```

```
## 
##  Paired t-test
## 
## data:  RIN_tail and RIN_jugular
## t = -0.1853, df = 4, p-value = 0.862
## alternative hypothesis: true difference in means is not equal to 0
## 95 percent confidence interval:
##  -0.6393521  0.5593521
## sample estimates:
## mean of the differences 
##                   -0.04
```

```
aggregate(RIN ~ location_blood_drawing, tail_jugular , function(x) c(mean = round(mean(x),digits = 2), sd = round(sd(x), digits = 2)))
```

```
##   location_blood_drawing RIN.mean RIN.sd
## 1                jugular     8.52   0.37
## 2                   tail     8.48   0.16
```

```
A260_tail<-tail_jugular[tail_jugular$location_blood_drawing=="tail",9]
A260_jugular<-tail_jugular[tail_jugular$location_blood_drawing=="jugular",9]
wilcox.test(A260_tail, A260_jugular, paired = TRUE, alternative = "two.sided", exact = FALSE)
```

```
## 
##  Wilcoxon signed rank test with continuity correction
## 
## data:  A260_tail and A260_jugular
## V = 0, p-value = 0.05906
## alternative hypothesis: true location shift is not equal to 0
```

```
t.test(A260_tail,A260_jugular , paired=TRUE, alternate="two.sided")
```

```
## 
##  Paired t-test
## 
## data:  A260_tail and A260_jugular
## t = -2.3729, df = 4, p-value = 0.07658
## alternative hypothesis: true difference in means is not equal to 0
## 95 percent confidence interval:
##  -5.9086253  0.4630253
## sample estimates:
## mean of the differences 
##                 -2.7228
```

```
aggregate(A260 ~ location_blood_drawing, tail_jugular , function(x) c(mean = round(mean(x),digits = 2), sd = round(sd(x), digits = 2)))
```

```
##   location_blood_drawing A260.mean A260.sd
## 1                jugular     15.99    2.75
## 2                   tail     13.27    4.02
```

```
A280_tail<-tail_jugular[tail_jugular$location_blood_drawing=="tail",10]
A280_jugular<-tail_jugular[tail_jugular$location_blood_drawing=="jugular",10]
wilcox.test(A280_tail, A280_jugular, paired = TRUE, alternative = "two.sided", exact = FALSE)
```

```
## 
##  Wilcoxon signed rank test with continuity correction
## 
## data:  A280_tail and A280_jugular
## V = 0, p-value = 0.05906
## alternative hypothesis: true location shift is not equal to 0
```

```
t.test(A280_tail, A280_jugular , paired=TRUE, alternate="two.sided")
```

```
## 
##  Paired t-test
## 
## data:  A280_tail and A280_jugular
## t = -2.4984, df = 4, p-value = 0.06688
## alternative hypothesis: true difference in means is not equal to 0
## 95 percent confidence interval:
##  -3.0035013  0.1583013
## sample estimates:
## mean of the differences 
##                 -1.4226
```

```
aggregate(A280 ~ location_blood_drawing, tail_jugular , function(x) c(mean = round(mean(x),digits = 2), sd = round(sd(x), digits = 2)))
```

```
##   location_blood_drawing A280.mean A280.sd
## 1                jugular      8.28    1.37
## 2                   tail      6.85    2.03
```

```
A260_A28_ratio_tail<-tail_jugular[tail_jugular$location_blood_drawing=="tail",8]
A260_A28_ratio_jugular<-tail_jugular[tail_jugular$location_blood_drawing=="jugular",8]
wilcox.test(A260_A28_ratio_tail, A260_A28_ratio_jugular, paired = TRUE, alternative = "two.sided", exact = FALSE)
```

```
## 
##  Wilcoxon signed rank test with continuity correction
## 
## data:  A260_A28_ratio_tail and A260_A28_ratio_jugular
## V = 7, p-value = 1
## alternative hypothesis: true location shift is not equal to 0
```

```
coin::wilcoxsign_test(ratio ~ location_blood_drawing | Heifer_ID, data = tail_jugular,alternative = "two.sided", distribution="exact", zero.method =  "Wilcoxon")
```

```
## 
##  Exact Wilcoxon Signed-Rank Test
## 
## data:  y by x (pos, neg) 
##   stratified by block
## Z = 0.13546, p-value = 1
## alternative hypothesis: true mu is not equal to 0
```

```
t.test(A260_A28_ratio_tail, A260_A28_ratio_jugular, paired = TRUE, alternative = "two.sided")
```

```
## 
##  Paired t-test
## 
## data:  A260_A28_ratio_tail and A260_A28_ratio_jugular
## t = -0.16116, df = 4, p-value = 0.8798
## alternative hypothesis: true difference in means is not equal to 0
## 95 percent confidence interval:
##  -0.03645478  0.03245478
## sample estimates:
## mean of the differences 
##                  -0.002
```

```
aggregate(ratio ~ location_blood_drawing, tail_jugular , function(x) c(mean = round(mean(x),digits = 2), sd = round(sd(x), digits = 2)))
```

```
##   location_blood_drawing ratio.mean ratio.sd
## 1                jugular       1.93     0.02
## 2                   tail       1.93     0.02
```

##### Processing Time Quality Parameters

```
processing_time <- RNA_data_frame[RNA_data_frame$location_blood_drawing=="jugular",]
processing_time$Heifer_ID <- as.factor(processing_time$Heifer_ID)
processing_time$Time_pre_processing <- factor(processing_time$Time_pre_processing, levels=c('1h', '3h', '6h', '8h','24h'))

##assuming normality of the sample
random_RIN_lme<-nlme::lme(RIN ~ Time_pre_processing,random=~ 1 |Heifer_ID,data=processing_time,method="REML")
summary(random_RIN_lme)
```

```
## Linear mixed-effects model fit by REML
##   Data: processing_time 
##        AIC      BIC    logLik
##   32.47105 39.44118 -9.235525
## 
## Random effects:
##  Formula: ~1 | Heifer_ID
##         (Intercept)  Residual
## StdDev: 4.68602e-06 0.3140064
## 
## Fixed effects:  RIN ~ Time_pre_processing 
##                        Value Std.Error DF  t-value p-value
## (Intercept)             8.52 0.1404279 16 60.67170  0.0000
## Time_pre_processing3h   0.08 0.1985951 16  0.40283  0.6924
## Time_pre_processing6h   0.14 0.1985951 16  0.70495  0.4910
## Time_pre_processing8h  -0.18 0.1985951 16 -0.90637  0.3782
## Time_pre_processing24h -0.52 0.1985951 16 -2.61839  0.0186
##  Correlation: 
##                        (Intr) Tm_p_3 Tm_p_6 Tm_p_8
## Time_pre_processing3h  -0.707                     
## Time_pre_processing6h  -0.707  0.500              
## Time_pre_processing8h  -0.707  0.500  0.500       
## Time_pre_processing24h -0.707  0.500  0.500  0.500
## 
## Standardized Within-Group Residuals:
##        Min         Q1        Med         Q3        Max 
## -1.9107893 -0.8280087  0.1273860  0.6369298  1.8470963 
## 
## Number of Observations: 25
## Number of Groups: 5
```

```
anova(random_RIN_lme)
```

```
##                     numDF denDF   F-value p-value
## (Intercept)             1    16 17992.844  <.0001
## Time_pre_processing     4    16     3.584  0.0286
```

```
dunnett_random_RIN_lme<-(glht(random_RIN_lme, mcp(Time_pre_processing="Dunnett")))
summary( dunnett_random_RIN_lme)
```

```
## 
##   Simultaneous Tests for General Linear Hypotheses
## 
## Multiple Comparisons of Means: Dunnett Contrasts
## 
## 
## Fit: lme.formula(fixed = RIN ~ Time_pre_processing, data = processing_time, 
##     random = ~1 | Heifer_ID, method = "REML")
## 
## Linear Hypotheses:
##               Estimate Std. Error z value Pr(>|z|)  
## 3h - 1h == 0    0.0800     0.1986   0.403    0.984  
## 6h - 1h == 0    0.1400     0.1986   0.705    0.890  
## 8h - 1h == 0   -0.1800     0.1986  -0.906    0.774  
## 24h - 1h == 0  -0.5200     0.1986  -2.618    0.031 *
## ---
## Signif. codes:  0 '***' 0.001 '**' 0.01 '*' 0.05 '.' 0.1 ' ' 1
## (Adjusted p values reported -- single-step method)
```

```
random_A260_lme<-nlme::lme(A260 ~ Time_pre_processing,random=~ 1 |Heifer_ID,data=processing_time,method="REML")
summary(random_A260_lme)
```

```
## Linear mixed-effects model fit by REML
##   Data: processing_time 
##        AIC      BIC   logLik
##   141.7994 148.7695 -63.8997
## 
## Random effects:
##  Formula: ~1 | Heifer_ID
##         (Intercept) Residual
## StdDev:     2.27162 4.442701
## 
## Fixed effects:  A260 ~ Time_pre_processing 
##                          Value Std.Error DF   t-value p-value
## (Intercept)            15.9912  2.231495 16  7.166138  0.0000
## Time_pre_processing3h   1.7446  2.809811 16  0.620896  0.5434
## Time_pre_processing6h  -2.2344  2.809811 16 -0.795214  0.4381
## Time_pre_processing8h  -3.5946  2.809811 16 -1.279303  0.2190
## Time_pre_processing24h -1.5124  2.809811 16 -0.538257  0.5978
##  Correlation: 
##                        (Intr) Tm_p_3 Tm_p_6 Tm_p_8
## Time_pre_processing3h  -0.63                      
## Time_pre_processing6h  -0.63   0.50               
## Time_pre_processing8h  -0.63   0.50   0.50        
## Time_pre_processing24h -0.63   0.50   0.50   0.50 
## 
## Standardized Within-Group Residuals:
##        Min         Q1        Med         Q3        Max 
## -2.5595668 -0.3423127  0.1180384  0.2958148  2.0742808 
## 
## Number of Observations: 25
## Number of Groups: 5
```

```
anova(random_A260_lme)
```

```
##                     numDF denDF   F-value p-value
## (Intercept)             1    16 121.41914  <.0001
## Time_pre_processing     4    16   1.07535  0.4012
```

```
random_A280_lme<-nlme::lme(A280 ~ Time_pre_processing,random=~ 1 |Heifer_ID,data=processing_time,method="REML")
summary(random_A280_lme)
```

```
## Linear mixed-effects model fit by REML
##   Data: processing_time 
##        AIC      BIC    logLik
##   115.1196 122.0897 -50.55978
## 
## Random effects:
##  Formula: ~1 | Heifer_ID
##         (Intercept) Residual
## StdDev:    1.116697 2.292509
## 
## Fixed effects:  A280 ~ Time_pre_processing 
##                          Value Std.Error DF   t-value p-value
## (Intercept)             8.2766  1.140404 16  7.257602  0.0000
## Time_pre_processing3h   0.9144  1.449910 16  0.630660  0.5372
## Time_pre_processing6h  -1.1530  1.449910 16 -0.795222  0.4381
## Time_pre_processing8h  -1.8780  1.449910 16 -1.295253  0.2136
## Time_pre_processing24h -0.7488  1.449910 16 -0.516446  0.6126
##  Correlation: 
##                        (Intr) Tm_p_3 Tm_p_6 Tm_p_8
## Time_pre_processing3h  -0.636                     
## Time_pre_processing6h  -0.636  0.500              
## Time_pre_processing8h  -0.636  0.500  0.500       
## Time_pre_processing24h -0.636  0.500  0.500  0.500
## 
## Standardized Within-Group Residuals:
##        Min         Q1        Med         Q3        Max 
## -2.5557700 -0.3469290  0.1199870  0.2990605  2.0861313 
## 
## Number of Observations: 25
## Number of Groups: 5
```

```
anova(random_A280_lme)
```

```
##                     numDF denDF   F-value p-value
## (Intercept)             1    16 129.11408  <.0001
## Time_pre_processing     4    16   1.09669  0.3917
```

```
random_ratio<-lme4::glmer(ratio ~ Time_pre_processing + (1|Heifer_ID), family = Gamma(link = "inverse"), data=processing_time)
summary(random_ratio)
```

```
## Generalized linear mixed model fit by maximum likelihood (Laplace
##   Approximation) [glmerMod]
##  Family: Gamma  ( inverse )
## Formula: ratio ~ Time_pre_processing + (1 | Heifer_ID)
##    Data: processing_time
## 
##      AIC      BIC   logLik deviance df.resid 
##    -97.7    -89.2     55.8   -111.7       18 
## 
## Scaled residuals: 
##     Min      1Q  Median      3Q     Max 
## -3.4659 -0.3746  0.0926  0.6006  1.4212 
## 
## Random effects:
##  Groups    Name        Variance  Std.Dev.
##  Heifer_ID (Intercept) 2.031e-05 0.004507
##  Residual              1.602e-04 0.012658
## Number of obs: 25, groups:  Heifer_ID, 5
## 
## Fixed effects:
##                         Estimate Std. Error t value Pr(>|z|)    
## (Intercept)            0.5177427  0.0046430 111.511   <2e-16 ***
## Time_pre_processing3h  0.0005362  0.0037581   0.143    0.887    
## Time_pre_processing6h  0.0016123  0.0037620   0.429    0.668    
## Time_pre_processing8h  0.0059612  0.0037778   1.578    0.115    
## Time_pre_processing24h 0.0032346  0.0037679   0.858    0.391    
## ---
## Signif. codes:  0 '***' 0.001 '**' 0.01 '*' 0.05 '.' 0.1 ' ' 1
## 
## Correlation of Fixed Effects:
##             (Intr) Tm_p_3 Tm_p_6 Tm_p_8
## Tm_pr_prcs3 -0.404                     
## Tm_pr_prcs6 -0.404  0.499              
## Tm_pr_prcs8 -0.402  0.497  0.496       
## Tm_pr_prc24 -0.403  0.498  0.498  0.496
```

```
anova(random_ratio)
```

```
## Analysis of Variance Table
##                     npar     Sum Sq    Mean Sq F value
## Time_pre_processing    4 0.00042579 0.00010645  0.6644
```

##### Additional file 1

Ribonucleic acid quality and abundance parameters from samples collected from the jugular and coccygeal veins and processed within one hour of sampling. Animals are indicated by shapes across charts.

```
tail_jugular$Heifer_ID <- as.factor(tail_jugular$Heifer_ID)
tail_jugular$location_blood_drawing <- as.factor(tail_jugular$location_blood_drawing)

font_size<-10
shape_size<-2

summary_RIN_location<-aggregate(RIN ~ location_blood_drawing, tail_jugular , function(x) c(mean = mean(x), sd = sd(x)))
summary_RIN_location<-do.call(data.frame,summary_RIN_location)
plot1<-ggplot(tail_jugular, aes(x=location_blood_drawing, y=RIN)) + 
  geom_point(data = summary_RIN_location, mapping = aes(x =location_blood_drawing, y= RIN.mean),size=1, color="darkgray") +
  geom_errorbar(data=summary_RIN_location, mapping = aes( x=location_blood_drawing, y=RIN.mean, ymin = RIN.mean - RIN.sd, ymax = RIN.mean + RIN.sd), width = 0.05,color="darkgray")+
  geom_jitter(position = position_jitter(w=0.3, h=0), aes(shape=Heifer_ID, fill=location_blood_drawing), size=shape_size) +
  scale_fill_manual(values = c("#9964cb","#5fa369")) +
  scale_shape_manual(values=c(21:25))+
  scale_color_manual(values="black")+
  scale_x_discrete(name=NULL,breaks=c("jugular","tail"), labels=c("Jugular","Tail")) +
  ylim(7,10) +
  theme_classic() + 
  theme(legend.position = "none",
        panel.grid = element_blank(),
        panel.background = element_blank(),
        axis.text = element_text( colour = 'black', size = font_size),
        axis.title = element_text( colour = 'black', size = font_size )
  )


summary_ratio_location<-aggregate(ratio ~ location_blood_drawing, tail_jugular , function(x) c(mean = mean(x), sd = sd(x)))
summary_ratio_location<-do.call(data.frame,summary_ratio_location)
plot2<-ggplot(tail_jugular, aes(x=location_blood_drawing, y=ratio)) + 
   geom_point(data = summary_ratio_location, mapping = aes(x =location_blood_drawing, y= ratio.mean),size=1, color="darkgray") +
  geom_errorbar(data=summary_ratio_location, mapping = aes( x=location_blood_drawing, y=ratio.mean, ymin = ratio.mean - ratio.sd, ymax = ratio.mean + ratio.sd), width = 0.1,color="darkgray")+
  geom_jitter(position = position_jitter(w=0.3, h=0), aes(shape=Heifer_ID, fill=location_blood_drawing), size=shape_size) +
  scale_fill_manual(values = c("#9964cb","#5fa369")) +
  scale_shape_manual(values=c(21:25))+
  scale_color_manual(values="black")+
  scale_x_discrete(name=NULL,breaks=c("jugular","tail"), labels=c("Jugular","Coccygeal")) +
  scale_y_continuous(name=expression(Ratio(A[260]/A[280])))+
  theme_classic() + 
  theme(legend.position = "none",
        panel.grid = element_blank(),
        panel.background = element_blank(),
        axis.text = element_text( colour = 'black', size = font_size),
        axis.title = element_text( colour = 'black', size = font_size )
  )

summary_A260_location<-aggregate(A260 ~ location_blood_drawing, tail_jugular , function(x) c(mean = mean(x), sd = sd(x)))
summary_A260_location<-do.call(data.frame,summary_A260_location)
plot3<-ggplot(tail_jugular, aes(x=location_blood_drawing, y=A260)) + 
  geom_point(data = summary_A260_location, mapping = aes(x =location_blood_drawing, y= A260.mean),size=1, color="darkgray") +
  geom_errorbar(data=summary_A260_location, mapping = aes( x=location_blood_drawing, y=A260.mean, ymin = A260.mean - A260.sd, ymax = A260.mean + A260.sd), width = 0.1,color="darkgray")+
   geom_jitter(position = position_jitter(w=0.3, h=0), aes(shape=Heifer_ID, fill=location_blood_drawing), size=shape_size) +
  scale_fill_manual(values = c("#9964cb","#5fa369")) +
  scale_shape_manual(values=c(21:25))+
  scale_color_manual(values="black")+
  scale_x_discrete(name=NULL,breaks=c("jugular","tail"), labels=c("Jugular","Coccygeal")) +
  scale_y_continuous(name="Absorbance at 260 nm")+
  theme_classic() + 
  theme(legend.position = "none",
        panel.grid = element_blank(),
        panel.background = element_blank(),
        axis.text = element_text( colour = 'black', size = font_size),
        axis.title = element_text( colour = 'black', size = font_size )
  )

summary_A280_location<-aggregate(A280 ~ location_blood_drawing, tail_jugular , function(x) c(mean = mean(x), sd = sd(x)))
summary_A280_location<-do.call(data.frame,summary_A280_location)
plot4<-ggplot(tail_jugular, aes(x=location_blood_drawing, y=A280)) + 
  geom_point(data = summary_A280_location, mapping = aes(x =location_blood_drawing, y= A280.mean),size=1, color="darkgray") +
  geom_errorbar(data=summary_A280_location, mapping = aes( x=location_blood_drawing, y=A280.mean, ymin = A280.mean - A280.sd, ymax = A280.mean + A280.sd), width = 0.1,color="darkgray")+
  geom_jitter(position = position_jitter(w=0.3, h=0), aes(shape=Heifer_ID, fill=location_blood_drawing), size=shape_size) +
  scale_fill_manual(values = c("#9964cb","#5fa369")) +
  scale_shape_manual(values=c(21:25))+
  scale_color_manual(values="black")+
  scale_x_discrete(name=NULL, breaks=c("jugular","tail"), labels=c("Jugular","Coccygeal")) +
  scale_y_continuous(name="Absorbance at 280 nm")+
  #ylim(7,10) +
  theme_classic() + 
  theme(legend.position = "none",
        panel.grid = element_blank(),
        panel.background = element_blank(),
        axis.text = element_text( colour = 'black', size = font_size),
        axis.title = element_text( colour = 'black', size = font_size )
  )

plot_grid( plot1,  plot3,plot4,plot2, nrow = 1, align = 'v')
```

##### Additional file 2

Ribonucleic acid quality and abundance parameters from samples collected from the jugular vein and processed after refrigeration (4°C) for multiple windows of time (x-axis). Animals are indicated by shapes across charts.

```
processing_time$Heifer_ID <- as.factor(processing_time$Heifer_ID)
processing_time$Time_pre_processing <- factor(processing_time$Time_pre_processing, levels=c('1h', '3h', '6h', '8h','24h'))

font_size<-10
shape_size<-2

summary_RIN<-aggregate(RIN ~ Time_pre_processing, processing_time , function(x) c(mean = mean(x), sd = sd(x)))
summary_RIN<-do.call(data.frame,summary_RIN)
timeplot1<-ggplot(processing_time, aes(x=Time_pre_processing, y=RIN)) + 
 geom_point(data = summary_RIN, mapping = aes(x =Time_pre_processing, y= RIN.mean),size=1, color="darkgray") +
  geom_errorbar(data=summary_RIN, mapping = aes( x=Time_pre_processing, y=RIN.mean, ymin = RIN.mean - RIN.sd, ymax = RIN.mean + RIN.sd), width = 0.1,color="darkgray")+
  geom_jitter(position = position_jitter(w=0.3, h=0), aes(shape=Heifer_ID, fill=Time_pre_processing), size=shape_size) +
  scale_fill_manual(values = c("#9964cb","#5fa369","#ca587a","#b38c3a","#798bc6")) +
  scale_shape_manual(values=c(21:25))+
  scale_color_manual(values="black")+
  scale_x_discrete(limits=c("1h", "3h", "6h", "8h", "24h"), labels=c("1hr", "3hr", "6hr", "8hr", "24hr")) + 
  ylim(7,10) + 
  xlab("Time Pre-Processing") +
  ggsignif::geom_signif(annotations = c("0.03"), y_position=9.7, xmin = c(1),xmax=c(5),tip_length = 0.1, textsize=2)+
  theme_classic() + 
  theme(legend.position = "none",
        panel.grid = element_blank(),
        panel.background = element_blank(),
        axis.text = element_text( colour = 'black', size = font_size),
        axis.title = element_text( colour = 'black', size = font_size ),
  )

summary_A260<-aggregate(A260 ~ Time_pre_processing, processing_time , function(x) c(mean = mean(x), sd = sd(x)))
summary_A260<-do.call(data.frame,summary_A260)
timeplot2<-ggplot(processing_time, aes(x=Time_pre_processing, y=A260)) + 
  geom_point(data = summary_A260, mapping = aes(x =Time_pre_processing, y= A260.mean),size=1, color="darkgray") +
  geom_errorbar(data=summary_A260, mapping = aes( x=Time_pre_processing, y=A260.mean, ymin = A260.mean - A260.sd, ymax = A260.mean + A260.sd), 
                width = 0.1,color="darkgray")+
  geom_jitter(position = position_jitter(w=0.3, h=0), aes(shape=Heifer_ID, fill=Time_pre_processing), size=shape_size) +
  scale_fill_manual(values = c("#9964cb","#5fa369","#ca587a","#b38c3a","#798bc6")) +
  scale_shape_manual(values=c(21:25))+
  scale_color_manual(values="black")+
   scale_x_discrete(limits=c("1h", "3h", "6h", "8h", "24h"), labels=c("1hr", "3hr", "6hr", "8hr", "24hr")) + 
  xlab("Time Pre-Processing") +
  theme_classic() + 
  theme(legend.position = "none",
        panel.grid = element_blank(),
        panel.background = element_blank(),
        axis.text = element_text( colour = 'black', size = font_size),
        axis.title = element_text( colour = 'black', size = font_size ),
  )

summary_A280<-aggregate(A280 ~ Time_pre_processing, processing_time , function(x) c(mean = mean(x), sd = sd(x)))
summary_A280<-do.call(data.frame,summary_A280)
timeplot3<-ggplot(processing_time, aes(x=Time_pre_processing, y=A280)) + 
  geom_errorbar(data=summary_A280, mapping = aes( x=Time_pre_processing, y=A280.mean, ymin = A280.mean - A280.sd, ymax = A280.mean + A280.sd), 
                width = 0.1,color="darkgray")+
  geom_jitter(position = position_jitter(w=0.3, h=0), aes(shape=Heifer_ID, fill=Time_pre_processing), size=shape_size) +
  scale_fill_manual(values = c("#9964cb","#5fa369","#ca587a","#b38c3a","#798bc6")) +
  scale_shape_manual(values=c(21:25))+
  scale_color_manual(values="black")+geom_point(data = summary_A280, mapping = aes(x =Time_pre_processing, y= A280.mean),size=1, color="darkgray") +
   scale_x_discrete(limits=c("1h", "3h", "6h", "8h", "24h"), labels=c("1hr", "3hr", "6hr", "8hr", "24hr")) +
  xlab("Time Pre-Processing") +
  theme_classic() + 
  theme(legend.position = "none",
        panel.grid = element_blank(),
        panel.background = element_blank(),
        axis.text = element_text( colour = 'black', size = font_size),
        axis.title = element_text( colour = 'black', size = font_size ),
  )

summary_ratio<-aggregate(ratio ~ Time_pre_processing, processing_time , function(x) c(mean = mean(x), sd = sd(x)))
summary_ratio<-do.call(data.frame,summary_ratio)
timeplot4<-ggplot(processing_time, aes(x=Time_pre_processing, y=ratio)) + 
  geom_point(data = summary_ratio, mapping = aes(x =Time_pre_processing, y= ratio.mean),size=1, color="darkgray") +
  geom_errorbar(data=summary_ratio, mapping = aes(x=Time_pre_processing, y=ratio.mean, ymin = ratio.mean - ratio.sd,ymax = ratio.mean + ratio.sd), 
                width = 0.1,color="darkgray")+
  scale_x_discrete(limits=c("1h", "3h", "6h", "8h", "24h"), labels=c("1hr", "3hr", "6hr", "8hr", "24hr")) +
  geom_jitter(position = position_jitter(w=0.3, h=0), aes(shape=Heifer_ID, fill=Time_pre_processing), size=shape_size) +
  scale_fill_manual(values = c("#9964cb","#5fa369","#ca587a","#b38c3a","#798bc6")) +
  scale_shape_manual(values=c(21:25))+
  scale_color_manual(values="black")+
  scale_y_continuous(name=expression(Ratio(A[260]/A[280])), limits = c(1.75,2))+
  xlab("Time Pre-Processing") +
  theme_classic() + 
  theme(legend.position = "none",
        panel.grid = element_blank(),
        panel.background = element_blank(),
        axis.text = element_text( colour = 'black', size = font_size),
        axis.title = element_text( colour = 'black', size = font_size ),
  )

plot_grid( timeplot1,timeplot2,timeplot3,timeplot4, nrow=2, align='v')
```

The next segment was not included in the paper, but it served for quality control of the RNA-sequencing data produced.

```
counting_summary_c<-reshape2::melt(counting_summary_b[,c(1,8,10:12)])
counting_summary_c$value<-counting_summary_c$value*100
counting_summary_c$variable<-factor(counting_summary_c$variable, levels=c( "perc_number_of_reads_discarted" ,"perc_unassigned_Ambiguity","perc_unassigned_NoFeatures" , "perc_number_of_reads_assigned"))
plot1<-ggplot() + 
  geom_bar(aes(y = value, x = sample, fill = variable), data = counting_summary_c,stat="identity")+
  scale_y_continuous(name="Percentage", breaks = seq(0,100,10))+
  scale_fill_hue(labels=c("Discarted", "Unassigned_Ambiguity","Unassigned_NoFeatures" , "Assigned"))+
  #ggtitle("Distribution of reads after alignment")+
  theme_bw(base_size = 12)+
  theme(legend.title = element_blank(),
        axis.title.x = element_blank(),
        axis.text.x = element_text(angle=75, hjust=1))

counting_summary_c<-reshape2::melt(counting_summary_b[,c(1,5)])
plot2<-ggplot() + 
  geom_bar(aes(y = value, x = sample, fill = variable), data = counting_summary_c,stat="identity")+
  scale_y_continuous(name="Read pairs sequenced")+
  theme_bw(base_size = 12)+
  theme(legend.title = element_blank(),
        axis.title.x = element_blank(),
        axis.text.x = element_blank())

counting_summary_c<-reshape2::melt(counting_summary_b[,c(1,2)])
plot3<-ggplot() + 
  geom_bar(aes(y = value, x = sample, fill = variable), data = counting_summary_c,stat="identity")+
  scale_y_continuous(name="Read pairs in annotation")+
  geom_hline(yintercept=10^7, linetype="dashed", color = "gray", size=1)+
  theme_bw(base_size = 12)+
  theme(legend.title = element_blank(),
        axis.title.x = element_blank(),
        axis.text.x = element_blank())

counting_summary_c<-reshape2::melt(counting_summary_b[,c(1,10)])
plot4<-ggplot() + 
  geom_bar(aes(y = value, x = sample, fill = variable), data = counting_summary_c,stat="identity")+
  scale_y_continuous(name="Proportion reads \n matching annotation")+
  theme_bw(base_size = 12)+
  theme(legend.title = element_blank(),
        axis.title.x = element_blank(),
        axis.text.x = element_blank())

plot_grid( plot2, plot3,plot4, plot1,  ncol = 1, align = 'v')
```

##### Test for sequencing bias

```
efficiency_data <- data.frame(Sample=counting_summary_b$sample,reads_sequenced=counting_summary_b$reads_sequenced,perc_number_of_reads_assigned= counting_summary_b$perc_number_of_reads_assigned)
processing_time_bias <- merge(RNA_data_frame, bias_metrics_b, by.x="Seq_name", by.y="sample")
processing_time_bias <- merge(processing_time_bias, efficiency_data, by.x="Seq_name", by.y="Sample")
processing_time_bias$Heifer_ID <- as.factor(processing_time_bias$Heifer_ID)
processing_time_bias$location_blood_drawing <- as.factor(processing_time_bias$location_blood_drawing)
processing_time_bias$Time_pre_processing <- as.factor(processing_time_bias$Time_pre_processing)

processing_time_bias_eff<-processing_time_bias[processing_time_bias$location_blood_drawing=="jugular",]


bias_fit <- glmmTMB(three_prime_bias_mean ~ Time_pre_processing + (1|Heifer_ID), data=processing_time_bias_eff, family = binomial(link="logit"))
summary(bias_fit)
```

```
##  Family: binomial  ( logit )
## Formula:          three_prime_bias_mean ~ Time_pre_processing + (1 | Heifer_ID)
## Data: processing_time_bias_eff
## 
##      AIC      BIC   logLik deviance df.resid 
##     46.6     54.0    -17.3     34.6       19 
## 
## Random effects:
## 
## Conditional model:
##  Groups    Name        Variance  Std.Dev. 
##  Heifer_ID (Intercept) 9.369e-10 3.061e-05
## Number of obs: 25, groups:  Heifer_ID, 5
## 
## Conditional model:
##                        Estimate Std. Error z value Pr(>|z|)
## (Intercept)             0.02603    0.89450   0.029    0.977
## Time_pre_processing24h  0.03569    1.26527   0.028    0.977
## Time_pre_processing3h  -0.01579    1.26497  -0.012    0.990
## Time_pre_processing6h   0.02775    1.26519   0.022    0.983
## Time_pre_processing8h   0.05637    1.26550   0.045    0.964
```

```
car::Anova(bias_fit, type="III")
```

```
## Analysis of Deviance Table (Type III Wald chisquare tests)
## 
## Response: three_prime_bias_mean
##                      Chisq Df Pr(>Chisq)
## (Intercept)         0.0008  1     0.9768
## Time_pre_processing 0.0041  4     1.0000
```

```
eff_fit <- glmmTMB(perc_number_of_reads_assigned ~ Time_pre_processing + (1|Heifer_ID), data=processing_time_bias_eff, family = binomial(link="logit"))
summary(eff_fit)
```

```
##  Family: binomial  ( logit )
## Formula:          
## perc_number_of_reads_assigned ~ Time_pre_processing + (1 | Heifer_ID)
## Data: processing_time_bias_eff
## 
##      AIC      BIC   logLik deviance df.resid 
##     46.2     53.5    -17.1     34.2       19 
## 
## Random effects:
## 
## Conditional model:
##  Groups    Name        Variance  Std.Dev. 
##  Heifer_ID (Intercept) 9.311e-10 3.051e-05
## Number of obs: 25, groups:  Heifer_ID, 5
## 
## Conditional model:
##                        Estimate Std. Error z value Pr(>|z|)
## (Intercept)             0.34111    0.90747   0.376    0.707
## Time_pre_processing24h -0.21252    1.27546  -0.167    0.868
## Time_pre_processing3h  -0.10993    1.27837  -0.086    0.931
## Time_pre_processing6h   0.03280    1.28522   0.025    0.980
## Time_pre_processing8h  -0.05785    1.28049  -0.045    0.964
```

```
car::Anova(eff_fit, type="III")
```

```
## Analysis of Deviance Table (Type III Wald chisquare tests)
## 
## Response: perc_number_of_reads_assigned
##                      Chisq Df Pr(>Chisq)
## (Intercept)         0.1413  1     0.7070
## Time_pre_processing 0.0461  4     0.9997
```

##### Summary of RNA-seq data

```
count_data_a<-count_data[rowSums(count_data)>0,]
count_data_annotated<-merge(count_data_a,annotation.ensembl.symbol, by.x="row.names", by.y="ensembl_gene_id", all.x=TRUE, all.y=FALSE)

n_reads_protein_coding<-sum(count_data_annotated[count_data_annotated$gene_biotype=='protein_coding',c(2:6)])
n_reads_lncRNA<-sum(count_data_annotated[count_data_annotated$gene_biotype=='lncRNA',c(2:6)])
n_reads_pseudogene<-sum(count_data_annotated[count_data_annotated$gene_biotype %in% c('pseudogene','processed_pseudogene'),c(2:6)])
n_reads_others<-sum(count_data_annotated[!(count_data_annotated$gene_biotype %in% c('pseudogene','processed_pseudogene','protein_coding','lncRNA')),c(2:6)])

summary_RNA_seq<-data.frame(class=c('protein_coding','lncRNA', 'pseudogene','others'), nreads=c(n_reads_protein_coding,n_reads_lncRNA,n_reads_pseudogene,n_reads_others))

summary_RNA_seq<-mutate(summary_RNA_seq,  prop = round(nreads/sum(nreads)*100,2))
summary_RNA_seq<-summary_RNA_seq[with(summary_RNA_seq, order(-nreads)),]
summary_RNA_seq$class<-factor(summary_RNA_seq$class, levels=c('protein_coding', 'lncRNA','pseudogene','others' ))
```

```
knitr::kable(summary_RNA_seq, format = 'pandoc')
```

| class | nreads | prop |
| --- | --- | --- |
| protein\_coding | 87462335 | 98.90 |
| lncRNA | 351164 | 0.40 |
| pseudogene | 328030 | 0.37 |
| others | 292673 | 0.33 |

We subset the counts to retain only gene biotypes ‘protein\_coding’, ‘lncRNA’, and ‘pseudogene’.

```
count_data_a<-count_data[rowSums(count_data)>0,]
count_data_annotated<-merge(count_data_a,annotation.ensembl.symbol, by.x="row.names", by.y="ensembl_gene_id", all.x=TRUE, all.y=FALSE)
count_data_annotated<-count_data_annotated[count_data_annotated$gene_biotype %in% c('protein_coding', 'lncRNA','pseudogene'),]
count_data_annotated_length<-gene.length[gene.length$ensembl_gene_id %in% count_data_annotated$Row.names, 2]
```

We calculate counts per million reads (CPM) and fragments per million per kilobase (FPKM) and subset the lowly expressed genes.

```
data_fpkm<-rpkm(count_data_annotated[,c(2:31)], count_data_annotated_length)
rownames(data_fpkm)<-count_data_annotated$Row.names
data_cpm<-edgeR::cpm(count_data_annotated[,c(2:31)])
rownames(data_cpm)<-count_data_annotated$Row.names

keep<-rowSums(data_fpkm>1) >=5
data_fpkm_filtered<-data_fpkm[keep,]
keep<-rowSums(data_cpm>1) >=5
data_cpm_filtered<-data_cpm[keep,]
genes_expressed<-intersect(rownames(data_fpkm_filtered), rownames(data_cpm_filtered))
data_fpkm_filtered<-data_fpkm_filtered[rownames(data_fpkm_filtered) %in% genes_expressed,]
data_cpm_filtered<-data_cpm_filtered[rownames(data_cpm_filtered) %in% genes_expressed,]
```

Number of genes retained for downstream analysis based on biotype.

```
table(annotation.ensembl.symbol[annotation.ensembl.symbol$ensembl_gene_id %in% genes_expressed, ]$gene_biotype)
```

```
## 
##         lncRNA protein_coding     pseudogene 
##            287          12414            109
```

##### Compare genes detected in Coccygeal vs Jugular Vein

```
tail_jugular<-merge(tail_jugular, data.frame(genes_detected_bef_filtering=colSums(data_cpm>1),genes_detected_post_filtering=colSums(data_cpm_filtered>1)), by.x="Seq_name", by.y="row.names")

wilcox.test( genes_detected_bef_filtering~location_blood_drawing, data= tail_jugular, paired = TRUE, alternative = "two.sided", exact = FALSE)
```

```
## 
##  Wilcoxon signed rank test with continuity correction
## 
## data:  genes_detected_bef_filtering by location_blood_drawing
## V = 2, p-value = 0.1775
## alternative hypothesis: true location shift is not equal to 0
```

```
t.test(genes_detected_bef_filtering~location_blood_drawing, data= tail_jugular, paired = TRUE, alternate="two.sided")
```

```
## 
##  Paired t-test
## 
## data:  genes_detected_bef_filtering by location_blood_drawing
## t = -1.5475, df = 4, p-value = 0.1966
## alternative hypothesis: true difference in means is not equal to 0
## 95 percent confidence interval:
##  -224.08684   63.68684
## sample estimates:
## mean of the differences 
##                   -80.2
```

```
aggregate(genes_detected_bef_filtering~location_blood_drawing, tail_jugular , function(x) c(mean = round(mean(x),digits = 0), sd = round(sd(x), digits = 0)))
```

```
##   location_blood_drawing genes_detected_bef_filtering.mean
## 1                jugular                             13055
## 2                   tail                             13135
##   genes_detected_bef_filtering.sd
## 1                             165
## 2                             167
```

```
wilcox.test( genes_detected_post_filtering~location_blood_drawing, data= tail_jugular, paired = TRUE, alternative = "two.sided", exact = FALSE)
```

```
## 
##  Wilcoxon signed rank test with continuity correction
## 
## data:  genes_detected_post_filtering by location_blood_drawing
## V = 2, p-value = 0.1775
## alternative hypothesis: true location shift is not equal to 0
```

```
t.test(genes_detected_post_filtering~location_blood_drawing, data= tail_jugular, paired = TRUE, alternate="two.sided")
```

```
## 
##  Paired t-test
## 
## data:  genes_detected_post_filtering by location_blood_drawing
## t = -1.4135, df = 4, p-value = 0.2304
## alternative hypothesis: true difference in means is not equal to 0
## 95 percent confidence interval:
##  -81.81253  26.61253
## sample estimates:
## mean of the differences 
##                   -27.6
```

```
aggregate(genes_detected_post_filtering~location_blood_drawing, tail_jugular , function(x) c(mean = round(mean(x),digits = 0), sd = round(sd(x), digits = 0)))
```

```
##   location_blood_drawing genes_detected_post_filtering.mean
## 1                jugular                              12462
## 2                   tail                              12490
##   genes_detected_post_filtering.sd
## 1                               74
## 2                               51
```

##### Compare genes detected between Processing Times

```
processing_time_bias<-merge(processing_time_bias, data.frame(genes_detected_bef_filtering=colSums(data_cpm>=1),genes_detected_post_filtering=colSums(data_cpm_filtered>=1)), by.x="Seq_name", by.y="row.names")

processing_time_bias_eff<-processing_time_bias[processing_time_bias$location_blood_drawing=="jugular",]
random_genes_bef_filt_lme<-nlme::lme(genes_detected_bef_filtering ~ Time_pre_processing,random=~ 1 |Heifer_ID,data=processing_time_bias_eff,method="REML")
summary(random_genes_bef_filt_lme)
```

```
## Linear mixed-effects model fit by REML
##   Data: processing_time_bias_eff 
##        AIC      BIC    logLik
##   254.0961 261.0663 -120.0481
## 
## Random effects:
##  Formula: ~1 | Heifer_ID
##         (Intercept) Residual
## StdDev:    121.7336 58.58959
## 
## Fixed effects:  genes_detected_bef_filtering ~ Time_pre_processing 
##                          Value Std.Error DF   t-value p-value
## (Intercept)            13054.8  60.41824 16 216.07381  0.0000
## Time_pre_processing24h    57.4  37.05531 16   1.54904  0.1409
## Time_pre_processing3h     63.0  37.05531 16   1.70016  0.1085
## Time_pre_processing6h     74.8  37.05531 16   2.01860  0.0606
## Time_pre_processing8h     47.0  37.05531 16   1.26837  0.2228
##  Correlation: 
##                        (Intr) Tm__24 Tm_p_3 Tm_p_6
## Time_pre_processing24h -0.307                     
## Time_pre_processing3h  -0.307  0.500              
## Time_pre_processing6h  -0.307  0.500  0.500       
## Time_pre_processing8h  -0.307  0.500  0.500  0.500
## 
## Standardized Within-Group Residuals:
##         Min          Q1         Med          Q3         Max 
## -1.88147395 -0.54828009  0.06741822  0.42925725  1.95197157 
## 
## Number of Observations: 25
## Number of Groups: 5
```

```
anova(random_genes_bef_filt_lme)
```

```
##                     numDF denDF  F-value p-value
## (Intercept)             1    16 55365.35  <.0001
## Time_pre_processing     4    16     1.21  0.3432
```

```
dunnett_random_genes_bef_filt_lme<-(glht(random_genes_bef_filt_lme, mcp(Time_pre_processing="Dunnett")))
summary( dunnett_random_genes_bef_filt_lme)
```

```
## 
##   Simultaneous Tests for General Linear Hypotheses
## 
## Multiple Comparisons of Means: Dunnett Contrasts
## 
## 
## Fit: lme.formula(fixed = genes_detected_bef_filtering ~ Time_pre_processing, 
##     data = processing_time_bias_eff, random = ~1 | Heifer_ID, 
##     method = "REML")
## 
## Linear Hypotheses:
##               Estimate Std. Error z value Pr(>|z|)
## 24h - 1h == 0    57.40      37.06   1.549    0.339
## 3h - 1h == 0     63.00      37.06   1.700    0.260
## 6h - 1h == 0     74.80      37.06   2.019    0.138
## 8h - 1h == 0     47.00      37.06   1.268    0.519
## (Adjusted p values reported -- single-step method)
```

```
random_genes_post_filt_lme<-nlme::lme(genes_detected_post_filtering ~ Time_pre_processing,random=~ 1 |Heifer_ID,data=processing_time_bias_eff,method="REML")
summary(random_genes_post_filt_lme)
```

```
## Linear mixed-effects model fit by REML
##   Data: processing_time_bias_eff 
##        AIC      BIC    logLik
##   226.2583 233.2284 -106.1291
## 
## Random effects:
##  Formula: ~1 | Heifer_ID
##         (Intercept) Residual
## StdDev:    46.45821 31.07467
## 
## Fixed effects:  genes_detected_post_filtering ~ Time_pre_processing 
##                          Value Std.Error DF  t-value p-value
## (Intercept)            12462.0  24.99600 16 498.5598  0.0000
## Time_pre_processing24h    39.8  19.65335 16   2.0251  0.0599
## Time_pre_processing3h     36.8  19.65335 16   1.8725  0.0795
## Time_pre_processing6h     35.4  19.65335 16   1.8012  0.0905
## Time_pre_processing8h     16.4  19.65335 16   0.8345  0.4163
##  Correlation: 
##                        (Intr) Tm__24 Tm_p_3 Tm_p_6
## Time_pre_processing24h -0.393                     
## Time_pre_processing3h  -0.393  0.500              
## Time_pre_processing6h  -0.393  0.500  0.500       
## Time_pre_processing8h  -0.393  0.500  0.500  0.500
## 
## Standardized Within-Group Residuals:
##         Min          Q1         Med          Q3         Max 
## -2.06962897 -0.57391186  0.08327253  0.52412359  1.66975127 
## 
## Number of Observations: 25
## Number of Groups: 5
```

```
anova(random_genes_post_filt_lme)
```

```
##                     numDF denDF  F-value p-value
## (Intercept)             1    16 331581.3  <.0001
## Time_pre_processing     4    16      1.5  0.2475
```

```
dunnett_random_genes_post_filt_lme<-(glht(random_genes_post_filt_lme, mcp(Time_pre_processing="Dunnett")))
summary( dunnett_random_genes_post_filt_lme)
```

```
## 
##   Simultaneous Tests for General Linear Hypotheses
## 
## Multiple Comparisons of Means: Dunnett Contrasts
## 
## 
## Fit: lme.formula(fixed = genes_detected_post_filtering ~ Time_pre_processing, 
##     data = processing_time_bias_eff, random = ~1 | Heifer_ID, 
##     method = "REML")
## 
## Linear Hypotheses:
##               Estimate Std. Error z value Pr(>|z|)
## 24h - 1h == 0    39.80      19.65   2.025    0.135
## 3h - 1h == 0     36.80      19.65   1.872    0.187
## 6h - 1h == 0     35.40      19.65   1.801    0.215
## 8h - 1h == 0     16.40      19.65   0.834    0.820
## (Adjusted p values reported -- single-step method)
```

##### Table 2

```
processing_time_bias_data_table<-as.data.table(processing_time_bias)

processing_time_bias_summary<- processing_time_bias_data_table[,. (ave_reads_sequenced=round(mean(reads_sequenced),0) ,
                                                                           sd_reads_sequenced=round(sd(reads_sequenced),0),
                                                                           ave_genes_detected_bef_filtering=round(mean(genes_detected_bef_filtering),0) ,
                                                                           sd_genes_detected_bef_filtering=round(sd(genes_detected_bef_filtering),0),
                                                                           ave_genes_detected_post_filtering=round(mean(genes_detected_post_filtering),0) ,
                                                                           sd_genes_detected_post_filtering=round(sd(genes_detected_post_filtering),0),
                                                                           ave_perc_number_of_reads_assigned=round(mean(perc_number_of_reads_assigned),2) ,
                                                                           sd_perc_number_of_reads_assigned=round(sd(perc_number_of_reads_assigned),2),
                                                                           ave_three_prime_bias_mean=round(mean(three_prime_bias_mean),2) ,
                                                                           sd_three_prime_bias_mean=round(sd(three_prime_bias_mean),2)
), by = c('location_blood_drawing','Time_pre_processing')]


processing_time_bias_summary[location_blood_drawing == "tail", location_blood_drawing := "coccygeal"  ]

#kbl(processing_time_bias_summary,
#    caption = "Table 2. Metrics obtained for RNA-sequencing libraries produced from peripheral white blood cells.",
#    col.names=c("Vein\n","Processing time","$\\overline{x}$","$\\hat{\\sigma}$","$\\bar{x}$","$\\hat{\\sigma}$","$\\bar{x}$","$\\hat{\\sigma}$","$\\bar{x}$","$\\hat{\\sigma}$","$\\bar{x}$","$\\hat{\\sigma}$")) %>%
#  kable_classic() %>%
#  add_header_above(c(" "=2, "Reads produced\n"=2, "Genes detected before filtering"=2, "Genes detected \n after filtering"=2,"Proportion of reads matching annotation"=2,"3'/5' bias"=2)) %>%
  #pack_rows(group_label="Coccygeal", 1, 1) %>%
  #pack_rows(group_label="Jugular", 2, 6) %>%
#  collapse_rows(columns = 1, valign = "top") %>%
#  column_spec(column = 3:11, width = "1.5in")


kbl(processing_time_bias_summary[,-1],
    caption = "Table 2. Metrics for RNA-sequencing data produced from peripheral white blood cells.",
    col.names=c("Processing time","$\\overline{x}$","$\\hat{\\sigma}$","$\\bar{x}$","$\\hat{\\sigma}$","$\\bar{x}$","$\\hat{\\sigma}$","$\\bar{x}$","$\\hat{\\sigma}$","$\\bar{x}$","$\\hat{\\sigma}$")) %>%
  kable_classic() %>%
  add_header_above(c(" "=1, "Reads produced\n"=2, "Genes detected before filtering"=2, "Genes detected \n after filtering"=2,"Proportion of reads matching annotation"=2,"3'/5' bias"=2)) %>%
  pack_rows(group_label="Coccygeal", 1, 1) %>%
  pack_rows(group_label="Jugular", 2, 6) %>%
  column_spec(column = 2:11, width = "1.5in")
```

Table 2. Metrics for RNA-sequencing data produced from peripheral white blood cells.

|  | Reads produced | | Genes detected before filtering | | Genes detected   after filtering | | Proportion of reads matching annotation | | 3’/5’ bias | |
| --- | --- | --- | --- | --- | --- | --- | --- | --- | --- | --- |
| Processing time | \(\overline{x}\) | \(\hat{\sigma}\) | \(\bar{x}\) | \(\hat{\sigma}\) | \(\bar{x}\) | \(\hat{\sigma}\) | \(\bar{x}\) | \(\hat{\sigma}\) | \(\bar{x}\) | \(\hat{\sigma}\) |
| **Coccygeal** | | | | | | | | | | |
| 1h | 30214519 | 4270887 | 13135 | 167 | 12490 | 51 | 0.56 | 0.04 | 0.51 | 0.02 |
| **Jugular** | | | | | | | | | | |
| 1h | 28452995 | 5098116 | 13055 | 165 | 12462 | 74 | 0.58 | 0.03 | 0.51 | 0.03 |
| 3h | 32132121 | 1968340 | 13118 | 128 | 12499 | 49 | 0.56 | 0.03 | 0.50 | 0.01 |
| 6h | 28263832 | 2338260 | 13130 | 139 | 12497 | 57 | 0.59 | 0.02 | 0.51 | 0.03 |
| 8h | 30483625 | 3040358 | 13102 | 132 | 12478 | 54 | 0.57 | 0.04 | 0.52 | 0.02 |
| 24h | 29683203 | 2546563 | 13112 | 105 | 12502 | 40 | 0.53 | 0.08 | 0.52 | 0.01 |

##### Evaluate genes detected after filtering - Processing Time

```
processing_time_bias_eff<-processing_time_bias[processing_time_bias$location_blood_drawing=="jugular",]
random_genes_bef_filt_lme<-nlme::lme(genes_detected_bef_filtering ~ Time_pre_processing,random=~ 1 |Heifer_ID,data=processing_time_bias_eff,method="REML")
summary(random_genes_bef_filt_lme)
```

```
## Linear mixed-effects model fit by REML
##   Data: processing_time_bias_eff 
##        AIC      BIC    logLik
##   254.0961 261.0663 -120.0481
## 
## Random effects:
##  Formula: ~1 | Heifer_ID
##         (Intercept) Residual
## StdDev:    121.7336 58.58959
## 
## Fixed effects:  genes_detected_bef_filtering ~ Time_pre_processing 
##                          Value Std.Error DF   t-value p-value
## (Intercept)            13054.8  60.41824 16 216.07381  0.0000
## Time_pre_processing24h    57.4  37.05531 16   1.54904  0.1409
## Time_pre_processing3h     63.0  37.05531 16   1.70016  0.1085
## Time_pre_processing6h     74.8  37.05531 16   2.01860  0.0606
## Time_pre_processing8h     47.0  37.05531 16   1.26837  0.2228
##  Correlation: 
##                        (Intr) Tm__24 Tm_p_3 Tm_p_6
## Time_pre_processing24h -0.307                     
## Time_pre_processing3h  -0.307  0.500              
## Time_pre_processing6h  -0.307  0.500  0.500       
## Time_pre_processing8h  -0.307  0.500  0.500  0.500
## 
## Standardized Within-Group Residuals:
##         Min          Q1         Med          Q3         Max 
## -1.88147395 -0.54828009  0.06741822  0.42925725  1.95197157 
## 
## Number of Observations: 25
## Number of Groups: 5
```

```
anova(random_genes_bef_filt_lme)
```

```
##                     numDF denDF  F-value p-value
## (Intercept)             1    16 55365.35  <.0001
## Time_pre_processing     4    16     1.21  0.3432
```

```
dunnett_random_genes_bef_filt_lme<-(glht(random_genes_bef_filt_lme, mcp(Time_pre_processing="Dunnett")))
summary( dunnett_random_genes_bef_filt_lme)
```

```
## 
##   Simultaneous Tests for General Linear Hypotheses
## 
## Multiple Comparisons of Means: Dunnett Contrasts
## 
## 
## Fit: lme.formula(fixed = genes_detected_bef_filtering ~ Time_pre_processing, 
##     data = processing_time_bias_eff, random = ~1 | Heifer_ID, 
##     method = "REML")
## 
## Linear Hypotheses:
##               Estimate Std. Error z value Pr(>|z|)
## 24h - 1h == 0    57.40      37.06   1.549    0.339
## 3h - 1h == 0     63.00      37.06   1.700    0.260
## 6h - 1h == 0     74.80      37.06   2.019    0.138
## 8h - 1h == 0     47.00      37.06   1.268    0.519
## (Adjusted p values reported -- single-step method)
```

```
random_genes_post_filt_lme<-nlme::lme(genes_detected_post_filtering ~ Time_pre_processing,random=~ 1 |Heifer_ID,data=processing_time_bias_eff,method="REML")
summary(random_genes_post_filt_lme)
```

```
## Linear mixed-effects model fit by REML
##   Data: processing_time_bias_eff 
##        AIC      BIC    logLik
##   226.2583 233.2284 -106.1291
## 
## Random effects:
##  Formula: ~1 | Heifer_ID
##         (Intercept) Residual
## StdDev:    46.45821 31.07467
## 
## Fixed effects:  genes_detected_post_filtering ~ Time_pre_processing 
##                          Value Std.Error DF  t-value p-value
## (Intercept)            12462.0  24.99600 16 498.5598  0.0000
## Time_pre_processing24h    39.8  19.65335 16   2.0251  0.0599
## Time_pre_processing3h     36.8  19.65335 16   1.8725  0.0795
## Time_pre_processing6h     35.4  19.65335 16   1.8012  0.0905
## Time_pre_processing8h     16.4  19.65335 16   0.8345  0.4163
##  Correlation: 
##                        (Intr) Tm__24 Tm_p_3 Tm_p_6
## Time_pre_processing24h -0.393                     
## Time_pre_processing3h  -0.393  0.500              
## Time_pre_processing6h  -0.393  0.500  0.500       
## Time_pre_processing8h  -0.393  0.500  0.500  0.500
## 
## Standardized Within-Group Residuals:
##         Min          Q1         Med          Q3         Max 
## -2.06962897 -0.57391186  0.08327253  0.52412359  1.66975127 
## 
## Number of Observations: 25
## Number of Groups: 5
```

```
anova(random_genes_post_filt_lme)
```

```
##                     numDF denDF  F-value p-value
## (Intercept)             1    16 331581.3  <.0001
## Time_pre_processing     4    16      1.5  0.2475
```

```
dunnett_random_genes_post_filt_lme<-(glht(random_genes_post_filt_lme, mcp(Time_pre_processing="Dunnett")))
summary( dunnett_random_genes_post_filt_lme)
```

```
## 
##   Simultaneous Tests for General Linear Hypotheses
## 
## Multiple Comparisons of Means: Dunnett Contrasts
## 
## 
## Fit: lme.formula(fixed = genes_detected_post_filtering ~ Time_pre_processing, 
##     data = processing_time_bias_eff, random = ~1 | Heifer_ID, 
##     method = "REML")
## 
## Linear Hypotheses:
##               Estimate Std. Error z value Pr(>|z|)
## 24h - 1h == 0    39.80      19.65   2.025    0.135
## 3h - 1h == 0     36.80      19.65   1.872    0.187
## 6h - 1h == 0     35.40      19.65   1.801    0.215
## 8h - 1h == 0     16.40      19.65   0.834    0.820
## (Adjusted p values reported -- single-step method)
```

##### Additional file 3

Parameters of library quality from samples collected from the jugular and coccygeal veins and processed within one hour of sampling. Animals are indicated by shapes across charts.

```
tail_jugular<-processing_time_bias[processing_time_bias$Time_pre_processing=="1h",]

font_size<-10
shape_size<-2

summary_three_prime_bias_mean_location<-aggregate(three_prime_bias_mean ~ location_blood_drawing, tail_jugular , function(x) c(mean = round(mean(x),2), sd = round(sd(x),2) ))
summary_three_prime_bias_mean_location<-do.call(data.frame,summary_three_prime_bias_mean_location)
plot1<-ggplot(tail_jugular, aes(x=location_blood_drawing, y=three_prime_bias_mean)) + 
  geom_point(data = summary_three_prime_bias_mean_location, mapping = aes(x =location_blood_drawing, y= three_prime_bias_mean.mean),size=2, color="darkgray") +
  geom_errorbar(data=summary_three_prime_bias_mean_location, mapping = aes( x=location_blood_drawing, y=three_prime_bias_mean.mean, ymin = three_prime_bias_mean.mean - three_prime_bias_mean.sd, ymax = three_prime_bias_mean.mean + three_prime_bias_mean.sd), width = 0.1,color="darkgray")+
  geom_jitter(position = position_jitter(w=0.2, h=0), aes(shape=Heifer_ID, fill=location_blood_drawing), size=shape_size) +
  scale_fill_manual(values = c("#9964cb","#5fa369")) +
  scale_shape_manual(values=c(21:25))+
  scale_color_manual(values="black")+
  
  scale_y_continuous(name="3'bias")+
  scale_x_discrete(name=NULL,breaks=c("jugular","tail"), labels=c("Jugular","Coccygeal")) +
  #ylim(7,10) +
  theme_classic() + 
  theme(legend.position = "none",
        panel.grid = element_blank(),
        panel.background = element_blank(),
        axis.text = element_text( colour = 'black', size = font_size),
        axis.title = element_text( colour = 'black', size = font_size )
  )

summary_perc_number_of_reads_assigned_location<-aggregate(perc_number_of_reads_assigned ~ location_blood_drawing, tail_jugular , function(x) c(mean = round(mean(x),2), sd = round(sd(x),2) ))
summary_perc_number_of_reads_assigned_location<-do.call(data.frame,summary_perc_number_of_reads_assigned_location)
plot2<-ggplot(tail_jugular, aes(x=location_blood_drawing, y=perc_number_of_reads_assigned)) + 
   geom_point(data = summary_perc_number_of_reads_assigned_location, mapping = aes(x =location_blood_drawing, y= perc_number_of_reads_assigned.mean),size=2, color="darkgray") +
  geom_errorbar(data=summary_perc_number_of_reads_assigned_location, mapping = aes( x=location_blood_drawing, y=perc_number_of_reads_assigned.mean, ymin = perc_number_of_reads_assigned.mean - perc_number_of_reads_assigned.sd, ymax = perc_number_of_reads_assigned.mean + perc_number_of_reads_assigned.sd), width = 0.1,color="darkgray")+
  geom_jitter(position = position_jitter(w=0.2, h=0), aes(shape=Heifer_ID, fill=location_blood_drawing), size=shape_size) +
  scale_fill_manual(values = c("#9964cb","#5fa369")) +
  scale_shape_manual(values=c(21:25))+
  scale_color_manual(values="black")+
  scale_y_continuous(name="Library efficiency")+
  scale_x_discrete(name=NULL,breaks=c("jugular","tail"), labels=c("Jugular","Coccygeal")) +
  #ylim(7,10) +
  theme_classic() + 
  theme(legend.position = "none",
        panel.grid = element_blank(),
        panel.background = element_blank(),
        axis.text = element_text( colour = 'black', size = font_size),
        axis.title = element_text( colour = 'black', size = font_size )
  )

summary_genes_detected_bef_filtering_location<-aggregate(genes_detected_bef_filtering ~ location_blood_drawing, tail_jugular , function(x) c(mean = round(mean(x),0), sd = round(sd(x),0) ))
summary_genes_detected_bef_filtering_location<-do.call(data.frame,summary_genes_detected_bef_filtering_location)
plot3<-ggplot(tail_jugular, aes(x=location_blood_drawing, y=genes_detected_bef_filtering)) + 
  geom_point(data = summary_genes_detected_bef_filtering_location, mapping = aes(x =location_blood_drawing, y= genes_detected_bef_filtering.mean),size=2, color="darkgray") +
  geom_errorbar(data=summary_genes_detected_bef_filtering_location, mapping = aes( x=location_blood_drawing, y=genes_detected_bef_filtering.mean, ymin = genes_detected_bef_filtering.mean - genes_detected_bef_filtering.sd, ymax = genes_detected_bef_filtering.mean + genes_detected_bef_filtering.sd), width = 0.1,color="darkgray")+
  geom_jitter(position = position_jitter(w=0.2, h=0), aes(shape=Heifer_ID, fill=location_blood_drawing), size=shape_size) +
  scale_fill_manual(values = c("#9964cb","#5fa369")) +
  scale_shape_manual(values=c(21:25))+
  scale_color_manual(values="black")+
  
  scale_y_continuous(name="Number of genes detected \n before filtering")+
  scale_x_discrete(name=NULL,breaks=c("jugular","tail"), labels=c("Jugular","Coccygeal")) +
  #ylim(7,10) +
  theme_classic() + 
  theme(legend.position = "none",
        panel.grid = element_blank(),
        panel.background = element_blank(),
        axis.text = element_text( colour = 'black', size = font_size),
        axis.title = element_text( colour = 'black', size = font_size )
  )

summary_genes_detected_post_filtering_location<-aggregate(genes_detected_post_filtering ~ location_blood_drawing, tail_jugular , function(x) c(mean = round(mean(x),0), sd = round(sd(x),0) ))
summary_genes_detected_post_filtering_location<-do.call(data.frame,summary_genes_detected_post_filtering_location)
plot4<-ggplot(tail_jugular, aes(x=location_blood_drawing, y=genes_detected_post_filtering)) + 
   geom_point(data = summary_genes_detected_post_filtering_location, mapping = aes(x =location_blood_drawing, y= genes_detected_post_filtering.mean),size=2, color="darkgray") +
  geom_errorbar(data=summary_genes_detected_post_filtering_location, mapping = aes( x=location_blood_drawing, y=genes_detected_post_filtering.mean, ymin = genes_detected_post_filtering.mean - genes_detected_post_filtering.sd, ymax = genes_detected_post_filtering.mean + genes_detected_post_filtering.sd), width = 0.1,color="darkgray")+
  geom_jitter(position = position_jitter(w=0.2, h=0), aes(shape=Heifer_ID, fill=location_blood_drawing), size=shape_size) +
  scale_fill_manual(values = c("#9964cb","#5fa369")) +
  scale_shape_manual(values=c(21:25))+
  scale_color_manual(values="black")+
  scale_y_continuous(name="Number of genes detected \n after filtering")+
  scale_x_discrete(name=NULL,breaks=c("jugular","tail"), labels=c("Jugular","Coccygeal")) +
  #ylim(7,10) +
  theme_classic() + 
  theme(legend.position = "none",
        panel.grid = element_blank(),
        panel.background = element_blank(),
        axis.text = element_text( colour = 'black', size = font_size),
        axis.title = element_text( colour = 'black', size = font_size )
  )

plot_grid( plot1,plot2, plot3, plot4,nrow=1, align='h')
```

##### Additional file 4

Parameters of library quality from samples collected from the jugular and processed after refrigeration (4°C) for multiple windows of time (x-axis). Animals are indicated by shapes across charts.

```
font_size<-10
shape_size<-2

summary_bias<-aggregate(three_prime_bias_mean ~ Time_pre_processing, processing_time_bias_eff , function(x) c(mean = mean(x), sd = sd(x)))
summary_bias<-do.call(data.frame,summary_bias)
timeplot1<-ggplot(processing_time_bias_eff, aes(x=Time_pre_processing, y=three_prime_bias_mean)) + 
   geom_point(data = summary_bias, mapping = aes(x =Time_pre_processing, y= three_prime_bias_mean.mean), size=1, color="darkgray") +
  geom_errorbar(data=summary_bias, mapping = aes( x=Time_pre_processing, y=three_prime_bias_mean.mean, ymin = three_prime_bias_mean.mean - three_prime_bias_mean.sd, ymax = three_prime_bias_mean.mean + three_prime_bias_mean.sd), width = 0.1,color="darkgray")+
  geom_jitter(position = position_jitter(w=0.2, h=0), aes(shape=Heifer_ID, fill=Time_pre_processing), size=shape_size) +
  scale_fill_manual(values = c("#9964cb","#5fa369","#ca587a","#b38c3a","#798bc6")) +
  scale_shape_manual(values=c(21:25))+
  scale_color_manual(values="black")+
  scale_x_discrete(limits=c("1h", "3h", "6h", "8h", "24h"),labels=c("1hr", "3hr", "6hr", "8hr", "24hr")) + 
  scale_y_continuous(name="3' bias")+
  xlab("Time Pre-Processing") +
  theme_classic() + 
  theme(legend.position = "none",
        panel.grid = element_blank(),
        panel.background = element_blank(),
        axis.text = element_text( colour = 'black', size = font_size),
        axis.title = element_text( colour = 'black', size = font_size ))

summary_reads_assigned<-aggregate( perc_number_of_reads_assigned ~ Time_pre_processing, processing_time_bias_eff , function(x) c(mean = mean(x), sd = sd(x)))
summary_reads_assigned<-do.call(data.frame,summary_reads_assigned)
timeplot2<-ggplot(processing_time_bias_eff, aes(x=Time_pre_processing, y=perc_number_of_reads_assigned)) + 
  geom_point(data = summary_reads_assigned, mapping = aes(x =Time_pre_processing, y= perc_number_of_reads_assigned.mean), size=1, color="darkgray") +
  geom_errorbar(data=summary_reads_assigned, mapping = aes( x=Time_pre_processing, y=perc_number_of_reads_assigned.mean, ymin = perc_number_of_reads_assigned.mean - perc_number_of_reads_assigned.sd, ymax = perc_number_of_reads_assigned.mean + perc_number_of_reads_assigned.sd), width = 0.1,color="darkgray")+
  geom_jitter(position = position_jitter(w=0.2, h=0), aes(shape=Heifer_ID, fill=Time_pre_processing), size=shape_size) +
  scale_fill_manual(values = c("#9964cb","#5fa369","#ca587a","#b38c3a","#798bc6")) +
  scale_shape_manual(values=c(21:25))+
  scale_color_manual(values="black")+
  scale_x_discrete(limits=c("1h", "3h", "6h", "8h", "24h"),labels=c("1hr", "3hr", "6hr", "8hr", "24hr")) + 
  scale_y_continuous(name="Library efficiency")+
  xlab("Time Pre-Processing") +
  theme_classic() + 
  theme(legend.position = "none",
        panel.grid = element_blank(),
        panel.background = element_blank(),
        axis.text = element_text( colour = 'black', size = font_size),
        axis.title = element_text( colour = 'black', size = font_size )  )


summary_genes_detected_bef_filtering<-aggregate(genes_detected_bef_filtering ~ Time_pre_processing, processing_time_bias_eff , function(x) c(mean = round(mean(x),0), sd = round(sd(x),0)))
summary_genes_detected_bef_filtering<-do.call(data.frame,summary_genes_detected_bef_filtering)
timeplot3<-ggplot(processing_time_bias_eff, aes(x=Time_pre_processing, y=genes_detected_bef_filtering)) + 
  geom_point(data = summary_genes_detected_bef_filtering, mapping = aes(x =Time_pre_processing, y= genes_detected_bef_filtering.mean), size=1, color="darkgray") +
  geom_errorbar(data=summary_genes_detected_bef_filtering, mapping = aes( x=Time_pre_processing, y=genes_detected_bef_filtering.mean, ymin = genes_detected_bef_filtering.mean - genes_detected_bef_filtering.sd, ymax = genes_detected_bef_filtering.mean + genes_detected_bef_filtering.sd), width = 0.1,color="darkgray")+
  geom_jitter(position = position_jitter(w=0.2, h=0), aes(shape=Heifer_ID, fill=Time_pre_processing), size=shape_size) +
  scale_fill_manual(values = c("#9964cb","#5fa369","#ca587a","#b38c3a","#798bc6")) +
  scale_shape_manual(values=c(21:25))+
  scale_color_manual(values="black")+
  scale_x_discrete(limits=c("1h", "3h", "6h", "8h", "24h"), labels=c("1hr", "3hr", "6hr", "8hr", "24hr")) + 
  scale_y_continuous(name="Number of genes detected \n before filtering")+
  xlab("Time Pre-Processing") +
  theme_classic() + 
  theme(legend.position = "none",
        panel.grid = element_blank(),
        panel.background = element_blank(),
        axis.text = element_text( colour = 'black', size = font_size),
        axis.title = element_text( colour = 'black', size = font_size ))


summary_genes_detected_post_filtering<-aggregate(genes_detected_post_filtering ~ Time_pre_processing, processing_time_bias_eff , function(x) c(mean = round(mean(x),0), sd = round(sd(x),0)))
summary_genes_detected_post_filtering<-do.call(data.frame,summary_genes_detected_post_filtering)
timeplot4<-ggplot(processing_time_bias_eff, aes(x=Time_pre_processing, y=genes_detected_post_filtering)) + 
    geom_point(data = summary_genes_detected_post_filtering, mapping = aes(x =Time_pre_processing, y= genes_detected_post_filtering.mean), size=1, color="darkgray") +
  geom_errorbar(data=summary_genes_detected_post_filtering, mapping = aes( x=Time_pre_processing, y=genes_detected_post_filtering.mean, ymin = genes_detected_post_filtering.mean - genes_detected_post_filtering.sd, ymax = genes_detected_post_filtering.mean + genes_detected_post_filtering.sd), width = 0.1,color="darkgray")+
  geom_jitter(position = position_jitter(w=0.2, h=0), aes(shape=Heifer_ID, fill=Time_pre_processing), size=shape_size) +
  scale_fill_manual(values = c("#9964cb","#5fa369","#ca587a","#b38c3a","#798bc6")) +
  scale_shape_manual(values=c(21:25))+
  scale_color_manual(values="black")+
  scale_x_discrete(limits=c("1h", "3h", "6h", "8h", "24h"), labels=c("1hr", "3hr", "6hr", "8hr", "24hr")) + 
  scale_y_continuous(name="Number of genes detected \n after filtering")+
  xlab("Time Pre-Processing") +
  theme_classic() + 
  theme(legend.position = "none",
        panel.grid = element_blank(),
        panel.background = element_blank(),
        axis.text = element_text( colour = 'black', size = font_size),
        axis.title = element_text( colour = 'black', size = font_size ))

plot_grid( timeplot1,timeplot2,timeplot3,timeplot4, nrow=2, align='v')
```

```
rm("A260_A28_ratio_jugular","A260_A28_ratio_tail","A260_jugular","A260_tail","A280_jugular","A280_tail","bias_fit","bias_metrics","bias_metrics_a","bias_metrics_b",
  "count","counting_summary","counting_summary_a","counting_summary_b","efficiency_data","fg","files",
 "flg","fln","fn","fw","keep","n","n_reads_lncRNA", "n_reads_others","n_reads_protein_coding","n_reads_pseudogene",
 "number_of_reads_assigned","number_of_reads_produced","plot.legend","plot1","plot2","plot3","plot4","processing_time","processing_time_bias",
 "processing_time_bias_eff","random_genes_bef_filt_lme","random_genes_post_filt_lme"  ,                  
 "random_ratio","random_RIN_lme","reads_produced","reads_produced_a","reads_produced_b","RIN_jugular",
 "RIN_tail","sample", "summary_A260","summary_A260_location","summary_A280",
 "summary_A280_location","summary_bias","summary_genes_detected_bef_filtering","summary_genes_detected_bef_filtering_location" ,       "summary_genes_detected_post_filtering","summary_genes_detected_post_filtering_location","summary_ratio","summary_ratio_location","summary_reads_assigned",
 "summary_RIN","summary_RIN_location","summary_RNA_seq","three_prime_bias_mean","three_prime_bias_median",
 "timeplot1","timeplot2","timeplot3", "timeplot4","unassigned_Ambiguity","unassigned_NoFeatures")
```

##### Differential Gene Expression Analysis

We analyzed the data using edgeR and DESeq2, merge the results and infer signifiance based on the results of both packages.

##### Coccygeal vs Jugular vein

```
data_frame_sample_information<- RNA_data_frame
data_frame_sample_information$Name <- sub("-", "_", data_frame_sample_information$Name)
data_frame_sample_information$Heifer_ID<-as.factor(data_frame_sample_information$Heifer_ID)
data_frame_sample_information$location_blood_drawing<-as.factor(data_frame_sample_information$location_blood_drawing)
data_frame_sample_information_a<-data_frame_sample_information[data_frame_sample_information$Time_pre_processing=="1h",]
```

- edgeR

```
count_data_annotated_a<-count_data_annotated[,c(2:31)]
rownames(count_data_annotated_a)<-count_data_annotated$Row.names

count_data_annotated_a<-count_data_annotated_a[rownames(count_data_annotated_a) %in% genes_expressed,]

Heifer_ID<-factor(data_frame_sample_information_a$Heifer_ID)
location_blood_drawing<-factor(data_frame_sample_information_a$location_blood_drawing, levels=c("tail", "jugular"))
design <- model.matrix(~ Heifer_ID + location_blood_drawing)

Jugular_Tail_vein_count_data_a<-count_data_annotated_a[,colnames(count_data_annotated_a) %in% data_frame_sample_information_a$Seq_name ]

Jugular_Tail_vein<-DGEList(count=Jugular_Tail_vein_count_data_a, group=data_frame_sample_information_a$location_blood_drawing)
Jugular_Tail_vein<-estimateDisp(Jugular_Tail_vein,design, robust=TRUE)

# LRT method
#Jugular_Tail_vein_fit <- glmFit(Jugular_Tail_vein, design)
#Jugular_Tail_vein_LRT <- glmLRT(Jugular_Tail_vein_fit,coef=6)
#Jugular_Tail_vein_edgeR_results_LRT<- topTags(Jugular_Tail_vein_LRT, adjust.method = "fdr", n=Inf)$table

#QLFTest method
Jugular_Tail_vein_QLFit <- glmQLFit(Jugular_Tail_vein, design,robust=TRUE)
Jugular_Tail_vein_QLF <- glmQLFTest(Jugular_Tail_vein_QLFit,coef=6 )
Jugular_Tail_vein_edgeR_results_QLF<- topTags(Jugular_Tail_vein_QLF, adjust.method = "fdr", n=Inf)$table
```

- DESeq2

```
Jugular_Tail_vein<-DESeqDataSetFromMatrix(countData=Jugular_Tail_vein_count_data_a,colData=data_frame_sample_information_a, design= ~Heifer_ID + location_blood_drawing)

# Wald method
Jugular_Tail_vein_Wald<-DESeq(Jugular_Tail_vein, test="Wald")
Jugular_Tail_vein_results_DESeq_Wald<-results(Jugular_Tail_vein_Wald, contrast=c("location_blood_drawing",  "tail", "jugular"), pAdjustMethod="fdr", tidy=TRUE)
Jugular_Tail_vein_results_DESeq_Wald<-Jugular_Tail_vein_results_DESeq_Wald[with(Jugular_Tail_vein_results_DESeq_Wald, order(pvalue)),]

# LRT method
#Jugular_Tail_vein_LRT<-DESeq(Jugular_Tail_vein, test="LRT", reduced = ~Heifer_ID)
#Jugular_Tail_vein_results_DESeq_LRT<-results(Jugular_Tail_vein_LRT, contrast=c("location_blood_drawing",  "tail", "jugular"), pAdjustMethod="fdr", tidy=TRUE)
#Jugular_Tail_vein_results_DESeq_LRT<-Jugular_Tail_vein_results_DESeq_LRT[with(Jugular_Tail_vein_results_DESeq_LRT, order(pvalue)),]
```

##### Figure 2

Transcript abundances obtained from the jugular and coccygeal veins from different subjects. The values presented are variance stabilizing read counts obtained from the “DESeq2” package.

```
deseq2_Jugular_Tail_vein_VST<-varianceStabilizingTransformation(Jugular_Tail_vein_Wald, blind=TRUE)
deseq2_Jugular_Tail_vein_VST<-as.data.frame(assay(deseq2_Jugular_Tail_vein_VST))

font_size<-10
point_size<-0.2

plot1<-ggscatter(deseq2_Jugular_Tail_vein_VST, x="GSC22335", y="GSC22336",
          size = point_size,
          title = "Subject A",
          cor.coef = TRUE,
          cor.coeff.args = list(method = "pearson", label.sep = ", ",digits = 4, cor.coef.name = "r",size = 2),
          cor.coef.coord=c(6,18))+
       scale_x_continuous(breaks = seq(6,18, 2), limits=c(6,18))+
       scale_y_continuous(breaks = seq(6,18, 2), limits=c(6,18))+
       theme(aspect.ratio = 1)+
       theme(axis.title = element_blank(),
             text = element_text(size = font_size),
             plot.title = element_text( size = font_size))

plot2<-ggscatter(deseq2_Jugular_Tail_vein_VST, x="GSC22341", y="GSC22342",
          size = point_size,
          title = "Subject B",
          cor.coef = TRUE,
          cor.coeff.args = list(method = "pearson", label.sep = ", ",digits = 4, cor.coef.name = "r",size = 2),
          cor.coef.coord=c(6,18))+
       scale_x_continuous(breaks = seq(6,18, 2), limits=c(6,18))+
       scale_y_continuous(breaks = seq(6,18, 2), limits=c(6,18))+
       theme(aspect.ratio = 1)+
       theme(axis.title = element_blank(),
             text = element_text(size = font_size),
             plot.title = element_text( size = font_size))

plot3<-ggscatter(deseq2_Jugular_Tail_vein_VST, x="GSC22347", y="GSC22348",
          size = point_size,
           title = "Subject C",
          cor.coef = TRUE,
          cor.coeff.args = list(method = "pearson",  label.sep = ", ",digits = 4, cor.coef.name = "r",size = 2),
          cor.coef.coord=c(6,18))+
       scale_x_continuous(breaks = seq(6,18, 2), limits=c(6,18))+
       scale_y_continuous(breaks = seq(6,18, 2), limits=c(6,18))+
       theme(aspect.ratio = 1)+
       theme(axis.title = element_blank(),
             text = element_text(size = font_size),
             plot.title = element_text( size = font_size))

plot4<-ggscatter(deseq2_Jugular_Tail_vein_VST, x="GSC22353", y="GSC22354",
          size = point_size,
          cor.coef = TRUE,
           title = "Subject D",
          cor.coeff.args = list(method = "pearson", label.sep = ", ",digits = 4, cor.coef.name = "r",size = 2),
          cor.coef.coord=c(6,18))+
       scale_x_continuous(breaks = seq(6,18, 2), limits=c(6,18))+
       scale_y_continuous(breaks = seq(6,18, 2), limits=c(6,18))+
       theme(aspect.ratio = 1)+
       theme(axis.title = element_blank(),
             text = element_text(size = font_size),
             plot.title = element_text( size = font_size))

plot5<-ggscatter(deseq2_Jugular_Tail_vein_VST, x="GSC22359", y="GSC22360",
          size = point_size,
          xlab = "",
          ylab = "",
          title = "Subject E",
          cor.coef = TRUE,
          cor.coeff.args = list(method = "pearson", label.sep = ", ",digits = 4, cor.coef.name = "r", size = 2),
          cor.coef.coord=c(6,18))+
       scale_x_continuous(breaks = seq(6,18, 2), limits=c(6,18))+
       scale_y_continuous(breaks = seq(6,18, 2), limits=c(6,18))+
       theme(aspect.ratio = 1)+
       theme(axis.title = element_blank(),
             text = element_text(size = font_size),
             plot.title = element_text( size = font_size))

plots<-plot_grid( plot1,plot2,plot3,plot4,plot5, nrow = 1,align='v')

y.grob <- textGrob("Jugular vein", gp=gpar(fontface="plain", col="black", fontsize=font_size), rot=90)

x.grob <- textGrob("Coccygeal vein", gp=gpar(fontface="plain", col="black", fontsize=font_size))

grid.arrange(arrangeGrob(plots, left = y.grob, bottom = x.grob))
```

##### Time of Processing

- edgeR

```
data_frame_sample_information_b<-data_frame_sample_information[data_frame_sample_information$location_blood_drawing !="tail",]
rownames(data_frame_sample_information_b)<-data_frame_sample_information_b$Name
data_frame_sample_information_b$Heifer_ID<-factor(data_frame_sample_information_b$Heifer_ID)
data_frame_sample_information_b$Time_pre_processing<-factor(data_frame_sample_information_b$Time_pre_processing, levels=c("1h", "3h", "6h", "8h", "24h"))

Heifer_ID<-factor(data_frame_sample_information_b$Heifer_ID)
time_blood_processing<-factor(data_frame_sample_information_b$Time_pre_processing, levels=c("1h", "3h", "6h", "8h", "24h"))
design <- model.matrix(~ 0 + time_blood_processing + Heifer_ID)

time_blood_processing_count_data_a<-count_data_annotated_a[,colnames(count_data_annotated_a) %in% data_frame_sample_information_b$Seq_name ]

time_blood_processing<-DGEList(count=time_blood_processing_count_data_a)
time_blood_processing<-estimateDisp(time_blood_processing,design, robust=TRUE)

#QLFTest method

time_blood_processing_QLFit <- glmQLFit(time_blood_processing, design,robust=TRUE)
#time_blood_processing_QLF <- glmQLFTest(time_blood_processing_QLFit, coef=1:5)
#time_blood_processing_edgeR_results_QLF<- topTags(time_blood_processing_QLF, adjust.method = "fdr", n=Inf)$table


my_contrasts<-makeContrasts('24hvs1h'=time_blood_processing24h-time_blood_processing1h,
                            '8hvs1h'=time_blood_processing8h-time_blood_processing1h  ,
                            '6hvs1h'=time_blood_processing6h-time_blood_processing1h ,
                            '3hvs1h'=time_blood_processing6h-time_blood_processing1h , levels = design)

time_blood_processing_QLF <- glmQLFTest(time_blood_processing_QLFit, contrast=my_contrasts[,'3hvs1h'])
time_blood_processing_edgeR_QLF_results_3hvs1h<- topTags(time_blood_processing_QLF, adjust.method = "fdr", n=Inf)$table
table(time_blood_processing_edgeR_QLF_results_3hvs1h$FDR < 0.05)
```

```
## 
## FALSE 
## 12810
```

```
rm(time_blood_processing_QLF)

time_blood_processing_QLF <- glmQLFTest(time_blood_processing_QLFit, contrast=my_contrasts[,'6hvs1h'])
time_blood_processing_edgeR_QLF_results_6hvs1h<- topTags(time_blood_processing_QLF, adjust.method = "fdr", n=Inf)$table
table(time_blood_processing_edgeR_QLF_results_6hvs1h$FDR < 0.05)
```

```
## 
## FALSE 
## 12810
```

```
rm(time_blood_processing_QLF)

time_blood_processing_QLF <- glmQLFTest(time_blood_processing_QLFit, contrast=my_contrasts[,'8hvs1h'])
time_blood_processing_edgeR_QLF_results_8hvs1h<- topTags(time_blood_processing_QLF, adjust.method = "fdr", n=Inf)$table
table(time_blood_processing_edgeR_QLF_results_8hvs1h$FDR < 0.05)
```

```
## 
## FALSE  TRUE 
## 12806     4
```

```
rm(time_blood_processing_QLF)

time_blood_processing_QLF <- glmQLFTest(time_blood_processing_QLFit, contrast=my_contrasts[,'24hvs1h'])
time_blood_processing_edgeR_QLF_results_24hvs1h<- topTags(time_blood_processing_QLF, adjust.method = "fdr", n=Inf)$table
table(time_blood_processing_edgeR_QLF_results_24hvs1h$FDR < 0.05)
```

```
## 
## FALSE  TRUE 
## 10528  2282
```

- DESeq2

```
time_blood_processing<-DESeqDataSetFromMatrix(countData=time_blood_processing_count_data_a,colData=data_frame_sample_information_b, design= ~Time_pre_processing +Heifer_ID )

#vsd_time_blood_processing <- varianceStabilizingTransformation(time_blood_processing)

# Wald method

time_blood_processing_Wald <- DESeq(time_blood_processing, test="Wald")

#time_blood_processing_results_DESeq_Wald<-results(time_blood_processing_Wald,pAdjustMethod="fdr", tidy=TRUE)
#time_blood_processing_results_DESeq_Wald<-time_blood_processing_results_DESeq_Wald[with(time_blood_processing_results_DESeq_Wald, order(pvalue)),]

#table(time_blood_processing_results_DESeq_Wald$padj < 0.05)

time_blood_processing_DESeq_Wald_result_3hvs1h<-results(time_blood_processing_Wald,pAdjustMethod="fdr", tidy=TRUE, contrast=c("Time_pre_processing", "3h","1h"))
time_blood_processing_DESeq_Wald_result_3hvs1h<-time_blood_processing_DESeq_Wald_result_3hvs1h[with(time_blood_processing_DESeq_Wald_result_3hvs1h, order(pvalue)),]
table(time_blood_processing_DESeq_Wald_result_3hvs1h$padj < 0.05)
```

```
## 
## FALSE 
## 12810
```

```
time_blood_processing_DESeq_Wald_result_6hvs1h<-results(time_blood_processing_Wald,pAdjustMethod="fdr", tidy=TRUE, contrast=c("Time_pre_processing", "6h","1h"))
time_blood_processing_DESeq_Wald_result_6hvs1h<-time_blood_processing_DESeq_Wald_result_6hvs1h[with(time_blood_processing_DESeq_Wald_result_6hvs1h, order(pvalue)),]
table(time_blood_processing_DESeq_Wald_result_6hvs1h$padj < 0.05)
```

```
## 
## FALSE  TRUE 
## 12768    42
```

```
time_blood_processing_DESeq_Wald_result_8hvs1h<-results(time_blood_processing_Wald,pAdjustMethod="fdr", tidy=TRUE, contrast=c("Time_pre_processing", "8h","1h") )
time_blood_processing_DESeq_Wald_result_8hvs1h<-time_blood_processing_DESeq_Wald_result_8hvs1h[with(time_blood_processing_DESeq_Wald_result_8hvs1h, order(pvalue)),]
table(time_blood_processing_DESeq_Wald_result_8hvs1h$padj < 0.05)
```

```
## 
## FALSE  TRUE 
## 12804     6
```

```
time_blood_processing_DESeq_Wald_result_24hvs1h<-results(time_blood_processing_Wald,pAdjustMethod="fdr", tidy=TRUE, contrast=c("Time_pre_processing", "24h","1h") )
time_blood_processing_DESeq_Wald_result_24hvs1h<-time_blood_processing_DESeq_Wald_result_24hvs1h[with(time_blood_processing_DESeq_Wald_result_24hvs1h, order(pvalue)),]
table(time_blood_processing_DESeq_Wald_result_24hvs1h$padj < 0.05)
```

```
## 
## FALSE  TRUE 
## 10266  2544
```

Overview of the overlap of the results between the two algorithms

```
venn_diagram<-venn.diagram(
  list("edgeR_8hvs1h"=rownames(time_blood_processing_edgeR_QLF_results_8hvs1h[time_blood_processing_edgeR_QLF_results_8hvs1h$FDR<0.05,]),
       "DESeq2_8hvs1h"=time_blood_processing_DESeq_Wald_result_8hvs1h[time_blood_processing_DESeq_Wald_result_8hvs1h$padj<0.05,]$row,
       "edgeR_24hvs1h"=rownames(time_blood_processing_edgeR_QLF_results_24hvs1h[time_blood_processing_edgeR_QLF_results_24hvs1h$FDR<0.05,]),
       "DESeq2_24hvs1h"=time_blood_processing_DESeq_Wald_result_24hvs1h[time_blood_processing_DESeq_Wald_result_24hvs1h$padj<0.05,]$row),
  cex=1,
  scaled= FALSE, 
  filename = NULL,
  #filename = "/2021_04_02_venndiagram_DEGs.tiff",
  height = 800, width = 1000,
  output=TRUE,
  col="transparent",
  fill=c("#56B4E9",  "#f0e442", "#0071b2","#E69F00"),
  #cat.pos=c(1,-1),
  cat.cex=c(1, 1, 1, 1),
  margin=c(0.2)
  )

grid.draw(venn_diagram)
```

##### Figure 3

Comparison of transcript abundance in PWBCs from blood samples processed after prolonged storage at 4°C. A. M-A plots of the contrasts between each of the prolonged storage times versus samples processed within one hour. B. Transcript abundance for genes with differential abundance in both the ‘8hr vs 1hr’ and ‘24hr vs 1hr’ contrasts.

##### Additional file 5

Results of differential transcript abundance for the contrast ‘24h vs 1h’.

```
overlap_3hvs1h<-merge(time_blood_processing_edgeR_QLF_results_3hvs1h , time_blood_processing_DESeq_Wald_result_3hvs1h, by.x='row.names',by.y="row", all=FALSE)

overlap_3hvs1h$result<-"neither"
overlap_3hvs1h$result[overlap_3hvs1h$FDR<0.05 & overlap_3hvs1h$padj<0.05 &  (overlap_3hvs1h$logFC < -0.5 | overlap_3hvs1h$logFC >  0.5)]<-"both"
overlap_3hvs1h$result[overlap_3hvs1h$FDR<0.05 & overlap_3hvs1h$padj>=0.05 & (overlap_3hvs1h$logFC < -0.5 | overlap_3hvs1h$logFC >  0.5)]<-"edger"
overlap_3hvs1h$result[overlap_3hvs1h$FDR>=0.05 & overlap_3hvs1h$padj<0.05 & (overlap_3hvs1h$logFC < -0.5 | overlap_3hvs1h$logFC >  0.5)]<-"deseq2"
overlap_3hvs1h$color<-"gray"
overlap_3hvs1h$color[overlap_3hvs1h$logFC < -0.5 & (overlap_3hvs1h$FDR<0.05 | overlap_3hvs1h$padj<0.05)]<-"down"
overlap_3hvs1h$color[overlap_3hvs1h$logFC >  0.5 & (overlap_3hvs1h$FDR<0.05 | overlap_3hvs1h$padj<0.05)]<-"up"
overlap_3hvs1h$color<-factor(overlap_3hvs1h$color, levels=c("up","gray","down"))

font_size<-10
legend_font_size<-6
shape_size<-0.7

plot1<-ggplot(data=overlap_3hvs1h , aes(x=logCPM,y=logFC, shape=result, color=color))+
geom_point(size=shape_size)+
#scale_y_continuous(name=c("LogFC"))+
#scale_x_continuous(name=c("LogCPM"))+
scale_shape_manual(values=c(3))+
scale_color_manual(values=c("gray"))+
ggtitle("3hr vs 1hr")+
theme_classic()+
theme(legend.position = "none",
       plot.title = element_text(hjust = 0.5, size = font_size, color="black"),
       axis.title = element_blank(),
       axis.text  = element_text(size = font_size, color="black")
      )


overlap_6hvs1h<-merge(time_blood_processing_edgeR_QLF_results_6hvs1h , time_blood_processing_DESeq_Wald_result_6hvs1h, by.x='row.names',by.y="row", all=FALSE)

overlap_6hvs1h$result<-"neither"
overlap_6hvs1h$result[overlap_6hvs1h$FDR<0.05 & overlap_6hvs1h$padj<0.05 &  (overlap_6hvs1h$logFC < -0.5 | overlap_6hvs1h$logFC >  0.5)]<-"both"
overlap_6hvs1h$result[overlap_6hvs1h$FDR<0.05 & overlap_6hvs1h$padj>=0.05 & (overlap_6hvs1h$logFC < -0.5 | overlap_6hvs1h$logFC >  0.5)]<-"edger"
overlap_6hvs1h$result[overlap_6hvs1h$FDR>=0.05 & overlap_6hvs1h$padj<0.05 & (overlap_6hvs1h$logFC < -0.5 | overlap_6hvs1h$logFC >  0.5)]<-"deseq2"
overlap_6hvs1h$color<-"gray"
overlap_6hvs1h$color[overlap_6hvs1h$logFC < -0.5 & (overlap_6hvs1h$FDR<0.05 | overlap_6hvs1h$padj<0.05)]<-"down"
overlap_6hvs1h$color[overlap_6hvs1h$logFC >  0.5 & (overlap_6hvs1h$FDR<0.05 | overlap_6hvs1h$padj<0.05)]<-"up"
overlap_6hvs1h$color<-factor(overlap_6hvs1h$color, levels=c("up", "down", "gray"))
overlap_6hvs1h$result<-factor(overlap_6hvs1h$result, levels=c("deseq2", "neither"))
overlap_6hvs1h<-overlap_6hvs1h[order(overlap_6hvs1h$color,decreasing = TRUE),]


plot2<-ggplot(data=overlap_6hvs1h , aes(x=logCPM,y=logFC, shape=result, color=color))+
geom_point(size=shape_size)+
#scale_y_continuous(name=c("LogFC"))+
#scale_x_continuous(name=c("LogCPM"))+
scale_shape_manual(values=c(15,3))+
scale_color_manual(values=c("blue",  "red", "gray"))+
ggtitle("6hr vs 1hr")+
theme_classic()+
theme(legend.position = "none",
       plot.title = element_text(hjust = 0.5, size = font_size, color="black"),
       axis.title = element_blank(),
       axis.text  = element_text(size = font_size, color="black")
      )


overlap_8hvs1h<-merge(time_blood_processing_edgeR_QLF_results_8hvs1h , time_blood_processing_DESeq_Wald_result_8hvs1h, by.x='row.names',by.y="row", all=FALSE)

overlap_8hvs1h$result<-"neither"
overlap_8hvs1h$result[overlap_8hvs1h$FDR<0.05 & overlap_8hvs1h$padj<0.05 &  (overlap_8hvs1h$logFC < -0.5 | overlap_8hvs1h$logFC >  0.5)]<-"both"
overlap_8hvs1h$result[overlap_8hvs1h$FDR<0.05 & overlap_8hvs1h$padj>0.05 & (overlap_8hvs1h$logFC < -0.5 | overlap_8hvs1h$logFC >  0.5)]<-"edger"
overlap_8hvs1h$result[overlap_8hvs1h$FDR>0.05 & overlap_8hvs1h$padj<0.05 & (overlap_8hvs1h$logFC < -0.5 | overlap_8hvs1h$logFC >  0.5)]<-"deseq2"
overlap_8hvs1h$result<-factor(overlap_8hvs1h$result, levels=c("both",  "deseq2", "neither"))

overlap_8hvs1h$color<-"gray"
overlap_8hvs1h$color[overlap_8hvs1h$logFC < -0.5 & (overlap_8hvs1h$FDR<0.05 | overlap_8hvs1h$padj<0.05)]<-"down"
overlap_8hvs1h$color[overlap_8hvs1h$logFC >  0.5 & (overlap_8hvs1h$FDR<0.05 | overlap_8hvs1h$padj<0.05)]<-"up"
overlap_8hvs1h$color<-factor(overlap_8hvs1h$color, levels=c("up","down","gray"))
overlap_8hvs1h<-overlap_8hvs1h[order(overlap_8hvs1h$color,decreasing = TRUE),]

plot3<-ggplot(data=overlap_8hvs1h , aes(x=logCPM,y=logFC, shape=result, color=color))+
geom_point(size=shape_size)+
#scale_y_continuous(name=c("LogFC"))+
#scale_x_continuous(name=c("LogCPM"))+
scale_shape_manual(values=c(16,15,3))+
scale_color_manual(values=c("blue", "gray","red"))+
ggtitle("8hr vs 1hr")+
theme_classic()+
theme(legend.position = "none",
       plot.title = element_text(hjust = 0.5, size = font_size, color="black"),
       axis.title = element_blank(),
       axis.text  = element_text(size = font_size, color="black")
      )

overlap_24hvs1h<-merge(time_blood_processing_edgeR_QLF_results_24hvs1h , time_blood_processing_DESeq_Wald_result_24hvs1h, by.x='row.names',by.y="row", all=FALSE)

#overlap_24hvs1h<-merge(overlap_24hvs1h, annotation.ensembl.symbol, by.x="Row.names", by.y="ensembl_gene_id", all.x=TRUE, all.y=FALSE)
#write_delim(overlap_24hvs1h, "/2021_12_21_Additional_file_5.txt", delim = "\t", quote =  "none")

overlap_24hvs1h$result<-"neither"
overlap_24hvs1h$result[overlap_24hvs1h$FDR<0.05 & overlap_24hvs1h$padj<0.05 &  (overlap_24hvs1h$logFC < -0.5 | overlap_24hvs1h$logFC >  0.5)]<-"both"
overlap_24hvs1h$result[overlap_24hvs1h$FDR<0.05 & overlap_24hvs1h$padj>=0.05 & (overlap_24hvs1h$logFC < -0.5 | overlap_24hvs1h$logFC >  0.5)]<-"edger"
overlap_24hvs1h$result[overlap_24hvs1h$FDR>=0.05 & overlap_24hvs1h$padj<0.05 & (overlap_24hvs1h$logFC < -0.5 | overlap_24hvs1h$logFC >  0.5)]<-"deseq2"
overlap_24hvs1h$color<-"gray"
overlap_24hvs1h$color[overlap_24hvs1h$logFC < -0.5 & (overlap_24hvs1h$FDR<0.05 | overlap_24hvs1h$padj<0.05)]<-"down"
overlap_24hvs1h$color[overlap_24hvs1h$logFC >  0.5 & (overlap_24hvs1h$FDR<0.05 | overlap_24hvs1h$padj<0.05)]<-"up"
overlap_24hvs1h$color<-factor(overlap_24hvs1h$color, levels=c(  "up","down","gray"))
overlap_24hvs1h$result<-factor(overlap_24hvs1h$result, levels=c(  "both",  "deseq2",   "edger", "neither" ))
overlap_24hvs1h<-overlap_24hvs1h[order(overlap_24hvs1h$color,decreasing = TRUE),]

plot4<-ggplot(data=overlap_24hvs1h , aes(x=logCPM,y=logFC, shape=result, color=color))+
geom_point(size=shape_size)+
scale_y_continuous(name=c("LogFC"))+
scale_x_continuous(name=c("LogCPM"))+
scale_shape_manual(name="Statistical significance" ,values=c(16,15,17,3), labels=c("FDR<0.05 edgeR and DESeq2","FDR<0.05 DESeq2","FDR<0.05 edgeR", "FDR>0.05"))+
scale_color_manual(name= "Direction of differential transcript abundance |Log(FC)|>0.5", values=c("blue",  "red", "gray"), labels=c("delayed processing > 1h","delayed processing < 1h","delayed processing ~ 1h"))+
ggtitle("24hr vs 1hr")+
theme_classic()+
theme(plot.title = element_text(hjust = 0.5, size = font_size, color="black"),
       axis.title = element_text(size = font_size, color="black"),
       axis.text  = element_text(size = font_size, color="black"),
      legend.text = element_text(size = legend_font_size, color="black", margin = margin(0,0,0,0)),
      legend.title = element_text(size = legend_font_size, color="black"),
      legend.position = "bottom",
      legend.text.align = 1,
      legend.spacing = unit(0,unit="pt"),
      legend.margin=margin(0,unit="pt"))+
      guides(colour = guide_legend(title.position="top", title.hjust = 0.5),
             shape = guide_legend(title.position="top", title.hjust = 0.5))

legend<-get_legend(plot4)

plot4<-ggplot(data=overlap_24hvs1h , aes(x=logCPM,y=logFC, shape=result, color=color))+
geom_point(size=shape_size)+
#scale_y_continuous(name=c("LogFC"))+
#scale_x_continuous(name=c("LogCPM"))+
scale_shape_manual(name=NULL,values=c(16,15,17,3), labels=c("FDR<0.05 edgeR and DESeq2","FDR<0.05 DESeq2","FDR<0.05 edgeR", "FDR>0.05"))+
scale_color_manual(name= "direction of differential\ntranscript abundance\n|Log(FC)|>0.5", values=c("blue",  "red", "gray"), labels=c("24h > 1h","24h < 1h","24h ~ 1h"))+
ggtitle("24hr vs 1hr")+
theme_classic()+
theme(legend.position = "none",
       plot.title = element_text(hjust = 0.5, size = font_size, color="black"),
       axis.title = element_blank(),
       axis.text  = element_text(size = font_size, color="black") )

#pdf(file="/2021_04_02_M_A_plots.pdf", width=12,height=4, bg="transparent", onefile=TRUE)

plots_a <- plot_grid(plot1, plot2, plot3, plot4, nrow = 1, ncol=4 , align = 'h', labels=c('A'), label_size=8, label_fontface="plain", label_x= -0.1)

y.grob <- textGrob("Log(FC)", gp=gpar(fontface="plain", col="black", fontsize=font_size), rot=90)

x.grob <- textGrob("Log(CPM)", gp=gpar(fontface="plain", col="black", fontsize=font_size))

plots_b<-plot_grid(  grid.arrange(arrangeGrob(plots_a, left = y.grob, bottom = x.grob)), legend, nrow = 2, ncol=1 ,rel_heights = c(1, .2))
```

```
data_chart_1<-data.frame( stringsAsFactors=FALSE)
data_chart_2<-data.frame( stringsAsFactors=FALSE)
overlap_8hvs1h_sig<-overlap_8hvs1h[overlap_8hvs1h$FDR < 0.05 & overlap_8hvs1h$padj < 0.05 & (overlap_8hvs1h$logFC < -0.5 | overlap_8hvs1h$logFC >  0.5), ]
overlap_8hvs1h_sig<-merge(overlap_8hvs1h_sig, annotation.ensembl.symbol, by.x="Row.names", by.y="ensembl_gene_id", all=FALSE)

data_frame_sample_information_c<-data_frame_sample_information_b[order(data_frame_sample_information_b$Time_pre_processing),]

data_fpkm_filtered_graph<-data_fpkm_filtered[rownames(data_fpkm_filtered) %in% overlap_8hvs1h_sig$Row.names,]
data_fpkm_filtered_graph<-data_fpkm_filtered_graph[,colnames(data_fpkm_filtered_graph) %in% data_frame_sample_information_c$Seq_name]
data_fpkm_filtered_graph<-data_fpkm_filtered_graph[,data_frame_sample_information_c$Seq_name]

for (i in  seq(dim(overlap_8hvs1h_sig)[1])){
  
  gene_id<-overlap_8hvs1h_sig[i,1]
  gene_symbol<-overlap_8hvs1h_sig$external_gene_name[i]
  if(gene_symbol==""){gene_symbol<-gene_id}
  
  data_chart_1<-data.frame(fpkm=data_fpkm_filtered_graph[rownames(data_fpkm_filtered_graph) %in% overlap_8hvs1h_sig[i,1],])
  data_chart_1$group<-as.factor( data_frame_sample_information_c$Time_pre_processing )
  data_chart_1$shape<-as.factor( data_frame_sample_information_c$Heifer_ID )
  data_chart_1$gene<-gene_symbol
  data_chart_1$chart<-i
  data_chart_1$fold_change<-overlap_8hvs1h_sig[overlap_8hvs1h_sig$Row.names==gene_id,]$logFC
  data_chart_1$pvalue<-overlap_8hvs1h_sig[overlap_8hvs1h_sig$Row.names==gene_id,]$FDR
  
  data_chart_2<-rbind(data_chart_2,data_chart_1)
}

plots_c <- list()
for (i in seq(1,dim(overlap_8hvs1h_sig)[1])) {
    data_chart_3<-data_chart_2[data_chart_2$chart %in% i , ]
    plot<-ggplot(data=data_chart_3, aes(x=group , y=fpkm))+
    stat_summary(fun = median,colour = "gray", size = 0.1,  geom = "crossbar", alpha=0.5)+
     geom_jitter(position = position_jitter(w=0.2, h=0), aes(shape=shape, fill=group), size=0.6) +
     scale_fill_manual(values = c("#9964cb","#5fa369","#ca587a","#b38c3a","#798bc6")) +
     scale_shape_manual(values=c(21:25))+
     scale_color_manual(values="black")+
     scale_y_continuous("")+
     scale_x_discrete(limits=c("1h", "3h", "6h", "8h", "24h"), labels=c("1hr", "3hr", "6hr", "8hr", "24hr")) +
     xlab("") +
     ggtitle(paste(data_chart_3$gene[1]))+
     #ggtitle(paste(data_chart_3$gene[1], "  ", "LogFC","=", round(data_chart_3$fold_change[1],1),"  ","P","=",round(data_chart_3$pvalue[1],4), sep=c("")))+
     theme_classic(base_size = 10) + 
     theme(legend.position = "none",
           plot.title = element_text(size = 8),
           panel.grid = element_blank(),
           panel.background = element_blank(),
           axis.text = element_text(size = font_size, colour = 'black'),
           axis.title = element_text( size = font_size, colour = 'black'),
           plot.margin = unit(c(0.5, 0, 0,0), "cm"))
      
    plots_c[[i]] <- plot
  }

plots_d<-plot_grid(  plotlist=plots_c, nrow=1, align = 'v', axis="l", labels = "B", label_x= -0.1,label_size=8, label_fontface = "plain")

y.grob <- textGrob("FPKM", gp=gpar(fontface="plain", col="black", fontsize=font_size), rot=90)

x.grob <- textGrob("Time of delayed processing", gp=gpar(fontface="plain", col="black", fontsize=font_size))

plots_e<-plot_grid(arrangeGrob(plots_d, left = y.grob, bottom = x.grob))
```

```
grid.arrange( plots_b,plots_e , nrow=2, heights=c(1.5,1.3))
```

##### Additional file 6

Charts of transcript abundance (fragment per kilobase per million reads, FPKM) for different times of refrigeration (4°C) prior to processing and cryopreservation of PWBCs. Animals are indicated by shapes across charts.

```
multiplot <- function(..., plotlist=NULL, file, cols=1, layout=NULL) {
  library(grid)
  
  # Make a list from the ... arguments and plotlist
  plots <- c(list(...), plotlist)
  
  numPlots = length(plots)
  
  # If layout is NULL, then use 'cols' to determine layout
  if (is.null(layout)) {
    # Make the panel
    # ncol: Number of columns of plots
    # nrow: Number of rows needed, calculated from # of cols
    layout <- matrix(seq(1, cols * ceiling(numPlots/cols)),
                     ncol = cols, nrow = ceiling(numPlots/cols))
  }
  
  if (numPlots==1) {
    print(plots[[1]])
    
  } else {
    # Set up the page
    grid.newpage()
    pushViewport(viewport(layout = grid.layout(nrow(layout), ncol(layout))))
    
    # Make each plot, in the correct location
    for (i in 1:numPlots) {
      # Get the i,j matrix positions of the regions that contain this subplot
      matchidx <- as.data.frame(which(layout == i, arr.ind = TRUE))
      
      print(plots[[i]], vp = viewport(layout.pos.row = matchidx$row,
                                      layout.pos.col = matchidx$col))
    }
  }
}

data_chart_1<-data.frame( stringsAsFactors=FALSE)
data_chart_2<-data.frame( stringsAsFactors=FALSE)
overlap_24hvs1h_sig<-overlap_24hvs1h[overlap_24hvs1h$FDR < 0.05 & overlap_24hvs1h$padj < 0.05 & (overlap_24hvs1h$logFC < -0.5 | overlap_24hvs1h$logFC >  0.5), ]
overlap_24hvs1h_sig<-merge(overlap_24hvs1h_sig, annotation.ensembl.symbol, by.x="Row.names", by.y="ensembl_gene_id", all=FALSE)
overlap_24hvs1h_sig<-overlap_24hvs1h_sig[order(overlap_24hvs1h_sig$logFC),]

data_frame_sample_information_c<-data_frame_sample_information_b[order(data_frame_sample_information_b$Time_pre_processing),]

data_fpkm_filtered_graph<-data_fpkm_filtered[rownames(data_fpkm_filtered) %in% overlap_24hvs1h_sig$Row.names,]
data_fpkm_filtered_graph<-data_fpkm_filtered_graph[,colnames(data_fpkm_filtered_graph) %in% data_frame_sample_information_c$Seq_name]
data_fpkm_filtered_graph<-data_fpkm_filtered_graph[,data_frame_sample_information_c$Seq_name]


for (i in  seq(dim(overlap_24hvs1h_sig)[1])){
  
  gene_id<-overlap_24hvs1h_sig[i,1]
  gene_symbol<-overlap_24hvs1h_sig$external_gene_name[i]
  if(gene_symbol==""){gene_symbol<-gene_id}
  
  data_chart_1<-data.frame(fpkm=data_fpkm_filtered_graph[rownames(data_fpkm_filtered_graph) %in% overlap_24hvs1h_sig[i,1],])
  data_chart_1$group<-as.factor( data_frame_sample_information_c$Time_pre_processing )
  data_chart_1$shape<-as.factor( data_frame_sample_information_c$Heifer_ID )
  data_chart_1$gene<-gene_symbol
  data_chart_1$chart<-i
  data_chart_1$fold_change<-overlap_24hvs1h_sig[overlap_24hvs1h_sig$Row.names==gene_id,]$logFC
  data_chart_1$pvalue<-overlap_24hvs1h_sig[overlap_24hvs1h_sig$Row.names==gene_id,]$FDR
  
  data_chart_2<-rbind(data_chart_2,data_chart_1)
}

#pdf(file="/2021_12_20_Additional_file_6.pdf", paper="letter", bg="transparent", onefile=TRUE)

for (i in seq(1,dim(overlap_24hvs1h_sig)[1], 12)) {
  plots <- list()
  k<-1
  for (j in c(i:(i+11))){
    data_chart_3<-data_chart_2[data_chart_2$chart %in% j , ]
    plot<-ggplot(data=data_chart_3, aes(x=group , y=fpkm))+
     geom_jitter(position = position_jitter(w=0.1, h=0), aes(shape=shape, fill=group), size=1) +
     scale_fill_manual(values = c("#9964cb","#5fa369","#ca587a","#b38c3a","#798bc6")) +
     scale_shape_manual(values=c(21:25))+
     scale_color_manual(values="black")+
     scale_y_continuous("FPKM")+
     scale_x_discrete(limits=c("1h", "3h", "6h", "8h", "24h"), labels=c("1hr", "3hr", "6hr", "8hr", "24hr")) +
     xlab("Time Pre-Processing") +
     stat_summary(fun = median,colour = "gray", size = 0.1,  geom = "crossbar", alpha=0.5)+
     ggtitle(paste(data_chart_3$gene[1], "  ", "LogFC","=", round(data_chart_3$fold_change[1],1), sep=c("")))+
     theme_classic(base_size = 10) + 
     theme(legend.position = "none",
           plot.title = element_text(size = 5),
           panel.grid = element_blank(),
           panel.background = element_blank(),
           axis.text = element_text( colour = 'black'),
           axis.title = element_text( colour = 'black'),
  )
    plots[[k]] <- plot
    k<-k+1
  }
  multiplot(plotlist = plots, cols = 3,layout = matrix(1:12, nrow=4, byrow = TRUE))
}
```

```
#dev.off()
```

##### Figure 4

Comparison of overall transcript sequencing coverage from samples that were processed within 1hr and 24hr after blood collection. A. Genes with no change in transcript abundance 24hr of storage at 4°C. B. Genes with lower transcript abundance after 24hr of storage at 4°C. C. Plots of the Delta D-statistic for all genes separated by their results of the contrast ‘24hr vs 1hr’.

```
#Figure 4A
genes_lower_abundance_24hvs1h<- overlap_24hvs1h[overlap_24hvs1h$result == "both" & overlap_24hvs1h$color=="down",]$Row.names
genes_greater_abundance_24hvs1h<- overlap_24hvs1h[overlap_24hvs1h$result == "both" & overlap_24hvs1h$color=="up",]$Row.names


GSC22336[,DEG:=ifelse(gene_id %in% genes_lower_abundance_24hvs1h, "DEG","noDEG")]
GSC22340[,DEG:=ifelse(gene_id %in% genes_lower_abundance_24hvs1h, "DEG","noDEG")]

GSC22342[,DEG:=ifelse(gene_id %in% genes_lower_abundance_24hvs1h, "DEG","noDEG")]
GSC22346[,DEG:=ifelse(gene_id %in% genes_lower_abundance_24hvs1h, "DEG","noDEG")]

GSC22348[,DEG:=ifelse(gene_id %in% genes_lower_abundance_24hvs1h, "DEG","noDEG")]
GSC22352[,DEG:=ifelse(gene_id %in% genes_lower_abundance_24hvs1h, "DEG","noDEG")]

GSC22354[,DEG:=ifelse(gene_id %in% genes_lower_abundance_24hvs1h, "DEG","noDEG")]
GSC22358[,DEG:=ifelse(gene_id %in% genes_lower_abundance_24hvs1h, "DEG","noDEG")]

GSC22360[,DEG:=ifelse(gene_id %in% genes_lower_abundance_24hvs1h, "DEG","noDEG")]
GSC22364[,DEG:=ifelse(gene_id %in% genes_lower_abundance_24hvs1h, "DEG","noDEG")]


temp_data1<-rbind(GSC22336[,time:="1hr"], GSC22340[,time:="24hr"])
temp_data2<-rbind(GSC22342[,time:="1hr"], GSC22346[,time:="24hr"])
temp_data3<-rbind(GSC22348[,time:="1hr"], GSC22352[,time:="24hr"])
temp_data4<-rbind(GSC22354[,time:="1hr"], GSC22358[,time:="24hr"])
temp_data5<-rbind(GSC22360[,time:="1hr"], GSC22364[,time:="24hr"])

font_size<-8
line_size<-0.5

plot1<-ggplot() +
geom_density(aes(x=rel_position, weight=prop, color=time), data=temp_data1[DEG=="noDEG",], adjust=0.4) +
scale_color_manual(name="Processing time",values=c("#00379c","#f04177"))+
scale_x_continuous(name="Relative transcript position (5' - 3')", breaks=c(0, 0.5, 1))+
ggtitle("Subject A")+
theme_classic()+
theme(
text=element_text(size=font_size, color="black", face="plain"),
axis.text = element_text( size=font_size, colour = 'black', face="plain"),
axis.title = element_blank(),
plot.title = element_text(size=font_size, hjust = 0.5),
legend.position="none",
axis.line.x = element_line(size=line_size),
axis.line.y = element_line(size=line_size),
axis.ticks = element_line(size=line_size))


plot2<-ggplot() +
geom_density(aes(x=rel_position, weight=prop,  color=time), data=temp_data2[DEG=="noDEG",], adjust=0.4) +
scale_color_manual(name="Processing time",values=c("#00379c","#f04177"))+
scale_x_continuous(name="Relative transcript position (5' - 3')", breaks=c(0, 0.5, 1))+
ggtitle("Subject B")+
theme_classic()+
theme(
text=element_text(size=font_size, color="black", face="plain"),
axis.text = element_text( size=font_size, colour = 'black', face="plain"),
axis.title = element_blank(),
plot.title = element_text(size=font_size, hjust = 0.5),
legend.position="none",
axis.line.x = element_line(size=line_size),
axis.line.y = element_line(size=line_size),
axis.ticks = element_line(size=line_size))


plot3<-ggplot() +
geom_density(aes(x=rel_position, weight=prop, color=time), data=temp_data3[DEG=="noDEG",], adjust=0.4) +
scale_color_manual(name="Processing time",values=c("#00379c","#f04177"))+
scale_x_continuous(name="Relative transcript position (5' - 3')", breaks=c(0, 0.5, 1))+
ggtitle("Subject C")+
theme_classic()+
theme(
text=element_text(size=font_size, color="black", face="plain"),
axis.text = element_text( size=font_size, colour = 'black', face="plain"),
axis.title = element_blank(),
plot.title = element_text(size=font_size, hjust = 0.5),
legend.position="none",
axis.line.x = element_line(size=line_size),
axis.line.y = element_line(size=line_size),
axis.ticks = element_line(size=line_size))


plot4<-ggplot() +
geom_density(aes(x=rel_position, weight=prop,  color=time), data=temp_data4[DEG=="noDEG",], adjust=0.4) +
scale_color_manual(name="Processing time",values=c("#00379c","#f04177"))+
scale_x_continuous(name="Relative transcript position (5' - 3')", breaks=c(0, 0.5, 1))+
ggtitle("Subject D")+
theme_classic()+
theme(
text=element_text(size=font_size, color="black", face="plain"),
axis.text = element_text( size=font_size, colour = 'black', face="plain"),
axis.title = element_blank(),
plot.title = element_text(size=font_size, hjust = 0.5),
legend.position="none",
axis.line.x = element_line(size=line_size),
axis.line.y = element_line(size=line_size),
axis.ticks = element_line(size=line_size))

plot5<-ggplot() +
geom_density(aes(x=rel_position, weight=prop, color=time), data=temp_data5[DEG=="noDEG",], adjust=0.4) +
scale_color_manual(name="Processing time",values=c("#00379c","#f04177"))+
scale_x_continuous(name="Relative transcript position (5' - 3')", breaks=c(0, 0.5, 1))+
ggtitle("Subject E")+
theme_classic()+
theme(
text=element_text(size=font_size, color="black", face="plain"),
axis.text = element_text( size=font_size, colour = 'black', face="plain"),
axis.title = element_blank(),
plot.title = element_text(size=font_size, hjust = 0.5),
legend.position="none",
axis.line.x = element_line(size=line_size),
axis.line.y = element_line(size=line_size),
axis.ticks = element_line(size=line_size))

y.grob <- textGrob("Density", gp=gpar(fontface="plain", col="black", fontsize=font_size), rot=90)
x.grob <- textGrob("Relative transcript position (5' - 3')", gp=gpar(fontface="plain", col="black", fontsize=font_size))

plot_a<-plot_grid( grid.arrange(arrangeGrob(plot_grid( plot1, plot2, plot3, plot4, plot5 ,nrow = 1 ), left = y.grob, bottom = x.grob)))
```

```
#Figure 4B

plot6<-ggplot() +
geom_density(aes(x=rel_position, weight=prop, color=time), data=temp_data1[DEG=="DEG",], adjust=0.4) +
scale_color_manual(name="Processing time",values=c("#00379c","#f04177"))+
scale_x_continuous(name="Relative transcript position (5' - 3')", breaks=c(0, 0.5, 1))+
theme_classic()+
theme(
text=element_text(size=font_size, color="black", face="plain"),
axis.text = element_text( size=font_size, colour = 'black', face="plain"),
axis.title = element_blank(),
plot.title = element_text(size=font_size),
legend.position="none",
axis.line.x = element_line(size=line_size),
axis.line.y = element_line(size=line_size),
axis.ticks = element_line(size=line_size))


plot7<-ggplot() +
geom_density(aes(x=rel_position, weight=prop,  color=time), data=temp_data2[DEG=="DEG",], adjust=0.4) +
scale_color_manual(name="Processing time",values=c("#00379c","#f04177"))+
scale_x_continuous(name="Relative transcript position (5' - 3')", breaks=c(0, 0.5, 1))+
theme_classic()+
theme(
text=element_text(size=font_size, color="black", face="plain"),
axis.text = element_text( size=font_size, colour = 'black', face="plain"),
axis.title = element_blank(),
plot.title = element_text(size=font_size),
legend.position="none",
axis.line.x = element_line(size=line_size),
axis.line.y = element_line(size=line_size),
axis.ticks = element_line(size=line_size))


plot8<-ggplot() +
geom_density(aes(x=rel_position, weight=prop, color=time), data=temp_data3[DEG=="DEG",], adjust=0.4) +
scale_color_manual(name="Processing time",values=c("#00379c","#f04177"))+
scale_x_continuous(name="Relative transcript position (5' - 3')", breaks=c(0, 0.5, 1))+
theme_classic()+
theme(
text=element_text(size=font_size, color="black", face="plain"),
axis.text = element_text( size=font_size, colour = 'black', face="plain"),
axis.title = element_blank(),
plot.title = element_text(size=font_size),
legend.position="none",
axis.line.x = element_line(size=line_size),
axis.line.y = element_line(size=line_size),
axis.ticks = element_line(size=line_size))


plot9<-ggplot() +
geom_density(aes(x=rel_position, weight=prop,  color=time), data=temp_data4[DEG=="DEG",], adjust=0.4) +
scale_color_manual(name="Processing time",values=c("#00379c","#f04177"))+
scale_x_continuous(name="Relative transcript position (5' - 3')", breaks=c(0, 0.5, 1))+
theme_classic()+
theme(
text=element_text(size=font_size, color="black", face="plain"),
axis.text = element_text( size=font_size, colour = 'black', face="plain"),
axis.title = element_blank(),
plot.title = element_text(size=font_size),
legend.position="none",
axis.line.x = element_line(size=line_size),
axis.line.y = element_line(size=line_size),
axis.ticks = element_line(size=line_size))


plot10<-ggplot() +
geom_density(aes(x=rel_position, weight=prop, color=time), data=temp_data5[DEG=="DEG",], adjust=0.4) +
scale_color_manual(name="Processing time",values=c("#00379c","#f04177"))+
scale_x_continuous(name="Relative transcript position (5' - 3')", breaks=c(0, 0.5, 1))+
theme_classic()+
theme(
text=element_text(size=font_size, color="black", face="plain"),
axis.text = element_text( size=font_size, colour = 'black', face="plain"),
axis.title = element_blank(),
plot.title = element_text(size=font_size),
legend.position="bottom",
legend.key.size = unit(0.2, 'cm'),
axis.line.x = element_line(size=line_size),
axis.line.y = element_line(size=line_size),
axis.ticks = element_line(size=line_size))

legend<-get_legend(plot10)

plot10<-ggplot() +
geom_density(aes(x=rel_position, weight=prop, color=time), data=temp_data5[DEG=="DEG",], adjust=0.4) +
scale_color_manual(name="Processing time",values=c("#00379c","#f04177"))+
scale_x_continuous(name="Relative transcript position (5' - 3')", breaks=c(0, 0.5, 1))+
theme_classic()+
theme(
text=element_text(size=font_size, color="black", face="plain"),
axis.text = element_text( size=font_size, colour = 'black', face="plain"),
axis.title = element_blank(),
plot.title = element_text(size=font_size),
legend.position="none",
axis.line.x = element_line(size=line_size),
axis.line.y = element_line(size=line_size),
axis.ticks = element_line(size=line_size))

y.grob <- textGrob("Density", gp=gpar(fontface="plain", col="black", fontsize=font_size), rot=90)
x.grob <- textGrob("Relative transcript position (5' - 3')", gp=gpar(fontface="plain", col="black", fontsize=font_size))

plot_b<-plot_grid( grid.arrange(arrangeGrob(plot_grid( plot6, plot7, plot8, plot9, plot10 ,nrow = 1 ), left = y.grob, bottom = x.grob)), legend, nrow = 2, ncol=1 ,rel_heights = c(1, .2))
```

```
#Figure 4C
ks_test_GSC22336_GSC22340_ks_test_GSC22336_GSC22335<-merge(ks_test_GSC22335_GSC22336,ks_test_GSC22336_GSC22340,by='gene_id', all=FALSE)
ks_test_GSC22336_GSC22340_ks_test_GSC22336_GSC22335$delta_D<-ks_test_GSC22336_GSC22340_ks_test_GSC22336_GSC22335$statistic.x  - ks_test_GSC22336_GSC22340_ks_test_GSC22336_GSC22335$statistic.y

ks_test_GSC22342_GSC22346_ks_test_GSC22341_GSC22342<-merge(ks_test_GSC22341_GSC22342,ks_test_GSC22342_GSC22346,by='gene_id', all=FALSE)
ks_test_GSC22342_GSC22346_ks_test_GSC22341_GSC22342$delta_D<-ks_test_GSC22342_GSC22346_ks_test_GSC22341_GSC22342$statistic.x  - ks_test_GSC22342_GSC22346_ks_test_GSC22341_GSC22342$statistic.y

ks_test_GSC22348_GSC22352_ks_test_GSC22347_GSC22348<-merge(ks_test_GSC22347_GSC22348,ks_test_GSC22348_GSC22352,by='gene_id', all=FALSE)
ks_test_GSC22348_GSC22352_ks_test_GSC22347_GSC22348$delta_D<-ks_test_GSC22348_GSC22352_ks_test_GSC22347_GSC22348$statistic.x  - ks_test_GSC22348_GSC22352_ks_test_GSC22347_GSC22348$statistic.y

ks_test_GSC22354_GSC22358_ks_test_GSC22353_GSC22354<-merge(ks_test_GSC22353_GSC22354,ks_test_GSC22354_GSC22358,by='gene_id', all=FALSE)
ks_test_GSC22354_GSC22358_ks_test_GSC22353_GSC22354$delta_D<-ks_test_GSC22354_GSC22358_ks_test_GSC22353_GSC22354$statistic.x  - ks_test_GSC22354_GSC22358_ks_test_GSC22353_GSC22354$statistic.y

ks_test_GSC22360_GSC22364_ks_test_GSC22359_GSC22360<-merge(ks_test_GSC22359_GSC22360,ks_test_GSC22360_GSC22364,by='gene_id', all=FALSE)
ks_test_GSC22360_GSC22364_ks_test_GSC22359_GSC22360$delta_D<-ks_test_GSC22360_GSC22364_ks_test_GSC22359_GSC22360$statistic.x  - ks_test_GSC22360_GSC22364_ks_test_GSC22359_GSC22360$statistic.y


ks_test_GSC22336_GSC22340_ks_test_GSC22336_GSC22335<-ks_test_GSC22336_GSC22340_ks_test_GSC22336_GSC22335[,heifer:=9108]
ks_test_GSC22342_GSC22346_ks_test_GSC22341_GSC22342<-ks_test_GSC22342_GSC22346_ks_test_GSC22341_GSC22342[,heifer:=9181]
ks_test_GSC22348_GSC22352_ks_test_GSC22347_GSC22348<-ks_test_GSC22348_GSC22352_ks_test_GSC22347_GSC22348[,heifer:=9135]
ks_test_GSC22354_GSC22358_ks_test_GSC22353_GSC22354<-ks_test_GSC22354_GSC22358_ks_test_GSC22353_GSC22354[,heifer:=9106]
ks_test_GSC22360_GSC22364_ks_test_GSC22359_GSC22360<-ks_test_GSC22360_GSC22364_ks_test_GSC22359_GSC22360[,heifer:=9129]

for_plotting<-do.call('rbind', list(ks_test_GSC22336_GSC22340_ks_test_GSC22336_GSC22335,ks_test_GSC22342_GSC22346_ks_test_GSC22341_GSC22342,ks_test_GSC22348_GSC22352_ks_test_GSC22347_GSC22348,ks_test_GSC22354_GSC22358_ks_test_GSC22353_GSC22354,ks_test_GSC22360_GSC22364_ks_test_GSC22359_GSC22360))

for_plotting[,DEG:=ifelse(gene_id %in% genes_lower_abundance_24hvs1h, "down_24h", ifelse(gene_id %in% genes_greater_abundance_24hvs1h,"up_24h","noDEG"))]

for_plotting[, ':=' (average_deltaD = mean(delta_D),count = .N) , by = gene_id]
for_plotting <- for_plotting[ count>4]
for_plotting <- for_plotting[order(for_plotting$average_deltaD),]
for_plotting <- for_plotting[, gene_id:=factor(gene_id, levels=c(unique(for_plotting[, gene_id])))]
for_plotting <- for_plotting[, heifer:=factor(heifer)]
for_plotting<-na.omit(for_plotting)


for_plotting<-for_plotting[order(heifer,DEG,delta_D),]

for_plotting_9106<-for_plotting[heifer==9106] %>%
group_by(DEG) %>%
mutate(row = row_number())

for_plotting_9108<-for_plotting[heifer==9108] %>%
group_by(DEG) %>%
mutate(row = row_number())

for_plotting_9129<-for_plotting[heifer==9129] %>%
group_by(DEG) %>%
mutate(row = row_number())

for_plotting_9135<-for_plotting[heifer==9135] %>%
group_by(DEG) %>%
mutate(row = row_number())

for_plotting_9181<-for_plotting[heifer==9181] %>%
group_by(DEG) %>%
mutate(row = row_number())

plot11<-ggplot( )+
geom_point(data=for_plotting_9106, aes(x=row, y=delta_D),size=0.1, shape=20)+
facet_wrap(~DEG,scales = "free_x" ,labeller=labeller(DEG = c(down_24h = "24hr < 1hr", noDEG =  paste("24hr", "~", "1hr"), up_24h="24hr > 1hr")))+
scale_x_discrete(name='Genes')+
scale_y_continuous(name='Delta D')+
#coord_flip()+
theme_classic(base_size = font_size)+
theme(
axis.title=element_blank(),
axis.text.x=element_blank(),
axis.ticks.x=element_blank(),
axis.text.y=element_text(size=font_size, color="black"),
plot.margin = unit(c(0.2, 0, 0,0), "cm"),
axis.line.x = element_line(size=line_size),
axis.line.y = element_line(size=line_size),
strip.background = element_blank()
)

plot12<-ggplot( )+
geom_point(data=for_plotting_9108, aes(x=row, y=delta_D),size=0.1, shape=20)+
facet_wrap(~DEG,scales = "free_x" ,labeller=labeller(DEG = c(down_24h = "24hr < 1hr", noDEG =  paste("24hr", "~", "1hr"), up_24h="24hr > 1hr")))+
scale_x_discrete(name='Genes')+
scale_y_continuous(name='Delta D')+
#coord_flip()+
theme_classic(base_size = font_size)+
theme(
axis.title=element_blank(),
axis.text.x=element_blank(),
axis.ticks.x=element_blank(),
axis.text.y=element_text(size=font_size, color="black"),
plot.margin = unit(c(0.2, 0, 0,0), "cm"),
axis.line.x = element_line(size=line_size),
axis.line.y = element_line(size=line_size),
strip.background = element_blank()
)

plot13<-ggplot( )+
geom_point(data=for_plotting_9129, aes(x=row, y=delta_D),size=0.1, shape=20)+
facet_wrap(~DEG,scales = "free_x" ,labeller=labeller(DEG = c(down_24h = "24hr < 1hr", noDEG =  paste("24hr", "~", "1hr"), up_24h="24hr > 1hr")))+
scale_x_discrete(name='Genes')+
scale_y_continuous(name='Delta D')+
#coord_flip()+
theme_classic(base_size = font_size)+
theme(
axis.title=element_blank(),
axis.text.x=element_blank(),
axis.ticks.x=element_blank(),
axis.text.y=element_text(size=font_size, color="black"),
plot.margin = unit(c(0.2, 0, 0,0), "cm"),
axis.line.x = element_line(size=line_size),
axis.line.y = element_line(size=line_size),
strip.background = element_blank()
)

plot14<-ggplot( )+
geom_point(data=for_plotting_9135, aes(x=row, y=delta_D),size=0.1, shape=20)+
facet_wrap(~DEG,scales = "free_x" ,labeller=labeller(DEG = c(down_24h = "24hr < 1hr", noDEG =  paste("24hr", "~", "1hr"), up_24h="24hr > 1hr")))+
scale_x_discrete(name='Genes')+
scale_y_continuous(name='Delta D')+
#coord_flip()+
theme_classic(base_size = font_size)+
theme(
axis.title=element_blank(),
axis.text.x=element_blank(),
axis.ticks.x=element_blank(),
axis.text.y=element_text(size=font_size, color="black"),
plot.margin = unit(c(0.2, 0, 0,0), "cm"),
axis.line.x = element_line(size=line_size),
axis.line.y = element_line(size=line_size),
strip.background = element_blank()
)

plot15<-ggplot( )+
geom_point(data=for_plotting_9181, aes(x=row, y=delta_D),size=0.1, shape=20) +
facet_wrap(~DEG,scales = "free_x" ,labeller=labeller(DEG = c(down_24h = "24hr < 1hr", noDEG = paste("24hr", "~", "1hr"), up_24h="24hr > 1hr")))+
scale_x_discrete(name='Genes')+
scale_y_continuous(name='Delta D')+
#coord_flip()+
theme_classic(base_size = font_size)+
theme(
axis.title=element_blank(),
axis.text.x=element_blank(),
axis.ticks.x=element_blank(),
axis.text.y=element_text(size=font_size, color="black"),
plot.margin = unit(c(0.2, 0, 0,0), "cm"),
axis.line.x = element_line(size=line_size),
axis.line.y = element_line(size=line_size),
strip.background = element_blank()
)

y.grob <- textGrob("Delta D", gp=gpar(fontface="plain", col="black", fontsize=font_size), rot=90)
x.grob <- textGrob("Genes", gp=gpar(fontface="plain", col="black", fontsize=font_size))

plot_c<-grid.arrange(arrangeGrob(plot_grid(plot11,plot12, plot13,  plot14, plot15, nrow = 1,  align = 'v'), left = y.grob, bottom = x.grob))
```

```
plot_grid(plot_a,plot_b,plot_c,nrow = 3, rel_heights=c(1,1,0.8), labels = c('A', 'B', 'C'), label_fontface="plain", label_size=10)
```

##### Gene Ontology enrichment analysis

For this enrichment analysis, we use the genes for which we quantified transcript abundance as the background list.

```
all_genes<-data.frame( gene=genes_expressed, stringsAsFactors=FALSE )
rownames(all_genes)<-all_genes$gene
N_expressed_genes<-length(all_genes$gene)

gene.length<-gene.length[gene.length$ensembl_gene_id %in% all_genes$gene,]

annotation.genelength.biomart_vector<-gene.length$transcript_length
names(annotation.genelength.biomart_vector)<-gene.length$ensembl_gene_id

annotation.GO.BP.biomart<-annotation.GO.biomart[annotation.GO.biomart$namespace_1003=="biological_process", c(1,3)]
annotation.GO.BP.biomart<-annotation.GO.BP.biomart[annotation.GO.BP.biomart$ensembl_gene_id %in% rownames(all_genes),]
annotation.GO.MF.biomart<-annotation.GO.biomart[annotation.GO.biomart$namespace_1003=="molecular_function", c(1,3)]
annotation.GO.MF.biomart<-annotation.GO.MF.biomart[annotation.GO.MF.biomart$ensembl_gene_id %in% rownames(all_genes),]
```

##### Additional file 7

Gene ontology enrichment analysis of biological processes for the genes with greater transcript abundance in PWBCs cryopreserved after 24 hours of refrigeration post collection versus within one hour of collection.

```
test.genes<-data.frame(a=unique(overlap_24hvs1h[overlap_24hvs1h$result=="both" & overlap_24hvs1h$logFC>0,1] ), stringsAsFactors=FALSE)
all_genes_numeric<-as.integer(all_genes$gene %in%test.genes$a)
names(all_genes_numeric)<-all_genes$gene

N_sig_genes<-length(test.genes$a)

set.seed(9830)
pwf<-nullp(all_genes_numeric, bias.data=annotation.genelength.biomart_vector, plot.fit=FALSE ) 
GO_BP_Cats_up_24h<-goseq(pwf,gene2cat=annotation.GO.BP.biomart, method ="Sampling", repcnt = 5000, use_genes_without_cat=FALSE)
GO_BP_Cats_up_24h<-GO_BP_Cats_up_24h[GO_BP_Cats_up_24h$numDEInCat>3,]
GO_BP_Cats_up_24h$FWER<-p.adjust(GO_BP_Cats_up_24h$over_represented_pvalue, method ="holm")
GO_BP_Cats_up_24h<-GO_BP_Cats_up_24h[with(GO_BP_Cats_up_24h, order(FWER,over_represented_pvalue, -numDEInCat)), ]
#head(GO_BP_Cats_up_24h, n=20)

GO_BP_Cats_up_24h$fold_enrichment<-(GO_BP_Cats_up_24h$numDEInCat/N_sig_genes)/(GO_BP_Cats_up_24h$numInCat/N_expressed_genes)
annotation.GO.BP.biomart_testgenes<-annotation.GO.BP.biomart[annotation.GO.BP.biomart$ensembl_gene_id %in% test.genes$a, ]
GO_BP_Cats_up_24h<-merge(GO_BP_Cats_up_24h,annotation.GO.BP.biomart_testgenes, by.x="category", by.y="go_id", all.x=TRUE, all.y=FALSE)
GO_BP_Cats_up_24h<-merge(GO_BP_Cats_up_24h, annotation.ensembl.symbol, by.x="ensembl_gene_id", by.y="ensembl_gene_id", all=FALSE, all.x=TRUE, all.y=FALSE)
GO_BP_Cats_up_24h<-GO_BP_Cats_up_24h[with(GO_BP_Cats_up_24h, order(FWER,term)), ]
GO_BP_Cats_up_24h<-GO_BP_Cats_up_24h[GO_BP_Cats_up_24h$FWER<0.1,]
#write.table(GO_BP_Cats_up_24h, file= "/2021_11_29_GO_BP_Cats_up_24h.txt", append = FALSE, quote = FALSE, sep = "\t" ,row.names = FALSE)
```

##### Additional file 8

Gene ontology enrichment analysis of molecular functions for the genes with greater transcript abundance in PWBCs cryopreserved after 24 hours of refrigeration post collection versus within one hour of collection.

```
set.seed(9830)
pwf<-nullp(all_genes_numeric, bias.data=annotation.genelength.biomart_vector, plot.fit=FALSE ) 
GO_MF_Cats_up_24h<-goseq(pwf,gene2cat=annotation.GO.MF.biomart, method ="Sampling", repcnt = 5000, use_genes_without_cat=FALSE)
GO_MF_Cats_up_24h<-GO_MF_Cats_up_24h[GO_MF_Cats_up_24h$numDEInCat>3,]
GO_MF_Cats_up_24h$FWER<-p.adjust(GO_MF_Cats_up_24h$over_represented_pvalue, method ="fdr")
GO_MF_Cats_up_24h<-GO_MF_Cats_up_24h[with(GO_MF_Cats_up_24h, order(FWER,over_represented_pvalue, -numDEInCat)), ]
#head(GO_MF_Cats_up_24h, n=20)

GO_MF_Cats_up_24h$fold_enrichment<-(GO_MF_Cats_up_24h$numDEInCat/N_sig_genes)/(GO_MF_Cats_up_24h$numInCat/N_expressed_genes)
annotation.GO.MF.biomart_testgenes<-annotation.GO.MF.biomart[annotation.GO.MF.biomart$ensembl_gene_id %in% test.genes$a, ]
GO_MF_Cats_up_24h<-merge(GO_MF_Cats_up_24h,annotation.GO.MF.biomart_testgenes, by.x="category", by.y="go_id", all.x=TRUE, all.y=FALSE)
GO_MF_Cats_up_24h<-merge(GO_MF_Cats_up_24h, annotation.ensembl.symbol, by.x="ensembl_gene_id", by.y="ensembl_gene_id", all=FALSE, all.x=TRUE, all.y=FALSE)
GO_MF_Cats_up_24h<-GO_MF_Cats_up_24h[with(GO_MF_Cats_up_24h, order(FWER,term)), ]
GO_MF_Cats_up_24h<-GO_MF_Cats_up_24h[GO_MF_Cats_up_24h$FWER<0.1,]
#write.table(GO_MF_Cats_up_24h, file= "/2021_11_29_GO_MF_Cats_up_24h.txt", append = FALSE, quote = FALSE, sep = "\t" ,row.names = FALSE)
```

##### Additional file 9

Gene ontology enrichment analysis of biological processes for the genes with lower transcript abundance in PWBCs cryopreserved after 24 hours of refrigeration post collection versus within one hour of collection.

```
test.genes<-data.frame(a=unique(overlap_24hvs1h[overlap_24hvs1h$result=="both" & overlap_24hvs1h$logFC<0,1] ), stringsAsFactors=FALSE)
all_genes_numeric<-as.integer(all_genes$gene %in%test.genes$a)
names(all_genes_numeric)<-all_genes$gene

N_sig_genes<-length(test.genes$a)

set.seed(9830)
pwf<-nullp(all_genes_numeric, bias.data=annotation.genelength.biomart_vector, plot.fit=FALSE ) 
GO_BP_Cats_down_24h<-goseq(pwf,gene2cat=annotation.GO.BP.biomart, method ="Sampling", repcnt = 5000, use_genes_without_cat=FALSE)
GO_BP_Cats_down_24h<-GO_BP_Cats_down_24h[GO_BP_Cats_down_24h$numDEInCat>3,]
GO_BP_Cats_down_24h$FWER<-p.adjust(GO_BP_Cats_down_24h$over_represented_pvalue, method ="fdr")
GO_BP_Cats_down_24h<-GO_BP_Cats_down_24h[with(GO_BP_Cats_down_24h, order(FWER,over_represented_pvalue, -numDEInCat)), ]
#head(GO_BP_Cats_down_24h, n=20)

GO_BP_Cats_down_24h$fold_enrichment<-(GO_BP_Cats_down_24h$numDEInCat/N_sig_genes)/(GO_BP_Cats_down_24h$numInCat/N_expressed_genes)
annotation.GO.BP.biomart_testgenes<-annotation.GO.BP.biomart[annotation.GO.BP.biomart$ensembl_gene_id %in% test.genes$a, ]
GO_BP_Cats_down_24h<-merge(GO_BP_Cats_down_24h,annotation.GO.BP.biomart_testgenes, by.x="category", by.y="go_id", all.x=TRUE, all.y=FALSE)
GO_BP_Cats_down_24h<-merge(GO_BP_Cats_down_24h, annotation.ensembl.symbol, by.x="ensembl_gene_id", by.y="ensembl_gene_id", all=FALSE, all.x=TRUE, all.y=FALSE)
GO_BP_Cats_down_24h<-GO_BP_Cats_down_24h[with(GO_BP_Cats_down_24h, order(FWER,term)), ]
GO_BP_Cats_down_24h<-GO_BP_Cats_down_24h[GO_BP_Cats_down_24h$FWER<0.1,]
#write.table(GO_BP_Cats_down_24h, file= "/2021_11_29_GO_BP_Cats_down_24h.txt", append = FALSE, quote = FALSE, sep = "\t" ,row.names = FALSE)
```

##### Additional file 10

Gene ontology enrichment analysis of molecular functions for the genes with lower transcript abundance in PWBCs cryopreserved after 24 hours of refrigeration post collection versus within one hour of collection.

```
pwf<-nullp(all_genes_numeric, bias.data=annotation.genelength.biomart_vector, plot.fit=FALSE ) 
GO_MF_Cats_down_24h<-goseq(pwf,gene2cat=annotation.GO.MF.biomart, method ="Sampling", repcnt = 5000, use_genes_without_cat=FALSE)
GO_MF_Cats_down_24h<-GO_MF_Cats_down_24h[GO_MF_Cats_down_24h$numDEInCat>3,]
GO_MF_Cats_down_24h$FWER<-p.adjust(GO_MF_Cats_down_24h$over_represented_pvalue, method ="fdr")
GO_MF_Cats_down_24h<-GO_MF_Cats_down_24h[with(GO_MF_Cats_down_24h, order(FWER,over_represented_pvalue, -numDEInCat)), ]
#head(GO_MF_Cats_down_24h, n=20)

GO_MF_Cats_down_24h$fold_enrichment<-(GO_MF_Cats_down_24h$numDEInCat/N_sig_genes)/(GO_MF_Cats_down_24h$numInCat/N_expressed_genes)
annotation.GO.MF.biomart_testgenes<-annotation.GO.MF.biomart[annotation.GO.MF.biomart$ensembl_gene_id %in% test.genes$a, ]
GO_MF_Cats_down_24h<-merge(GO_MF_Cats_down_24h,annotation.GO.MF.biomart_testgenes, by.x="category", by.y="go_id", all.x=TRUE, all.y=FALSE)
GO_MF_Cats_down_24h<-merge(GO_MF_Cats_down_24h, annotation.ensembl.symbol, by.x="ensembl_gene_id", by.y="ensembl_gene_id", all=FALSE, all.x=TRUE, all.y=FALSE)
GO_MF_Cats_down_24h<-GO_MF_Cats_down_24h[with(GO_MF_Cats_down_24h, order(FWER,term)), ]
GO_MF_Cats_down_24h<-GO_MF_Cats_down_24h[GO_MF_Cats_down_24h$FWER<0.1,]
#write.table(GO_MF_Cats_down_24h, file= "/2021_11_29_GO_MF_Cats_down_24h.txt", append = FALSE, quote = FALSE, sep = "\t" ,row.names = FALSE)
```

##### Figure 5

Gene ontology categories enriched in the genes with greater transcript abundance at 24hr versus one hr. A. Biological processes and B. Molecular functions. To improve readability, only categories with more than five genes are displayed on the graphs, please, see Additional files 7 and 8 for a full list of categories and the associated genes with their annotation.

```
GO_BP_Cats_up_24h_plotting<-GO_BP_Cats_up_24h[!duplicated(GO_BP_Cats_up_24h[,2:9]),c(2:10)]
GO_BP_Cats_up_24h_plotting<-GO_BP_Cats_up_24h_plotting[GO_BP_Cats_up_24h_plotting$FWER <= 0.1 & GO_BP_Cats_up_24h_plotting$numDEInCat>5,]
GO_BP_Cats_up_24h_plotting<-GO_BP_Cats_up_24h_plotting[order(-GO_BP_Cats_up_24h_plotting$FWER),]
GO_BP_Cats_up_24h_plotting$term<-factor(GO_BP_Cats_up_24h_plotting$term, levels=GO_BP_Cats_up_24h_plotting$term)

point_size<-2

plot1<-ggplot(data=GO_BP_Cats_up_24h_plotting, aes(x=term, y=-log10(FWER)))+
  geom_point(aes(color=fold_enrichment, size=point_size))+
  scale_y_continuous(name=bquote(-Log[10](FWER)), limits=c(1,3))+
  coord_flip()+
  scale_colour_gradient(low = "#B8B8B8", high = "#080808", na.value = NA,limits = c(1, 23))+
  scale_x_discrete(name=NULL)+
  ggtitle("Biological Processes")+
  theme_classic(base_size = 10) + 
  theme(
    legend.position = "none",
    axis.text = element_text(color="black"),
    axis.title.y = element_blank(),
    axis.title.x = element_text(vjust=1,margin = margin(t = 0, r = 0, b = 0, l = 0, unit = "cm")),
    plot.margin=unit(c(-0.30,0,0,0), "null"),
    plot.title = element_text(size = 10,margin = margin(t=0.2, r=0, b=0.2, l=0, unit = "cm"))
  )


GO_MF_Cats_up_24h_plotting<-GO_MF_Cats_up_24h[!duplicated(GO_MF_Cats_up_24h[,2:9]),c(2:10)]
GO_MF_Cats_up_24h_plotting<-GO_MF_Cats_up_24h_plotting[GO_MF_Cats_up_24h_plotting$FWER <= 0.1 & GO_MF_Cats_up_24h_plotting$numDEInCat>5,]
GO_MF_Cats_up_24h_plotting<-GO_MF_Cats_up_24h_plotting[order(-GO_MF_Cats_up_24h_plotting$FWER),]
GO_MF_Cats_up_24h_plotting$term<-factor(GO_MF_Cats_up_24h_plotting$term, levels=GO_MF_Cats_up_24h_plotting$term)

plot2<-ggplot(data=GO_MF_Cats_up_24h_plotting, aes(x=term, y=-log10(FWER)),size=10)+
 geom_point(aes(color=fold_enrichment))+
  scale_y_continuous(name=bquote(-Log[10](FWER)), limits=c(1,3))+
  coord_flip()+
   scale_colour_gradient(low = "#B8B8B8", high = "#080808", na.value = NA,limits = c(1, 23))+
  ggtitle("Molecular Functions")+
  theme_classic(base_size = 10) + 
  theme(
    axis.text = element_text(color="black"),
    axis.title.y = element_blank(),
    legend.position = "bottom",
    legend.key.width=unit(1,"cm"),
    legend.key.height = unit(0.2,"cm")
  )

legend<-get_legend(plot2)


plot2<-ggplot(data=GO_MF_Cats_up_24h_plotting, aes(x=term, y=-log10(FWER)))+
 geom_point(aes(color=fold_enrichment, size=point_size))+
  scale_y_continuous(name=bquote(-Log[10](FWER)), limits=c(1,3))+
  coord_flip()+
   scale_colour_gradient(low = "#B8B8B8", high = "#080808", na.value = NA,limits = c(1, 23))+
  ggtitle("Molecular Functions")+
  theme_classic(base_size = 10) + 
  theme(
    legend.position = "none",
    axis.text = element_text(color="black"),
     axis.title.y = element_blank(),
    axis.title.x = element_text(vjust=1,margin = margin(t = 0, r = 0.3, b = 0, l = 0.2, unit = "cm")),
    plot.margin = unit(c(0, 0, 0,0), "cm"),
    plot.title = element_text(size = 10,margin = margin(t=0.2, r=0, b=0.2, l=0, unit = "cm"))
  )


grid.arrange(arrangeGrob(plot_grid(plot1, plot2, nrow=1, ncol=2, align='hv', axis='l', labels=c('A', 'B'),hjust=c(-4,-1), label_fontface="plain", label_size=10), legend, heights=c(1,0.2)))
```

##### Figure 6

Gene ontology categories enriched in the genes with lower transcript abundance at 24hr versus 1hr. A. Biological processes and B. Molecular functions. To improve readability, only categories with more than five genes are displayed on the graphs, please, see Additional files 9 and 10 for a full list of categories and the associated genes with their annotation.

```
GO_MF_Cats_down_24h_plotting<-GO_MF_Cats_down_24h[!duplicated(GO_MF_Cats_down_24h[,2:9]),c(2:10)]
GO_MF_Cats_down_24h_plotting<-GO_MF_Cats_down_24h_plotting[GO_MF_Cats_down_24h_plotting$FWER <= 0.1 & GO_MF_Cats_down_24h_plotting$numDEInCat>5,]
GO_MF_Cats_down_24h_plotting<-GO_MF_Cats_down_24h_plotting[order(-GO_MF_Cats_down_24h_plotting$FWER),]
GO_MF_Cats_down_24h_plotting$term<-factor(GO_MF_Cats_down_24h_plotting$term, levels=GO_MF_Cats_down_24h_plotting$term)

GO_BP_Cats_down_24h_plotting<-GO_BP_Cats_down_24h[!duplicated(GO_BP_Cats_down_24h[,2:9]),c(2:10)]
GO_BP_Cats_down_24h_plotting<-GO_BP_Cats_down_24h_plotting[GO_BP_Cats_down_24h_plotting$FWER <= 0.1 & GO_BP_Cats_down_24h_plotting$numDEInCat>5,]
GO_BP_Cats_down_24h_plotting<-GO_BP_Cats_down_24h_plotting[order(-GO_BP_Cats_down_24h_plotting$FWER),]
GO_BP_Cats_down_24h_plotting$term<-factor(GO_BP_Cats_down_24h_plotting$term, levels=GO_BP_Cats_down_24h_plotting$term)

plot3<-ggplot(data=GO_BP_Cats_down_24h_plotting, aes(x=term, y=-log10(FWER)))+
  geom_point(aes(color=fold_enrichment, size=point_size))+
  scale_x_discrete(label = function(x) stringr::str_trunc(x, 40))+
  coord_flip()+
  scale_colour_gradient(low = "#B8B8B8", high = "#080808", na.value = NA,limits = c(1, 23))+
  scale_y_continuous(name=bquote(-Log[10](FWER)), limits=c(1,3))+
  scale_x_discrete(name=NULL)+
  ggtitle("Biological Processes")+
  theme_classic(base_size = 10) + 
  theme(
    axis.text = element_text(color="black"),
    axis.title.y = element_blank(),
    axis.title.x = element_text(vjust=1,margin = margin(t = 0, r = 0.3, b = 0, l = 0.2, unit = "cm")),
    legend.position = "none",
    plot.margin=unit(c(-0.30,0,0,0), "null"),
    plot.title = element_text(size = 10, margin = margin(t=0.2, r=0, b=0.2, l=0, unit = "cm"))
  )


plot4<-ggplot(data=GO_MF_Cats_down_24h_plotting, aes(x=term, y=-log10(FWER)),size=10)+
  geom_point(aes(color=fold_enrichment ,size=point_size))+
  scale_y_continuous(name=bquote(-Log[10](FWER)), limits=c(1,3))+
  coord_flip()+
  scale_colour_gradient(name="Fold enrichment", low = "#B8B8B8", high = "#080808", na.value = NA,limits = c(1, 23))+
  scale_size_continuous(guide="none")+
  #ylim(c(1,3))+
  theme_classic(base_size = 10) + 
  theme(
    axis.text = element_text(color="black"),
    axis.title.y = element_blank(),
    legend.position = "bottom",
    legend.key.width=unit(1,"cm"),
    legend.key.height = unit(0.2,"cm")
  )

legend<-get_legend(plot4)

plot4<-ggplot(data=GO_MF_Cats_down_24h_plotting, aes(x=term, y=-log10(FWER)),size=10)+
  geom_point(aes(color=fold_enrichment ,size=point_size))+
  scale_y_continuous(name=bquote(-Log[10](FWER)), limits=c(1,3))+
  scale_x_discrete(label = function(x) stringr::str_trunc(x, 40))+
  coord_flip()+
  scale_colour_gradient(low = "#B8B8B8", high = "#080808", na.value = NA,limits = c(1, 23))+
  #ylim(c(1,3))+
  ggtitle("Molecular Functions")+
  theme_classic(base_size = 10) + 
  theme(
    axis.text = element_text(color="black"),
    axis.title.y = element_blank(),
    axis.title.x = element_text(vjust=1,margin = margin(t = 0, r = 0.3, b = 0, l = 0.2, unit = "cm")),
    legend.position = "none",
    plot.margin = unit(c(0, 0, 0,0), "cm"),
    plot.title = element_text(size = 7,margin = margin(t=0.2, r=0, b=0.2, l=0, unit = "cm"))
  )


grid.arrange(arrangeGrob(plot_grid(plot3, plot4, nrow=1, ncol=2, align='hv', axis='l', labels=c('A', 'B'),hjust=c(-4,-1), label_fontface="plain", label_size=8), legend, heights=c(1,0.1)))
```

##### Additional file 11

Differential transcript abundance for genes previously detected as potential biomarkers of heifer fertility. A. Genes with no significant variation of transcript abundance following three, six, eight or 24 hours of refrigeration post collection versus within one hour of collection. B. Genes with significantly differential transcript abundance following 24 hours of refrigeration post collection versus within one hour of collection.

```
#table(overlap_8hvs1h[(overlap_8hvs1h$Row.names %in% DEGs_pregnancy_outcome_past_publications$Ensembl_ID ),]$result)
#table(overlap_24hvs1h[(overlap_24hvs1h$Row.names %in% DEGs_pregnancy_outcome_past_publications$Ensembl_ID ),]$result)
#overlap_24hvs1h[(overlap_24hvs1h$Row.names %in% DEGs_pregnancy_outcome_past_publications$Ensembl_ID & overlap_24hvs1h$result =="both"),]$Row.names

overlap_3hvs1h_DEG_past_pub<-overlap_3hvs1h[(overlap_3hvs1h$Row.names %in% DEGs_pregnancy_outcome_past_publications$Ensembl_ID),]
overlap_6hvs1h_DEG_past_pub<-overlap_6hvs1h[(overlap_6hvs1h$Row.names %in% DEGs_pregnancy_outcome_past_publications$Ensembl_ID),]
overlap_8hvs1h_DEG_past_pub<-overlap_8hvs1h[(overlap_8hvs1h$Row.names %in% DEGs_pregnancy_outcome_past_publications$Ensembl_ID),]
overlap_24hvs1h_DEG_past_pub<-overlap_24hvs1h[(overlap_24hvs1h$Row.names %in% DEGs_pregnancy_outcome_past_publications$Ensembl_ID),]


overlap_3hvs1h_DEG_past_pub$time<-"3h"
overlap_6hvs1h_DEG_past_pub$time<-"6h"
overlap_8hvs1h_DEG_past_pub$time<-"8h"
overlap_24hvs1h_DEG_past_pub$time<-"24h"

forplotting_DEGs_past_pubs<-rbind(overlap_3hvs1h_DEG_past_pub,overlap_6hvs1h_DEG_past_pub,overlap_8hvs1h_DEG_past_pub,overlap_24hvs1h_DEG_past_pub)
forplotting_DEGs_past_pubs<-forplotting_DEGs_past_pubs[,c("Row.names","logFC", "time")]
forplotting_DEGs_past_pubs$time<-factor(forplotting_DEGs_past_pubs$time, levels=c("3h", "6h",  "8h", "24h" ))
forplotting_DEGs_past_pubs$color<-ifelse(forplotting_DEGs_past_pubs$Row.names %in% c("ENSBTAG00000010606" , "ENSBTAG00000052658"), "red", "black")
forplotting_DEGs_past_pubs$color<-factor(forplotting_DEGs_past_pubs$color, levels=c("black","red"))

forplotting_DEGs_past_pubs<-merge(forplotting_DEGs_past_pubs, annotation.ensembl.symbol, by.x="Row.names", by.y="ensembl_gene_id", all=FALSE)
forplotting_DEGs_past_pubs<-forplotting_DEGs_past_pubs[,c( "Row.names","logFC","time", "color" ,"external_gene_name")]


forplotting_DEGs_past_pubs$symbol<-ifelse(forplotting_DEGs_past_pubs$Row.names %in% c("ENSBTAG00000010606" ,"ENSBTAG00000052658") , forplotting_DEGs_past_pubs$external_gene_name , ifelse(forplotting_DEGs_past_pubs$logFC < -1.5, forplotting_DEGs_past_pubs$external_gene_name, ""))

forplotting_DEGs_past_pubs$symbol<-ifelse(forplotting_DEGs_past_pubs$Row.names %in% c("ENSBTAG00000010606" ,"ENSBTAG00000052658") , forplotting_DEGs_past_pubs$external_gene_name , "")


font_size<-10
plot1<-ggplot(data=forplotting_DEGs_past_pubs[forplotting_DEGs_past_pubs$symbol=="",], aes(x=time, y=logFC, group=Row.names,color="gray", label=symbol))+
geom_line()+
ggrepel::geom_text_repel(max.overlaps=100)+
scale_color_manual(name=NULL,values=c("lightgray"))+
scale_y_continuous(limits=c(-3,3))+
theme_classic(base_size=font_size)+
theme(
legend.position = "none",
axis.text=element_text(size=font_size, color="black"),
axis.title=element_text(size=font_size, color="black")
)

plot2<-ggplot(data=forplotting_DEGs_past_pubs[!(forplotting_DEGs_past_pubs$symbol==""),], aes(x=time, y=logFC, group=Row.names,color="red", label=symbol))+
geom_point()+
geom_line(linetype=3)+
scale_y_continuous(limits=c(-3,3))+
ggrepel::geom_text_repel(max.overlaps=100)+
theme_classic()+
theme(
legend.position = "none",
axis.text=element_text(size=font_size, color="black"),
axis.title=element_text(size=font_size, color="black")
)

plot_grid( plot1,plot2,nrow=1, align='h', labels=c("A","B"), label_fontface="plain")
```

#### sessionInfo

```
sessionInfo()
```

```
## R version 4.0.3 (2020-10-10)
## Platform: x86_64-pc-linux-gnu (64-bit)
## Running under: Ubuntu 20.04.3 LTS
## 
## Matrix products: default
## BLAS:   /usr/lib/x86_64-linux-gnu/blas/libblas.so.3.9.0
## LAPACK: /usr/lib/x86_64-linux-gnu/lapack/liblapack.so.3.9.0
## 
## locale:
##  [1] LC_CTYPE=en_US.UTF-8       LC_NUMERIC=C              
##  [3] LC_TIME=en_US.UTF-8        LC_COLLATE=en_US.UTF-8    
##  [5] LC_MONETARY=en_US.UTF-8    LC_MESSAGES=en_US.UTF-8   
##  [7] LC_PAPER=en_US.UTF-8       LC_NAME=C                 
##  [9] LC_ADDRESS=C               LC_TELEPHONE=C            
## [11] LC_MEASUREMENT=en_US.UTF-8 LC_IDENTIFICATION=C       
## 
## attached base packages:
##  [1] grid      parallel  stats4    stats     graphics  grDevices utils    
##  [8] datasets  methods   base     
## 
## other attached packages:
##  [1] gridExtra_2.3               kableExtra_1.3.4.9000      
##  [3] ggsignif_0.6.3              data.table_1.14.2          
##  [5] dgof_1.2                    Rsamtools_2.4.0            
##  [7] Biostrings_2.56.0           XVector_0.28.0             
##  [9] goseq_1.40.0                geneLenDataBase_1.24.0     
## [11] BiasedUrn_1.07              car_3.0-12                 
## [13] carData_3.0-4               glmmTMB_1.1.2.3            
## [15] ggpubr_0.4.0                betareg_3.1-4              
## [17] logspline_2.1.16            fitdistrplus_1.1-6         
## [19] predictmeans_1.0.6          lmeInfo_0.1.3              
## [21] lme4_1.1-27.1               Matrix_1.4-0               
## [23] reshape2_1.4.4              htmlwidgets_1.5.4          
## [25] forcats_0.5.1               stringr_1.4.0              
## [27] purrr_0.3.4                 readr_2.1.1                
## [29] tidyr_1.1.4                 tibble_3.1.6               
## [31] tidyverse_1.3.1             plotly_4.10.0              
## [33] emmeans_1.7.1-1             multcomp_1.4-17            
## [35] TH.data_1.1-0               MASS_7.3-54                
## [37] survival_3.2-13             mvtnorm_1.1-3              
## [39] nlme_3.1-153                circlize_0.4.13            
## [41] flashClust_1.01-2           ComplexHeatmap_2.4.3       
## [43] VennDiagram_1.7.1           futile.logger_1.4.3        
## [45] DEXSeq_1.34.1               RColorBrewer_1.1-2         
## [47] AnnotationDbi_1.50.3        BiocParallel_1.22.0        
## [49] readxl_1.3.1                DESeq2_1.28.1              
## [51] SummarizedExperiment_1.18.2 DelayedArray_0.14.1        
## [53] matrixStats_0.61.0          Biobase_2.48.0             
## [55] GenomicRanges_1.40.0        GenomeInfoDb_1.24.2        
## [57] IRanges_2.22.2              S4Vectors_0.26.1           
## [59] BiocGenerics_0.34.0         GGally_2.1.2               
## [61] cowplot_1.1.1               edgeR_3.30.3               
## [63] limma_3.44.3                ggplot2_3.3.5              
## [65] dplyr_1.0.7                
## 
## loaded via a namespace (and not attached):
##   [1] utf8_1.2.2               tidyselect_1.1.1         RSQLite_2.2.9           
##   [4] munsell_0.5.0            codetools_0.2-18         statmod_1.4.36          
##   [7] withr_2.4.3              colorspace_2.0-2         highr_0.9               
##  [10] knitr_1.37               rstudioapi_0.13          labeling_0.4.2          
##  [13] GenomeInfoDbData_1.2.3   hwriter_1.3.2            farver_2.1.0            
##  [16] bit64_4.0.5              coda_0.19-4              vctrs_0.3.8             
##  [19] generics_0.1.1           lambda.r_1.2.4           xfun_0.29               
##  [22] BiocFileCache_1.12.1     R6_2.5.1                 clue_0.3-60             
##  [25] locfit_1.5-9.4           flexmix_2.3-17           bitops_1.0-7            
##  [28] cachem_1.0.6             reshape_0.8.8            assertthat_0.2.1        
##  [31] scales_1.1.1             nnet_7.3-16              gtable_0.3.0            
##  [34] sandwich_3.0-1           rlang_0.4.12             genefilter_1.70.0       
##  [37] systemfonts_1.0.3        GlobalOptions_0.1.2      splines_4.0.3           
##  [40] rtracklayer_1.48.0       TMB_1.7.22               rstatix_0.7.0           
##  [43] lazyeval_0.2.2           broom_0.7.10             abind_1.4-5             
##  [46] yaml_2.2.1               modelr_0.1.8             GenomicFeatures_1.40.1  
##  [49] backports_1.4.1          tools_4.0.3              ellipsis_0.3.2          
##  [52] jquerylib_0.1.4          Rcpp_1.0.7               plyr_1.8.6              
##  [55] progress_1.2.2           zlibbioc_1.34.0          RCurl_1.98-1.5          
##  [58] prettyunits_1.1.1        openssl_1.4.6            GetoptLong_1.0.5        
##  [61] zoo_1.8-9                ggrepel_0.9.1            haven_2.4.3             
##  [64] cluster_2.1.2            fs_1.5.2                 magrittr_2.0.1          
##  [67] futile.options_1.0.1     lmtest_0.9-39            reprex_2.0.1            
##  [70] hms_1.1.1                evaluate_0.14            xtable_1.8-4            
##  [73] pbkrtest_0.5.1           XML_3.99-0.8             shape_1.4.6             
##  [76] compiler_4.0.3           biomaRt_2.44.4           crayon_1.4.2            
##  [79] minqa_1.2.4              htmltools_0.5.2          mgcv_1.8-38             
##  [82] tzdb_0.2.0               Formula_1.2-4            libcoin_1.0-9           
##  [85] geneplotter_1.66.0       lubridate_1.8.0          DBI_1.1.2               
##  [88] formatR_1.11             dbplyr_2.1.1             rappdirs_0.3.3          
##  [91] boot_1.3-28              cli_3.1.0                pkgconfig_2.0.3         
##  [94] GenomicAlignments_1.24.0 coin_1.4-2               numDeriv_2016.8-1.1     
##  [97] xml2_1.3.3               svglite_2.0.0            annotate_1.66.0         
## [100] bslib_0.3.1              webshot_0.5.2            estimability_1.3        
## [103] rvest_1.0.2              digest_0.6.29            rmarkdown_2.11          
## [106] cellranger_1.1.0         curl_4.3.2               modeltools_0.2-23       
## [109] rjson_0.2.20             nloptr_1.2.2.3           lifecycle_1.0.1         
## [112] jsonlite_1.7.2           viridisLite_0.4.0        askpass_1.1             
## [115] fansi_0.5.0              pillar_1.6.4             lattice_0.20-45         
## [118] GO.db_3.11.4             fastmap_1.1.0            httr_1.4.2              
## [121] glue_1.6.0               png_0.1-7                bit_4.0.4               
## [124] stringi_1.7.6            sass_0.4.0               blob_1.2.2              
## [127] memoise_2.0.1
```
